# Estimating clade-specific diversification rates and palaeodiversity dynamics from reconstructed phylogenies

**DOI:** 10.1101/2022.05.10.490920

**Authors:** Nathan Mazet, Hélène Morlon, Pierre-Henri Fabre, Fabien L. Condamine

## Abstract

Understanding palaeodiversity dynamics through time and space is a central goal of macroevolution. Estimating palaeodiversity dynamics has been historically addressed with fossil data because it directly reflects the past variations of biodiversity. Unfortunately, some groups or regions lack a good fossil record, and dated phylogenies can be useful to estimate diversification dynamics. Recent methodological developments have unlocked the possibility to investigate palaeodiversity dynamics by using phylogenetic birth-death models with non-homogeneous rates through time and across clades. One of them seems particularly promising to detect clades whose diversity has declined through time. However, empirical applications of the method have been hampered by the lack of a robust, accessible implementation of the whole procedure, therefore requiring users to conduct all the steps of the analysis by hand in a time-consuming and error-prone way.

Here we propose an automation of Morlon et al. (2011) clade-shift model with additional features accounting for recent developments and we implement it in the R package RPANDA. We also test the approach with simulations focusing on its ability to detect shifts of diversification and to infer palaeodiversity dynamics. Finally, we illustrate the automation by investigating the palaeodiversity dynamics of Cetacea, Vangidae, Parnassiinae, and Cycadales.

Simulations showed that we accurately detected shifts of diversification although false shift detections were higher for time-dependent diversification models with extinction. The median global error of palaeodiversity dynamics estimated with the automated model is low showing that the method can capture diversity declines. We detected shifts of diversification for three of the four empirical examples considered (Cetacea, Parnassiinae and Cycadales). Our analyses unveil a waxing-and-waning pattern due to a phase of negative net diversification rate embedded in the trees after isolating recent radiations.

Our work makes possible to easily apply non-homogeneous models of diversification in which rates can vary through time and across clades to reconstruct palaeodiversity dynamics. By doing so, we detected palaeodiversity declines among three of the four groups tested, highlighting that such periods of negative net diversification might be common. We discuss the extent to which this approach might provide reliable estimates of extinction rates and we provide guidelines for users.

## 1 Introduction

Estimating palaeodiversity dynamics over geological timescales is a challenging goal of macroevolution (Alroy et al. 2008; Quental and Marshall 2010; Morlon et al. 2011). This question is generally addressed with paleontological data because fossil records are direct witnesses of past diversity dynamics (Marx and Uhen 2010; Ezard et al. 2011). Unfortunately, the fossil record is not homogeneous over time, space, and groups (Close et al. 2020). Consequently, investigating evolutionary history of poorly fossilised groups relies on diversification analyses based on dated phylogenies (time-trees) mainly composed of extant species. Methodological developments in the field have focused on estimating the speciation rate (λ) and extinction rate (μ), which represent the pace at which lineages appear and go extinct, respectively, as well as the net diversification rate (denoted r) defined as the difference between them (Ricklefs 2007; Stadler 2013; Morlon 2014). A large variety of phylogenetic models allows testing evolutionary hypotheses on whether diversification rates are constant, vary over time (Rabosky and Lovette 2008; Morlon et al. 2010), across clades (Morlon et al. 2011; Rabosky et al. 2013; Maliet et al. 2019) or in association with species’ traits (Maddison et al. 2007; FitzJohn 2012; Beaulieu and O’Meara 2016) or with environmental changes (Condamine et al. 2013, Condamine, Rolland & Morlon 2019). However, early studies have shown that reconstructed dated phylogenies can display heterogeneity of diversification rates across clades (Rabosky et al. 2007).

Among the methods developed to detect this heterogeneity (Alfaro et al. 2009; Stadler and Bokma 2013; Rabosky et al. 2013; Maliet et al. 2019; Barido-Sottani et al. 2020; Maliet & Morlon 2022), the model developed by Morlon et al. (2011) considers heterogeneity in diversification rate by testing for shifts in rates of diversification at specific branching points in the phylogeny. The diversification of subclades identified *a priori* based on taxonomy or ecological traits, and the diversification of the ancestral group leading to these clades, hereafter called the backbone, can be analysed separately. The main advantage of this method is that each part of a phylogeny (subclades and backbone) can follow a different birth-death model, offering the opportunity to test biological hypotheses that create heterogeneity of diversification rate and thus reconstruct palaeodiversity dynamics for each part of the tree (Nee et al. 1992; Morlon et al. 2011; Morlon 2014). Recent advances have improved palaeodiversity estimation with a probabilistic approach providing confidence intervals of palaeodiversity values (Billaud et al. 2020). Subclades might experience recent adaptive radiations due to the acquisition of key innovations (Jønsson et al. 2012; Drummond et al. 2012) while the backbone tree might display a different dynamic such as a waxing-waning pattern of diversification. The resulting heterogeneity of diversification rates cannot be detected in clade-homogeneous birth-death models because recently radiating clades can overshadow the estimation of extinction in other parts of the tree, and likely hamper the inference of a negative net diversification rate (r < 0, μ > λ) that may explain such a period of diversity decline (Morlon et al. 2011). The waxing-waning pattern detected by Morlon et al. (2011) was promising, yet the approach developed in that paper, part of which was made accessible in the R package RPANDA (Morlon et al. 2016), has been applied to only a few empirical cases since it was proposed (Bacon et al. 2013; Legendre and Condamine 2018; Billaud et al. 2020; Chazot et al. 2021) because of the lack of automation.

The method developed by Morlon et al. (2011) requires manually isolating subclades and backbones, accounting for all pruned speciation events, applying diversification models, and testing all backbone(s)/subclade(s) combinations by hand. This is time-consuming, error-prone, and makes the method underutilized. In addition to making this method more automatic and user-friendly, several aspects of its performance remain to be tested. It was shown that the method recovers unbiased rates of diversification and can detect periods of diversity decline (Morlon et al. 2011), but its ability to detect clade-specific shifts of diversification remains untested and can make the palaeodiversity dynamic challenging to recover (Burin et al. 2019). More generally, several studies have questioned our ability to estimate extinction rate from molecular phylogenies (Rabosky 2010, 2016; Louca and Pennell 2020; but see Nee et al. 1994; Morlon 2014; Beaulieu and O’Meara 2015). Historically, Nee et al. (1994) have demonstrated how to estimate constant extinction rate from molecular-dated phylogenies with extant species only. Some studies relaxed the assumption of constant rates (Rabosky and Lovette 2008, Stadler 2011a), but these models are homogeneous across clades. By being non-homogeneous across clades, the model of Morlon et al. (2011) brings better extinction rate estimates by revealing ancient periods of palaeodiversity decline (Stadler 2011b) in line with the fossil record (Marx and Uhen 2010). This model opens the possibility to investigate palaeodiversity dynamics for groups without good fossil records.

Here, we automatize the approach developed by Morlon el al. (2011) and implement it in the R package RPANDA (Morlon et al. 2016). In addition, we provide several advances compared to the original method. In particular, we implement the possibility to test for a shift at the crown age of subclades, whereas the original version of the method tested shifts at the stem age of subclades (Morlon et al. 2011; Billaud et al. 2020). This rationale is motivated by the idea that clade radiations – may it be adaptive or simply rapid compared to the rest of the tree – are characterized by long stem branches at the crown of subclades due to extinction, with fast speciation starting at the crown (Reznick and Ricklefs 2009, Budd and Mann 2020). We provide functions to calculate rates and diversity dynamics through time, including recent developments from Billaud et al. (2020) with new functions using automation outputs. Using simulations, we assess the performance of the approach to detect shifts and to infer palaeodiversity dynamics under various diversification scenarios. We further illustrate the approach empirically by investigating the evolutionary history of four groups with well-sampled molecular-dated phylogenies (Cetacea, Vangidae, Parnassiinae and Cycadales), which have been previously analysed with other methods. We finally discuss both simulations and empirical cases to provide guidelines and recommendations when using this approach.

## 2 Methods

### 2.1 The clade-dependent time-dependent diversification model of Morlon et al. (2011)

Morlon et al. (2011) developed an exact analytical expression of a birth-death model of cladogenesis that estimates rate variation through time while accounting for undersampling thanks to a sampling fraction (denoted by *f*, *f*=1 if all described species are included in the phylogeny). The expression is defined backwards in time such that values for λ and μ for time-dependent models correspond to rate values at present (*t*=0) and α and β are their dependency parameters, respectively. Then, the assumed changes of diversification rates occur at specific branching points (nodes) of the reconstructed phylogeny, meaning that potential rate shifts on unobserved (extinct or unsampled) lineages are ignored (Moore et al. 2016; Barido-Sottani et al. 2020). Under such assumptions, their analytical expression holds and in particular their expressions of the probability that a lineage alive at time *t* has no descendant in the sample at *t*=0 (Φ(*t*)), and the probability that a lineage alive at time *t* leaves exactly one descendant lineage at time *s* < *t* (Ψ(*s*, *t*), *i.e.,* the probability of survival from *t* to *s*, see Appendix S8 and S9 from Morlon et al. 2011). These two expressions allow expressing the likelihood of the backbone phylogeny by including the speciation event corresponding to the subclade. The probability 1 - Φ(*t*_1_^cl^) that the subclade survives from this branch point *t*_1_^cl^ is then included in the likelihood of the subclade expressed with the same formula. Finally, the product of the likelihood of the subtree and the backbone phylogeny gives the global likelihood of the phylogeny while accounting for heterogeneity of diversification rates across clades. This global likelihood can be compared to the likelihood of a diversification model with a homogeneous model of diversification to see if the shift of diversification rate is supported. This can be extended to a phylogeny with multiple subclades following their own diversification rates and even shifts of diversification rates inside the backbone creating multiple backbones.

### 2.2 Automation

#### 2.2.1 Implementation in RPANDA

Few empirical studies have applied the approach using a manual and sequential approach (Morlon et al. 2011; Bacon et al. 2013; Billaud et al. 2020; Chazot et al. 2021). Each shift was selected one by one among subclades defined *a priori* until either the inclusion of a new shift was no longer statistically supported or shifts at all nodes were supported. Our automated approach still relies on the *a priori* definition of a limited number of subclades in which shifts may have occurred, but contrary to the sequential approach it compares all combinations of these potential shifts at the same time. The simplest way to make an *a priori* selection of subclades is to follow the approach of Morlon et al. (2011) by choosing taxonomic subclades such as families. Moreover, taxonomic databases often provide the information required to compute sampling fractions (*i.e.,* clade-level taxonomic diversity) that are required by phylogenetic diversification models. The tested subclades can also correspond to clades of species sharing key-innovations or colonisation events to test the role of these biological events on the clade’s diversification. Once subclades have been chosen *a priori*, the whole analysis is divided in five steps: (1) calculating sampling fractions for the subclades from the taxonomic database, (2) calculating all potential combinations of subclade(s) and corresponding backbone(s) from the set of preselected subclades, (3) applying a set of diversification models (see section 2.2.2) to each part (subclades and backbone) of each combination to calculate global likelihoods and the corrected Akaike Information Criterion (AICc) for each combination and comparing their statistical support, (4) computing the palaeodiversity dynamics with deterministic or probabilistic approaches, and (5) testing model adequacy by simulating phylogenetic trees using the best combination(s) of shifts to check whether simulated data are close to empirical ones. The whole procedure is illustrated in **Figure 1**. By default, two constraints are applied: (1) λ and μ must be strictly positive, and (2) r (λ - μ) must be positive (r > 0) at the beginning of the diversification history (otherwise the clade goes extinct immediately).

**Figure 1.**
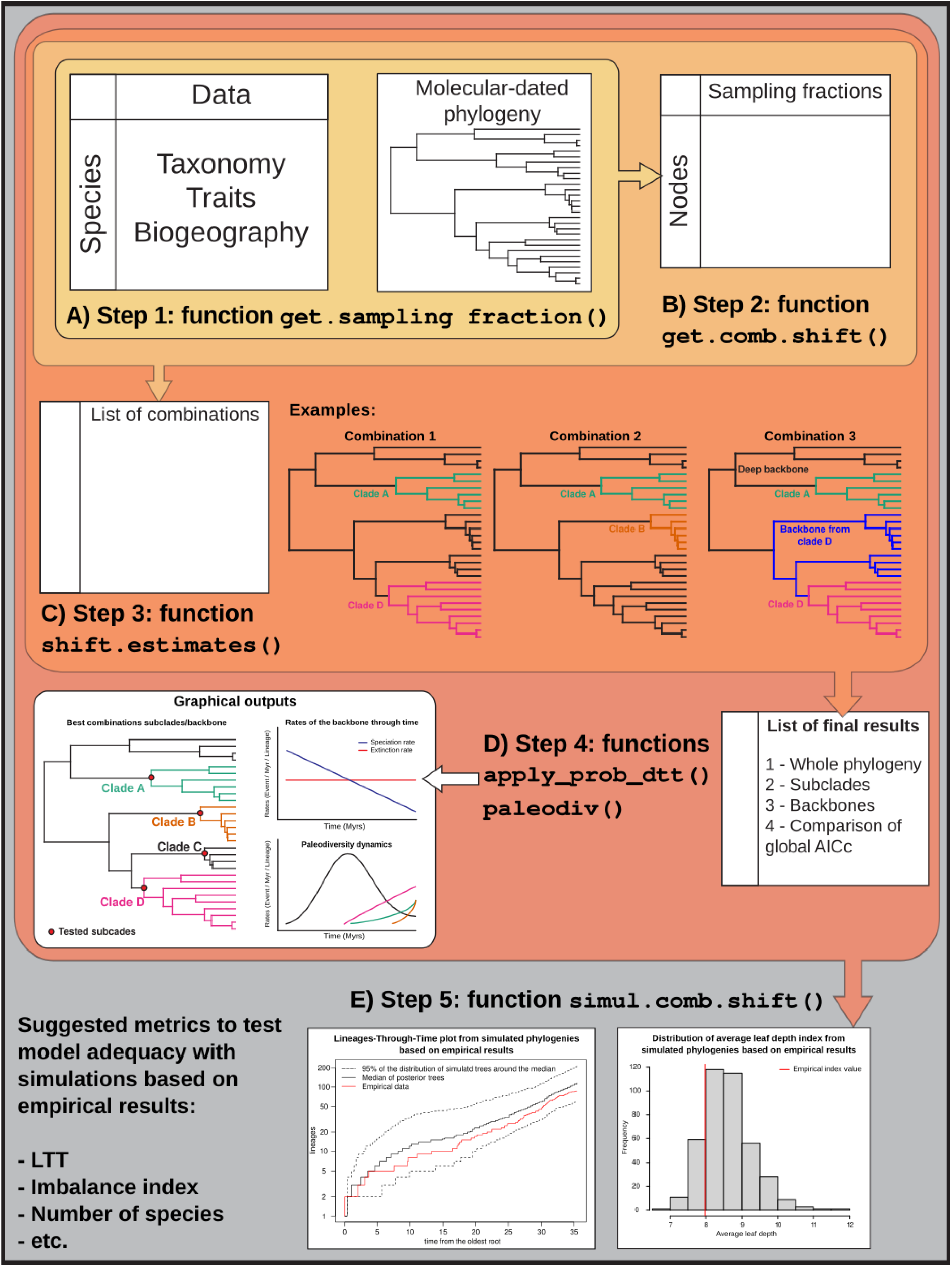
Conceptualization of the analytical pipeline for the automation of the method presented in Morlon et al. (2011). A) Step 1 calculates sampling fractions (*f*) from a database that contains all described species of the clade of interest and monophyletic subgroups (based on taxonomy or ecology); B) Step 2 calculates all the possible combinations of shifts (*e.g.* combination 1 has two shifts Clade A and D, the rest of the tree is included into the backbone tree); C) Step 3 applies a model comparison to select the best model for each part (subclades and backbone) of each combination. Then, it computes the likelihood and AICc of each combination, when using the best models. Finally, it selects the best supported combination. The final result is a list containing four elements: the model comparison for (1) the phylogeny analysed as a whole, (2) all subclades apart, (3) all backbones apart, and (4) the comparison of all combinations of heterogeneous diversification models based on their global AICc (see Appendix S4 for example). D) Step 4 computes palaeodiversity dynamics either with the deterministic (function *paleodiv*) or probabilistic approach (function *apply_prob_dtt*). E) Step 5 tests for model adequacy by simulating phylogenetic trees according to the best combination(s) of shifts to check whether empirical data matches with simulated data. This match can be evaluated with LTT plots, species richness or imbalance indices (see Appendix S4 for more details).

#### 2.2.2 Set of diversification models

Several features have been implemented to perform different analyses. The minimum clade size for applying diversification can be defined at each step and should be the same during the application of the whole pipeline (argument clade.size, default is 5 tips). The set of diversification models applied by default in the model comparison corresponds to the six following birth-death models: (1) constant pure-birth model (BCST), (2) constant birth-death model (BCST_DCST), (3) pure-birth model with exponential time-dependency such as λ(t) = λ_0_.e^α.t^ (BVAR), (4) birth-death model with exponential time-dependency for speciation rate and constant extinction rate (BVAR_DCST), (5) birth-death model with constant speciation rate and exponential time-dependency for extinction rate such as μ(t) = μ_0_.e^β.t^ (BCST_DVAR), and (6) birth-death model with exponential time-dependency for both speciation rate and extinction rate (BVAR_DVAR). A subset of these models can be used for all parts (subclades and backbones) with the argument models or only for subclades if, for example, extinction is not expected among subclades but only in the backbone(s) (see help files in RPANDA).

#### 2.2.3 Testing shifts at crown or stem age

Originally, the probability of the speciation event at the stem age was included in the likelihood of the backbone, while the probability that the stem of the subclade survives to the crown age was included in the likelihood of the subclade. Hence, the shift of diversification is located at the stem age. However, when phylogenies are usually analysed with diversification models, the stem branch is rarely included. Moreover, adaptive, or rapid radiations are expected to start at the crown age of the clade (Reznick and Ricklefs 2009). We implemented the possibility to test for shifts at crown ages of subclades by including the probability that the stem of the subclade survives to the crown age in the likelihood of the backbone. In other words, it means that the change in diversification rates synchronously occurs with the cladogenesis event as it could be expected under an adaptive radiation (Reznick and Ricklefs 2009). Users can choose to test for a shift at the stem age (backbone.option=“stem.shift”) as in Morlon et al. (2011), or at the crown age of the subclade (backbone.option=“crown.shift”, see Appendix S1 for more details).

#### 2.2.4 Multiple backbones

Shifts of diversification can also be nested creating combinations of shifts with more than one backbone (multiple backbones) in the phylogeny (Figure 1C, combination 3). We recommend to start investigating the heterogeneity of diversification with a simple backbone (no shift in the backbone) to first capture the main tendency of the backbone (option multi.backbone=F in function shift.estimates). This might be interesting if there are a lot of combinations with multiple backbones. Combinations with multiple backbones can then be analysed using the option multi.backbone=T or all combinations can be analysed at once with the option multi.backbone=“all”.

#### 2.2.5 Estimation of paleodiversity dynamics and constraints

Morlon et al. (2011) used a deterministic calculation of the palaeodiversity dynamics corresponding to the solution of the differential equation relating the time variation in the number of lineages to the rate values of the birth-death model (see Box 4 in Morlon 2014 for more details). Our automation also includes recent developments on palaeodiversity dynamic estimates (Billaud et al. 2020), a probabilistic approach to infer a confidence interval around the palaeodiversity curve. Functions that are already included in the R package RPANDA 2.0 (Morlon et al. 2016) have been slightly modified to be used with outputs of the shift.estimates() through a new function called apply_prob_dtt(). Palaeodiversity dynamics are computed independently for subclades and backbones. Two optional constraints are embedded in the optimisation process (in function shift.estimates): maximum rate value and maximum number of species estimates can be constrained to set an upper bound with a maximum value for each part of the combinations (argument rate.max and n.max, respectively). To constrain the number of species, we calculate the deterministic palaeodiversity curve during the optimisation process, and set the likelihood to -Inf if the maximum palaeodiversity value exceeds the constraint.

### 2.3 Simulations

We used simulations to test whether the approach successfully detects diversification shifts and correctly estimates palaeodiversity dynamics (**Figure 2**). We simulated four diversification scenarios following Billaud et al. (2020). All trees were simulated forward in time over 100 million years (Myrs, T_max_=100) with the R package TESS 2.1.0 (Höhna et al. 2016). For both waxing-waning models, α and β values were defined such that the diversity decline starts between the last third and the last quarter of the phylogenetic tree (*i.e.,* between 33 and 25 Myrs ago in our case). Rate values were selected such as they respect the following conditions:

1. Expanding diversity: constant pure-birth model (BCST) with *λ* ∊ [0.05; 0.25] and *μ*=0 (**Figure 2A**).
2. Saturing diversity: pure-birth model with exponential time-dependency (BVAR) with *λ_(t)_*=*λ_0_*exp(-*αt*) with *λ_0_* ∊ [0.05; 0.25] and *α* such that T_eq_ satisfies exp(-*α*T_eq_)=0.5 is in [T_max_2/3; T_max_3/4] and μ=0 (**Figure 2B**).
3. Waxing-waning (with decreasing speciation): birth-death model with exponential time-dependency for speciation rate and constant extinction rate (BVAR_DCST) with *λ_(t)_*=*λ_0_*exp(-*αt*) with *λ_0_* ∊ [0.2; 0.4] and *μ=λ_0_*/2 and *α* such that T_eq_ satisfies exp(*α*T_eq_)=*μ*/*λ* in [T_max_2/3; T_max_3/4] (**Figure 2C**).
4. Waxing-waning (with increasing extinction): birth-death model with constant extinction rate and exponential time-dependency for speciation rate (BCST_DVAR) *λ* ∊ [0.2; 0.4] and *μ*(*t*)=*μ_0_*exp(*βt*) with *μ_0_*=*λ*/2 and *β* such that T_eq_ satisfies exp(*β*T_eq_)=*λ*/*μ* in [T_max_ 2/3; T_max_3/4] (**Figure 2D**).

**Figure 2.**
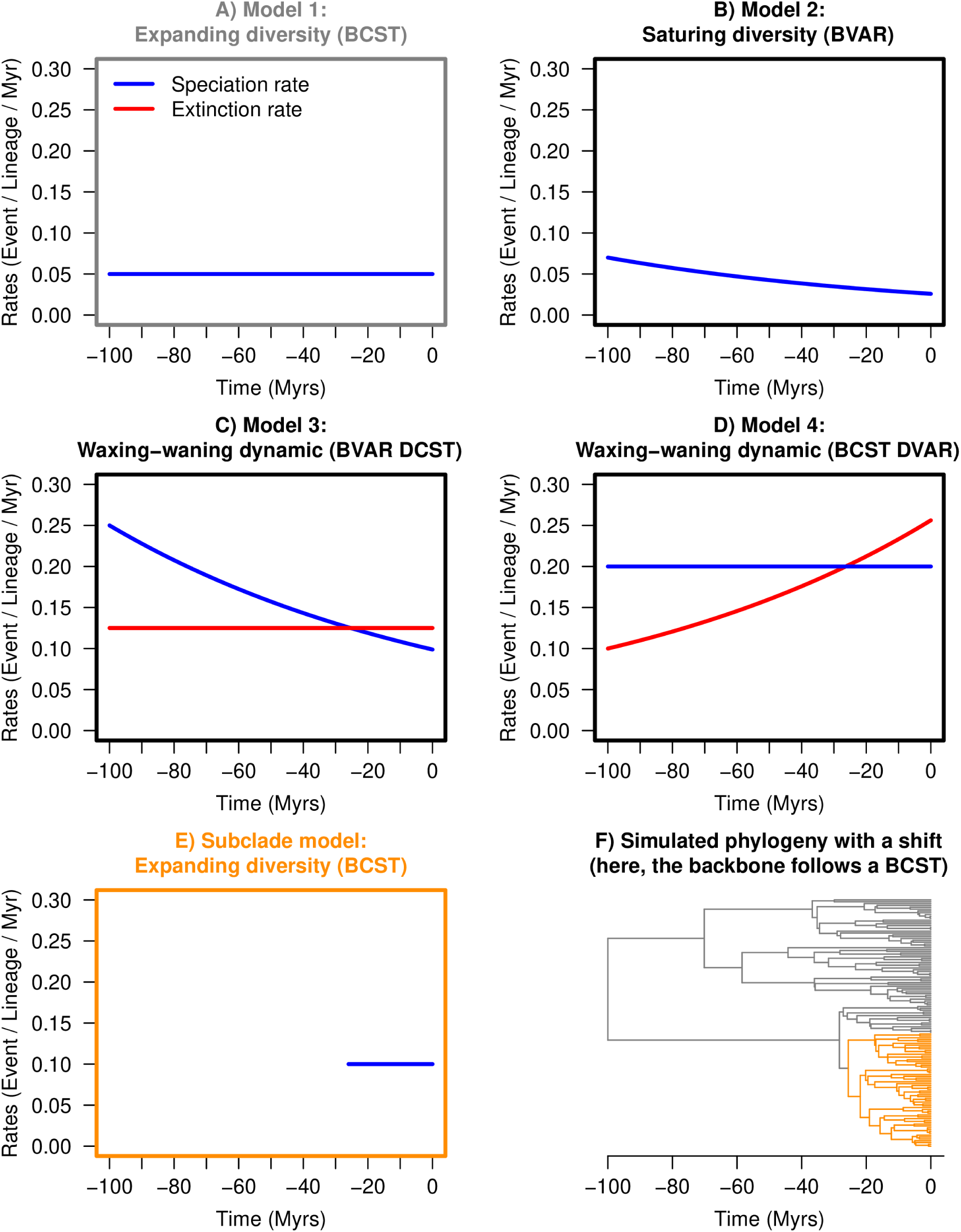
Simulation pipeline. A) to D) Diversification rates through time for the models used to simulate the backbone (grey). E) Diversification model used for the subclade (orange). F) Example of a simulated phylogeny with a rate shift shown in orange (the backbone is simulated with the Model 1).

We retained the simulated trees containing at least 40 species. We simulated the diversification shift by grafting a new simulated tree at a specific node defined as the more recent node among nodes between 33 and 25 Myrs ago (**Figure 2F**). This node must have a minimum of five extant descendant species to avoid applying diversification models on a species-poor clade when testing for the detection of a shift on raw phylogenies (*i.e.,* when there is no shift). For trees with a shift of diversification, the grafted tree follows a pure-birth model (BCST) with *λ*=0.1 (**Figure 2E**). This tree was grafted at its crown age and must contain at least 15 species. Because the shift was simulated at the crown age, the search for shift was also conducted at crown ages.

We quantified whether the model detected a shift of diversification while no shift was simulated (*i.e.,* false detection of shifts). Following Billaud et al. (2020), we also calculated the global error *D* between the estimated and the expected diversity curves, calculated with the deterministic approach for each simulation (see Appendix S3 for details). Simulations were set up with Snakemake 5.2.4 and R 3.6.3 using the R packages ape 5.5 (Paradis et al. 2019), phytools 0.7-80 (Revell 2012), TESS 2.1.0 (Höhna et al. 2016), phangorn 2.7.0 (Schliep 2011), and RPANDA 2.0 (Morlon et al. 2016, see Appendix S3 for more details).

Additionally, we tested whether extinction rate is properly detected and estimated in subclades. We performed a set of simulations (500 trees) following the same pipeline with Model 3 (BVAR_DCST) but with a diversification shift simulated by grafting a subclade following a constant birth-death model (BCST_DCST) instead of a pure-birth model (BCST). All the other criteria of simulations were the same, including the net diversification rate of the subclade kept at r=0.1 (λ=0.2 and μ=0.1).

### 2.4 Applications to empirical phylogenies

We tested the automated pipeline on four clades representing a large diversity of taxa to illustrate the potential of the approach. First, we applied the approach to the original example of Morlon et al. (2011), the cetacean phylogeny published by Steeman et al. (2009). To compare with previous results, we used the same taxonomy as in Morlon et al. (2011) with a global sampling fraction of 98% (87/89 species). We looked for diversification shifts for the four main families (Balaenopteridae, Delphinidae, Phocoenidae and Ziphiidae) and for the two parvorders (Odontoceti and Mysticeti). We tested shifts at the crown age (backbone.option=“crown.shift”) of subclades but also at the stem age (backbone.option=“stem.shift” see Appendix S2) as in Morlon et al. (2011) to compare both approaches.

Second, we tested a group without expected diversity decline, and studied the phylogeny of Vangidae (Jønsson et al. 2012), which is a small but fully sampled group (32/32 species). In their study, the authors detected two diversification shifts and concluded that this group has experienced adaptive radiation involving changes in beak size and body mass. We thus tested these hypotheses.

Third, we applied the approach to Parnassiinae, a subfamily of swallowtail butterflies (Condamine et al. 2018) with a good sampling fraction (96.6%, 85/88 species). Shifts of diversification can also come from ecological changes such as colonisation of new environments (Yoder et al. 2010) or host-plant shifts (Ehrlich and Raven 1964). We tested hypotheses related to host-plant shifts, Himalayan Mountain colonisations, and taxonomic groups.

Finally, we investigated the diversification history of Cycadales (Nagalingum et al. 2011). The sampling fraction of this group is equal to 72.5% (231/327 species, IUCN 2021). This group originated 275 Myr ago and almost all extant species emerge from recent radiations during the Miocene (Condamine et al. 2015). We tested shifts for all genera with more than five species described except *Dioon*, which is the least sampled (8/13 species). To account for uncertainty in divergence time estimates, we applied the approach to posterior trees and calculated the corresponding range of the palaeodiversity dynamics with the deterministic approach. This allows measuring variation of palaeodiversity estimates on a large panel of posterior trees and therefore accounting for uncertainties on divergence time estimates. We also calculated the palaeodiversity dynamics with the probabilistic approach to account for uncertainty of diversity estimates (Billaud et al. 2020).

For all empirical cases, we did not use the BVAR_DVAR model because it often produced unrealistic estimates and has already been discussed as problematic (Burin et al. 2019). We also detailed an example with the BVAR_DVAR model in Appendix S4 (Part II-5) to illustrate our choice.

### 2.5 Testing model adequacy

While model selection provides the best model among a set, it does not mean that this model provides a good representation of the data. Testing model adequacy is therefore an important step of phylogenetic comparative analyses (Pennell et al. 2015). We performed model adequacy tests on each empirical phylogeny by simulating 500 phylogenies under the best fit model and parameters and comparing simulated to empirical phylogenies. We developed the simul.comb.shifts() function to simulate phylogenies with a given combination of shifts. To compare simulated and empirical trees, we recorded the proportion of time points for which the empirical LTT falls within the 95% of the distribution of the simulated LTTs (explained in detail in Appendix S4 for each case). We also used the average leaf depth index, which quantifies the imbalance of a rooted tree (higher indices for more imbalanced trees, Fischer et al. 2021).

## 3 Results

### 3.1 Simulations

The simulation plan was set up to test both shift detections and palaeodiversity dynamic estimates. The analyses of the simulations initially revealed high proportions of false detections of shift (>11%, **Figure 3**, light orange). These false detections of shift led us to look at what they would be without the model selection procedure, *i.e.,* if we fitted only the true (simulated) models rather than selecting the best model from the set of models applied to subclades, backbones, and the whole phylogeny. By doing this, proportions of false detections of shift dropped below 10% (**Figure 3**, light blue). This finding sheds light on two problems: (1) the model selection, and (2) even with the simulated models, proportions of false detections of shift are higher than 5%. This latter problem shows that when a homogeneous model is simulated, splitting the optimisation of this model into a subclade and a backbone provides a gain of likelihood at the subclade level that is not penalised enough to avoid selecting a shift. We solved this by arbitrarily adding an extra penalisation with an additional parameter for the shift in the AICc calculation. This additional parameter can be thought as corresponding to the shift location as in previous methods such as MEDUSA (Alfaro et al. 2009); although the locations of the shifts are less free to vary in our approach than in MEDUSA, where all nodes are tested, they still need to be selected among a set. With this additional parameter and when fitting the models that were simulated, proportions of false detections of shift decreased below 5% (**Figure 3**, dark blue). If we use this additional parameter but used models that were selected rather than those that were simulated, proportions of false detections of shift are equal to 3.3% for BCST, 2.4% for BVAR, 13.8% for BVAR_DCST, and 22.7% for BCST_BVAR (**Figure 3**, dark orange). These results are satisfactory for simple models (BCST and BVAR) but false detections of shift are still important for more complex models.

**Figure 3.**
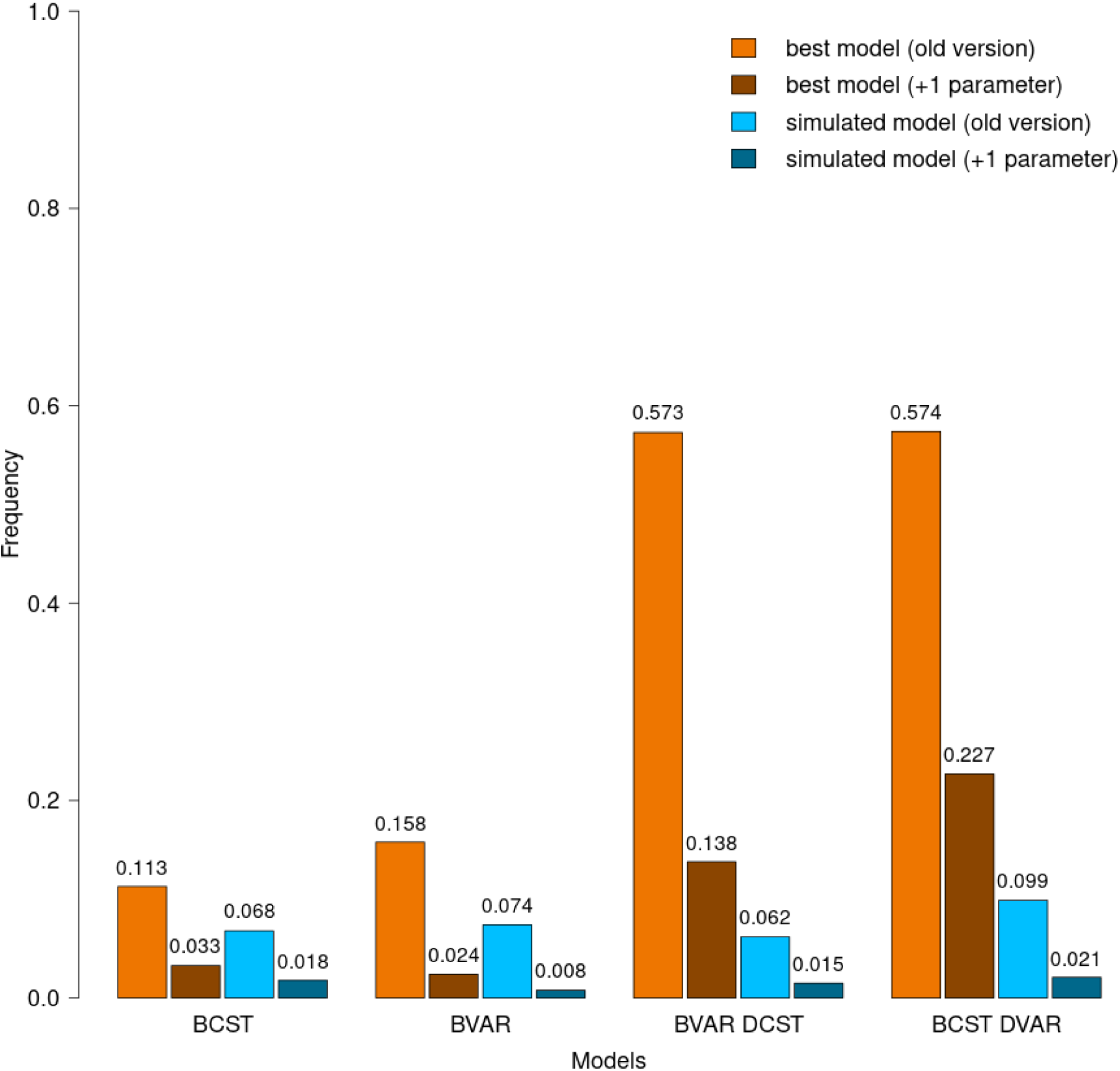
Proportions of false detection of shifts by models used to simulate the backbone depending on the method. Values for best-fitting models (in shades of orange) include the step of model selection for each part (subclade and backbone) while values for simulated models (in shades of blue) are calculated using the simulated models (no model selection). “Old version” corresponds to the AICc calculation from Morlon et al. (2011) and “+1 parameter” corresponds to the addition of one parameter for the location of the shift.

Concerning palaeodiversity dynamics, the main tendency (decline or not) is well recovered (BCST=97.35%, BVAR=84.6%, BVAR_DCST=89.5%, BCST_DVAR=92.65%) More precisely, we compared palaeodiversity curves from simulated rate values with palaeodiversity curves from estimated rate values. Figure 4 represents these comparisons for simulations that get a global median error *D* equal to the three quartiles of *D* for each model. We found rather low global median errors *D* for simpler models (BCST and BVAR models) (**Figure 4**). More complex models have slightly higher median values of *D* (BVAR_DCST and BCST_DVAR models), **Figure 4**). First and third quartiles follow the same tendency as the median across models, with higher values of *D* for more complex models. The mean of median values of *D* across the different models is equal to 0.233. There is no specific bias neither towards overestimation nor underestimation of palaeodiversity (Appendix S3).

**Figure 4.**
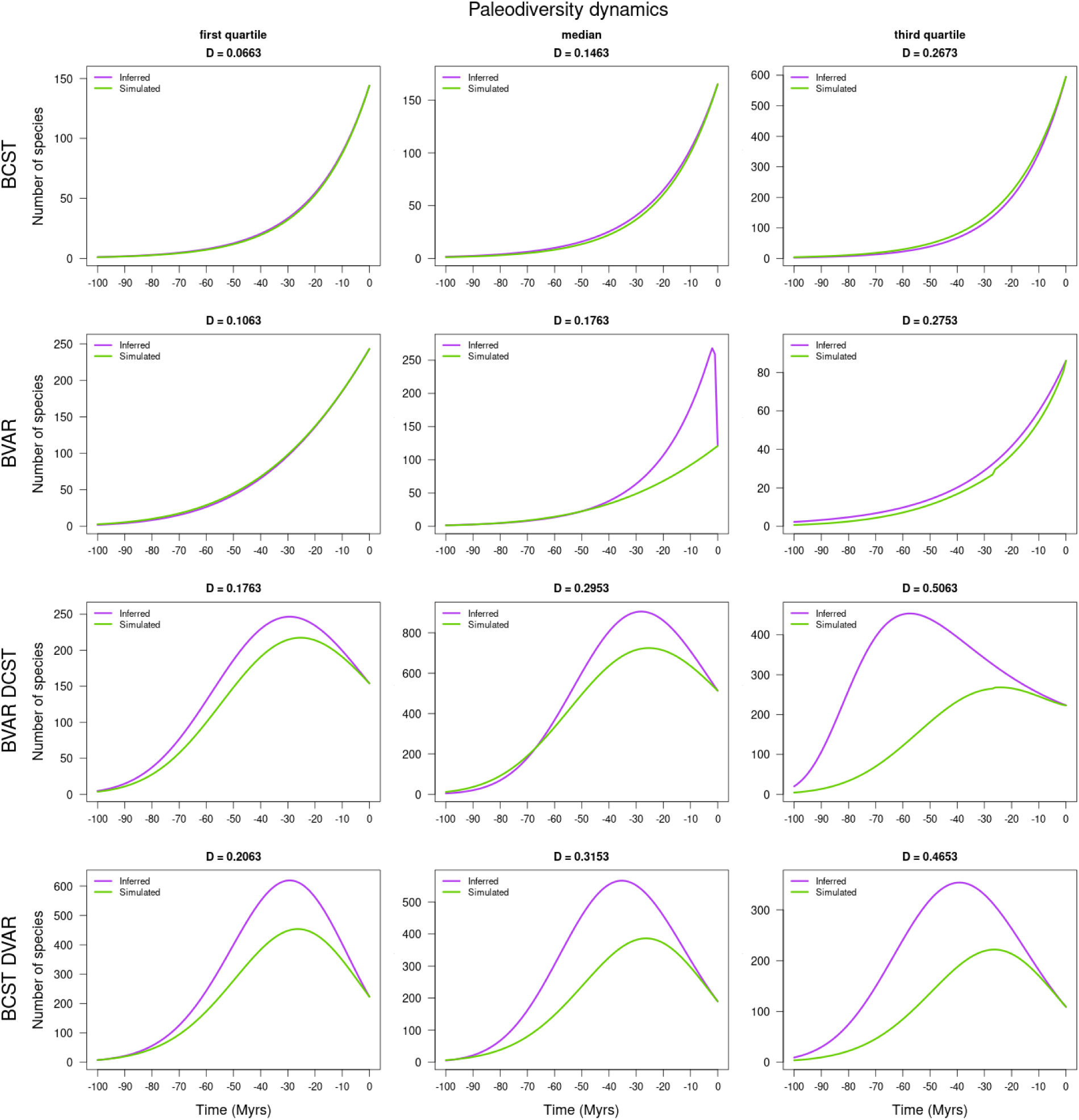
Comparison of inferred versus simulated palaeodiversity dynamics by model, with increasing global error *D*. For each model (in row) inferred palaeodiversity (purple) and simulated palaeodiversity (in green) are compared for the three quartiles of the global error *D* distributions by model (in column); see Appendix S3 for details. Note that the inferred palaeodiversity with the BVAR model (median) corresponds to a false detection of shift with the selection of a BVAR DCST model for the backbone creating this decline.

Concerning the additional simulations with extinction in subclades, the shifts were recovered in 99.2% of the simulations. However, the best supported model for 76.6% of the simulated subclades was the simplest BCST model (instead of the simulated BCST_DCST model), reflecting the loss of statistical power with decreasing tree size. Speciation and extinction rate estimates under BCST_DCST were unbiased, while the speciation rate was overestimated under BCST (Appendix S3, III-4).

### 3.2 Empirical evidence of extinction from phylogenies

#### 3.2.1 Cetacea

As in Morlon et al. (2011), we tested the presence of shifts in diversification at the base of the four main and recently diverged families (Balaenopteridae, Delphinidae, Phocoenidae and Ziphiidae) and the two parvorders (Mysteceti and Odotonceti) of Cetacea. This selection of subclades creates 25 combinations of shifts with a single backbone and 47 combinations of shifts if we add combinations with multiple backbones. When testing the shifts at the crown of subclades, we detected a single best combination with shifts for the four families tested (Table S1, ΔAICc=8.50). All the subclades follow a model with a decreasing speciation rate through time (see Appendix S4 for details). The simple backbone of this combination follows a model of diversification with a constant speciation rate and an increasing extinction rate leading to a phase of decline starting at 10 Ma before present, with a peak of palaeodiversity at 596 species (**Figure 5**). The same analysis with shifts tested at stem ages provides a best combination with a lower global AICc (Appendix S1, Table S5).

**Figure 5.**
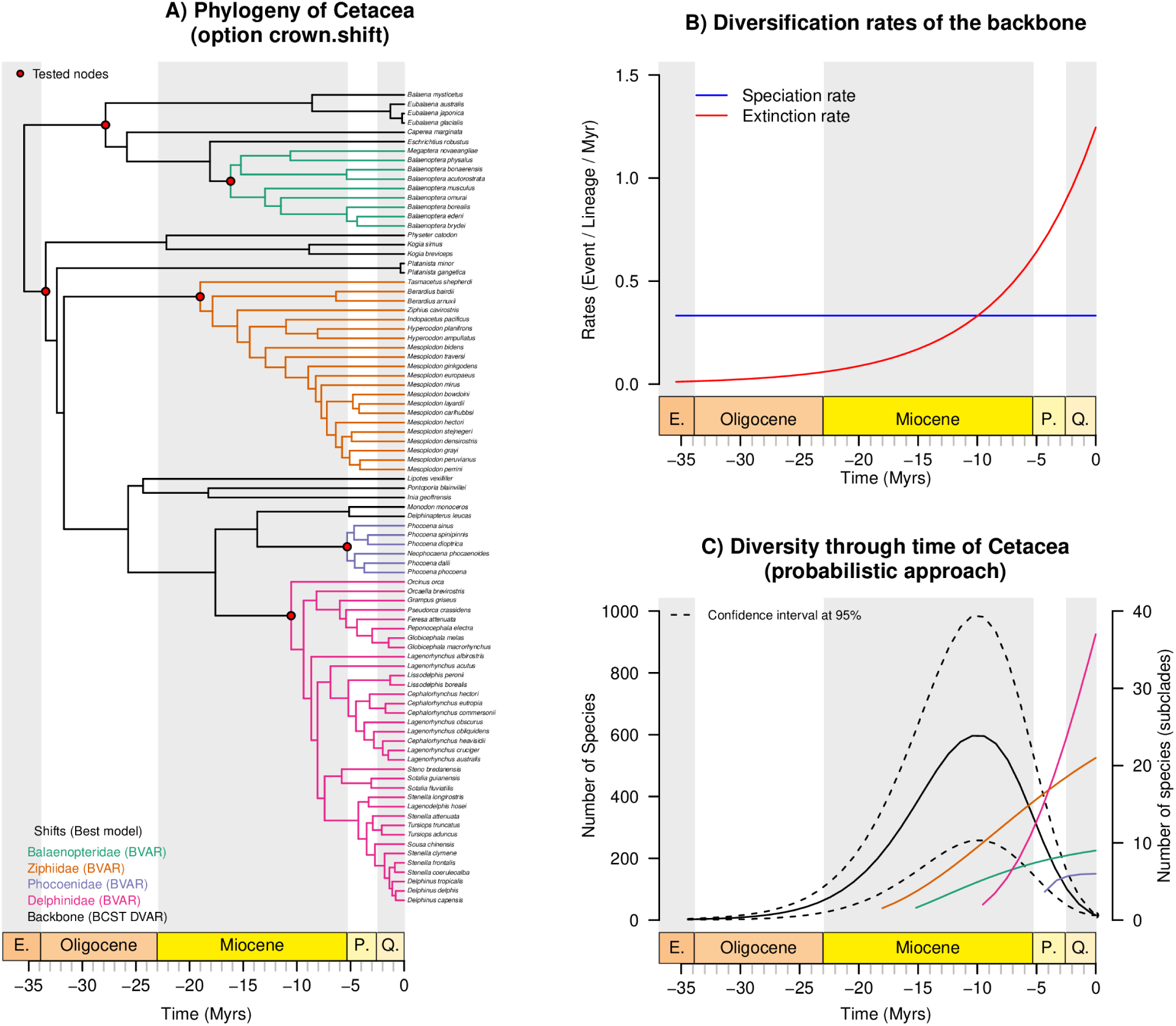
Diversification shifts and palaeodiversity dynamics estimated for Cetacea with shifts tested at crown ages. A) The phylogeny of Cetacea with shifts highlighted in colours and best models in parenthesis. Red dots correspond to all tested nodes. B) The evolution of diversification rates through time. C) The palaeodiversity dynamic estimated with the probabilistic approach for all parts of the best combination. To better distinguish subclade paleodiversity dynamics, a different y-axis is used on the right of the plot. Dotted line represents the confidence interval of diversity estimates for the backbone at 95% calculated with the probabilistic approach. For the sake of clarity, confidence intervals of diversity estimates for subclades are not represented.

#### 3.2.2 Vangidae

We tested shifts at three nodes in the Vangidae phylogeny corresponding to the Malagasy Vangidae clade and two nodes where shifts of morphological evolution have been detected in a previous study (Jønsson et al. 2012). Because these nodes are nested and there are only 32 species, only five combinations result from this initial selection of subclades. We did not detect any shift of diversification, and a model for the phylogeny without shift remains better (Table S2, ΔAICc=3.38) than other combinations. The whole phylogeny follows a model with a decreasing speciation rate resulting in a palaeodiversity increase through time, which slows down to present and reaches the current diversity of 32 species (**Figure 6**).

**Figure 6.**
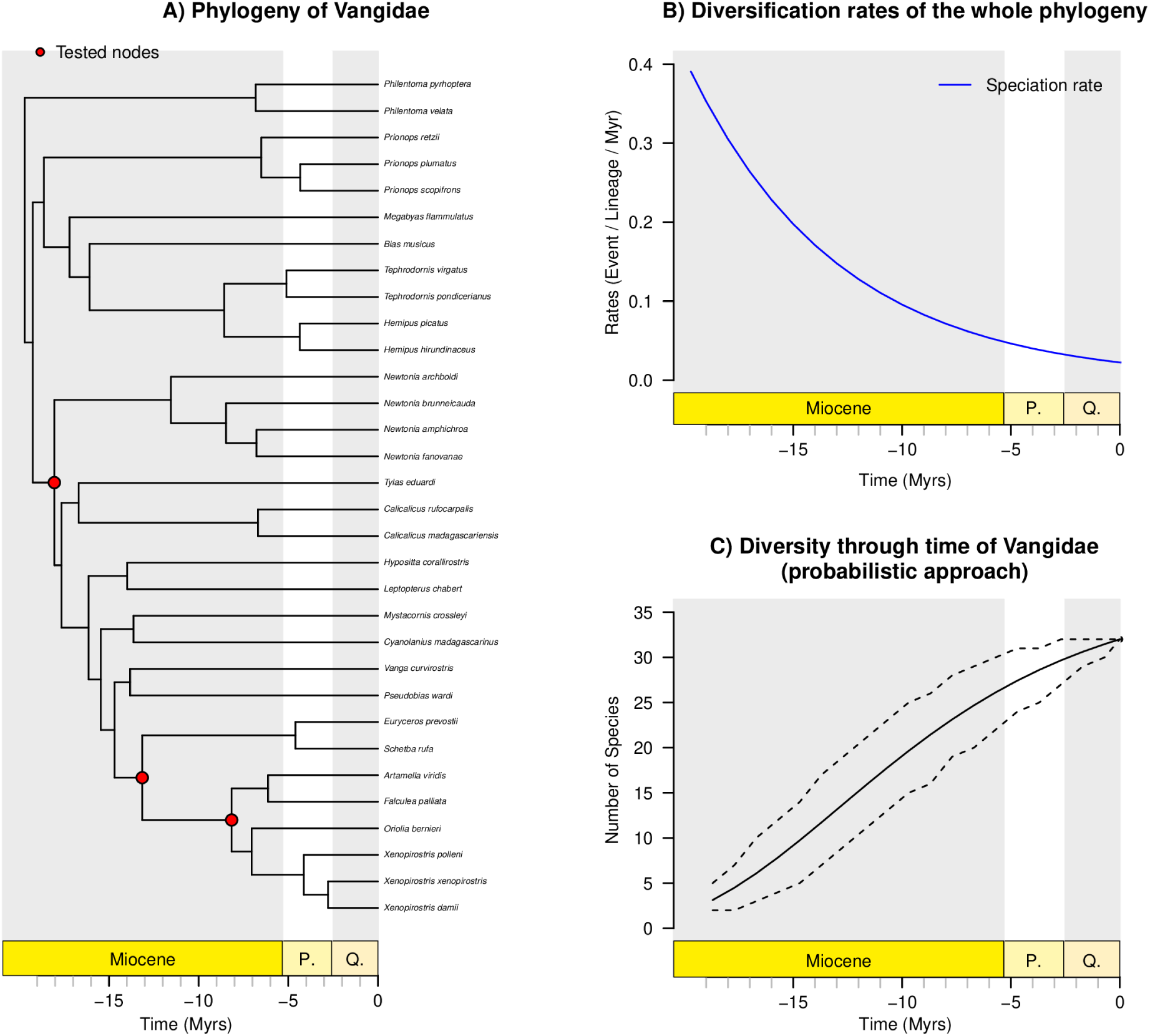
Diversification shifts and palaeodiversity dynamics estimated for Vangidae with stems included in backbone analysis. A) The phylogeny of Vangidae without shift detected (crown shifts). B) Diversification rates through time. C) The palaeodiversity dynamic estimated with the probabilistic approach. Dotted line represents the confidence interval of diversity estimates for the backbone at 95% calculated with the probabilistic approach.

#### 3.2.3 Parnassiinae

We tested 11 shifts corresponding to tribes (Luehdorfiini, Parnassiini, Zerynthiini), genera (*Allancastria*, *Parnassius*), subgenera (*Driopa*, *Tadumia*, *Kailasius*, *Koralius*, *Parnassius*), of which some match with host-plant shifts (genus *Parnassius*) or colonisation of Himalayan mountains (*Kailasius*, *Koramius*). This selection creates 392 combinations to compare with a single backbone and 1079 by adding combinations with multiple backbones. First results showed unrealistically high rates of diversification leading to unlikely palaeodiversity curves highlighting limits of some models (see Appendix S4 for details). We repeated the analysis by constraining the rates to a maximum value of 2 and found three best combinations among the 1079 combinations (Table S3, ΔAICc=2.13). Among these three best combinations, the first and the third combination of shifts showed highly unrealistic palaeodiversity dynamics despite the applied constraint on rate values. Given that the three combinations are statistically equivalent (ΔAICc < 2), we only kept the second combination that was more biologically realistic (ΔAICc=1.11 with the first combination, see Appendix S4 for details). Among the tested shifts, seven shifts compose the best combination with tribes (Luehdorfiini, Zerynthiini) and subgenera (*Driopa*, *Tadumia*, *Koramius*, *Kailasius* and *Parnassius,* **Figure 7**). The tribes Luehdorfiini, Zerynthiini and both subgenera *Parnassius* and *Tadumia* follow a model with a constant speciation rate while subgenera *Driopa*, *Kaliasius* and *Koramius* follow a model with a decreasing speciation rate through time. The backbone follows a model with a constant speciation rate and an increasing extinction rate through time (**Figure 7b**), leading to a period of negative net diversification rate from the beginning of the Miocene (19 Myrs ago) from the peak of the palaeodiversity of 324 species to the diversity of 88 extant species. This creates a palaeodiversity decline at the beginning of the Miocene.

**Figure 7.**
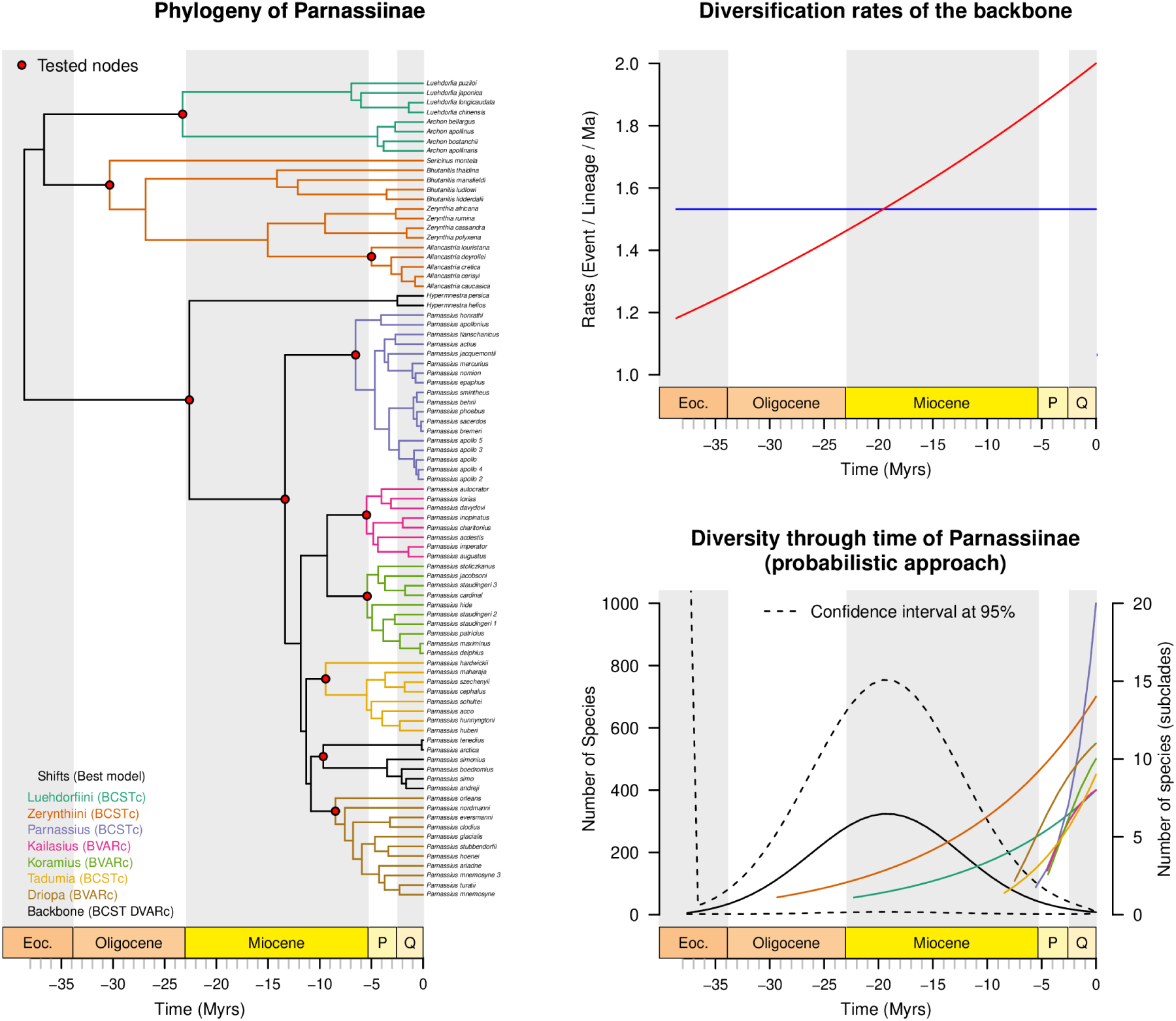
Diversification shifts and palaeodiversity dynamics estimated for Parnassiinae with stems included in backbone analysis. A) The phylogeny of Parnassiinae with shifts highlighted (crown shifts) in colours and best models in parenthesis. Red dots correspond to all tested nodes. B) Diversification rates through time. C) The palaeodiversity dynamic estimated with the probabilistic approach for all parts of the best combination. To better distinguish subclade paleodiversity dynamics, a different y-axis is used on the right of the plot. Dotted line represents the confidence interval of diversity estimates for the backbone at 95% calculated with the probabilistic approach. For the sake of clarity, confidence intervals of diversity estimates for subclades are not represented.

#### 3.2.4 Cycadales

We tested five shifts with the genera *Cycas*, *Ceratozamia*, *Zamia*, *Macrozamia*, and *Encephalartos*. This selection creates 32 combinations with a simple backbone and none with multiple backbones. The best combination is composed of all the tested subclades (Table S4, ΔAICc=11.96). The genera *Cycas*, *Ceratozamia* and *Macrozamia* follow pure-birth models with a constant speciation rate, while *Zamia* and *Encephalartos* follow models with a decreasing speciation rate (**Figure 8a**, Appendix S4). The diversification model of the backbone has a constant speciation rate and an increasing extinction rate, creating a negative net diversification rate starting 130 Myrs ago during the Early Cretaceous (**Figure 8b**). The global palaeodiversity increased again and reached the extant diversity thanks to the radiations of extant genera during the Miocene. The peak of palaeodiversity calculated with the deterministic approach reaches 200 species and ranges on posterior trees from 500 to 100 species (**Figure 8c**). The probabilistic approach estimates 150 species at the palaeodiversity with a confidence interval ranging from 400 to 0 species (**Figure 8d**).

**Figure 8.**
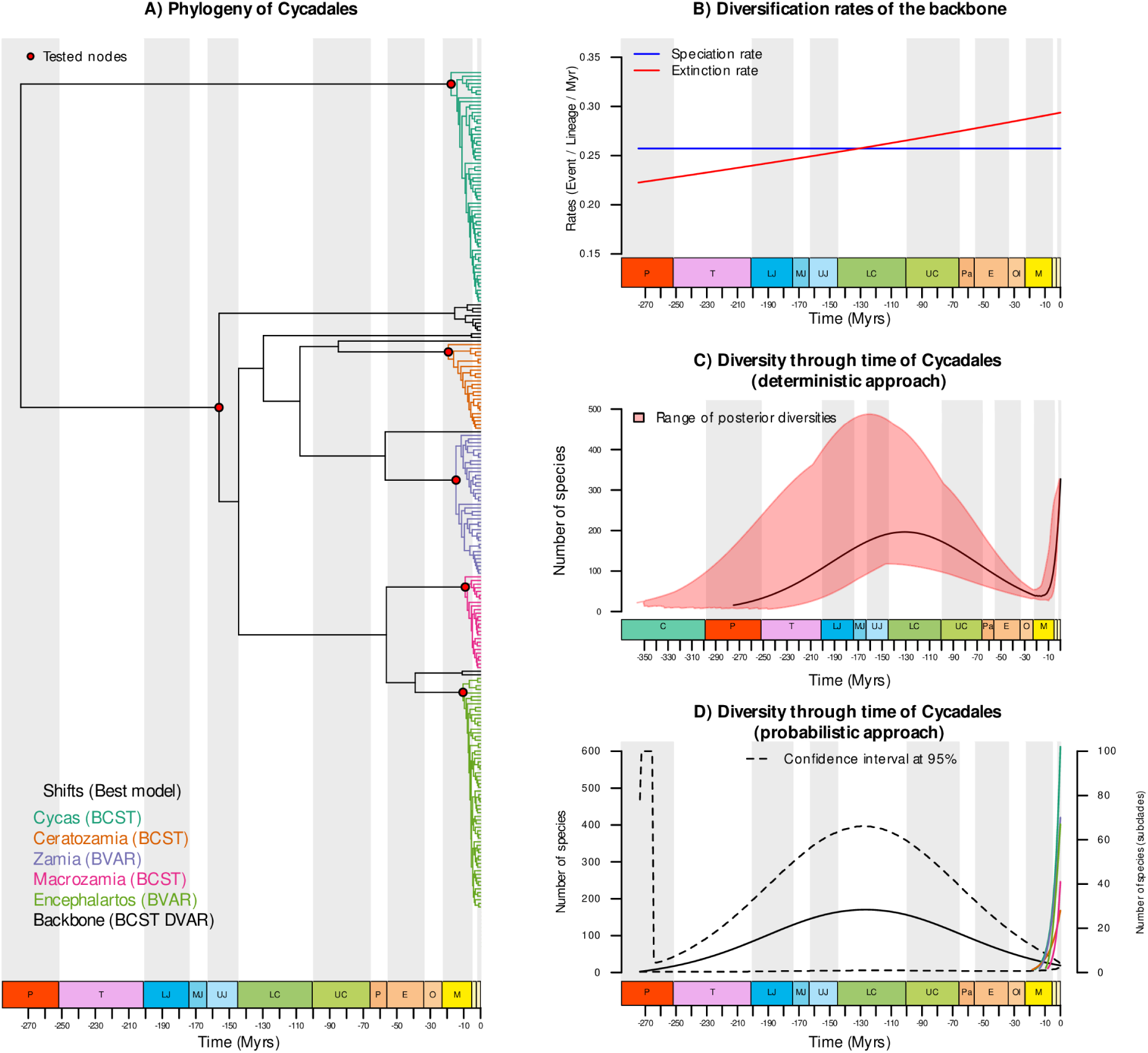
Diversification shifts and palaeodiversity dynamics estimated for Cycadales with stems included in backbone analysis. A) The phylogeny of cycads with shifts highlighted (crown shifts) in colours and best models in parenthesis. Red dots correspond to all tested nodes. B) The evolution of diversification rates through time. C) The palaeodiversity estimated with the deterministic approach. The red area represents the range of palaeodiversity estimates calculated on posterior trees. D) The palaeodiversity dynamic estimated with the probabilistic approach for all parts of the best combination. To better distinguish subclade paleodiversity dynamics, a different y-axis is used on the right of the plot. Dotted line represents the confidence interval of diversity estimates for the backbone at 95% calculated with the probabilistic approach. For the sake of clarity, confidence intervals of diversity estimates for subclades and species names are not represented.

### 3.3 Model adequacy

The model adequacy analyses allowed checking how well trees simulated under the best fit model and parameters mimic empirical trees. We found a good adequacy in terms of both branch length distribution and imbalance. 98.9% of the empirical LTT of the Cetacea fell within the 95% distribution of the simulated LTTs, 84.8.% for Parnassiinae, and 99.6% for Cycadales (Appendix S5, Appendix S4 III for details). The empirical values of the node depth index of imbalance fell in the 95% distribution of the simulated values for the three groups (Appendix S4, III for details).

## 4 Discussion

In this study, we automated and implemented in RPANDA, tested with simulations, and illustrated with empirical cases the approach developed by Morlon et al. (2011) that allows detecting shifts of diversification and estimating palaeodiversity dynamics through time from dated phylogenies. Several options and functions have been added to make this approach largely accessible to address an array of questions about heterogeneity of diversification rates and palaeodiversity dynamics by consolidating the challenging field of macroevolution in adopting hypothesis-driven approaches (Morlon et al. 2022). This emphasises the importance of additional information from biological knowledge on the focal groups to compare scenarios motivated by biological hypotheses (*e.g.* Legendre and Condamine 2018). In this sense, integrative taxonomic work is fundamental as it summarises broad morphological, ecological, and geographical divergences without neglecting historical divergence of clades thanks to phylogenetic analyses. Beyond taxonomy, ecological data, or other analyses such as ancestral state estimations can ground hypotheses for selecting subclades if the sampling fraction is known.

From a biological point of view, diversification histories are more complex than birth-death models. Accordingly, one should keep in mind that underlying assumptions of models influence the results. Among time- and clade-heterogeneous approaches, the most popular model, BAMM, investigates shifts along branches and does not allow periods of decline of diversity (Rabosky et al. 2013). The Multi-Type Birth-Death model developed by Barido-Sottani et al. (2020) provides a Bayesian framework that investigates shifts thanks to type-specific birth and death rate without defining location of the type changes. ClaDS models rate heterogeneity with small but frequent shifts (Maliet et al. 2019). MiSSE is an extension of HiSSE models that focuses on trait-free shifts and investigates heterogeneity of diversification rates at the tips of phylogenies (Vasconcelos et al. 2022). These alternative approaches can be used in a complementary way to study heterogeneity of diversification but will not necessarily provide similar results because of their underlying assumptions on diversification rates. Globally, each model is better at finding what it is made for: BAMM performs better to find large discrete shifts and will underestimate finer heterogeneity while ClaDS and, to a lesser degree MiSSE, will perform relatively worst in looking for large shifts but will be more accurate for small and more frequent changes of diversification rates (see Vasconcelos et al. 2022 for a comparison between BAMM, ClaDS and MiSSE). In our model, rates of birth-death models are either constant or exponentially variable over time and shifts occur at specific branch points. Despite the uncertainty being higher deep in the past, the main advantage of our approach is to provide inference of palaeodiversity dynamics by better estimating palaeodiversity values during deep periods, with the probabilistic approach to calculate palaeodiversity dynamics (see Billaud et al. 2020).

Previous studies on Vangidae (Jønsson et al. 2012), on Cetacea (Rabosky 2014), on Parnassiinae (Condamine et al. 2018), and on Cycadales (Condamine et al. 2015) have investigated heterogeneity of diversification with other methods. The cladogenesis test used for the Vangidae found one significant and one marginal shift among the nodes we tested. Contrary to the cladogenesis test that looks at the number of descendants from a node relatively to what is expected under a null birth-death model, our method applies and compares various birth-death models. Finding no shift can then come from the small size of the phylogeny but might also highlight that our approach is more conservative in detecting shifts in small phylogenies. Interestingly, BAMM analyses on the three other empirical cases estimated less diversification shifts (one in Cetacea vs. four here, one in Parnassiinae vs. seven here, and four in Cycadales vs. five here). This confirms that BAMM might be conservative in rate shift detection, although it has been shown to concern shifts with small clade sizes (Mitchell et al. 2019). Moreover, BAMM did not detect palaeodiversity decline contrary to our inferences. For instance, in Parnassiinae, we found a decline starting at 19 Ma from a peak at 324 species to the current diversity of the backbone (8 species). This is consistent with Condamine et al. (2018) who applied models assuming instantaneous drops of diversity through time at specific time points (Stadler 2011a; May et al. 2016). This study found a sudden extinction around 15 Ma, 4 Myrs later, with a loss of 89% of the diversity (from 90 species to 10). Both methods recover a decline of diversity, although different assumptions behind each model (mass extinction vs. exponential dependency to time) influence the inferred timing and the scale of the decline.

The approach we automated has already been recognized for its ability to better estimate extinction rates (Stadler 2011b). Our simulations confirm the capacity of the model to recover non-zero extinction rate in the backbone that can exceed speciation rate, resulting in a diversity decline. Despite the high values of false detection of shifts for complex models, the difference between observed and simulated palaeodiversity was reasonable with a mean of median global error *D*=0.233 on average over models and close to the value found in Billaud et al. (2020) with a *D*=0.22. This result shows that even with an error on the detection of shift, the model estimates a trustworthy palaeodiversity dynamic (**Figure 4**). Moreover, our empirical analyses, confirmed by posterior predictive analyses, suggest diversity declines are more widespread than previously considered. Cetacea, Parnassiinae, and Cycadales show a common pattern with a single period of decline, but we can address the question to what extent patterns with many diversity declines might be detected. Indeed, the main limit of our approach is that important levels of extinction are likely to erode both diversity and statistical power. Splitting a phylogeny into backbones and subclades requires enough species to be shared in different parts that will be analysed separately. Dated phylogenies with only a few extant species cannot be investigated by our approach although we sometimes know from the fossil record that they experienced extinction. The clade of Ginkgophyta is a caricatural example of this problem as it only contains one lineage (*Ginkgo biloba*) despite 275 Ma of evolution, and probably experienced a lot of extinction events (Royer et al. 2003). An interesting perspective will be to simulate more complex patterns of extinction with multiple backbones and multiple periods of decline to see whether we can properly recover these patterns.

Concerning subclades, we recovered only models without extinction, although some subclades are quite ancient (*e.g.* in Parnassiinae). We ran some simulations that tend to confirm the difficulty to select models with extinction in subclades. This is not surprising given the small size of subclades (*e.g.* six species of Phocoenidae in Cetacea), which do not provide sufficient sample size to select parameter-rich models (*i.e.*, with extinction). Low extinction rates in subclades can also be a real pattern in some cases. For the Cetacea, the phylogenetic placement of fossils indicates that extinct taxa are concentrated in the backbone, in particular along stems of the four main families (Lloyd and Slater 2021), which is consistent with our results indicating that the backbone bears most of the extinction signal in the tree.

Given the difficulty to estimate extinction rates (Morlon 2014; Beaulieu and O’Meara 2015; Rabosky 2016), studies have repeatedly raised debates about models in macroevolution (Kubo and Iwasa 1995; Rabosky and Lovette 2008; Crisp and Cook 2009; Burin et al. 2019; Pannetier et al. 2021). Recently, Louca and Pennell (2020) argued that diversification rates should not be estimated from birth-death models with clade-homogeneous rates because any reconstructed phylogeny can be equally well explained with an infinite number of alternative models. Here we adopt a hypothesis-testing approach, in which we consider a limited set of alternative models chosen *a priori*, some of which will explain the phylogeny better than others (Helmstetter et al. 2022, Morlon et al. 2022). Even when allowing for potential rate shifts in only few clades, our approach sometimes compared thousands of combinations of shifts; still, cases with many statistically equivalent combinations were rare. Methodological developments and collecting more data (*e.g.* to update taxonomy, to increase sampling fractions) are required to investigate questions in such a challenging field while keeping in mind that rates inferred from extant time-trees are tentative estimates of a long history and should be interpreted cautiously (Helmstetter et al. 2022).

In conclusion, the automated implementation of the approach originally developed by Morlon et al. (2011) provides a user-friendly tool to investigate heterogeneity of diversification. We improved this approach by (1) adding the possibility to test shifts at crown age that seems to drastically improve the global likelihood of the approach (see Cetacea for example), (2) select a set of constant or time-dependent diversification models, (3) easily set up multiple backbone investigation, (4) make it possible to constrain rate estimates, (5) infer palaeodiversity dynamics while considering uncertainty both within rate estimation (probabilistic approach) and across many trees that can differ in terms of topology or divergence times, and (6) confirm empirical results with model adequacy tests. Nonetheless, we encourage users to adopt a hypothesis-driven approach and to be parsimonious concerning the number of nodes to test. We also recommend being critical on results of inferred palaeodiversity of the backbone in terms of raw number of species that results from rate values. We also caution that the most complex model where both speciation and extinction rates vary exponentially (BVAR_DVAR) is prone to provide highly unrealistic patterns of palaeodiversity dynamics as shown in an example with Cetacea (Appendix S4, II-5). Echoing the results from Burin et al. (2019), the BVAR_DVAR model should be avoided or constrained. Keeping in mind these limitations, our study highlights that extinction signature and palaeodiversity declines can be investigated in groups without species-level fossil records.

## Acknowledgements

We thank three anonymous reviewers who provided constructive comments that improved the study. We thank Arthur Weyna who helped us setting up simulations with Snakemake. This project benefited from the Montpellier Bioinformatics Biodiversity platform supported by the LabEx CeMEB, an “Investissements d’Avenir” program managed by the Agence Nationale de la Recherche (ANR-10-LABX-04-01). N.M. was supported by the LabEx CEBA, an “Investissements d’Avenir” program managed by the Agence Nationale de la Recherche (ANR-10-LABX-25-01). H.M. was supported by funding from the European Research Council (ERC) under the European Union’s Horizon 2020 research and innovation programme (project PANDA, agreement No. 616419). F.L.C. was supported by funding from the European Research Council (ERC) under the European Union’s Horizon 2020 research and innovation programme (project GAIA, agreement No. 851188).

## Authors’ contribution

N.M. and F.L.C. designed the research with advice from P.-H.F.; N.M. performed the research and analysed the data with advice from H.M. and F.L.C.; and N.M. wrote the manuscript, and all authors edited the text.

## Data availability statement

This open-source software is written entirely in the R language and is freely available through the GitHub repository of PANDA package on the branch named clade.shift.model at: https://github.com/hmorlon/PANDA/tree/clade.shift.model. The scripts used, simulation results, model fits and case study data are detailed and available at: https://figshare.com/s/6e8924774165b5ed81ec

