## Appendix S1 for "Estimating clade-specific diversification rates and palaeodiversity dynamics from reconstructed phylogenies"

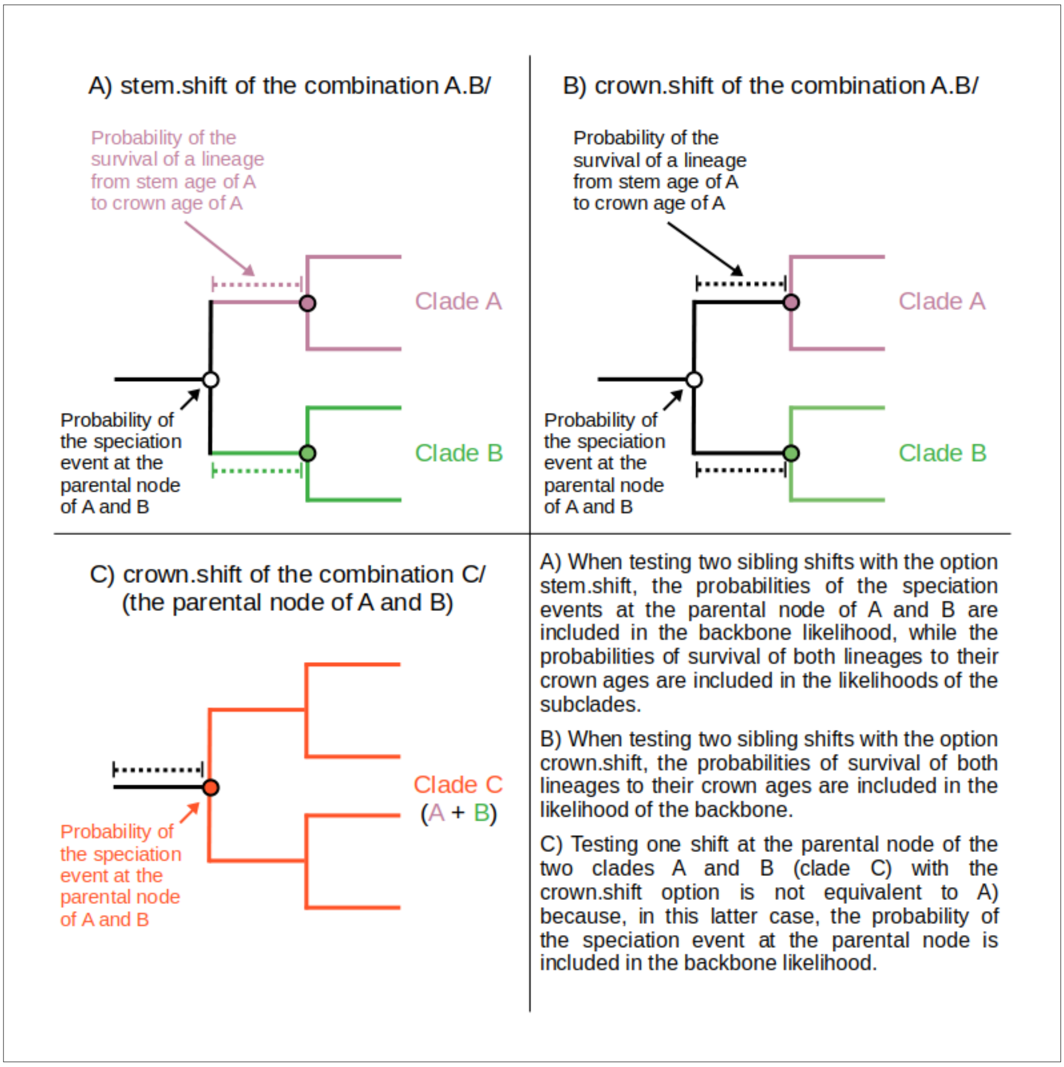


**Figure S1: Differences between backbone.option A) "stem.shift" and B) "crown.shift" and a case where testing with backbone.option="crown.shift" at the parental node of two sister clades is not equivalent to test simultaneous shifts at both clade nodes with backbone.option="stem.shift".**

The difference of likelihood computation between stem.shift (Figure 2A) and crown.shift (Figure 2B, C) is that the probability of the lineage survival is included in the backbone model such as the diversification rates of the subclade start at the crown age. In other words, it means that the change in diversification synchronously occurs with the cladogenesis event as it could be expected under an adaptive radiation (Figure 2B). Accordingly, crown.shift option can be seen as a case in which the shift of diversification rate leads to or creates the cladogeneis event because the speciation event at the crown is included in the likelihood of the shift. In addition, stem ages are rarely available, or their length are less reliable depending on the quality of the outgroups used in the phylogenetic reconstruction.
