## Appendix S2 for "Estimating clade-specific diversification rates and palaeodiversity dynamics from reconstructed phylogenies"

**Appendix S2.** Results on Cetacea when testing at stem age as in Morlon et al. (2011)

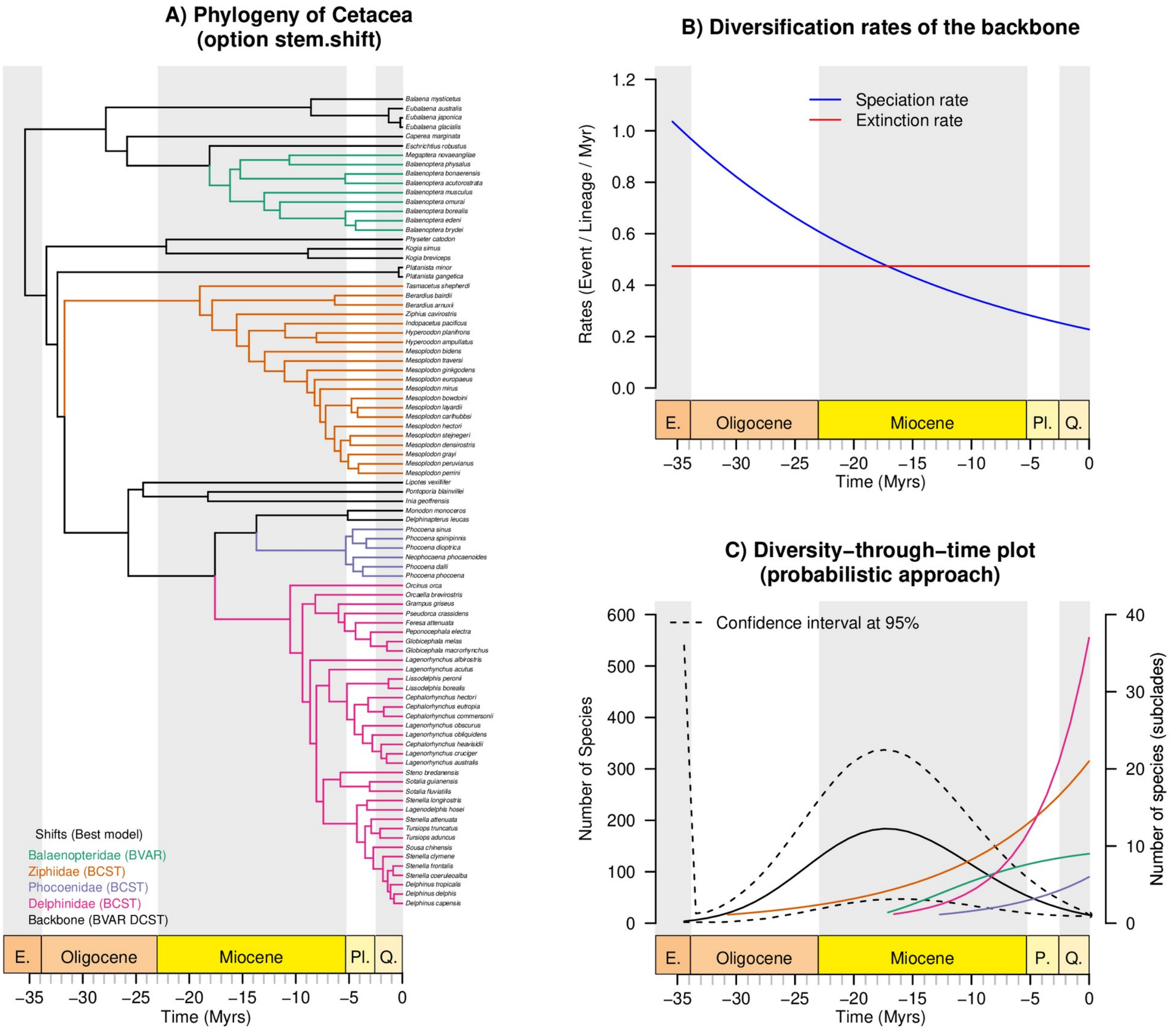


**Figure S2. Diversification shifts and palaeodiversity dynamics estimated for Cetacea with stems included in subclade analyses.** A) The phylogeny of Cetacea with shifts highlighted (stem shifts) in colours and best models in parenthesis. Red dots correspond to all tested nodes. B) The evolution of diversification rates through time. C) The palaeodiversity dynamic estimated with the probabilistic approach. Dotted line represents the confidence interval of diversity estimates for the backbone at 95% calculated with the probabilistic approach. For the sake of clarity, confidence intervals of diversity estimates for subclades are not represented.

As in Morlon et al. (2011), we tested the four main and recently diverged families (Balaenopteridae, Delphinidae, Phocoenidae and Ziphiidae) and the two parvorders (Mysteceti and Odotonceti) of Cetacea. This selection of subclades creates 25 combinations of shifts with a single backbone and 47 combinaisons of shifts with multiple backbones included. When testing the shifts at the stem of subclades, three combinations were the best in terms of AICc (Table S5). We only represented the best of them in **Figure S2** but other combinations can be reproduced by following **Appendix S4**. This combination is composed of the four tested families without any shift inside the backbone. Three of the subclades follow a model with constant speciation rate except the Balaenopteridae who experienced a model with a decreasing speciation rate. The backbone of this combination follows a model with a decreasing speciation rate through time and a constant extinction rate. The extinction rate exceeds the speciation rate 18 Myrs ago, creating a decline of diversity from this age with a maximum of 174 species. At the peak, the confidence interval range is approximately from 50 species to almost 350 species.
