## Supplementary material for "Estimating clade-specific diversification rates and palaeodiversity dynamics from reconstructed phylogenies": Tables_S1-S5

**Table S1:** A) Global comparison of combinations of diversification shifts for Cetacea (shifts are tested at the crown) with B) the rates of the diversification model for their backbones. When the combination has multiple backbones, only the deepest is displayed (more details in Appendix S4). The best combination of shift and the phylogeny analysed with no shift are highlighted in bold. NP = Number of parameters, logL = log(Likelihood), λ = speciation rate (at present if variable), α = dependency parameter of speciation rate, μ = extinction rate, β = dependency parameter of extinction rate

| **Total** | | | | | **Backbone (only deep backbone when multiple backbones)** | | | | | | | |
|---|---|---|---|---|---|---|---|---|---|---|---|---|
| **Combination** | **NP** | **logL** | **AICc** | **ΔAICc** | **Model** | **NP** | **logL** | **AICc** | **λ** | **α** | **μ** | **β** |
| **95.109.135.140/** | **15** | **-233.477** | **517.94** | **0** | **BCST DVAR** | **3** | **-57.119** | **122.237** | **0.332** | **–** | **1.245** | **-0.133** |
| 95.109.140/ | 12 | -247.062 | 526.443 | 8.503 | BVAR DCST | 3 | -78.2 | 163.734 | 0.186 | 0.044 | 0.392 | – |
| 89.109.135.140/ | 14 | -240.927 | 529.039 | 11.099 | BVAR DCST | 3 | -38.952 | 87.905 | 0.325 | 0.037 | 0.647 | – |
| 109.135.140/ | 12 | -246.105 | 531.538 | 13.598 | BVAR DCST | 3 | -94.728 | 196.6 | 0.091 | 0.052 | 0.2 | – |
| 89.109.140/ | 11 | -252.469 | 532.125 | 14.184 | BVAR DCST | 3 | -57.992 | 123.984 | 0.19 | 0.042 | 0.407 | – |
| 109.140/ | 9 | -255.542 | 532.158 | 14.218 | BVAR DCST | 3 | -111.662 | 230.213 | 0.096 | 0.046 | 0.176 | – |
| 95.109.135.140/103 | 19 | -228.97 | 534.927 | 16.986 | BVAR DCST | 3 | -15.561 | 49.122 | 1.165 | 0.017 | 1.627 | – |
| 140/ | 5 | -262.311 | 535.641 | 17.7 | BVAR | 2 | -180.635 | 365.515 | 0.059 | 0.024 | – | – |
| 109.135.140/103 | 14 | -242.896 | 536.285 | 18.345 | BCST | 1 | -54.469 | 111.245 | 0.072 | – | – | – |
| 95.109.135.140/89 | 17 | -237.787 | 536.56 | 18.62 | BVAR DCST | 3 | -38.952 | 87.905 | 0.325 | 0.037 | 0.647 | – |
| 135.140/ | 8 | -254.329 | 537.712 | 19.772 | BVAR | 2 | -165.157 | 334.592 | 0.051 | 0.03 | – | – |
| 95.109.140/89 | 14 | -249.33 | 539.645 | 21.705 | BVAR DCST | 3 | -38.952 | 87.905 | 0.325 | 0.037 | 0.647 | – |
| 95.109.140/103 | 16 | -242.805 | 540.233 | 22.292 | BVAR DCST | 3 | -15.561 | 49.122 | 1.165 | 0.017 | 1.627 | – |
| 95.140/ | 7 | -260.389 | 540.45 | 22.51 | BCST | 1 | -153.731 | 309.56 | 0.076 | – | – | – |
| 109.140/103 | 11 | -256.731 | 541.591 | 23.651 | BCST | 1 | -54.469 | 111.245 | 0.072 | – | – | – |
| 95.135.140/ | 10 | -252.848 | 543.384 | 25.443 | BCST | 1 | -138.693 | 279.499 | 0.071 | – | – | – |
| 95.140/103 | 11 | -252.602 | 545.132 | 27.192 | BVAR DCST | 3 | -15.561 | 49.122 | 1.165 | 0.017 | 1.627 | – |
| 89.140/ | 6 | -266.111 | 546.111 | 28.171 | BCST | 1 | -133.838 | 269.79 | 0.08 | – | – | – |
| 140/103 | 6 | -266.528 | 546.491 | 28.55 | BCST | 1 | -54.469 | 111.245 | 0.072 | – | – | – |
| 95.135.140/103 | 15 | -243.499 | 547.461 | 29.52 | BVAR DCST | 3 | -15.561 | 49.122 | 1.165 | 0.017 | 1.627 | – |
| 89.135.140/ | 10 | -257.105 | 548.413 | 30.473 | BVAR | 2 | -117.335 | 239.098 | 0.05 | 0.033 | – | – |
| 135.140/103 | 10 | -257.424 | 548.82 | 30.879 | BVAR | 3 | -113.783 | 234.455 | 0.052 | 0.029 | – | – |
| 95.140/89 | 9 | -262.972 | 553.632 | 35.692 | BVAR DCST | 3 | -38.952 | 87.905 | 0.325 | 0.037 | 0.647 | – |
| 109/ | 4 | -272.737 | 554.948 | 37.008 | BCST | 1 | -210.533 | 423.129 | 0.108 | – | – | – |
| 95.109/ | 8 | -266.269 | 554.972 | 37.031 | BCST DCST | 2 | -179.083 | 362.388 | 0.187 | – | 0.13 | – |
| **No shift** | **1** | **-276.789** | **555.626** | **37.685** | **–** | **–** | **–** | **–** | **–** | **–** | **–** | **–** |
| 95.109.135/ | 12 | -256.583 | 555.889 | 37.948 | BVAR DCST | 3 | -161.9 | 330.311 | 0.247 | 0.024 | 0.328 | – |

| | | | | | | | | | | | | |
|---|---|---|---|---|---|---|---|---|---|---|---|---|
| 95.135.140/89 | 13 | -253.966 | 555.934 | 37.994 | BVAR DCST | 3 | -38.952 | 87.905 | 0.325 | 0.037 | 0.647 | – |
| 89.109/ | 7 | -270.335 | 557.331 | 39.391 | BCST DCST | 2 | -157.533 | 319.317 | 0.212 | – | 0.153 | – |
| 109.135/ | 8 | -264.643 | 558.907 | 40.967 | BCST DCST | 2 | -194.942 | 394.094 | 0.15 | – | 0.082 | – |
| 89.109.135/ | 11 | -261.342 | 559.681 | 41.74 | BVAR DCST | 3 | -141.043 | 288.672 | 0.261 | 0.018 | 0.314 | – |
| 95/ | 4 | -273.489 | 559.831 | 41.891 | BCST | 1 | -248.507 | 499.067 | 0.109 | – | – | – |
| 135/ | 4 | -270.139 | 560.328 | 42.387 | BCST | 1 | -262.642 | 527.334 | 0.103 | – | – | – |
| 95.109/103 | 11 | -259.957 | 560.376 | 42.435 | BVAR DCST | 3 | -15.561 | 49.122 | 1.165 | 0.017 | 1.627 | – |
| 95.103/ | 8 | -264.338 | 561.65 | 43.709 | BVAR DCST | 3 | -15.561 | 49.122 | 1.165 | 0.017 | 1.627 | – |
| 109/103 | 6 | -273.882 | 561.734 | 43.794 | BCST | 1 | -54.469 | 111.245 | 0.072 | – | – | – |
| 89/ | 3 | -277.896 | 562.849 | 44.908 | BCST | 1 | -227.298 | 456.653 | 0.117 | – | – | – |
| 103/ | 3 | -278.263 | 563.008 | 45.068 | BCST | 1 | -54.469 | 111.245 | 0.072 | – | – | – |
| 95.135/ | 7 | -266.902 | 564.662 | 46.721 | BCST | 1 | -234.423 | 470.903 | 0.106 | – | – | – |
| 95.109/89 | 10 | -267.195 | 564.852 | 46.912 | BVAR DCST | 3 | -38.952 | 87.905 | 0.325 | 0.037 | 0.647 | – |
| 95.109.135/103 | 15 | -252.378 | 565.553 | 47.613 | BVAR DCST | 3 | -15.561 | 49.122 | 1.165 | 0.017 | 1.627 | – |
| 95.135/103 | 11 | -257.904 | 566.799 | 48.859 | BVAR DCST | 3 | -15.561 | 49.122 | 1.165 | 0.017 | 1.627 | – |
| 109.135/103 | 10 | -266.304 | 566.912 | 48.972 | BCST | 1 | -54.469 | 111.245 | 0.072 | – | – | – |
| 95.109.135/89 | 14 | -258.202 | 567.202 | 49.261 | BVAR DCST | 3 | -38.952 | 87.905 | 0.325 | 0.037 | 0.647 | – |
| 89.135/ | 6 | -271.452 | 567.967 | 50.027 | BCST | 1 | -213.358 | 428.778 | 0.114 | – | – | – |
| 135/103 | 6 | -271.83 | 568.158 | 50.218 | BCST | 1 | -54.469 | 111.245 | 0.072 | – | – | – |
| 95/89 | 6 | -274.756 | 570.37 | 52.429 | BVAR DCST | 3 | -38.952 | 87.905 | 0.325 | 0.037 | 0.647 | – |
| 95.135/89 | 9 | -268.313 | 575.488 | 57.548 | BVAR DCST | 3 | -38.952 | 87.905 | 0.325 | 0.037 | 0.647 | – |

**Table S2:** A) Global comparison of combinations of diversification shifts for Vangidae (shifts are tested at the crown) with B) the rates of the diversification model for their backbones. Best combination (no shift) is highlighted in bold. NP = Number of parameters, logL = log(Likelihood), λ = speciation rate (at present if variable), α = dependency parameter of speciation rate, μ = extinction rate, β = dependency parameter of extinction rate

| **A) Total** | | | | | **B) Backbone** | | | | | | | |
|---|---|---|---|---|---|---|---|---|---|---|---|---|
| **Combination** | **NP** | **logL** | **AICc** | **ΔAICc** | **Model** | **NP** | **logL** | **AICc** | **λ** | **α** | **μ** | **β** |
| **No shift** | **2** | **-96.566** | **197.547** | **0** | **–** | **–** | **–** | **–** | **–** | **–** | **–** | **–** |
| 50/ | 4 | -94.979 | 200.93 | 3.383 | BVAR | 2 | -73.041 | 150.653 | 0.013 | 0.18 | – | – |
| 52/ | 4 | -94.706 | 201.934 | 4.388 | BVAR | 2 | -79.779 | 164.08 | 0.014 | 0.175 | – | – |
| 43/ | 4 | -96.906 | 203.667 | 6.121 | BCST | 1 | -35.511 | 73.467 | 0.078 | – | – | – |
| 50/43 | 6 | -93.829 | 205.168 | 7.622 | BVAR | 3 | -36.379 | 81.425 | 0.004 | 0.277 | – | – |
| 52/43 | 6 | -93.954 | 206.533 | 8.987 | BVAR | 3 | -43.515 | 95.213 | 0.006 | 0.251 | – | – |

**Table S3:** A) Global comparison of combinations of diversification shifts for Parnassiinae (shifts are tested a
equal to 2) with B) the rates of the diversification model for their backbones. When the combination has mult
(more details in Appendix S4). The best combination of shift and the phylogeny analysed with no shift a
parameters, logL = log(Likelihood), λ = speciation rate (at present if variable), α = dependency parameter
dependency parameter of extinction rate

| A) Total | | | | | B) Backbone (only deep b | | |
|---|---|---|---|---|---|---|---|
| **Combination** | **NP** | **logL** | **AICc** | **ΔAICc** | **Model** | **NP** | **logL** |
| **95.111/108** | **9** | **-215.807** | **457.79** | **0** | **BVAR DCSTc** | **3** | **-15.158** |
| **88.95.111.130.137.147.160/** | **20** | **-196.388** | **458.896** | **1.106** | **BCST DVARc** | **3** | **-19.12** |
| **95.108/** | **7** | **-218.938** | **459.167** | **1.376** | **BVAR DCSTc** | **3** | **-15.158** |
| 95.111.160/108 | 12 | -212.109 | 459.924 | 2.134 | BVAR DCSTc | 3 | -15.158 |
| 95.111.137/108 | 12 | -212.318 | 460.901 | 3.111 | BVAR DCSTc | 3 | -15.158 |
| 88.95.130.137.147.155.160/ | 20 | -198.092 | 461.004 | 3.214 | BVAR DCSTc | 3 | -40.366 |
| 95.160/108 | 10 | -215.145 | 461.055 | 3.265 | BVAR DCSTc | 3 | -15.158 |
| 95.111.130/108 | 12 | -212.313 | 462.869 | 5.079 | BVAR DCSTc | 3 | -15.158 |
| 95.111.137.160/108 | 15 | -208.634 | 463.158 | 5.368 | BVAR DCSTc | 3 | -15.158 |
| 95.137/108 | 10 | -216.077 | 463.485 | 5.695 | BVAR DCSTc | 3 | -15.158 |
| 104.111/95.108 | 11 | -213.379 | 463.844 | 6.053 | BVAR DCSTc | 3 | -15.158 |
| 95.111.147/108 | 11 | -215.72 | 464.083 | 6.293 | BVAR DCSTc | 3 | -15.158 |
| 110/ | 5 | -226.786 | 464.565 | 6.774 | BCST DCSTc | 2 | -80.506 |
| 95.130/108 | 10 | -215.825 | 464.971 | 7.181 | BVAR DCSTc | 3 | -15.158 |
| 95.111.130.160/108 | 15 | -208.631 | 465.103 | 7.313 | BVAR DCSTc | 3 | -15.158 |
| 95.137.160/108 | 13 | -212.175 | 465.176 | 7.386 | BVAR DCSTc | 3 | -15.158 |
| 95.147/108 | 9 | -218.859 | 465.439 | 7.649 | BVAR DCSTc | 3 | -15.158 |
| 88.104.111.130.137.147.160/95 | 22 | -194.291 | 465.61 | 7.82 | BCST DVARc | 3 | -19.12 |
| 95.111.155/108 | 11 | -215.801 | 465.826 | 8.036 | BVAR DCSTc | 3 | -15.158 |
| 104.108/95 | 9 | -216.84 | 465.881 | 8.09 | BVAR DCSTc | 3 | -15.158 |
| 160/110 | 7 | -223.923 | 466.102 | 8.311 | BCST DCSTc | 2 | -80.506 |
| 95.111.147.160/108 | 14 | -212.013 | 466.268 | 8.478 | BVAR DCSTc | 3 | -15.158 |
| 104.111.160/95.108 | 14 | -209.851 | 466.317 | 8.527 | BVAR DCSTc | 3 | -15.158 |
| 95.110/ | 7 | -224.554 | 466.334 | 8.544 | BCST DCSTc | 2 | -32.891 |
| 95.130.160/108 | 13 | -211.865 | 466.543 | 8.752 | BVAR DCSTc | 3 | -15.158 |
| 95.147.160/108 | 12 | -214.77 | 466.752 | 8.962 | BVAR DCSTc | 3 | -15.158 |
| 95.111.137.147/108 | 14 | -212.238 | 467.266 | 9.476 | BVAR DCSTc | 3 | -15.158 |
| 104.111.137/95.108 | 14 | -210.057 | 467.289 | 9.499 | BVAR DCSTc | 3 | -15.158 |
| 88.110/ | 8 | -223.287 | 467.396 | 9.605 | BVAR DCSTc | 3 | -53.71 |
| 88.95.130.137.147.160/ | 18 | -205.528 | 467.466 | 9.676 | BVAR DCSTc | 3 | -61.169 |
| 95.155/108 | 9 | -219.087 | 467.488 | 9.697 | BVAR DCSTc | 3 | -15.158 |
| 88.104.130.137.147.155.160/95 | 22 | -195.995 | 467.718 | 9.928 | BVAR DCSTc | 3 | -40.366 |
| 104.160/95.108 | 12 | -213.062 | 467.798 | 10.008 | BVAR DCSTc | 3 | -15.158 |
| 95.130.137.147.155.160/108 | 22 | -197.154 | 467.894 | 10.104 | BVAR DCSTc | 3 | -15.158 |
| 88.95.111.130.137.155.160/ | 20 | -201.12 | 467.96 | 10.17 | BVAR DCSTc | 3 | -29.779 |
| 95.111.130.137.147.160/108 | 22 | -193.455 | 467.963 | 10.173 | BVAR DCSTc | 3 | -15.158 |
| 95.160/110 | 9 | -221.936 | 468.361 | 10.571 | BCST DCSTc | 2 | -32.891 |
| 95.111.137.155/108 | 14 | -212.278 | 468.908 | 11.118 | BVAR DCSTc | 3 | -15.158 |

| | | | | | | | |
|---|---|---|---|---|---|---|---|
| 88.95.111.137.147.155.160/ | 19 | -204.396 | 468.912 | 11.122 | BCST DVARc | 3 | -27.824 |
| 104.111.130/95.108 | 14 | -209.882 | 468.917 | 11.127 | BVAR DCSTc | 3 | -15.158 |
| 88.95.111.130.147.155.160/ | 19 | -203.945 | 469.009 | 11.219 | BVAR DCSTc | 3 | -31.603 |
| 95.130.137.147.160/108 | 20 | -202.164 | 469.152 | 11.362 | BVAR DCSTc | 3 | -15.158 |
| 95.111.130.147/108 | 14 | -212.235 | 469.223 | 11.433 | BVAR DCSTc | 3 | -15.158 |
| 95.111.130.137/108 | 15 | -210.506 | 469.403 | 11.612 | BVAR DCSTc | 3 | -15.158 |
| 88.160/110 | 10 | -220.669 | 469.423 | 11.633 | BVAR DCSTc | 3 | -53.71 |
| 95.137.147/108 | 12 | -215.862 | 469.5 | 11.71 | BVAR DCSTc | 3 | -15.158 |
| 137/110 | 7 | -225.346 | 469.513 | 11.722 | BCST DCSTc | 2 | -80.506 |
| 95.111.137.147.160/108 | 17 | -208.551 | 469.744 | 11.954 | BVAR DCSTc | 3 | -15.158 |
| 104.111.137.160/95.108 | 17 | -206.477 | 469.754 | 11.964 | BVAR DCSTc | 3 | -15.158 |
| 130.137.147.155.160/110 | 19 | -204.142 | 469.761 | 11.97 | BCST DCSTc | 2 | -80.506 |
| 95.137.147.160/108 | 16 | -210.106 | 469.931 | 12.141 | BVAR DCSTc | 3 | -15.158 |
| 104.137/95.108 | 12 | -213.893 | 470.025 | 12.235 | BVAR DCSTc | 3 | -15.158 |
| 130.137.147.160/110 | 17 | -208.826 | 470.157 | 12.367 | BCST DCSTc | 2 | -80.506 |
| 104.111.147/95.108 | 13 | -213.372 | 470.297 | 12.506 | BVAR DCSTc | 3 | -15.158 |
| 88.95.111.130.147.160/ | 17 | -209.307 | 470.447 | 12.657 | BVAR DCSTc | 3 | -50.332 |
| 130/110 | 7 | -224.843 | 470.497 | 12.706 | BCST DCSTc | 2 | -80.506 |
| 147/110 | 6 | -227.661 | 470.533 | 12.742 | BCST DCSTc | 2 | -80.506 |
| 95.111.130.137.160/108 | 20 | -202.804 | 470.564 | 12.774 | BVAR DCSTc | 3 | -15.158 |
| 88.95.111.137.147.160/ | 17 | -210.309 | 470.737 | 12.946 | BVAR DCSTc | 3 | -47.103 |
| 137.160/110 | 10 | -221.232 | 470.788 | 12.998 | BCST DCSTc | 2 | -80.506 |
| 95.111.130.155/108 | 14 | -212.293 | 470.905 | 13.115 | BVAR DCSTc | 3 | -15.158 |
| 95.130.147/108 | 12 | -215.573 | 470.909 | 13.119 | BVAR DCSTc | 3 | -15.158 |
| 95.130.147.160/108 | 16 | -209.628 | 470.925 | 13.135 | BVAR DCSTc | 3 | -15.158 |
| 88.111/108 | 9 | -223.532 | 470.95 | 13.159 | BVAR DCSTc | 3 | -44.968 |
| 104.110/ | 6 | -226.245 | 471.147 | 13.357 | BCSTc | 1 | -70.448 |
| 147.160/110 | 9 | -223.262 | 471.233 | 13.442 | BCST DCSTc | 2 | -80.506 |
| 95.111.130.147.160/108 | 19 | -205.277 | 471.351 | 13.56 | BVAR DCSTc | 3 | -15.158 |
| 95.111.155.160/108 | 14 | -213.798 | 471.396 | 13.606 | BVAR DCSTc | 3 | -15.158 |
| 104.130/95.108 | 12 | -213.615 | 471.46 | 13.67 | BVAR DCSTc | 3 | -15.158 |
| 95.137.155/108 | 12 | -216.057 | 471.477 | 13.687 | BVAR DCSTc | 3 | -15.158 |
| 95.111.130.137.147.155.160/ | 20 | -202.885 | 471.49 | 13.7 | BVAR DCSTc | 3 | -35.548 |
| 104.111.130.160/95.108 | 17 | -206.391 | 471.532 | 13.741 | BVAR DCSTc | 3 | -15.158 |
| 155/110 | 7 | -226.248 | 471.537 | 13.747 | BCST DCSTc | 2 | -80.506 |
| 104.111.155/95.108 | 13 | -213.269 | 471.671 | 13.881 | BVAR DCSTc | 3 | -15.158 |
| 95.137/110 | 10 | -222.21 | 471.736 | 13.946 | BCST DCSTc | 2 | -32.891 |
| 88.95.111.130.137.160/ | 18 | -208.014 | 471.747 | 13.957 | BVAR DCSTc | 3 | -50.039 |
| 130.160/110 | 10 | -220.726 | 471.76 | 13.969 | BCST DCSTc | 2 | -80.506 |
| 111/108 | 7 | -227.739 | 471.897 | 14.106 | BVAR DCSTc | 3 | -72.804 |
| 95.155.160/108 | 12 | -216.549 | 471.897 | 14.107 | BVAR DCSTc | 3 | -15.158 |
| 104.147/95.108 | 11 | -216.662 | 471.955 | 14.165 | BVAR DCSTc | 3 | -15.158 |
| 95.130.137.160/108 | 17 | -208.32 | 471.959 | 14.169 | BVAR DCSTc | 3 | -15.158 |

| | | | | | | | |
|---|---|---|---|---|---|---|---|
| 104.137.160/95.108 | 15 | -210.128 | 471.992 | 14.202 | BVAR DCSTc | 3 | -15.158 |
| 95.130.137/108 | 13 | -214.347 | 472.072 | 14.281 | BVAR DCSTc | 3 | -15.158 |
| 95.130.137.147.160/110 | 19 | -206.688 | 472.116 | 14.326 | BCST DCSTc | 2 | -32.891 |
| 95.111.147.155/108 | 13 | -215.716 | 472.152 | 14.361 | BVAR DCSTc | 3 | -15.158 |
| 88.108/ | 7 | -226.663 | 472.326 | 14.536 | BVAR DCSTc | 3 | -44.968 |
| 104.160/110 | 8 | -223.383 | 472.684 | 14.894 | BCSTc | 1 | -70.448 |
| 95.137.160/110 | 12 | -219.089 | 472.736 | 14.946 | BCST DCSTc | 2 | -32.891 |
| 88.137/110 | 11 | -220.944 | 472.798 | 15.008 | BVAR DCSTc | 3 | -53.71 |
| 104.111.147.160/95.108 | 16 | -209.839 | 472.827 | 15.037 | BVAR DCSTc | 3 | -15.158 |
| 95.130.137.147.155.160/110 | 21 | -202.568 | 472.847 | 15.057 | BCST DCSTc | 2 | -32.891 |
| 88.104.110/ | 10 | -220.315 | 472.879 | 15.089 | BVAR DCSTc | 3 | -41.22 |
| 95.130.155/108 | 12 | -215.793 | 472.937 | 15.147 | BVAR DCSTc | 3 | -15.158 |
| 95.147/110 | 9 | -224.65 | 472.995 | 15.205 | BCST DCSTc | 2 | -32.891 |
| 95.130/110 | 9 | -222.987 | 473.018 | 15.228 | BCST DCSTc | 2 | -32.891 |
| 104.110/95 | 9 | -222.456 | 473.048 | 15.258 | BCST DCSTc | 2 | -32.891 |
| 88.111.160/108 | 12 | -219.834 | 473.083 | 15.293 | BVAR DCSTc | 3 | -44.968 |
| 88.130.137.147.160/110 | 20 | -205.422 | 473.177 | 15.387 | BVAR DCSTc | 3 | -53.71 |
| 95.147.160/110 | 11 | -221.176 | 473.294 | 15.504 | BCST DCSTc | 2 | -32.891 |
| 130.137.160/110 | 15 | -213.923 | 473.328 | 15.538 | BCST DCSTc | 2 | -80.506 |
| 95.147.155/108 | 11 | -218.797 | 473.345 | 15.555 | BVAR DCSTc | 3 | -15.158 |
| 95.155/110 | 9 | -224.121 | 473.517 | 15.727 | BCST DCSTc | 2 | -32.891 |
| 104.130.160/95.108 | 15 | -209.902 | 473.525 | 15.735 | BVAR DCSTc | 3 | -15.158 |
| 104.155/95.108 | 11 | -216.729 | 473.68 | 15.89 | BVAR DCSTc | 3 | -15.158 |
| 104.147.160/95.108 | 14 | -212.784 | 473.69 | 15.9 | BVAR DCSTc | 3 | -15.158 |
| 104.130.137.147.155.160/95.108 | 24 | -194.603 | 473.702 | 15.912 | BVAR DCSTc | 3 | -15.158 |
| 88.137.160/110 | 13 | -217.822 | 473.798 | 16.008 | BVAR DCSTc | 3 | -53.71 |
| 95.130.160/110 | 12 | -218.63 | 473.802 | 16.012 | BCST DCSTc | 2 | -32.891 |
| 104.111.137.147/95.108 | 16 | -210.054 | 473.808 | 16.018 | BVAR DCSTc | 3 | -15.158 |
| 88.130.137.147.155.160/110 | 22 | -201.301 | 473.908 | 16.118 | BVAR DCSTc | 3 | -53.71 |
| 108/ | 5 | -231.2 | 473.934 | 16.143 | BVAR DCSTc | 3 | -72.804 |
| 111/ | 4 | -232.514 | 474.016 | 16.226 | BVARc | 2 | -199.605 |
| 88.147/110 | 10 | -223.384 | 474.057 | 16.267 | BVAR DCSTc | 3 | -53.71 |
| 88.111.137/108 | 12 | -220.043 | 474.061 | 16.27 | BVAR DCSTc | 3 | -44.968 |
| 88.130/110 | 10 | -221.72 | 474.08 | 16.29 | BVAR DCSTc | 3 | -53.71 |
| 88.104.130.137.147.160/95 | 20 | -203.43 | 474.18 | 16.39 | BVAR DCSTc | 3 | -61.169 |
| 88.160/108 | 10 | -222.871 | 474.215 | 16.424 | BVAR DCSTc | 3 | -44.968 |
| 88.147.160/110 | 12 | -219.91 | 474.356 | 16.566 | BVAR DCSTc | 3 | -53.71 |
| 111.160/108 | 10 | -224.21 | 474.37 | 16.58 | BVAR DCSTc | 3 | -72.804 |
| 111.137.160/ | 10 | -222.948 | 474.404 | 16.613 | BCST DCSTc | 2 | -147.719 |
| 88.104.111/108 | 11 | -219.093 | 474.473 | 16.682 | BVAR DCSTc | 3 | -31.012 |
| 88.155/110 | 10 | -222.854 | 474.578 | 16.788 | BVAR DCSTc | 3 | -53.71 |
| 104.160/95.110 | 11 | -219.593 | 474.585 | 16.795 | BCST DCSTc | 2 | -32.891 |
| 88.104.111.130.137.155.160/95 | 22 | -199.023 | 474.674 | 16.884 | BVAR DCSTc | 3 | -29.779 |

| | | | | | | | |
|---|---|---|---|---|---|---|---|
| 104.111.130.137.147.160/95.108 | 24 | -191.358 | 474.677 | 16.887 | BVAR DCSTc | 3 | -15.158 |
| 111.160/ | 7 | -228.129 | 474.712 | 16.922 | BVARc | 2 | -171.193 |
| 88.130.160/110 | 13 | -217.364 | 474.864 | 17.074 | BVAR DCSTc | 3 | -53.71 |
| 95.111.137.155.160/108 | 17 | -210.393 | 474.905 | 17.114 | BVAR DCSTc | 3 | -15.158 |
| 88.104.160/110 | 12 | -217.697 | 474.907 | 17.116 | BVAR DCSTc | 3 | -41.22 |
| 104.111.137.155/95.108 | 16 | -209.951 | 475.163 | 17.373 | BVAR DCSTc | 3 | -15.158 |
| 95.111.130.147.155.160/108 | 21 | -201.873 | 475.179 | 17.389 | BVAR DCSTc | 3 | -15.158 |
| 95.111.137.160/ | 12 | -220.746 | 475.226 | 17.436 | BCST DCSTc | 2 | -100.134 |
| 137.147/110 | 9 | -224.987 | 475.245 | 17.454 | BCST DCSTc | 2 | -80.506 |
| 95.111.130.137.147.160/ | 18 | -209.774 | 475.267 | 17.477 | BVAR DCSTc | 3 | -55.803 |
| 104.130.137.147.160/95.108 | 22 | -199.797 | 475.327 | 17.537 | BVAR DCSTc | 3 | -15.158 |
| 95.111.137.147.155/108 | 16 | -212.192 | 475.342 | 17.551 | BVAR DCSTc | 3 | -15.158 |
| 111.137/108 | 10 | -224.417 | 475.342 | 17.552 | BVAR DCSTc | 3 | -72.804 |
| 95.130.137.160/110 | 17 | -211.844 | 475.403 | 17.613 | BCST DCSTc | 2 | -32.891 |
| 104.111.130.147/95.108 | 16 | -209.883 | 475.428 | 17.638 | BVAR DCSTc | 3 | -15.158 |
| 111.137/ | 7 | -228.251 | 475.524 | 17.733 | BVARc | 2 | -177.047 |
| 95.111.137.147.155.160/108 | 21 | -201.952 | 475.624 | 17.834 | BVAR DCSTc | 3 | -15.158 |
| 88.104.111.137.147.155.160/95 | 21 | -202.299 | 475.626 | 17.836 | BCST DVARc | 3 | -27.824 |
| 95.111.160/ | 9 | -226.007 | 475.641 | 17.85 | BCST DCSTc | 2 | -123.688 |
| 104.111.130.137/95.108 | 17 | -208.18 | 475.661 | 17.871 | BVAR DCSTc | 3 | -15.158 |
| 88.104.111.130.147.155.160/95 | 21 | -201.847 | 475.723 | 17.933 | BVAR DCSTc | 3 | -31.603 |
| 95.111/ | 6 | -230.819 | 475.768 | 17.978 | BVARc | 2 | -152.526 |
| 88.104.108/ | 9 | -222.225 | 475.849 | 18.059 | BVAR DCSTc | 3 | -31.012 |
| 160/108 | 8 | -227.422 | 475.851 | 18.061 | BVAR DCSTc | 3 | -72.804 |
| 95.111.130.137.147/108 | 17 | -210.415 | 475.871 | 18.08 | BVAR DCSTc | 3 | -15.158 |
| 137.147.160/110 | 12 | -220.552 | 475.918 | 18.128 | BCST DCSTc | 2 | -80.506 |
| 155.160/110 | 9 | -224.855 | 476.002 | 18.212 | BCST DCSTc | 2 | -80.506 |
| 88.111.130/108 | 12 | -220.038 | 476.029 | 18.238 | BVAR DCSTc | 3 | -44.968 |
| 104.137/110 | 8 | -224.805 | 476.095 | 18.305 | BCSTc | 1 | -70.448 |
| 95.137.155.160/108 | 15 | -213.634 | 476.15 | 18.36 | BVAR DCSTc | 3 | -15.158 |
| 95.111.137.147.160/ | 14 | -217.93 | 476.15 | 18.36 | BCST DCSTc | 2 | -78.023 |
| 130.147/110 | 9 | -224.449 | 476.155 | 18.365 | BCST DCSTc | 2 | -80.506 |
| 104.137.147/95.108 | 14 | -213.741 | 476.168 | 18.378 | BVAR DCSTc | 3 | -15.158 |
| 88.111.137.160/108 | 15 | -216.359 | 476.318 | 18.527 | BVAR DCSTc | 3 | -44.968 |
| 104.130.137.147.155.160/110 | 20 | -203.601 | 476.343 | 18.553 | BCSTc | 1 | -70.448 |
| 95.111.130.137.155.160/108 | 22 | -199.512 | 476.344 | 18.554 | BVAR DCSTc | 3 | -15.158 |
| 111.130.160/ | 10 | -222.931 | 476.357 | 18.567 | BCST DCSTc | 2 | -151.932 |
| 95.111.130.160/ | 12 | -220.339 | 476.384 | 18.594 | BCST DCSTc | 2 | -103.956 |
| 104.137.147.160/95.108 | 18 | -207.894 | 476.416 | 18.626 | BVAR DCSTc | 3 | -15.158 |
| 104.111.137.147.160/95.108 | 19 | -206.453 | 476.458 | 18.668 | BVAR DCSTc | 3 | -15.158 |
| 88.130.137.160/110 | 18 | -210.577 | 476.465 | 18.675 | BVAR DCSTc | 3 | -53.71 |
| 95.111.130.147.160/ | 14 | -217.158 | 476.556 | 18.766 | BCST DCSTc | 2 | -81.481 |
| 88.104.111.160/108 | 14 | -215.395 | 476.606 | 18.816 | BVAR DCSTc | 3 | -31.012 |

| | | | | | | | |
|---|---|---|---|---|---|---|---|
| 88.137/108 | 10 | -223.802 | 476.644 | 18.854 | BVAR DCSTc | 3 | -44.968 |
| 104.111.130.155/95.108 | 16 | -209.73 | 476.689 | 18.899 | BVAR DCSTc | 3 | -15.158 |
| 104.130.137.147.160/110 | 18 | -208.286 | 476.74 | 18.95 | BCSTc | 1 | -70.448 |
| 95.111.130.155.160/108 | 17 | -210.361 | 476.747 | 18.957 | BVAR DCSTc | 3 | -15.158 |
| 130.147.160/110 | 12 | -220.015 | 476.817 | 19.027 | BCST DCSTc | 2 | -80.506 |
| 104.111.130.137.160/95.108 | 22 | -200.483 | 476.831 | 19.041 | BVAR DCSTc | 3 | -15.158 |
| 111.130/108 | 10 | -224.242 | 476.97 | 19.18 | BVAR DCSTc | 3 | -72.804 |
| 104.130/110 | 8 | -224.302 | 477.079 | 19.289 | BCSTc | 1 | -70.448 |
| 137.155/110 | 10 | -223.979 | 477.115 | 19.324 | BCST DCSTc | 2 | -80.506 |
| 104.147/110 | 7 | -227.12 | 477.115 | 19.325 | BCSTc | 1 | -70.448 |
| 104.130.147.160/95.108 | 18 | -207.27 | 477.119 | 19.328 | BVAR DCSTc | 3 | -15.158 |
| 88.104.111.130.147.160/95 | 19 | -207.209 | 477.161 | 19.371 | BVAR DCSTc | 3 | -50.332 |
| 95.111.147.160/ | 11 | -223.556 | 477.219 | 19.428 | BCST DCSTc | 2 | -101.943 |
| 104.111.130.147.160/95.108 | 21 | -202.756 | 477.218 | 19.428 | BVAR DCSTc | 3 | -15.158 |
| 111.137.147.160/ | 12 | -221.126 | 477.224 | 19.434 | BCST DCSTc | 2 | -126.603 |
| 88.111.147/108 | 11 | -223.445 | 477.243 | 19.453 | BVAR DCSTc | 3 | -44.968 |
| 95.111.130.147.155/108 | 16 | -212.216 | 477.322 | 19.532 | BVAR DCSTc | 3 | -15.158 |
| 95.137.147/110 | 11 | -222.913 | 477.331 | 19.541 | BCST DCSTc | 2 | -32.891 |
| 104.111/108 | 9 | -225.202 | 477.336 | 19.545 | BVAR DCSTc | 3 | -60.749 |
| 104.137.160/110 | 11 | -220.691 | 477.371 | 19.581 | BCSTc | 1 | -70.448 |
| 95.130.155.160/108 | 15 | -213.273 | 477.409 | 19.619 | BVAR DCSTc | 3 | -15.158 |
| 95.111.130.137.155/108 | 17 | -210.443 | 477.443 | 19.653 | BVAR DCSTc | 3 | -15.158 |
| 88.104.111.137.147.160/95 | 19 | -208.211 | 477.451 | 19.66 | BVAR DCSTc | 3 | -47.103 |
| 104.111.155.160/95.108 | 16 | -211.379 | 477.466 | 19.676 | BVAR DCSTc | 3 | -15.158 |
| 95.137.147.155/108 | 14 | -215.823 | 477.471 | 19.681 | BVAR DCSTc | 3 | -15.158 |
| 95.111.130.137.160/ | 15 | -215.825 | 477.54 | 19.75 | BCST DCSTc | 2 | -81.148 |
| 88.104.111.137/108 | 14 | -215.604 | 477.584 | 19.793 | BVAR DCSTc | 3 | -31.012 |
| 104.130.147/95.108 | 14 | -213.467 | 477.607 | 19.817 | BVAR DCSTc | 3 | -15.158 |
| 95.130.137.147.160/ | 15 | -215.202 | 477.71 | 19.92 | BCST DCSTc | 2 | -94.141 |
| 95.137.147.160/110 | 14 | -218.334 | 477.717 | 19.926 | BCST DCSTc | 2 | -32.891 |
| 111.130/ | 7 | -228.36 | 477.733 | 19.943 | BVARc | 2 | -181.387 |
| 88.104.160/108 | 12 | -218.432 | 477.738 | 19.947 | BVAR DCSTc | 3 | -31.012 |
| 111.147.160/ | 9 | -226.429 | 477.753 | 19.963 | BCST DCSTc | 2 | -150.199 |
| 130.137.155.160/110 | 17 | -211.942 | 477.788 | 19.998 | BCST DCSTc | 2 | -80.506 |
| 111.137.160/108 | 13 | -220.837 | 477.807 | 20.017 | BVAR DCSTc | 3 | -72.804 |
| 104.147.160/110 | 10 | -222.722 | 477.815 | 20.025 | BCSTc | 1 | -70.448 |
| 104.137.155/95.108 | 14 | -213.825 | 477.922 | 20.132 | BVAR DCSTc | 3 | -15.158 |
| 147.155/110 | 8 | -227.355 | 477.954 | 20.164 | BCST DCSTc | 2 | -80.506 |
| 130.137/110 | 10 | -223.544 | 477.959 | 20.169 | BCST DCSTc | 2 | -80.506 |
| 104.137/95.110 | 11 | -221.016 | 477.996 | 20.206 | BCST DCSTc | 2 | -32.891 |
| 130.155/110 | 9 | -224.598 | 478.04 | 20.249 | BCST DCSTc | 2 | -80.506 |
| 137/108 | 8 | -228.252 | 478.078 | 20.288 | BVAR DCSTc | 3 | -72.804 |
| 95.130.137.147/108 | 15 | -214.12 | 478.084 | 20.293 | BVAR DCSTc | 3 | -15.158 |

| | | | | | | | |
|---|---|---|---|---|---|---|---|
| 88.104.111.130.137.147.155.160/ | 22 | -200.827 | 478.112 | 20.322 | BCST DVARc | 3 | -46.058 |
| 104.155/110 | 8 | -225.707 | 478.12 | 20.33 | BCSTc | 1 | -70.448 |
| 88.130/108 | 10 | -223.55 | 478.13 | 20.34 | BVAR DCSTc | 3 | -44.968 |
| 104.111.147.155/95.108 | 15 | -213.266 | 478.16 | 20.37 | BVAR DCSTc | 3 | -15.158 |
| 104.130.137.160/95.108 | 19 | -205.969 | 478.166 | 20.376 | BVAR DCSTc | 3 | -15.158 |
| 104.111.130.137.147.155.160/95 | 22 | -200.788 | 478.204 | 20.414 | BVAR DCSTc | 3 | -35.548 |
| 88.111.137.160/ | 12 | -221.62 | 478.211 | 20.421 | BCST DCSTc | 2 | -123.093 |
| 95.130.137.155.160/108 | 20 | -205.986 | 478.231 | 20.44 | BVAR DCSTc | 3 | -15.158 |
| 88.95.137.147.155.160/ | 17 | -212.955 | 478.23 | 20.44 | BVAR DCSTc | 3 | -69.292 |
| 95.137.147.155.160/108 | 19 | -208.792 | 478.242 | 20.452 | BVAR DCSTc | 3 | -15.158 |
| 104.130.137.147.155.160/95.110 | 23 | -199.812 | 478.244 | 20.454 | BCST DCSTc | 2 | -32.891 |
| 95/ | 4 | -234.49 | 478.247 | 20.457 | BVARc | 2 | -189.106 |
| 88.111.130.160/108 | 15 | -216.356 | 478.263 | 20.473 | BVAR DCSTc | 3 | -44.968 |
| 88.104.137/110 | 13 | -217.971 | 478.282 | 20.492 | BVAR DCSTc | 3 | -41.22 |
| 95.111.137/ | 9 | -227.061 | 478.312 | 20.522 | BVARc | 2 | -130.474 |
| 88.137.160/108 | 13 | -219.9 | 478.336 | 20.546 | BVAR DCSTc | 3 | -44.968 |
| 104.130.160/110 | 11 | -220.185 | 478.342 | 20.552 | BCSTc | 1 | -70.448 |
| 111.147/108 | 9 | -227.732 | 478.35 | 20.559 | BVAR DCSTc | 3 | -72.804 |
| 95.160/ | 7 | -229.811 | 478.351 | 20.561 | BVARc | 2 | -160.401 |
| 88.137.147/110 | 12 | -221.646 | 478.393 | 20.602 | BVAR DCSTc | 3 | -53.71 |
| 88.95.130.147.155.160/ | 17 | -212.078 | 478.399 | 20.608 | BVAR DCSTc | 3 | -72.646 |
| 104.155.160/95.108 | 14 | -214.35 | 478.408 | 20.618 | BVAR DCSTc | 3 | -15.158 |
| 88.104.111.130.137.160/95 | 20 | -205.916 | 478.461 | 20.671 | BVAR DCSTc | 3 | -50.039 |
| 95.130.147/110 | 11 | -222.486 | 478.463 | 20.673 | BCST DCSTc | 2 | -32.891 |
| 95.130.147.160/110 | 14 | -217.784 | 478.588 | 20.798 | BCST DCSTc | 2 | -32.891 |
| 95.155.160/110 | 11 | -223.034 | 478.595 | 20.805 | BCST DCSTc | 2 | -32.891 |
| 88.147/108 | 9 | -226.584 | 478.598 | 20.808 | BVAR DCSTc | 3 | -44.968 |
| 104.130.137/95.108 | 15 | -212.17 | 478.626 | 20.836 | BVAR DCSTc | 3 | -15.158 |
| 104.130.137.147.160/95.110 | 21 | -204.496 | 478.641 | 20.851 | BCST DCSTc | 2 | -32.891 |
| 88.111.130.137.147.160/ | 18 | -211.148 | 478.659 | 20.869 | BVAR DCSTc | 3 | -79.263 |
| 88.104.130.137.147.160/110 | 22 | -202.449 | 478.661 | 20.871 | BVAR DCSTc | 3 | -41.22 |
| 88.111.160/ | 9 | -226.884 | 478.664 | 20.874 | BCST DCSTc | 2 | -146.651 |
| 88.137.147.160/110 | 15 | -217.068 | 478.778 | 20.988 | BVAR DCSTc | 3 | -53.71 |
| 111.130.147.160/ | 12 | -220.919 | 478.791 | 21.001 | BCST DCSTc | 2 | -130.626 |
| 95.137.147.160/ | 12 | -221.799 | 478.825 | 21.034 | BCST DCSTc | 2 | -114.801 |
| 95.130.147.155/108 | 14 | -215.516 | 478.839 | 21.049 | BVAR DCSTc | 3 | -15.158 |
| **No shift** | **2** | **-237.347** | **478.841** | **21.051** | – | – | – |
| 111.130.137.160/ | 13 | -219.163 | 478.897 | 21.106 | BCST DCSTc | 2 | -129.87 |
| 95.137.160/ | 10 | -225.083 | 478.941 | 21.15 | BCST DCSTc | 2 | -137.379 |
| 104.130/95.110 | 11 | -220.513 | 478.98 | 21.19 | BCST DCSTc | 2 | -32.891 |
| 88.111.155/108 | 11 | -223.526 | 478.986 | 21.195 | BVAR DCSTc | 3 | -44.968 |
| 104.147/95.110 | 10 | -223.331 | 479.016 | 21.226 | BCST DCSTc | 2 | -32.891 |
| 95.137.155/110 | 12 | -221.815 | 479.021 | 21.231 | BCST DCSTc | 2 | -32.891 |

| | | | | | | | |
|---|---|---|---|---|---|---|---|
| 88.111.130.137.147.155.160/ | 20 | -206.997 | 479.022 | 21.232 | BVAR DCSTc | 3 | -61.745 |
| 160/ | 5 | -232.834 | 479.267 | 21.476 | BVARc | 2 | -208.808 |
| 104.137.160/95.110 | 14 | -216.902 | 479.271 | 21.481 | BCST DCSTc | 2 | -32.891 |
| 104.130.155/95.108 | 14 | -213.51 | 479.28 | 21.49 | BVAR DCSTc | 3 | -15.158 |
| 88.104.137.160/110 | 15 | -214.85 | 479.282 | 21.492 | BVAR DCSTc | 3 | -41.22 |
| 95.130.147.155.160/108 | 19 | -208.408 | 479.335 | 21.544 | BVAR DCSTc | 3 | -15.158 |
| 104.108/ | 7 | -228.663 | 479.373 | 21.583 | BVAR DCSTc | 3 | -60.749 |
| 88.104.130.137.147.155.160/110 | 24 | -198.329 | 479.392 | 21.602 | BVAR DCSTc | 3 | -41.22 |
| 88.111.147.160/108 | 14 | -219.739 | 479.427 | 21.637 | BVAR DCSTc | 3 | -44.968 |
| 111.130.137.147.160/ | 16 | -214.965 | 479.482 | 21.692 | BVAR DCSTc | 3 | -106.378 |
| 130/108 | 8 | -227.974 | 479.513 | 21.723 | BVAR DCSTc | 3 | -72.804 |
| 88.130.147/110 | 12 | -221.22 | 479.525 | 21.735 | BVAR DCSTc | 3 | -53.71 |
| 95.147.155.160/108 | 15 | -215.955 | 479.535 | 21.745 | BVAR DCSTc | 3 | -15.158 |
| 88.104.147/110 | 12 | -220.411 | 479.541 | 21.751 | BVAR DCSTc | 3 | -41.22 |
| 88.104.111.130/108 | 14 | -215.599 | 479.552 | 21.761 | BVAR DCSTc | 3 | -31.012 |
| 88.104.130/110 | 12 | -218.748 | 479.564 | 21.774 | BVAR DCSTc | 3 | -41.22 |
| 111.130.160/108 | 13 | -220.75 | 479.585 | 21.794 | BVAR DCSTc | 3 | -72.804 |
| 111.147/ | 6 | -232.102 | 479.618 | 21.828 | BVARc | 2 | -179.899 |
| 88.95.130.137.155.160/ | 18 | -210.849 | 479.618 | 21.828 | BVAR DCSTc | 3 | -72.417 |
| 95.130.147.160/ | 12 | -221.208 | 479.629 | 21.839 | BCST DCSTc | 2 | -118.441 |
| 88.130.147.160/110 | 15 | -216.517 | 479.65 | 21.859 | BVAR DCSTc | 3 | -53.71 |
| 88.155.160/110 | 12 | -221.768 | 479.657 | 21.867 | BVAR DCSTc | 3 | -53.71 |
| 88.111.130.160/ | 12 | -221.363 | 479.678 | 21.888 | BCST DCSTc | 2 | -127.066 |
| 88.130.160/108 | 13 | -219.59 | 479.702 | 21.912 | BVAR DCSTc | 3 | -44.968 |
| 104.147.160/95.110 | 13 | -218.933 | 479.716 | 21.926 | BCST DCSTc | 2 | -32.891 |
| 111.155/108 | 9 | -227.629 | 479.724 | 21.934 | BVAR DCSTc | 3 | -72.804 |
| 104.147.155/95.108 | 13 | -216.535 | 479.731 | 21.941 | BVAR DCSTc | 3 | -15.158 |
| 88.95.137.147.160/ | 14 | -219.015 | 479.736 | 21.945 | BCST DCSTc | 2 | -88.719 |
| 88.95.130.147.160/ | 14 | -218.032 | 479.747 | 21.957 | BCST DCSTc | 2 | -91.967 |
| 104.111.160/108 | 12 | -221.674 | 479.809 | 22.019 | BVAR DCSTc | 3 | -60.749 |
| 88.104.147.160/110 | 14 | -216.938 | 479.84 | 22.05 | BVAR DCSTc | 3 | -41.22 |
| 88.104.111.137.160/108 | 17 | -211.92 | 479.84 | 22.05 | BVAR DCSTc | 3 | -31.012 |
| 104.130.137.160/110 | 16 | -213.383 | 479.911 | 22.12 | BCSTc | 1 | -70.448 |
| 88.147.160/108 | 12 | -222.495 | 479.911 | 22.121 | BVAR DCSTc | 3 | -44.968 |
| 104.111.130.147.155.160/95.108 | 23 | -198.796 | 479.935 | 22.145 | BVAR DCSTc | 3 | -15.158 |
| 95.130.137/110 | 14 | -219.056 | 479.969 | 22.179 | BCST DCSTc | 2 | -32.891 |
| 95.111.130/ | 9 | -226.913 | 480.002 | 22.212 | BVARc | 2 | -134.557 |
| 147/108 | 7 | -231.022 | 480.008 | 22.218 | BVAR DCSTc | 3 | -72.804 |
| 104.155/95.110 | 11 | -221.918 | 480.021 | 22.23 | BCST DCSTc | 2 | -32.891 |
| 95.147.155/110 | 11 | -224.138 | 480.039 | 22.248 | BCST DCSTc | 2 | -32.891 |
| 137.160/108 | 11 | -224.488 | 480.045 | 22.255 | BVAR DCSTc | 3 | -72.804 |
| 95.130.137.155.160/110 | 19 | -209.96 | 480.058 | 22.268 | BCST DCSTc | 2 | -32.891 |
| 88.111/ | 6 | -232.324 | 480.062 | 22.272 | BVARc | 2 | -176.117 |

| | | | | | | | |
|---|---|---|---|---|---|---|---|
| 88.104.155/110 | 12 | -219.882 | 480.062 | 22.272 | BVAR DCSTc | 3 | -41.22 |
| 95.130.137.155/108 | 15 | -214.324 | 480.073 | 22.283 | BVAR DCSTc | 3 | -15.158 |
| 88.137.155/110 | 13 | -220.549 | 480.083 | 22.293 | BVAR DCSTc | 3 | -53.71 |
| 95.130.160/ | 10 | -224.663 | 480.09 | 22.3 | BCST DCSTc | 2 | -141.19 |
| 88.111.137.147.160/ | 14 | -219.31 | 480.092 | 22.302 | BCST DCSTc | 2 | -101.488 |
| 88.95.111.147.155.160/ | 16 | -216.247 | 480.113 | 22.323 | BVAR DCSTc | 3 | -57.969 |
| 111.137.147/ | 9 | -227.343 | 480.146 | 22.356 | BCST DCSTc | 2 | -156.845 |
| 88.104.137/108 | 12 | -219.364 | 480.167 | 22.377 | BVAR DCSTc | 3 | -31.012 |
| 95.111.147.155.160/108 | 18 | -211.919 | 480.225 | 22.435 | BVAR DCSTc | 3 | -15.158 |
| 104.130.160/95.110 | 14 | -216.396 | 480.243 | 22.453 | BCST DCSTc | 2 | -32.891 |
| 95.130.137.160/ | 13 | -219.733 | 480.293 | 22.503 | BCST DCSTc | 2 | -117.966 |
| 130.137.147/110 | 14 | -219 | 480.306 | 22.515 | BCST DCSTc | 2 | -80.506 |
| 88.104.130.160/110 | 15 | -214.391 | 480.348 | 22.558 | BVAR DCSTc | 3 | -41.22 |
| 137.147.155.160/110 | 16 | -216.051 | 480.407 | 22.616 | BCST DCSTc | 2 | -80.506 |
| 88.111.137.147/108 | 14 | -219.963 | 480.426 | 22.635 | BVAR DCSTc | 3 | -44.968 |
| 95.130.155/110 | 12 | -221.547 | 480.458 | 22.668 | BCST DCSTc | 2 | -32.891 |
| 95.147.160/ | 9 | -227.664 | 480.492 | 22.702 | BCST DCSTc | 2 | -138.96 |
| 137.160/ | 8 | -228.498 | 480.621 | 22.831 | BCST DCSTc | 2 | -186.178 |
| 88.95.130.137.160/ | 16 | -215.463 | 480.646 | 22.856 | BVAR DCSTc | 3 | -90.398 |
| 88.155/108 | 9 | -226.812 | 480.647 | 22.857 | BVAR DCSTc | 3 | -44.968 |
| 88.111.147.160/ | 11 | -224.704 | 480.761 | 22.971 | BCST DCSTc | 2 | -125.177 |
| 88.104.111.147/108 | 13 | -219.006 | 480.766 | 22.976 | BVAR DCSTc | 3 | -31.012 |
| 104.111.137/108 | 12 | -221.88 | 480.781 | 22.991 | BVAR DCSTc | 3 | -60.749 |
| 137.155.160/110 | 12 | -222.245 | 480.877 | 23.087 | BCST DCSTc | 2 | -80.506 |
| 111.147.160/108 | 12 | -224.198 | 480.88 | 23.09 | BVAR DCSTc | 3 | -72.804 |
| 88.111.130.147.160/ | 14 | -218.765 | 480.972 | 23.182 | BCST DCSTc | 2 | -105.174 |
| 88.130.137/110 | 15 | -217.789 | 481.031 | 23.241 | BVAR DCSTc | 3 | -53.71 |
| 88.130.137.147.155.160/108 | 22 | -204.879 | 481.054 | 23.264 | BVAR DCSTc | 3 | -44.968 |
| 88.95.111.137.160/ | 15 | -219.078 | 481.075 | 23.285 | BVAR DCSTc | 3 | -75.167 |
| 95.130.137.147.155.160/ | 18 | -211.581 | 481.082 | 23.292 | BVAR DCSTc | 3 | -77.153 |
| 88.147.155/110 | 12 | -222.871 | 481.1 | 23.31 | BVAR DCSTc | 3 | -53.71 |
| 88.130.137.155.160/110 | 20 | -208.693 | 481.119 | 23.329 | BVAR DCSTc | 3 | -53.71 |
| 88.111.130.137.147.160/108 | 22 | -201.18 | 481.123 | 23.332 | BVAR DCSTc | 3 | -44.968 |
| 88.95.111.147.160/ | 14 | -220.963 | 481.136 | 23.346 | BVAR DCSTc | 3 | -76.052 |
| 130.147.155.160/110 | 16 | -215.511 | 481.162 | 23.372 | BCST DCSTc | 2 | -80.506 |
| 88.111.137/ | 9 | -227.885 | 481.231 | 23.441 | BCST DCSTc | 2 | -153.384 |
| 104.160/108 | 10 | -224.885 | 481.29 | 23.5 | BVAR DCSTc | 3 | -60.749 |
| 88.95.111.130.160/ | 14 | -219.545 | 481.332 | 23.542 | BCST DCSTc | 2 | -79.865 |
| 137/ | 5 | -233.588 | 481.342 | 23.552 | BVARc | 2 | -215.294 |
| 104.111.137.155.160/95.108 | 19 | -208.162 | 481.353 | 23.563 | BVAR DCSTc | 3 | -15.158 |
| 95.137/ | 7 | -231.094 | 481.485 | 23.695 | BVARc | 2 | -167.416 |
| 88.130.155/110 | 13 | -220.281 | 481.519 | 23.729 | BVAR DCSTc | 3 | -53.71 |
| 95.111.147/ | 8 | -230.485 | 481.546 | 23.756 | BVARc | 2 | -132.898 |

| | | | | | | | |
|---|---|---|---|---|---|---|---|
| 130.155.160/110 | 12 | -221.597 | 481.558 | 23.768 | BCST DCSTc | 2 | -80.506 |
| 88.111.130.137.160/ | 16 | -216.006 | 481.563 | 23.773 | BVAR DCSTc | 3 | -103.415 |
| 130.160/108 | 11 | -224.262 | 481.578 | 23.788 | BVAR DCSTc | 3 | -72.804 |
| 95.111.137.147/ | 11 | -225.486 | 481.638 | 23.848 | BCST DCSTc | 2 | -109.605 |
| 88.104.130/108 | 12 | -219.111 | 481.653 | 23.863 | BVAR DCSTc | 3 | -31.012 |
| 111.155/ | 6 | -232.354 | 481.714 | 23.924 | BVARc | 2 | -186.078 |
| 155/108 | 7 | -231.089 | 481.733 | 23.943 | BVAR DCSTc | 3 | -72.804 |
| 147.160/108 | 10 | -227.144 | 481.743 | 23.953 | BVAR DCSTc | 3 | -72.804 |
| 130.137.147.155.160/108 | 20 | -208.963 | 481.755 | 23.965 | BVAR DCSTc | 3 | -72.804 |
| 104.111.137.147.155/95.108 | 18 | -209.96 | 481.786 | 23.996 | BVAR DCSTc | 3 | -15.158 |
| 88.104.111.130.160/108 | 17 | -211.918 | 481.786 | 23.996 | BVAR DCSTc | 3 | -31.012 |
| 104.130.137.160/95.110 | 19 | -209.593 | 481.811 | 24.021 | BCST DCSTc | 2 | -32.891 |
| 104.137.147/110 | 10 | -224.446 | 481.827 | 24.037 | BCSTc | 1 | -70.448 |
| 111.130.137/ | 10 | -225.397 | 481.855 | 24.065 | BCST DCSTc | 2 | -160.13 |
| 88.104.137.160/108 | 15 | -215.461 | 481.859 | 24.069 | BVAR DCSTc | 3 | -31.012 |
| 111.137.147/108 | 12 | -224.414 | 481.861 | 24.071 | BVAR DCSTc | 3 | -72.804 |
| 88.95.111.130.155.160/ | 17 | -214.347 | 481.914 | 24.124 | BVAR DCSTc | 3 | -61.3 |
| 104.111.137.160/95 | 14 | -218.649 | 481.94 | 24.15 | BCST DCSTc | 2 | -100.134 |
| 88.104.130.137.160/110 | 20 | -207.605 | 481.949 | 24.159 | BVAR DCSTc | 3 | -41.22 |
| 104.111.137.160/ | 12 | -221.703 | 481.951 | 24.161 | BCST DCSTc | 2 | -136.956 |
| 104.111.130.137.147.160/95 | 20 | -207.676 | 481.981 | 24.191 | BVAR DCSTc | 3 | -55.803 |
| 88.111.137.155/108 | 14 | -220.003 | 482.067 | 24.277 | BVAR DCSTc | 3 | -44.968 |
| 104.111.130.137.155.160/95.108 | 24 | -196.939 | 482.106 | 24.316 | BVAR DCSTc | 3 | -15.158 |
| 88.104.147/108 | 11 | -222.145 | 482.121 | 24.331 | BVAR DCSTc | 3 | -31.012 |
| 111.130.147/ | 9 | -227.371 | 482.192 | 24.402 | BCST DCSTc | 2 | -161.104 |
| 88.95.111.137.155.160/ | 17 | -215.403 | 482.239 | 24.449 | BVAR DCSTc | 3 | -58.125 |
| 130.160/ | 8 | -228.32 | 482.259 | 24.469 | BCST DCSTc | 2 | -190.231 |
| 104.111.130.137.147/95.108 | 19 | -208.179 | 482.308 | 24.518 | BVAR DCSTc | 3 | -15.158 |
| 88.130.137.147.160/108 | 20 | -209.889 | 482.312 | 24.521 | BVAR DCSTc | 3 | -44.968 |
| 104.111.137.147.155.160/95.108 | 23 | -199.855 | 482.338 | 24.548 | BVAR DCSTc | 3 | -15.158 |
| 104.111.160/95 | 11 | -223.909 | 482.355 | 24.564 | BCST DCSTc | 2 | -123.688 |
| 95.130.137.147/110 | 16 | -216.913 | 482.365 | 24.575 | BCST DCSTc | 2 | -32.891 |
| 88.104.111.130.137.147.160/ | 20 | -207.746 | 482.367 | 24.577 | BVAR DCSTc | 3 | -66.344 |
| 88.111.130.147/108 | 14 | -219.96 | 482.382 | 24.592 | BVAR DCSTc | 3 | -44.968 |
| 104.111.130/108 | 12 | -221.705 | 482.409 | 24.619 | BVAR DCSTc | 3 | -60.749 |
| 104.111.160/ | 9 | -226.987 | 482.452 | 24.662 | BCST DCSTc | 2 | -160.534 |
| 104.111/95 | 8 | -228.721 | 482.482 | 24.692 | BVARc | 2 | -152.526 |
| 104.137.147.160/110 | 13 | -220.012 | 482.501 | 24.711 | BCSTc | 1 | -70.448 |
| 137.147.160/ | 10 | -226.224 | 482.502 | 24.712 | BCST DCSTc | 2 | -164.609 |
| 88.104.111.155/108 | 13 | -219.088 | 482.509 | 24.718 | BVAR DCSTc | 3 | -31.012 |
| 95.137.147.155.160/110 | 18 | -214 | 482.538 | 24.748 | BCST DCSTc | 2 | -32.891 |
| 88.111.130.137/108 | 15 | -218.231 | 482.562 | 24.772 | BVAR DCSTc | 3 | -44.968 |
| 104.155.160/110 | 10 | -224.314 | 482.585 | 24.794 | BCSTc | 1 | -70.448 |

| | | | | | | | |
|---|---|---|---|---|---|---|---|
| 88.95.111.130.137.147.155/ | 19 | -210.647 | 482.651 | 24.861 | BVAR DCSTc | 3 | -44.038 |
| 88.137.147/108 | 12 | -223.587 | 482.659 | 24.869 | BVAR DCSTc | 3 | -44.968 |
| 111.130.137.147.160/108 | 20 | -205.717 | 482.73 | 24.94 | BVAR DCSTc | 3 | -72.804 |
| 104.130.147/110 | 10 | -223.908 | 482.738 | 24.948 | BCSTc | 1 | -70.448 |
| 104.111.130.155.160/95.108 | 19 | -207.91 | 482.755 | 24.965 | BVAR DCSTc | 3 | -15.158 |
| 104.137.155.160/95.108 | 18 | -210.309 | 482.796 | 25.006 | BVAR DCSTc | 3 | -15.158 |
| 95.130/ | 7 | -230.758 | 482.806 | 25.016 | BVARc | 2 | -171.311 |
| 137.147.155/110 | 11 | -224.743 | 482.811 | 25.021 | BCST DCSTc | 2 | -80.506 |
| 147.155.160/110 | 13 | -222.576 | 482.841 | 25.051 | BCST DCSTc | 2 | -80.506 |
| 104.111.137.147.160/95 | 16 | -215.832 | 482.864 | 25.074 | BCST DCSTc | 2 | -78.023 |
| 104.111/ | 6 | -231.937 | 482.877 | 25.087 | BVARc | 2 | -189.51 |
| 88.111.137.147.160/108 | 17 | -216.276 | 482.903 | 25.113 | BVAR DCSTc | 3 | -44.968 |
| 88.104.111.147.160/108 | 16 | -215.3 | 482.95 | 25.16 | BVAR DCSTc | 3 | -31.012 |
| 95.111.130.147/ | 11 | -225.158 | 482.96 | 25.169 | BCST DCSTc | 2 | -113.508 |
| 88.137.160/ | 10 | -226.464 | 482.983 | 25.193 | BCST DCSTc | 2 | -160.846 |
| 95.137.155.160/110 | 14 | -220.197 | 483.015 | 25.225 | BCST DCSTc | 2 | -32.891 |
| 111.137.155/ | 9 | -227.991 | 483.032 | 25.241 | BCST DCSTc | 2 | -163.421 |
| 88.137.147.160/108 | 16 | -217.831 | 483.09 | 25.3 | BVAR DCSTc | 3 | -44.968 |
| 104.111.130.160/95 | 14 | -218.241 | 483.098 | 25.308 | BCST DCSTc | 2 | -103.956 |
| 130/ | 5 | -233.473 | 483.108 | 25.318 | BVARc | 2 | -219.409 |
| 147.160/ | 7 | -231.546 | 483.112 | 25.321 | BCST DCSTc | 2 | -188.226 |
| 88.111.130/ | 9 | -227.864 | 483.178 | 25.387 | BCST DCSTc | 2 | -157.593 |
| 111.137.155/108 | 12 | -224.311 | 483.216 | 25.426 | BVAR DCSTc | 3 | -72.804 |
| 95.111.130.137/ | 12 | -223.475 | 483.217 | 25.426 | BCST DCSTc | 2 | -112.825 |
| 88.104.130.160/108 | 15 | -215.152 | 483.225 | 25.435 | BVAR DCSTc | 3 | -31.012 |
| 104.111.130.147.155/95.108 | 18 | -209.723 | 483.245 | 25.455 | BVAR DCSTc | 3 | -15.158 |
| 104.111.137.160/108 | 15 | -218.3 | 483.246 | 25.456 | BVAR DCSTc | 3 | -60.749 |
| 104.111.130.147.160/95 | 16 | -215.06 | 483.27 | 25.48 | BCST DCSTc | 2 | -81.481 |
| 95.111.130.147.155.160/ | 16 | -216.461 | 483.348 | 25.558 | BCST DCSTc | 2 | -67.418 |
| 95.111.137.155.160/ | 14 | -220.766 | 483.373 | 25.583 | BCST DCSTc | 2 | -86.786 |
| 130.137.147.160/108 | 18 | -214.157 | 483.38 | 25.59 | BVAR DCSTc | 3 | -72.804 |
| 104.130.147.160/110 | 13 | -219.475 | 483.4 | 25.61 | BCSTc | 1 | -70.448 |
| 88.130.137.147/110 | 17 | -215.647 | 483.427 | 25.636 | BVAR DCSTc | 3 | -53.71 |
| 88.104.147.160/108 | 14 | -218.056 | 483.434 | 25.644 | BVAR DCSTc | 3 | -31.012 |
| 104.130.155.160/95.108 | 18 | -209.662 | 483.459 | 25.669 | BVAR DCSTc | 3 | -15.158 |
| 104.111.130.137.155/95.108 | 19 | -208.004 | 483.475 | 25.685 | BVAR DCSTc | 3 | -15.158 |
| 111.130.147/108 | 12 | -224.243 | 483.481 | 25.69 | BVAR DCSTc | 3 | -72.804 |
| 104.137/108 | 10 | -225.716 | 483.517 | 25.727 | BVAR DCSTc | 3 | -60.749 |
| 88.160/ | 7 | -231.751 | 483.522 | 25.731 | BCST DCSTc | 2 | -184.427 |
| 88.95.111.160/ | 11 | -226.708 | 483.522 | 25.732 | BCST DCSTc | 2 | -101.091 |
| 130.147.155/110 | 11 | -224.122 | 483.549 | 25.759 | BCST DCSTc | 2 | -80.506 |
| 88.137.147.155.160/110 | 19 | -212.733 | 483.6 | 25.81 | BVAR DCSTc | 3 | -53.71 |
| 95.147/ | 6 | -233.972 | 483.635 | 25.845 | BVARc | 2 | -169.295 |

| | | | | | | | |
|---|---|---|---|---|---|---|---|
| 95.111.155/ | 8 | -230.749 | 483.661 | 25.871 | BVARc | 2 | -139.089 |
| 88.137.147.160/ | 12 | -223.598 | 483.692 | 25.902 | BCST DCSTc | 2 | -138.686 |
| 104.137.155/110 | 11 | -223.438 | 483.697 | 25.907 | BCSTc | 1 | -70.448 |
| 111.130.137/108 | 13 | -222.54 | 483.714 | 25.924 | BVAR DCSTc | 3 | -72.804 |
| 88.111.130.137.160/108 | 20 | -210.529 | 483.724 | 25.934 | BVAR DCSTc | 3 | -44.968 |
| 104.137.147/95.110 | 13 | -220.657 | 483.728 | 25.938 | BCST DCSTc | 2 | -32.891 |
| 88.95.137.160/ | 12 | -224.261 | 483.749 | 25.959 | BCST DCSTc | 2 | -113.259 |
| 104.111.130.160/ | 12 | -221.614 | 483.756 | 25.966 | BCST DCSTc | 2 | -141.097 |
| 104.111.147/108 | 11 | -225.195 | 483.789 | 25.998 | BVAR DCSTc | 3 | -60.749 |
| 95.130.147.155.160/110 | 18 | -213.708 | 483.79 | 25.999 | BCST DCSTc | 2 | -32.891 |
| 95.111.130.155.160/ | 14 | -220.01 | 483.819 | 26.029 | BCST DCSTc | 2 | -90.26 |
| 88.130.137.147.160/ | 16 | -216.482 | 483.859 | 26.069 | BVAR DCSTc | 3 | -117.506 |
| 88.104.137.147/110 | 14 | -218.674 | 483.877 | 26.086 | BVAR DCSTc | 3 | -41.22 |
| 130.147.160/ | 10 | -225.916 | 483.879 | 26.089 | BCST DCSTc | 2 | -168.532 |
| 111.137.155.160/ | 12 | -223.674 | 483.901 | 26.111 | BCST DCSTc | 2 | -135.078 |
| 104.111.147.160/95 | 13 | -221.458 | 483.933 | 26.142 | BCST DCSTc | 2 | -101.943 |
| 95.130.155.160/110 | 14 | -219.669 | 483.935 | 26.144 | BCST DCSTc | 2 | -32.891 |
| 88.104.111.137.147/108 | 16 | -215.524 | 483.948 | 26.158 | BVAR DCSTc | 3 | -31.012 |
| 95.111.130.137.155.160/ | 17 | -214.934 | 483.988 | 26.198 | BCST DCSTc | 2 | -66.891 |
| 104.130.137.155.160/95.108 | 22 | -203.443 | 484.053 | 26.263 | BVAR DCSTc | 3 | -15.158 |
| 88.111.130.155/108 | 14 | -220.018 | 484.065 | 26.274 | BVAR DCSTc | 3 | -44.968 |
| 104.137.147.155/95.108 | 16 | -213.667 | 484.068 | 26.278 | BVAR DCSTc | 3 | -15.158 |
| 88.130.147/108 | 12 | -223.298 | 484.069 | 26.278 | BVAR DCSTc | 3 | -44.968 |
| 88.137.155.160/110 | 15 | -218.931 | 484.077 | 26.287 | BVAR DCSTc | 3 | -53.71 |
| 88.130.147.160/108 | 16 | -217.353 | 484.084 | 26.294 | BVAR DCSTc | 3 | -44.968 |
| 88/ | 4 | -236.765 | 484.093 | 26.303 | BVARc | 2 | -213.467 |
| 95.111.155.160/ | 11 | -226.206 | 484.095 | 26.304 | BCST DCSTc | 2 | -110.52 |
| 88.95.147.160/ | 11 | -226.247 | 484.106 | 26.315 | BCST DCSTc | 2 | -114.245 |
| 95.111.130.137.147.155/108 | 19 | -210.334 | 484.158 | 26.368 | BVAR DCSTc | 3 | -15.158 |
| 104.111.137/ | 9 | -227.558 | 484.161 | 26.37 | BVARc | 2 | -166.837 |
| 88.104.155/108 | 11 | -222.374 | 484.17 | 26.38 | BVAR DCSTc | 3 | -31.012 |
| 130.137.160/ | 11 | -224.262 | 484.179 | 26.389 | BCST DCSTc | 2 | -167.878 |
| 88.95.130.160/ | 12 | -223.501 | 484.215 | 26.425 | BCST DCSTc | 2 | -116.73 |
| 137.147/108 | 10 | -228.101 | 484.221 | 26.431 | BVAR DCSTc | 3 | -72.804 |
| 104.111.130.137.160/95 | 17 | -213.727 | 484.254 | 26.464 | BCST DCSTc | 2 | -81.148 |
| 88.104.137.147.160/110 | 17 | -214.096 | 484.262 | 26.472 | BVAR DCSTc | 3 | -41.22 |
| 111.155.160/ | 9 | -228.893 | 484.27 | 26.48 | BCST DCSTc | 2 | -158.591 |
| 95.111.137.147.155.160/ | 17 | -216.423 | 484.28 | 26.49 | BVAR DCSTc | 3 | -63.149 |
| 88.130.160/ | 10 | -226.13 | 484.306 | 26.516 | BCST DCSTc | 2 | -164.742 |
| 88.111.137.147/ | 11 | -226.223 | 484.363 | 26.572 | BCST DCSTc | 2 | -132.428 |
| 104.130.137.155.160/110 | 18 | -211.401 | 484.371 | 26.58 | BCSTc | 1 | -70.448 |
| 104.137.147.155.160/95.108 | 21 | -206.402 | 484.372 | 26.581 | BVAR DCSTc | 3 | -15.158 |
| 147/ | 4 | -236.908 | 484.378 | 26.588 | BVARc | 2 | -217.614 |

| | | | | | | | |
|---|---|---|---|---|---|---|---|
| 104.137.147.160/95.110 | 16 | -216.223 | 484.402 | 26.612 | BCST DCSTc | 2 | -32.891 |
| 104.130.137.147.160/95 | 17 | -213.104 | 484.424 | 26.634 | BCST DCSTc | 2 | -94.141 |
| 137.147.160/108 | 14 | -222.254 | 484.469 | 26.679 | BVAR DCSTc | 3 | -72.804 |
| 104.155.160/95.110 | 13 | -220.525 | 484.485 | 26.695 | BCST DCSTc | 2 | -32.891 |
| 88.111.130.147.160/108 | 19 | -213.002 | 484.51 | 26.72 | BVAR DCSTc | 3 | -44.968 |
| 111.137.147.160/108 | 15 | -220.813 | 484.511 | 26.721 | BVAR DCSTc | 3 | -72.804 |
| 104.147.155/110 | 9 | -226.814 | 484.537 | 26.746 | BCSTc | 1 | -70.448 |
| 104.130.137/110 | 11 | -223.003 | 484.542 | 26.752 | BCSTc | 1 | -70.448 |
| 88.111.155.160/108 | 14 | -221.523 | 484.555 | 26.765 | BVAR DCSTc | 3 | -44.968 |
| 104.130.137.147/95.108 | 18 | -210.716 | 484.56 | 26.77 | BVAR DCSTc | 3 | -15.158 |
| 88.104.130.137.147.155.160/108 | 24 | -200.441 | 484.577 | 26.787 | BVAR DCSTc | 3 | -31.012 |
| 88.111.147/ | 8 | -231.38 | 484.61 | 26.82 | BCST DCSTc | 2 | -155.879 |
| 104.130.155/110 | 10 | -224.057 | 484.622 | 26.832 | BCSTc | 1 | -70.448 |
| 88.137.155/108 | 12 | -223.782 | 484.637 | 26.847 | BVAR DCSTc | 3 | -44.968 |
| 104.130.147/95.110 | 13 | -220.119 | 484.639 | 26.848 | BCST DCSTc | 2 | -32.891 |
| 104.111.137.147.160/ | 14 | -219.807 | 484.642 | 26.852 | BCST DCSTc | 2 | -115.765 |
| 88.104.111.130.137.147.160/108 | 24 | -196.742 | 484.645 | 26.855 | BVAR DCSTc | 3 | -31.012 |
| 130.137.155/110 | 14 | -220.432 | 484.685 | 26.895 | BCST DCSTc | 2 | -80.506 |
| 88.130.147.160/ | 12 | -223.104 | 484.693 | 26.903 | BCST DCSTc | 2 | -142.423 |
| 104.130.147.155.160/95.108 | 21 | -205.648 | 484.725 | 26.935 | BVAR DCSTc | 3 | -15.158 |
| 111.130.155/108 | 12 | -224.09 | 484.742 | 26.952 | BVAR DCSTc | 3 | -72.804 |
| 88.130.147.155.160/110 | 19 | -212.442 | 484.851 | 27.061 | BVAR DCSTc | 3 | -53.71 |
| 95.111.130.137.147/ | 14 | -221.043 | 484.876 | 27.086 | BCST DCSTc | 2 | -91.098 |
| 111.130.137.160/108 | 18 | -214.843 | 484.884 | 27.094 | BVAR DCSTc | 3 | -72.804 |
| 88.147.160/ | 9 | -229.221 | 484.889 | 27.098 | BCST DCSTc | 2 | -162.603 |
| 88.104.137.147.155.160/95 | 19 | -210.858 | 484.944 | 27.154 | BVAR DCSTc | 3 | -69.292 |
| 104.111.130.137.147.160/ | 18 | -212.588 | 484.947 | 27.157 | BVAR DCSTc | 3 | -94.483 |
| 104.130/108 | 10 | -225.438 | 484.952 | 27.162 | BVAR DCSTc | 3 | -60.749 |
| 104/95 | 6 | -232.392 | 484.961 | 27.171 | BVARc | 2 | -189.106 |
| 111.130.137.147/ | 12 | -223.739 | 484.994 | 27.204 | BCST DCSTc | 2 | -139.179 |
| 88.130.155.160/110 | 15 | -218.402 | 484.996 | 27.206 | BVAR DCSTc | 3 | -53.71 |
| 130.137.147.160/ | 13 | -221.452 | 484.999 | 27.209 | BCST DCSTc | 2 | -145.774 |
| 88.104.130.147/110 | 14 | -218.247 | 485.009 | 27.218 | BVAR DCSTc | 3 | -41.22 |
| 95.130.137.147.155/108 | 18 | -212.317 | 485.015 | 27.225 | BVAR DCSTc | 3 | -15.158 |
| 95.137.147.155/110 | 13 | -222.732 | 485.022 | 27.232 | BCST DCSTc | 2 | -32.891 |
| 104.111.130.160/108 | 15 | -218.214 | 485.024 | 27.233 | BVAR DCSTc | 3 | -60.749 |
| 104.111.137/95 | 11 | -224.963 | 485.026 | 27.236 | BVARc | 2 | -130.474 |
| 88.95.111.130.137.147/ | 17 | -216.373 | 485.038 | 27.247 | BVAR DCSTc | 3 | -63.131 |
| 88.155.160/108 | 12 | -224.274 | 485.056 | 27.266 | BVAR DCSTc | 3 | -44.968 |
| 104.160/95 | 9 | -227.713 | 485.065 | 27.275 | BVARc | 2 | -160.401 |
| 95.147.155.160/110 | 15 | -220.593 | 485.11 | 27.319 | BCST DCSTc | 2 | -32.891 |
| 88.104.130.147.155.160/95 | 19 | -209.98 | 485.113 | 27.322 | BVAR DCSTc | 3 | -72.646 |
| 88.130.137.160/108 | 17 | -216.045 | 485.119 | 27.329 | BVAR DCSTc | 3 | -44.968 |

| | | | | | | | |
|---|---|---|---|---|---|---|---|
| 88.104.130.147.160/110 | 17 | -213.545 | 485.133 | 27.343 | BVAR DCSTc | 3 | -41.22 |
| 88.104.155.160/110 | 14 | -218.795 | 485.141 | 27.351 | BVAR DCSTc | 3 | -41.22 |
| 95.111.137.155/ | 11 | -226.455 | 485.154 | 27.363 | BCST DCSTc | 2 | -116.501 |
| 104.111.155/108 | 11 | -225.092 | 485.163 | 27.373 | BVAR DCSTc | 3 | -60.749 |
| 130.147.160/108 | 14 | -221.63 | 485.172 | 27.381 | BVAR DCSTc | 3 | -72.804 |
| 104.111.147.160/ | 11 | -225.124 | 485.177 | 27.387 | BCST DCSTc | 2 | -139.377 |
| 111.130.155/ | 9 | -228.091 | 485.222 | 27.432 | BCST DCSTc | 2 | -167.752 |
| 88.130.137/108 | 13 | -222.073 | 485.231 | 27.441 | BVAR DCSTc | 3 | -44.968 |
| 88.130.137.160/ | 13 | -221.578 | 485.251 | 27.46 | BCST DCSTc | 2 | -141.896 |
| 104.147.155.160/95.108 | 17 | -213.363 | 485.26 | 27.47 | BVAR DCSTc | 3 | -15.158 |
| 111.130.147.160/108 | 17 | -217.116 | 485.271 | 27.481 | BVAR DCSTc | 3 | -72.804 |
| 104.130.147.160/95.110 | 16 | -215.685 | 485.301 | 27.511 | BCST DCSTc | 2 | -32.891 |
| 88.111.147.155/108 | 13 | -223.441 | 485.311 | 27.521 | BVAR DCSTc | 3 | -44.968 |
| 104.130.147.155/95.108 | 16 | -213.351 | 485.418 | 27.627 | BVAR DCSTc | 3 | -15.158 |
| 111.130.155.160/ | 12 | -223.441 | 485.419 | 27.629 | BCST DCSTc | 2 | -139.076 |
| 104.147/108 | 9 | -228.485 | 485.447 | 27.657 | BVAR DCSTc | 3 | -60.749 |
| 104.137.160/108 | 13 | -221.951 | 485.484 | 27.694 | BVAR DCSTc | 3 | -60.749 |
| 111.155.160/108 | 12 | -225.739 | 485.519 | 27.729 | BVAR DCSTc | 3 | -72.804 |
| 104.137.147.160/95 | 14 | -219.701 | 485.539 | 27.748 | BCST DCSTc | 2 | -114.801 |
| 88.104.111.137.160/ | 14 | -220.267 | 485.563 | 27.773 | BCST DCSTc | 2 | -112.222 |
| 88.104.137.155/110 | 15 | -217.576 | 485.567 | 27.777 | BVAR DCSTc | 3 | -41.22 |
| 88.104.111.137.155/108 | 16 | -215.564 | 485.59 | 27.8 | BVAR DCSTc | 3 | -31.012 |
| 104.137.155/95.110 | 14 | -219.649 | 485.598 | 27.808 | BCST DCSTc | 2 | -32.891 |
| 104.137.160/95 | 12 | -222.985 | 485.655 | 27.865 | BCST DCSTc | 2 | -137.379 |
| 130.147/108 | 10 | -227.827 | 485.66 | 27.87 | BVAR DCSTc | 3 | -72.804 |
| 95.137.147/ | 9 | -229.994 | 485.72 | 27.929 | BCST DCSTc | 2 | -147.023 |
| 104.111.147.155.160/95.108 | 20 | -209.261 | 485.817 | 28.026 | BVAR DCSTc | 3 | -15.158 |
| 88.104.130.137.147.160/108 | 22 | -205.451 | 485.835 | 28.044 | BVAR DCSTc | 3 | -31.012 |
| 88.104.111.130.147/108 | 16 | -215.522 | 485.905 | 28.115 | BVAR DCSTc | 3 | -31.012 |
| 88.104.111.137.147.160/ | 17 | -215.873 | 485.917 | 28.127 | BVAR DCSTc | 3 | -88.533 |
| 104.111.130.147.160/ | 14 | -219.477 | 485.957 | 28.167 | BCST DCSTc | 2 | -119.666 |
| 88.111.130.147/ | 11 | -226.032 | 485.964 | 28.174 | BCST DCSTc | 2 | -136.467 |
| 137.155/108 | 10 | -228.185 | 485.975 | 28.185 | BVAR DCSTc | 3 | -72.804 |
| 88.111.130.137/ | 12 | -224.24 | 485.996 | 28.206 | BCST DCSTc | 2 | -135.676 |
| 95.137.147.155.160/ | 14 | -221.357 | 485.998 | 28.207 | BCST DCSTc | 2 | -100.992 |
| 95.130.137.155.160/ | 15 | -218.579 | 486.042 | 28.251 | BCST DCSTc | 2 | -103.445 |
| 95.130.147.155/110 | 13 | -222.268 | 486.074 | 28.284 | BCST DCSTc | 2 | -32.891 |
| 88.137.147.155/110 | 14 | -221.465 | 486.083 | 28.293 | BVAR DCSTc | 3 | -53.71 |
| 88.104.111.130.137/108 | 17 | -213.792 | 486.085 | 28.295 | BVAR DCSTc | 3 | -31.012 |
| 88.104.111.160/ | 11 | -225.582 | 486.093 | 28.302 | BCST DCSTc | 2 | -135.831 |
| 88.130.155/108 | 12 | -223.518 | 486.096 | 28.306 | BVAR DCSTc | 3 | -44.968 |
| 95.130.147.155.160/ | 14 | -220.447 | 486.156 | 28.366 | BCST DCSTc | 2 | -104.313 |
| 104.111.130.137.160/ | 16 | -216.554 | 486.164 | 28.374 | BVAR DCSTc | 3 | -117.744 |

| | | | | | | | |
|---|---|---|---|---|---|---|---|
| 88.147.155.160/110 | 16 | -219.326 | 486.171 | 28.381 | BVAR DCSTc | 3 | -53.71 |
| 88.104.137.147/108 | 14 | -219.148 | 486.182 | 28.392 | BVAR DCSTc | 3 | -31.012 |
| 95.111.147.155.160/ | 13 | -224.003 | 486.205 | 28.415 | BCST DCSTc | 2 | -89.023 |
| 111.147.155/108 | 11 | -227.625 | 486.213 | 28.423 | BVAR DCSTc | 3 | -72.804 |
| 130.137.160/108 | 15 | -220.329 | 486.219 | 28.429 | BVAR DCSTc | 3 | -72.804 |
| 88.95.160/ | 9 | -230.529 | 486.223 | 28.433 | BCST DCSTc | 2 | -137.822 |
| 95.155/ | 6 | -234.488 | 486.26 | 28.47 | BVARc | 2 | -175.737 |
| 104.130.137.155.160/95.110 | 21 | -207.612 | 486.271 | 28.481 | BCST DCSTc | 2 | -32.891 |
| 104.111.147.160/108 | 14 | -221.661 | 486.319 | 28.529 | BVAR DCSTc | 3 | -60.749 |
| 88.104.130.137.155.160/95 | 20 | -208.752 | 486.332 | 28.542 | BVAR DCSTc | 3 | -72.417 |
| 104.130.147.160/95 | 14 | -219.111 | 486.343 | 28.553 | BCST DCSTc | 2 | -118.441 |
| 95.137.155.160/ | 12 | -224.786 | 486.383 | 28.593 | BCST DCSTc | 2 | -123.715 |
| 88.104.111.137.147.160/108 | 19 | -211.837 | 486.426 | 28.636 | BVAR DCSTc | 3 | -31.012 |
| 104.111.130/ | 9 | -227.698 | 486.431 | 28.641 | BVARc | 2 | -171.208 |
| 104.147.155/95.110 | 12 | -223.025 | 486.437 | 28.647 | BCST DCSTc | 2 | -32.891 |
| 104.130.137/95.110 | 14 | -219.214 | 486.443 | 28.653 | BCST DCSTc | 2 | -32.891 |
| 88.104.137.147.160/95 | 16 | -216.917 | 486.45 | 28.659 | BCST DCSTc | 2 | -88.719 |
| 88.104.130.147.160/95 | 16 | -215.934 | 486.461 | 28.671 | BCST DCSTc | 2 | -91.967 |
| 155.160/108 | 10 | -228.71 | 486.461 | 28.671 | BVAR DCSTc | 3 | -72.804 |
| 88.147.155/108 | 11 | -226.522 | 486.504 | 28.714 | BVAR DCSTc | 3 | -44.968 |
| 88.104.130.137/110 | 17 | -214.817 | 486.515 | 28.725 | BVAR DCSTc | 3 | -41.22 |
| 104.130.137.155/95.108 | 17 | -212.094 | 486.52 | 28.73 | BVAR DCSTc | 3 | -15.158 |
| 104.130.155/95.110 | 13 | -220.268 | 486.523 | 28.733 | BCST DCSTc | 2 | -32.891 |
| 111.137.147.155/ | 11 | -226.513 | 486.526 | 28.735 | BCST DCSTc | 2 | -142.649 |
| 130.137.147.155/110 | 16 | -217.949 | 486.536 | 28.745 | BCST DCSTc | 2 | -80.506 |
| 88.104.147.155/110 | 14 | -219.899 | 486.584 | 28.794 | BVAR DCSTc | 3 | -41.22 |
| 88.104.111.130.147.160/ | 17 | -215.257 | 486.585 | 28.795 | BVAR DCSTc | 3 | -92.148 |
| 88.104.130.137.155.160/110 | 22 | -205.721 | 486.603 | 28.813 | BVAR DCSTc | 3 | -41.22 |
| 88.104.137.147.160/108 | 18 | -213.392 | 486.613 | 28.823 | BVAR DCSTc | 3 | -31.012 |
| 130.137/108 | 11 | -226.53 | 486.679 | 28.889 | BVAR DCSTc | 3 | -72.804 |
| 95.111.130.155/ | 11 | -226.234 | 486.692 | 28.902 | BCST DCSTc | 2 | -120.511 |
| 104.111.130/95 | 11 | -224.815 | 486.716 | 28.926 | BVARc | 2 | -134.557 |
| 137.147/ | 7 | -233.07 | 486.728 | 28.938 | BCST DCSTc | 2 | -195.482 |
| 88.104.111.130.160/ | 14 | -219.886 | 486.775 | 28.985 | BCST DCSTc | 2 | -116.071 |
| 104.130.160/95 | 12 | -222.565 | 486.804 | 29.014 | BCST DCSTc | 2 | -141.19 |
| 95.130.137.155/110 | 16 | -218.379 | 486.813 | 29.022 | BCST DCSTc | 2 | -32.891 |
| 88.104.111.147.155.160/95 | 18 | -214.149 | 486.827 | 29.037 | BVAR DCSTc | 3 | -57.969 |
| 155/ | 4 | -237.342 | 486.842 | 29.052 | BVARc | 2 | -223.975 |
| 88.137/ | 7 | -233.14 | 486.868 | 29.078 | BVARc | 2 | -191.548 |
| 104.130.137.147/110 | 15 | -218.46 | 486.888 | 29.098 | BCSTc | 1 | -70.448 |
| 111.147.155/ | 8 | -231.73 | 486.899 | 29.109 | BCST DCSTc | 2 | -166.159 |
| 111.130.137.155.160/ | 16 | -217.933 | 486.952 | 29.162 | BVAR DCSTc | 3 | -115.274 |
| 95.130.147/ | 9 | -229.617 | 486.956 | 29.165 | BCST DCSTc | 2 | -150.876 |

| | | | | | | | |
|---|---|---|---|---|---|---|---|
| 104.137.147.155.160/110 | 17 | -215.51 | 486.989 | 29.199 | BCSTc | 1 | -70.448 |
| 88.111.137.155.160/ | 14 | -221.977 | 486.997 | 29.206 | BCST DCSTc | 2 | -110.083 |
| 88.104.130.155/110 | 15 | -217.308 | 487.003 | 29.213 | BVAR DCSTc | 3 | -41.22 |
| 104.130.137.160/95 | 15 | -217.635 | 487.007 | 29.217 | BCST DCSTc | 2 | -117.966 |
| 104.130.160/108 | 13 | -221.725 | 487.017 | 29.227 | BVAR DCSTc | 3 | -60.749 |
| 88.111.130.137.155.160/ | 18 | -214.597 | 487.022 | 29.232 | BVAR DCSTc | 3 | -88.639 |
| 88.104.111.130.137.160/ | 18 | -213.654 | 487.079 | 29.289 | BVAR DCSTc | 3 | -91.545 |
| 88.130.147.155/110 | 14 | -221.001 | 487.136 | 29.345 | BVAR DCSTc | 3 | -53.71 |
| 95.130.155.160/ | 12 | -224.18 | 487.158 | 29.368 | BCST DCSTc | 2 | -127.34 |
| 104.155/108 | 9 | -228.552 | 487.172 | 29.382 | BVAR DCSTc | 3 | -60.749 |
| 104.147.160/108 | 12 | -224.607 | 487.182 | 29.392 | BVAR DCSTc | 3 | -60.749 |
| 104.130.137.147.155.160/108 | 22 | -206.426 | 487.194 | 29.404 | BVAR DCSTc | 3 | -60.749 |
| 104.147.160/95 | 11 | -225.566 | 487.206 | 29.416 | BCST DCSTc | 2 | -138.96 |
| 88.104.111.130.137.160/108 | 22 | -206.091 | 487.247 | 29.456 | BVAR DCSTc | 3 | -31.012 |
| 104.111.137.147/108 | 14 | -221.877 | 487.3 | 29.51 | BVAR DCSTc | 3 | -60.749 |
| 88.111.137.147.155.160/ | 17 | -217.547 | 487.323 | 29.533 | BVAR DCSTc | 3 | -86.359 |
| 130.155/108 | 10 | -227.869 | 487.333 | 29.543 | BVAR DCSTc | 3 | -72.804 |
| 95.130.137/ | 10 | -228.007 | 487.345 | 29.555 | BCST DCSTc | 2 | -150.266 |
| 88.104.130.137.160/95 | 18 | -213.366 | 487.36 | 29.57 | BVAR DCSTc | 3 | -90.398 |
| 88.111.130.147.155.160/ | 17 | -216.632 | 487.384 | 29.594 | BVAR DCSTc | 3 | -89.674 |
| 104.137.155.160/110 | 13 | -221.704 | 487.46 | 29.67 | BCSTc | 1 | -70.448 |
| 88.111.155.160/ | 11 | -227.274 | 487.485 | 29.695 | BCST DCSTc | 2 | -133.674 |
| 88.130.137.147.155.160/ | 18 | -214.253 | 487.534 | 29.744 | BVAR DCSTc | 3 | -101.91 |
| 95.155.160/ | 9 | -230.401 | 487.556 | 29.766 | BCST DCSTc | 2 | -147.624 |
| 88.104.111.130.155/108 | 16 | -215.58 | 487.588 | 29.797 | BVAR DCSTc | 3 | -31.012 |
| 88.104.130.147/108 | 14 | -218.859 | 487.592 | 29.801 | BVAR DCSTc | 3 | -31.012 |
| 88.104.130.147.160/108 | 18 | -212.914 | 487.607 | 29.817 | BVAR DCSTc | 3 | -31.012 |
| 88.111.137.155/ | 11 | -227.084 | 487.667 | 29.877 | BCST DCSTc | 2 | -139.215 |
| 95.111.137.147.155/ | 13 | -224.479 | 487.71 | 29.92 | BCST DCSTc | 2 | -95.231 |
| 104.130.147.155.160/110 | 17 | -214.971 | 487.745 | 29.954 | BCSTc | 1 | -70.448 |
| 147.155/108 | 9 | -230.895 | 487.784 | 29.994 | BVAR DCSTc | 3 | -72.804 |
| 104.111.137.147/ | 11 | -226.147 | 487.786 | 29.996 | BCST DCSTc | 2 | -146.132 |
| 88.104.111.137.160/95 | 17 | -216.98 | 487.789 | 29.999 | BVAR DCSTc | 3 | -75.167 |
| 104.130.137.147.155.160/95 | 20 | -209.484 | 487.796 | 30.006 | BVAR DCSTc | 3 | -77.153 |
| 111.137.147.155.160/ | 15 | -221.156 | 487.798 | 30.008 | BVAR DCSTc | 3 | -113.266 |
| 88.111.155/ | 8 | -232.186 | 487.812 | 30.021 | BCST DCSTc | 2 | -162.612 |
| 88.104.111.147.160/ | 13 | -223.221 | 487.846 | 30.055 | BCST DCSTc | 2 | -114.176 |
| 88.104.111.147.160/95 | 16 | -218.865 | 487.85 | 30.06 | BVAR DCSTc | 3 | -76.052 |
| 88.130.137.155/110 | 17 | -217.112 | 487.874 | 30.084 | BVAR DCSTc | 3 | -53.71 |
| 88.111.130.155.160/ | 14 | -221.432 | 487.88 | 30.09 | BCST DCSTc | 2 | -113.768 |
| 88.104.130.137.147.160/ | 18 | -213.466 | 487.935 | 30.145 | BVAR DCSTc | 3 | -104.973 |
| 95.111.147.155/ | 10 | -229.672 | 487.968 | 30.178 | BCST DCSTc | 2 | -118.718 |
| 111.130.147.155.160/108 | 19 | -213.156 | 487.988 | 30.198 | BVAR DCSTc | 3 | -72.804 |

| | | | | | | | |
|---|---|---|---|---|---|---|---|
| 95.130.137.147/ | 12 | -225.109 | 487.994 | 30.204 | BCST DCSTc | 2 | -128.074 |
| 88.104.111.130.147.160/108 | 21 | -208.564 | 488.033 | 30.243 | BVAR DCSTc | 3 | -31.012 |
| 111.130.137.147.155.160/ | 18 | -215.106 | 488.041 | 30.251 | BVAR DCSTc | 3 | -93.152 |
| 88.104.111.130.160/95 | 16 | -217.448 | 488.046 | 30.256 | BCST DCSTc | 2 | -79.865 |
| 88.111.137.155.160/108 | 17 | -218.118 | 488.064 | 30.274 | BVAR DCSTc | 3 | -44.968 |
| 88.104.111.155.160/108 | 16 | -217.085 | 488.078 | 30.288 | BVAR DCSTc | 3 | -31.012 |
| 104.130.155.160/110 | 13 | -221.057 | 488.141 | 30.35 | BCSTc | 1 | -70.448 |
| 104.111.147/ | 8 | -231.361 | 488.156 | 30.366 | BCST DCSTc | 2 | -169.639 |
| 104/ | 4 | -237.001 | 488.157 | 30.367 | BVARc | 2 | -227.483 |
| 88.104.137.155/108 | 14 | -219.344 | 488.16 | 30.37 | BVAR DCSTc | 3 | -31.012 |
| 104.137.160/ | 10 | -227.263 | 488.169 | 30.378 | BCST DCSTc | 2 | -175.425 |
| 104.111.130.137.147.160/108 | 22 | -203.181 | 488.169 | 30.379 | BVAR DCSTc | 3 | -60.749 |
| 111.130.137.155/ | 12 | -224.536 | 488.171 | 30.381 | BCST DCSTc | 2 | -145.902 |
| 104.137/95 | 9 | -228.996 | 488.2 | 30.409 | BVARc | 2 | -167.416 |
| 104.111.147/95 | 10 | -228.387 | 488.26 | 30.47 | BVARc | 2 | -132.898 |
| 88.111.130.137.147/ | 14 | -222.162 | 488.324 | 30.533 | BCST DCSTc | 2 | -114.303 |
| 88.111.130.147.155.160/108 | 21 | -209.598 | 488.338 | 30.548 | BVAR DCSTc | 3 | -44.968 |
| 104.111.137.147/95 | 13 | -223.388 | 488.352 | 30.562 | BCST DCSTc | 2 | -109.605 |
| 111.130.147.155/ | 11 | -226.468 | 488.421 | 30.631 | BCST DCSTc | 2 | -146.834 |
| 130.147/ | 7 | -232.928 | 488.437 | 30.647 | BCST DCSTc | 2 | -199.57 |
| 130.137/ | 8 | -231.136 | 488.459 | 30.669 | BCST DCSTc | 2 | -198.778 |
| 88.95.137.155.160/ | 14 | -222.601 | 488.484 | 30.694 | BCST DCSTc | 2 | -98.232 |
| 88.95.130.155.160/ | 14 | -221.617 | 488.495 | 30.705 | BCST DCSTc | 2 | -101.478 |
| 88.130/ | 7 | -232.96 | 488.501 | 30.711 | BVARc | 2 | -195.598 |
| 88.111.137.147.155/108 | 16 | -219.917 | 488.501 | 30.711 | BVAR DCSTc | 3 | -44.968 |
| 104.160/ | 7 | -232.452 | 488.514 | 30.724 | BCST DCSTc | 2 | -198.908 |
| 88.104.155.160/108 | 14 | -219.836 | 488.579 | 30.789 | BVAR DCSTc | 3 | -31.012 |
| 88.104.111.130.155.160/95 | 19 | -212.25 | 488.628 | 30.838 | BVAR DCSTc | 3 | -61.3 |
| 88.104.130.137.160/108 | 19 | -211.607 | 488.642 | 30.852 | BVAR DCSTc | 3 | -31.012 |
| 104.111.137.155/108 | 14 | -221.774 | 488.655 | 30.865 | BVAR DCSTc | 3 | -60.749 |
| 111.130.147.155.160/ | 15 | -220.625 | 488.678 | 30.888 | BVAR DCSTc | 3 | -116.965 |
| 95.111.130.147.155/ | 13 | -223.992 | 488.703 | 30.913 | BCST DCSTc | 2 | -98.974 |
| 95.130.137.147.155/110 | 18 | -215.939 | 488.75 | 30.959 | BCST DCSTc | 2 | -32.891 |
| 88.104.130.137/108 | 15 | -217.634 | 488.754 | 30.964 | BVAR DCSTc | 3 | -31.012 |
| 88.95.147.155.160/ | 13 | -224.553 | 488.769 | 30.979 | BCST DCSTc | 2 | -99.184 |
| 88.111.137.147.155.160/108 | 21 | -209.678 | 488.784 | 30.994 | BVAR DCSTc | 3 | -44.968 |
| 104.130.137.147/95.110 | 18 | -214.671 | 488.789 | 30.999 | BCST DCSTc | 2 | -32.891 |
| 88.104.130.137.147.155.160/ | 20 | -209.808 | 488.804 | 31.013 | BVAR DCSTc | 3 | -87.947 |
| 104.130.137.147.160/108 | 20 | -211.62 | 488.819 | 31.029 | BVAR DCSTc | 3 | -60.749 |
| 88.104.111.147.155/108 | 15 | -219.003 | 488.834 | 31.044 | BVAR DCSTc | 3 | -31.012 |
| 104.137.147.155.160/95.110 | 20 | -211.721 | 488.89 | 31.1 | BCST DCSTc | 2 | -32.891 |
| 88.104.130.137.147/110 | 19 | -212.674 | 488.91 | 31.12 | BVAR DCSTc | 3 | -41.22 |
| 95.147.155.160/ | 11 | -227.859 | 488.917 | 31.127 | BCST DCSTc | 2 | -125.789 |

| | | | | | | | |
|---|---|---|---|---|---|---|---|
| 104.111.130.147/108 | 14 | -221.706 | 488.92 | 31.13 | BVAR DCSTc | 3 | -60.749 |
| 88.104.111.137/ | 11 | -226.721 | 488.935 | 31.145 | BCST DCSTc | 2 | -142.702 |
| 88.104.111.137.155.160/95 | 19 | -213.305 | 488.953 | 31.163 | BVAR DCSTc | 3 | -58.125 |
| 88.104.111/ | 8 | -231.797 | 489.028 | 31.238 | BVARc | 2 | -166.072 |
| 88.111.130.137.147/108 | 17 | -218.14 | 489.03 | 31.24 | BVAR DCSTc | 3 | -44.968 |
| 88.104.137.147.155.160/110 | 21 | -209.761 | 489.084 | 31.294 | BVAR DCSTc | 3 | -41.22 |
| 104.111.130.137/108 | 15 | -220.003 | 489.153 | 31.363 | BVAR DCSTc | 3 | -60.749 |
| 95.111.130.137.155/ | 14 | -222.407 | 489.166 | 31.376 | BCST DCSTc | 2 | -98.39 |
| 88.137.155.160/108 | 15 | -221.359 | 489.31 | 31.519 | BVAR DCSTc | 3 | -44.968 |
| 137.155/ | 7 | -233.574 | 489.329 | 31.539 | BVARc | 2 | -201.913 |
| 104.137.155.160/95.110 | 16 | -217.915 | 489.36 | 31.57 | BCST DCSTc | 2 | -32.891 |
| 88.104.111.130.137.147.155/95 | 21 | -208.549 | 489.365 | 31.575 | BVAR DCSTc | 3 | -44.038 |
| 88.95.111.155.160/ | 13 | -225.588 | 489.376 | 31.586 | BCST DCSTc | 2 | -86.604 |
| 104.137.147.155/110 | 12 | -224.203 | 489.394 | 31.604 | BCSTc | 1 | -70.448 |
| 137.155.160/ | 10 | -228.879 | 489.404 | 31.614 | BCST DCSTc | 2 | -173.192 |
| 111.137.155.160/108 | 15 | -222.522 | 489.406 | 31.616 | BVAR DCSTc | 3 | -72.804 |
| 104.147.155.160/110 | 14 | -222.035 | 489.424 | 31.634 | BCSTc | 1 | -70.448 |
| 104.111.130.137/ | 12 | -224.172 | 489.436 | 31.646 | BCST DCSTc | 2 | -149.387 |
| 88.111.130.155/ | 11 | -226.986 | 489.457 | 31.667 | BCST DCSTc | 2 | -143.348 |
| 111.147.155.160/ | 11 | -228.261 | 489.458 | 31.668 | BCST DCSTc | 2 | -138.664 |
| 88.111.130.137.155.160/108 | 22 | -207.237 | 489.503 | 31.713 | BVAR DCSTc | 3 | -44.968 |
| 88.95.111.137/ | 11 | -229.424 | 489.513 | 31.723 | BCST DCSTc | 2 | -109.539 |
| 104.130/95 | 9 | -228.66 | 489.52 | 31.73 | BVARc | 2 | -171.311 |
| 95.137.155/ | 9 | -231.114 | 489.55 | 31.76 | BVARc | 2 | -154.07 |
| 88.137.147/ | 9 | -231.27 | 489.554 | 31.764 | BCST DCSTc | 2 | -170.384 |
| 88.104.137.155.160/110 | 17 | -215.958 | 489.561 | 31.771 | BVAR DCSTc | 3 | -41.22 |
| 88.147/ | 6 | -236.316 | 489.614 | 31.824 | BVARc | 2 | -193.724 |
| 88.104.130.155/108 | 14 | -219.079 | 489.619 | 31.829 | BVAR DCSTc | 3 | -31.012 |
| 104.130.147.155.160/95.110 | 20 | -211.182 | 489.645 | 31.855 | BCST DCSTc | 2 | -32.891 |
| 104.137.147/108 | 12 | -225.564 | 489.66 | 31.87 | BVAR DCSTc | 3 | -60.749 |
| 104.111.130.147/95 | 13 | -223.06 | 489.674 | 31.883 | BCST DCSTc | 2 | -113.508 |
| 104.130.160/ | 10 | -227.023 | 489.682 | 31.892 | BCST DCSTc | 2 | -179.416 |
| 104.111.130.147/ | 11 | -226.113 | 489.705 | 31.915 | BCST DCSTc | 2 | -150.329 |
| 88.95.111.137.147/ | 13 | -226.263 | 489.716 | 31.926 | BCST DCSTc | 2 | -87.084 |
| 104.137.147.160/ | 12 | -224.824 | 489.727 | 31.937 | BCST DCSTc | 2 | -153.692 |
| 88.95.111.130.147/ | 13 | -225.312 | 489.777 | 31.987 | BCST DCSTc | 2 | -90.364 |
| 155.160/ | 7 | -234.086 | 489.785 | 31.994 | BCST DCSTc | 2 | -196.693 |
| 88.130.137.147.155/110 | 19 | -214.672 | 489.811 | 32.021 | BVAR DCSTc | 3 | -53.71 |
| 111.137.147.155/108 | 14 | -224.319 | 489.839 | 32.049 | BVAR DCSTc | 3 | -72.804 |
| 88.95.111/ | 8 | -234.652 | 489.88 | 32.09 | BCST DCSTc | 2 | -133.061 |
| 88.111.130.155.160/108 | 17 | -218.086 | 489.906 | 32.116 | BVAR DCSTc | 3 | -44.968 |
| 104.137.147.160/108 | 16 | -219.717 | 489.908 | 32.118 | BVAR DCSTc | 3 | -60.749 |
| 104.111.130.137/95 | 14 | -221.378 | 489.931 | 32.14 | BCST DCSTc | 2 | -112.825 |

| | | | | | | | |
|---|---|---|---|---|---|---|---|
| 104.111.137.147.160/108 | 17 | -218.276 | 489.95 | 32.16 | BVAR DCSTc | 3 | -60.749 |
| 88.104.147.155/108 | 13 | -222.083 | 490.027 | 32.237 | BVAR DCSTc | 3 | -31.012 |
| 104.130.155.160/95.110 | 16 | -217.267 | 490.041 | 32.251 | BCST DCSTc | 2 | -32.891 |
| 104.111.130.147.155.160/95 | 18 | -214.364 | 490.062 | 32.272 | BCST DCSTc | 2 | -67.418 |
| 104.111.137.155.160/95 | 16 | -218.669 | 490.087 | 32.297 | BCST DCSTc | 2 | -86.786 |
| 104.130.147.155/110 | 12 | -223.582 | 490.132 | 32.342 | BCSTc | 1 | -70.448 |
| 111.130.137.155.160/108 | 20 | -211.299 | 490.159 | 32.369 | BVAR DCSTc | 3 | -72.804 |
| 88.104.137.160/ | 12 | -225.042 | 490.162 | 32.372 | BCST DCSTc | 2 | -149.906 |
| 95.111.130.137.147.155/ | 16 | -219.609 | 490.175 | 32.385 | BCST DCSTc | 2 | -76.298 |
| 104.111.130.155/108 | 14 | -221.553 | 490.181 | 32.391 | BVAR DCSTc | 3 | -60.749 |
| 88.104.111.160/95 | 13 | -224.61 | 490.236 | 32.446 | BCST DCSTc | 2 | -101.091 |
| 88.95.111.130/ | 11 | -228.817 | 490.277 | 32.487 | BCST DCSTc | 2 | -113.162 |
| 104.111.130.137.160/108 | 20 | -212.306 | 490.324 | 32.533 | BVAR DCSTc | 3 | -60.749 |
| 88.104.137.147.160/ | 14 | -221.901 | 490.331 | 32.541 | BCST DCSTc | 2 | -127.471 |
| 88.104.130.147.155.160/110 | 21 | -209.469 | 490.335 | 32.545 | BVAR DCSTc | 3 | -41.22 |
| 104.147/95 | 8 | -231.875 | 490.349 | 32.559 | BVARc | 2 | -169.295 |
| 111.130.137.147/108 | 15 | -222.539 | 490.361 | 32.571 | BVAR DCSTc | 3 | -72.804 |
| 88.95.130.137.147/ | 14 | -223.064 | 490.372 | 32.582 | BCST DCSTc | 2 | -102.731 |
| 104.111.155/95 | 10 | -228.651 | 490.375 | 32.585 | BVARc | 2 | -139.089 |
| 104.111.130.137.147.155/95.108 | 21 | -207.988 | 490.376 | 32.586 | BVAR DCSTc | 3 | -15.158 |
| 111.137.147.155.160/108 | 19 | -214.215 | 490.391 | 32.601 | BVAR DCSTc | 3 | -72.804 |
| 88.104.137.160/95 | 14 | -222.164 | 490.463 | 32.673 | BCST DCSTc | 2 | -113.259 |
| 104.111.155/ | 8 | -231.726 | 490.478 | 32.688 | BVARc | 2 | -175.932 |
| 88.104.130.155.160/110 | 17 | -215.43 | 490.48 | 32.69 | BVAR DCSTc | 3 | -41.22 |
| 88.111.130.147.155/108 | 16 | -219.941 | 490.482 | 32.692 | BVAR DCSTc | 3 | -44.968 |
| 104.147.160/ | 9 | -230.233 | 490.5 | 32.71 | BCST DCSTc | 2 | -177.394 |
| 104.111.130.155.160/95 | 16 | -217.912 | 490.533 | 32.743 | BCST DCSTc | 2 | -90.26 |
| 88.111.137.147.155/ | 13 | -225.282 | 490.539 | 32.749 | BCST DCSTc | 2 | -118.12 |
| 88.130.155.160/108 | 15 | -220.999 | 490.569 | 32.778 | BVAR DCSTc | 3 | -44.968 |
| 104.137/ | 7 | -233.203 | 490.585 | 32.795 | BVARc | 2 | -205.391 |
| 88.111.130.137.155/108 | 17 | -218.168 | 490.603 | 32.812 | BVAR DCSTc | 3 | -44.968 |
| 104.130.147.160/108 | 16 | -219.093 | 490.611 | 32.82 | BVAR DCSTc | 3 | -60.749 |
| 88.137.147.155/108 | 14 | -223.549 | 490.631 | 32.84 | BVAR DCSTc | 3 | -44.968 |
| 104.111.130.137.147.155.160/ | 20 | -211.225 | 490.678 | 32.887 | BVAR DCSTc | 3 | -79.753 |
| 104.111.137.155/ | 11 | -226.806 | 490.691 | 32.901 | BCST DCSTc | 2 | -152.718 |
| 104.111.130.137.155.160/95 | 19 | -212.837 | 490.702 | 32.912 | BCST DCSTc | 2 | -66.891 |
| 88.104.111.130.147.155.160/ | 19 | -213.134 | 490.707 | 32.917 | BVAR DCSTc | 3 | -76.658 |
| 104.111.130.147.160/108 | 19 | -214.579 | 490.71 | 32.92 | BVAR DCSTc | 3 | -60.749 |
| 104.130.137.147.160/ | 16 | -218.143 | 490.729 | 32.939 | BVAR DCSTc | 3 | -132.947 |
| 130.155.160/ | 10 | -228.555 | 490.748 | 32.958 | BCST DCSTc | 2 | -177.098 |
| 130.137.147/ | 10 | -229.067 | 490.748 | 32.958 | BCST DCSTc | 2 | -177.415 |
| 88.104.111.130/ | 11 | -226.634 | 490.748 | 32.958 | BCST DCSTc | 2 | -146.846 |
| 104.111.155.160/95 | 13 | -224.108 | 490.809 | 33.018 | BCST DCSTc | 2 | -110.52 |

| | | | | | | | |
|---|---|---|---|---|---|---|---|
| 111.130.155.160/108 | 15 | -222.27 | 490.808 | 33.018 | BVAR DCSTc | 3 | -72.804 |
| 88.104.147.160/95 | 13 | -224.149 | 490.82 | 33.029 | BCST DCSTc | 2 | -114.245 |
| 95.130.155/ | 9 | -230.753 | 490.819 | 33.029 | BVARc | 2 | -157.939 |
| 137.155.160/108 | 14 | -224.669 | 490.849 | 33.059 | BVAR DCSTc | 3 | -72.804 |
| 88.111.147.155/ | 10 | -230.503 | 490.891 | 33.101 | BCST DCSTc | 2 | -141.634 |
| 88.95.111.130.137/ | 14 | -224.057 | 490.905 | 33.115 | BCST DCSTc | 2 | -90.109 |
| 88.104.130.160/95 | 14 | -221.404 | 490.929 | 33.139 | BCST DCSTc | 2 | -116.73 |
| 88.104.160/ | 9 | -230.447 | 490.93 | 33.14 | BCST DCSTc | 2 | -173.605 |
| 104.130.147.160/ | 12 | -224.432 | 490.933 | 33.143 | BCST DCSTc | 2 | -157.531 |
| 88.137.155.160/ | 12 | -226.437 | 490.958 | 33.167 | BCST DCSTc | 2 | -147.452 |
| 104.111.155.160/108 | 14 | -223.202 | 490.958 | 33.168 | BVAR DCSTc | 3 | -60.749 |
| 88.130.147/ | 9 | -230.977 | 490.961 | 33.171 | BCST DCSTc | 2 | -174.322 |
| 104.130.137.147.155/95.108 | 20 | -209.85 | 490.99 | 33.2 | BVAR DCSTc | 3 | -15.158 |
| 88.95.111.147/ | 10 | -231.974 | 490.992 | 33.202 | BCST DCSTc | 2 | -111.089 |
| 104.111.137.147.155.160/95 | 19 | -214.326 | 490.994 | 33.204 | BVAR DCSTc | 3 | -63.149 |
| 88.111.147.155.160/ | 13 | -225.809 | 491.034 | 33.244 | BCST DCSTc | 2 | -112.915 |
| 130.155/ | 7 | -233.456 | 491.089 | 33.298 | BVARc | 2 | -206.026 |
| 104.130.147/108 | 12 | -225.29 | 491.099 | 33.309 | BVAR DCSTc | 3 | -60.749 |
| 88.104.130.147.160/ | 14 | -221.294 | 491.103 | 33.313 | BCST DCSTc | 2 | -131.095 |
| 111.130.137.147.155/ | 14 | -222.772 | 491.119 | 33.328 | BCST DCSTc | 2 | -124.844 |
| 88.130.137/ | 10 | -229.301 | 491.215 | 33.425 | BCST DCSTc | 2 | -173.645 |
| 88.130.137.147/108 | 15 | -221.845 | 491.243 | 33.453 | BVAR DCSTc | 3 | -44.968 |
| 104.130.137.155/110 | 15 | -219.891 | 491.268 | 33.478 | BCSTc | 1 | -70.448 |
| 104.137.147.155/95.110 | 15 | -220.414 | 491.294 | 33.504 | BCST DCSTc | 2 | -32.891 |
| 111.130.147.155/108 | 14 | -224.083 | 491.298 | 33.508 | BVAR DCSTc | 3 | -72.804 |
| 104.111.137.155.160/ | 14 | -222.353 | 491.31 | 33.52 | BCST DCSTc | 2 | -124.239 |
| 88.104.130.160/ | 12 | -224.625 | 491.318 | 33.528 | BCST DCSTc | 2 | -153.719 |
| 104.147.155.160/95.110 | 17 | -218.246 | 491.325 | 33.535 | BCST DCSTc | 2 | -32.891 |
| 104.130.137.160/ | 13 | -222.831 | 491.341 | 33.551 | BCST DCSTc | 2 | -156.93 |
| 88.104.111.130.137.155.160/ | 20 | -211.564 | 491.356 | 33.566 | BVAR DCSTc | 3 | -76.088 |
| 88.130.137.155.160/108 | 20 | -213.711 | 491.39 | 33.6 | BVAR DCSTc | 3 | -44.968 |
| 88.137.147.155.160/108 | 19 | -216.517 | 491.402 | 33.612 | BVAR DCSTc | 3 | -44.968 |
| 104.137.155/108 | 12 | -225.648 | 491.414 | 33.624 | BVAR DCSTc | 3 | -60.749 |
| 88.104.111.137.147.155.160/ | 19 | -214.407 | 491.442 | 33.652 | BVAR DCSTc | 3 | -73.701 |
| 130.155.160/108 | 14 | -224.022 | 491.512 | 33.722 | BVAR DCSTc | 3 | -72.804 |
| 111.130.137.155/108 | 15 | -222.364 | 491.528 | 33.738 | BVAR DCSTc | 3 | -72.804 |
| 88.104.137.147.155/110 | 16 | -218.493 | 491.567 | 33.777 | BVAR DCSTc | 3 | -41.22 |
| 88.104.111.137.155.160/108 | 19 | -213.679 | 491.587 | 33.797 | BVAR DCSTc | 3 | -31.012 |
| 104.111.130.137.147/95 | 16 | -218.945 | 491.59 | 33.8 | BCST DCSTc | 2 | -91.098 |
| 95.137.147.155/ | 11 | -228.932 | 491.628 | 33.838 | BCST DCSTc | 2 | -132.594 |
| 104.111.147.155/108 | 13 | -225.089 | 491.652 | 33.862 | BVAR DCSTc | 3 | -60.749 |
| 95.147.155/ | 8 | -233.97 | 491.654 | 33.864 | BVARc | 2 | -155.926 |
| 88.104.147.155.160/110 | 18 | -216.354 | 491.655 | 33.865 | BVAR DCSTc | 3 | -41.22 |

| | | | | | | | |
|---|---|---|---|---|---|---|---|
| 104.130.137.160/108 | 17 | -217.792 | 491.658 | 33.868 | BVAR DCSTc | 3 | -60.749 |
| 104.111.155.160/ | 11 | -227.605 | 491.723 | 33.933 | BCST DCSTc | 2 | -147.785 |
| 88.104.111.130.137.147/95 | 19 | -214.276 | 491.752 | 33.961 | BVAR DCSTc | 3 | -63.131 |
| 88.104.130.137.160/ | 15 | -219.832 | 491.792 | 34.002 | BCST DCSTc | 2 | -130.633 |
| 88.155.160/ | 9 | -231.899 | 491.837 | 34.047 | BCST DCSTc | 2 | -171.208 |
| 88.104.147.160/ | 11 | -227.685 | 491.838 | 34.048 | BCST DCSTc | 2 | -151.549 |
| 88.104.111.137.147/ | 13 | -224.946 | 491.856 | 34.066 | BCST DCSTc | 2 | -121.633 |
| 88.104.111.130.147.155.160/108 | 23 | -205.159 | 491.861 | 34.071 | BVAR DCSTc | 3 | -31.012 |
| 104.111.137.155/95 | 13 | -224.357 | 491.868 | 34.077 | BCST DCSTc | 2 | -116.501 |
| 88.95/ | 7 | -236.995 | 491.888 | 34.098 | BCST DVARc | 3 | -168.314 |
| 88.111.130.147.155/ | 13 | -224.971 | 491.896 | 34.105 | BCST DCSTc | 2 | -122.04 |
| 104.155.160/108 | 12 | -226.173 | 491.9 | 34.11 | BVAR DCSTc | 3 | -60.749 |
| 88.130.155.160/ | 12 | -225.931 | 491.935 | 34.144 | BCST DCSTc | 2 | -151.176 |
| 88.130.147.155/108 | 14 | -223.241 | 491.998 | 34.208 | BVAR DCSTc | 3 | -44.968 |
| 88.104.111.137.147.155/108 | 18 | -215.479 | 492.024 | 34.234 | BVAR DCSTc | 3 | -31.012 |
| 104.130.147.155/95.110 | 15 | -219.793 | 492.033 | 34.243 | BCST DCSTc | 2 | -32.891 |
| 88.95.137.147/ | 11 | -229.935 | 492.046 | 34.256 | BCST DCSTc | 2 | -123.665 |
| 88.130.137.155.160/ | 15 | -220.964 | 492.064 | 34.274 | BCST DCSTc | 2 | -127.915 |
| 88.111.130.137.155/ | 14 | -223.253 | 492.081 | 34.29 | BCST DCSTc | 2 | -121.321 |
| 130.137.155.160/108 | 18 | -217.803 | 492.106 | 34.316 | BVAR DCSTc | 3 | -72.804 |
| 104.130.137/108 | 13 | -223.993 | 492.118 | 34.328 | BVAR DCSTc | 3 | -60.749 |
| 88.155/ | 6 | -236.772 | 492.121 | 34.331 | BVARc | 2 | -200.107 |
| 137.147.155/108 | 12 | -228.027 | 492.121 | 34.331 | BVAR DCSTc | 3 | -72.804 |
| 88.95.155.160/ | 11 | -229.479 | 492.156 | 34.365 | BCST DCSTc | 2 | -123.404 |
| 88.104.111.147/ | 10 | -230.158 | 492.195 | 34.405 | BCST DCSTc | 2 | -145.139 |
| 130.137.155.160/ | 13 | -224.258 | 492.2 | 34.41 | BCST DCSTc | 2 | -154.507 |
| 88.137.147.155.160/ | 14 | -223.884 | 492.305 | 34.515 | BCST DCSTc | 2 | -125.605 |
| 88.104.111.137.147.155.160/108 | 23 | -205.239 | 492.307 | 34.516 | BVAR DCSTc | 3 | -31.012 |
| 104.130/ | 7 | -233.101 | 492.377 | 34.587 | BVARc | 2 | -209.52 |
| 147.155/ | 6 | -236.912 | 492.4 | 34.61 | BVARc | 2 | -204.251 |
| 137.147.155.160/108 | 17 | -220.762 | 492.425 | 34.634 | BVAR DCSTc | 3 | -72.804 |
| 104.111.130.137.147/ | 14 | -222.433 | 492.43 | 34.64 | BCST DCSTc | 2 | -128.355 |
| 104.137.147/95 | 11 | -227.897 | 492.434 | 34.643 | BCST DCSTc | 2 | -147.023 |
| 88.130.137.147/ | 12 | -226.71 | 492.47 | 34.68 | BCST DCSTc | 2 | -151.76 |
| 88.130.147.155.160/108 | 19 | -216.133 | 492.494 | 34.704 | BVAR DCSTc | 3 | -44.968 |
| 88.95.137/ | 9 | -233.392 | 492.515 | 34.725 | BCST DCSTc | 2 | -146.417 |
| 88.95.130.137.147.155/ | 17 | -218.885 | 492.551 | 34.76 | BVAR DCSTc | 3 | -85.185 |
| 88.104.111.130.137.147/108 | 19 | -213.701 | 492.553 | 34.763 | BVAR DCSTc | 3 | -31.012 |
| 104.111.130.155.160/ | 14 | -222.008 | 492.597 | 34.807 | BCST DCSTc | 2 | -128.124 |
| 88.95.130.147/ | 11 | -229.224 | 492.611 | 34.821 | BCST DCSTc | 2 | -127.185 |
| 104.111.130.137.155.160/ | 18 | -215.67 | 492.611 | 34.821 | BVAR DCSTc | 3 | -103.492 |
| 130.137.147/108 | 14 | -225.076 | 492.613 | 34.823 | BVAR DCSTc | 3 | -72.804 |
| 88.104.130.147.155/110 | 16 | -218.029 | 492.62 | 34.829 | BVAR DCSTc | 3 | -41.22 |

| | | | | | | | |
|---|---|---|---|---|---|---|---|
| 130.137.147.155.160/ | 16 | -220.084 | 492.629 | 34.838 | BVAR DCSTc | 3 | -131.04 |
| 95.130.147.155/ | 11 | -228.463 | 492.678 | 34.888 | BCST DCSTc | 2 | -136.355 |
| 88.147.155.160/108 | 15 | -223.68 | 492.694 | 34.904 | BVAR DCSTc | 3 | -44.968 |
| 88.130.147.155.160/ | 14 | -223.089 | 492.7 | 34.909 | BCST DCSTc | 2 | -129.04 |
| 104.137.147.155.160/95 | 16 | -219.26 | 492.712 | 34.921 | BCST DCSTc | 2 | -100.992 |
| 104.130.137.155.160/95 | 17 | -216.482 | 492.756 | 34.965 | BCST DCSTc | 2 | -103.445 |
| 104.130.155/108 | 12 | -225.332 | 492.772 | 34.982 | BVAR DCSTc | 3 | -60.749 |
| 130.147.155.160/108 | 17 | -220.008 | 492.778 | 34.988 | BVAR DCSTc | 3 | -72.804 |
| 104.111.130.155/ | 11 | -226.858 | 492.782 | 34.992 | BCST DCSTc | 2 | -157 |
| 137.147.155.160/ | 12 | -227.362 | 492.808 | 35.018 | BCST DCSTc | 2 | -152.381 |
| 88.104.137.155.160/108 | 17 | -216.92 | 492.833 | 35.042 | BVAR DCSTc | 3 | -31.012 |
| 104.130.147.155.160/95 | 16 | -218.349 | 492.87 | 35.08 | BCST DCSTc | 2 | -104.313 |
| 104.111.147.155.160/95 | 15 | -221.905 | 492.919 | 35.129 | BCST DCSTc | 2 | -89.023 |
| 88.104.160/95 | 11 | -228.432 | 492.937 | 35.147 | BCST DCSTc | 2 | -137.822 |
| 104.155/95 | 8 | -232.39 | 492.974 | 35.184 | BVARc | 2 | -175.737 |
| 95.130.137.147.155/ | 14 | -223.586 | 492.996 | 35.206 | BCST DCSTc | 2 | -113.184 |
| 88.104.111.130.137.155.160/108 | 24 | -202.799 | 493.026 | 35.236 | BVAR DCSTc | 3 | -31.012 |
| 104.137.155.160/95 | 14 | -222.688 | 493.097 | 35.307 | BCST DCSTc | 2 | -123.715 |
| 137.147.155/ | 9 | -232.247 | 493.101 | 35.311 | BCST DCSTc | 2 | -181.292 |
| 104.130.137.147.155/110 | 17 | -217.408 | 493.118 | 35.328 | BCSTc | 1 | -70.448 |
| 88.95.111.137.147.155/ | 16 | -222.524 | 493.151 | 35.361 | BVAR DCSTc | 3 | -69.978 |
| 95.130.137.155/ | 12 | -226.9 | 493.164 | 35.374 | BCST DCSTc | 2 | -135.792 |
| 104.130.137.155/95.110 | 18 | -216.102 | 493.168 | 35.378 | BCST DCSTc | 2 | -32.891 |
| 104.147.155/108 | 11 | -228.358 | 493.223 | 35.433 | BVAR DCSTc | 3 | -60.749 |
| 88.130.137.155/108 | 15 | -222.049 | 493.232 | 35.442 | BVAR DCSTc | 3 | -44.968 |
| 88.104.111.130.147/ | 13 | -224.65 | 493.242 | 35.452 | BCST DCSTc | 2 | -125.567 |
| 88.95.111.130.147.155/ | 16 | -221.664 | 493.281 | 35.491 | BVAR DCSTc | 3 | -73.348 |
| 147.155.160/108 | 13 | -227.723 | 493.313 | 35.523 | BVAR DCSTc | 3 | -72.804 |
| 88.104.130.137.155/110 | 19 | -214.14 | 493.358 | 35.568 | BVAR DCSTc | 3 | -41.22 |
| 104.111.137.147.155.160/ | 17 | -218.846 | 493.365 | 35.575 | BVAR DCSTc | 3 | -101.438 |
| 88.111.147.155.160/108 | 18 | -219.645 | 493.385 | 35.594 | BVAR DCSTc | 3 | -44.968 |
| 88.95.130.137/ | 12 | -227.806 | 493.388 | 35.598 | BCST DCSTc | 2 | -126.767 |
| 104.111.130.155/95 | 13 | -224.137 | 493.406 | 35.616 | BCST DCSTc | 2 | -120.511 |
| 88.104.111.130.137/ | 14 | -222.922 | 493.407 | 35.617 | BCST DCSTc | 2 | -124.839 |
| 88.95.130/ | 9 | -232.848 | 493.417 | 35.627 | BCST DCSTc | 2 | -150.103 |
| 104.111.130.147.155.160/108 | 21 | -210.619 | 493.427 | 35.637 | BVAR DCSTc | 3 | -60.749 |
| 88.104.111.130.155.160/108 | 19 | -213.647 | 493.429 | 35.639 | BVAR DCSTc | 3 | -31.012 |
| 130.147.155/108 | 12 | -227.711 | 493.471 | 35.68 | BVAR DCSTc | 3 | -72.804 |
| 88.104/ | 6 | -236.467 | 493.507 | 35.717 | BVARc | 2 | -203.651 |
| 88.104.111.137.155.160/ | 17 | -218.969 | 493.611 | 35.82 | BVAR DCSTc | 3 | -97.557 |
| 88.137.155/ | 9 | -232.511 | 493.629 | 35.839 | BCST DCSTc | 2 | -177.552 |
| 130.147.155.160/ | 12 | -226.79 | 493.653 | 35.863 | BCST DCSTc | 2 | -156.039 |
| 104.130.147/95 | 11 | -227.519 | 493.67 | 35.879 | BCST DCSTc | 2 | -150.876 |

| | | | | | | | |
|---|---|---|---|---|---|---|---|
| 104.147/ | 6 | -236.569 | 493.711 | 35.921 | BVARc | 2 | -207.757 |
| 88.111.130.137.147.155/ | 17 | -219.549 | 493.742 | 35.951 | BVAR DCSTc | 3 | -98.323 |
| 147.155.160/ | 9 | -232.853 | 493.746 | 35.956 | BCST DCSTc | 2 | -176.166 |
| 88.95.147/ | 8 | -235.828 | 493.778 | 35.988 | BCST DCSTc | 2 | -147.852 |
| 104.111.130.147.155.160/ | 17 | -218.11 | 493.809 | 36.019 | BVAR DCSTc | 3 | -104.933 |
| 111.147.155.160/108 | 16 | -223.62 | 493.87 | 36.079 | BVAR DCSTc | 3 | -72.804 |
| 104.130.155.160/95 | 14 | -222.082 | 493.872 | 36.082 | BCST DCSTc | 2 | -127.34 |
| 88.104.111.130.147.155/108 | 18 | -215.502 | 494.005 | 36.215 | BVAR DCSTc | 3 | -31.012 |
| 104.111.137.147.155/ | 13 | -225.247 | 494.038 | 36.247 | BCST DCSTc | 2 | -131.865 |
| 104.130.137/95 | 12 | -225.909 | 494.059 | 36.269 | BCST DCSTc | 2 | -150.266 |
| 88.104.130.155.160/108 | 17 | -216.56 | 494.091 | 36.301 | BVAR DCSTc | 3 | -31.012 |
| 88.104.111.130.137.155/108 | 19 | -213.729 | 494.126 | 36.335 | BVAR DCSTc | 3 | -31.012 |
| 88.104.137.147.155/108 | 16 | -219.11 | 494.154 | 36.363 | BVAR DCSTc | 3 | -31.012 |
| 88.95.111.130.137.155/ | 17 | -220.242 | 494.188 | 36.397 | BVAR DCSTc | 3 | -72.927 |
| 88.147.155.160/ | 11 | -229.869 | 494.211 | 36.421 | BCST DCSTc | 2 | -149.883 |
| 104.155.160/95 | 11 | -228.303 | 494.27 | 36.48 | BCST DCSTc | 2 | -147.624 |
| 88.104.111.130.155.160/ | 17 | -218.379 | 494.346 | 36.556 | BVAR DCSTc | 3 | -101.197 |
| 104.137.147/ | 9 | -231.909 | 494.422 | 36.632 | BCST DCSTc | 2 | -184.803 |
| 104.111.137.147.155/95 | 15 | -222.381 | 494.424 | 36.634 | BCST DCSTc | 2 | -95.231 |
| 104.111.147.155/ | 10 | -230.518 | 494.503 | 36.712 | BCST DCSTc | 2 | -155.43 |
| 130.137.155/108 | 13 | -226.453 | 494.573 | 36.783 | BVAR DCSTc | 3 | -72.804 |
| 88.104.111.155.160/ | 13 | -225.82 | 494.621 | 36.831 | BCST DCSTc | 2 | -122.702 |
| 104.111.147.155/95 | 12 | -227.574 | 494.682 | 36.892 | BCST DCSTc | 2 | -118.718 |
| 130.147.155/ | 9 | -232.043 | 494.685 | 36.895 | BCST DCSTc | 2 | -185.318 |
| 104.130.137.147/95 | 14 | -223.011 | 494.708 | 36.918 | BCST DCSTc | 2 | -128.074 |
| 88.104.130.137.147/108 | 17 | -217.407 | 494.766 | 36.976 | BVAR DCSTc | 3 | -31.012 |
| 130.137.155/ | 10 | -230.288 | 494.782 | 36.992 | BCST DCSTc | 2 | -184.563 |
| 104.111.137.155.160/108 | 17 | -219.985 | 494.845 | 37.055 | BVAR DCSTc | 3 | -60.749 |
| 88.104.130.137.155.160/108 | 22 | -209.273 | 494.913 | 37.123 | BVAR DCSTc | 3 | -31.012 |
| 88.104.137.147.155.160/108 | 21 | -212.079 | 494.925 | 37.135 | BVAR DCSTc | 3 | -31.012 |
| 88.104.137/ | 9 | -232.176 | 494.956 | 37.166 | BCST DCSTc | 2 | -181.066 |
| 88.95.111.137.155/ | 13 | -228.119 | 494.99 | 37.2 | BCST DCSTc | 2 | -94.867 |
| 104.130.137.147.155/95.110 | 20 | -213.619 | 495.019 | 37.229 | BCST DCSTc | 2 | -32.891 |
| 88.130.155/ | 9 | -232.276 | 495.152 | 37.362 | BCST DCSTc | 2 | -181.548 |
| 88.104.137.155.160/95 | 16 | -220.503 | 495.198 | 37.408 | BCST DCSTc | 2 | -98.232 |
| 88.104.130.155.160/95 | 16 | -219.519 | 495.209 | 37.419 | BCST DCSTc | 2 | -101.478 |
| 88.104.111.137.155/ | 13 | -225.835 | 495.213 | 37.423 | BCST DCSTc | 2 | -128.449 |
| 88.104.111.130.137.147/ | 17 | -219.317 | 495.235 | 37.445 | BVAR DCSTc | 3 | -101.941 |
| 104.111.137.147.155/108 | 16 | -221.783 | 495.278 | 37.488 | BVAR DCSTc | 3 | -60.749 |
| 88.104.130.137.147.155/110 | 21 | -211.7 | 495.295 | 37.505 | BVAR DCSTc | 3 | -41.22 |
| 88.95.111.130.155/ | 13 | -227.326 | 495.371 | 37.581 | BCST DCSTc | 2 | -98.305 |
| 104.111.130.147.155/95 | 15 | -221.894 | 495.417 | 37.627 | BCST DCSTc | 2 | -98.974 |
| 88.104.147.155.160/95 | 15 | -222.456 | 495.483 | 37.693 | BCST DCSTc | 2 | -99.184 |

| | | | | | | | |
|---|---|---|---|---|---|---|---|
| 88.104.111.155/ | 10 | -231.015 | 495.496 | 37.706 | BCST DCSTc | 2 | -151.923 |
| 88.104.130.147.155/108 | 16 | -218.803 | 495.521 | 37.731 | BVAR DCSTc | 3 | -31.012 |
| 88.137.147.155/ | 11 | -230.275 | 495.59 | 37.799 | BCST DCSTc | 2 | -156.022 |
| 104.111.130.137.155.160/108 | 22 | -208.762 | 495.598 | 37.808 | BVAR DCSTc | 3 | -60.749 |
| 104.111.130.137.155/ | 14 | -223.233 | 495.608 | 37.818 | BCST DCSTc | 2 | -135.081 |
| 104.147.155.160/95 | 13 | -225.762 | 495.631 | 37.841 | BCST DCSTc | 2 | -125.789 |
| 88.95.111.155/ | 10 | -233.545 | 495.715 | 37.924 | BCST DCSTc | 2 | -118.587 |
| 104.111.130.147.155/ | 13 | -225.123 | 495.77 | 37.98 | BCST DCSTc | 2 | -135.971 |
| 104.111.130.137.147/108 | 17 | -220.002 | 495.8 | 38.01 | BVAR DCSTc | 3 | -60.749 |
| 104.111.137.147.155.160/108 | 21 | -211.678 | 495.83 | 38.04 | BVAR DCSTc | 3 | -60.749 |
| 104.111.130.137.155/95 | 16 | -220.309 | 495.88 | 38.09 | BCST DCSTc | 2 | -98.39 |
| 88.95.111.147.155/ | 12 | -230.444 | 496.008 | 38.218 | BCST DCSTc | 2 | -96.192 |
| 88.104.130.147.155.160/108 | 21 | -211.694 | 496.017 | 38.227 | BVAR DCSTc | 3 | -31.012 |
| 104.130.147/ | 9 | -231.71 | 496.017 | 38.227 | BCST DCSTc | 2 | -188.835 |
| 88.147.155/ | 8 | -235.515 | 496.031 | 38.241 | BCST DCSTc | 2 | -179.556 |
| 88.104.111.155.160/95 | 15 | -223.491 | 496.09 | 38.3 | BCST DCSTc | 2 | -86.604 |
| 104.130.137/ | 10 | -229.952 | 496.107 | 38.317 | BCST DCSTc | 2 | -188.076 |
| 104.155/ | 6 | -236.982 | 496.133 | 38.343 | BVARc | 2 | -214.097 |
| 88.104.147.155.160/108 | 17 | -219.241 | 496.217 | 38.427 | BVAR DCSTc | 3 | -31.012 |
| 88.104.111.137/95 | 13 | -227.326 | 496.227 | 38.437 | BCST DCSTc | 2 | -109.539 |
| 104.111.130.155.160/108 | 17 | -219.733 | 496.247 | 38.457 | BVAR DCSTc | 3 | -60.749 |
| 104.137.155/95 | 11 | -229.017 | 496.264 | 38.474 | BVARc | 2 | -154.07 |
| 104.137.155.160/108 | 16 | -222.132 | 496.288 | 38.498 | BVAR DCSTc | 3 | -60.749 |
| 88.104.111.147.155.160/ | 16 | -222.189 | 496.367 | 38.576 | BVAR DCSTc | 3 | -99.777 |
| 104.130.137.147.155.160/ | 18 | -216.913 | 496.382 | 38.592 | BVAR DCSTc | 3 | -118.351 |
| 88.104.111.137.147/95 | 15 | -224.165 | 496.43 | 38.64 | BCST DCSTc | 2 | -87.084 |
| 88.95.137.147.155/ | 13 | -228.114 | 496.452 | 38.662 | BCST DCSTc | 2 | -108.477 |
| 88.104.130/ | 9 | -231.947 | 496.49 | 38.7 | BCST DCSTc | 2 | -185.067 |
| 88.104.111.130.147/95 | 15 | -223.215 | 496.491 | 38.701 | BCST DCSTc | 2 | -90.364 |
| 104.111.147.155.160/ | 14 | -225.568 | 496.492 | 38.702 | BVAR DCSTc | 3 | -126.454 |
| 88.104.111/95 | 10 | -232.554 | 496.594 | 38.804 | BCST DCSTc | 2 | -133.061 |
| 104.137.155.160/ | 12 | -227.52 | 496.709 | 38.919 | BCST DCSTc | 2 | -162.315 |
| 104.111.130.147.155/108 | 16 | -221.546 | 496.737 | 38.947 | BVAR DCSTc | 3 | -60.749 |
| 130.137.147.155/ | 12 | -228.049 | 496.739 | 38.949 | BCST DCSTc | 2 | -163.031 |
| 88.95.130.147.155/ | 13 | -227.273 | 496.752 | 38.962 | BCST DCSTc | 2 | -111.867 |
| 88.104.130.137.155/108 | 17 | -217.611 | 496.755 | 38.965 | BVAR DCSTc | 3 | -31.012 |
| 88.104.137.147.155.160/ | 17 | -219.9 | 496.757 | 38.966 | BVAR DCSTc | 3 | -112.103 |
| 88.104.111.130.155/ | 13 | -225.654 | 496.832 | 39.042 | BCST DCSTc | 2 | -132.499 |
| 88.130.147.155/ | 11 | -229.901 | 496.833 | 39.043 | BCST DCSTc | 2 | -159.878 |
| 88.104.130.137.155.160/ | 18 | -217.153 | 496.862 | 39.072 | BVAR DCSTc | 3 | -114.587 |
| 104.111.130.137.147.155/95 | 18 | -217.511 | 496.889 | 39.099 | BCST DCSTc | 2 | -76.298 |
| 88.104.111.147.155.160/108 | 20 | -215.206 | 496.907 | 39.117 | BVAR DCSTc | 3 | -31.012 |
| 88.104.137.147/ | 11 | -229.956 | 496.948 | 39.158 | BCST DCSTc | 2 | -159.552 |

| | | | | | | | |
|---|---|---|---|---|---|---|---|
| 104.130.155.160/108 | 16 | -221.485 | 496.951 | 39.161 | BVAR DCSTc | 3 | -60.749 |
| 104.111.130.137.155/108 | 17 | -219.827 | 496.967 | 39.177 | BVAR DCSTc | 3 | -60.749 |
| 88.104.111.130/95 | 13 | -226.719 | 496.991 | 39.201 | BCST DCSTc | 2 | -113.162 |
| 88.104.130.137.147/95 | 16 | -220.967 | 497.086 | 39.296 | BCST DCSTc | 2 | -102.731 |
| 88.104.130.147.155.160/ | 17 | -219.12 | 497.154 | 39.364 | BVAR DCSTc | 3 | -115.553 |
| 88.130.137.155/ | 12 | -228.271 | 497.183 | 39.393 | BCST DCSTc | 2 | -159.249 |
| 104.155.160/ | 9 | -232.811 | 497.251 | 39.461 | BCST DCSTc | 2 | -185.9 |
| 88.111.130.137.147.155/108 | 19 | -218.059 | 497.317 | 39.527 | BVAR DCSTc | 3 | -44.968 |
| 88.104.147/ | 8 | -235.19 | 497.376 | 39.586 | BCST DCSTc | 2 | -183.08 |
| 104.130.155/95 | 11 | -228.655 | 497.533 | 39.743 | BVARc | 2 | -157.939 |
| 104.130.137.155.160/108 | 20 | -215.266 | 497.545 | 39.755 | BVAR DCSTc | 3 | -60.749 |
| 104.137.147.155/108 | 14 | -225.49 | 497.56 | 39.77 | BVAR DCSTc | 3 | -60.749 |
| 88.104.111.130.137/95 | 16 | -221.96 | 497.619 | 39.829 | BCST DCSTc | 2 | -90.109 |
| 88.95.130.137.155/ | 14 | -225.907 | 497.637 | 39.847 | BCST DCSTc | 2 | -111.501 |
| 88.104.111.147/95 | 12 | -229.877 | 497.706 | 39.916 | BCST DCSTc | 2 | -111.089 |
| 88.104.137.155.160/ | 14 | -224.816 | 497.746 | 39.956 | BCST DCSTc | 2 | -136.313 |
| 104.137.147.155.160/108 | 19 | -218.225 | 497.864 | 40.073 | BVAR DCSTc | 3 | -60.749 |
| 88.104.111.137.147.155/ | 15 | -223.918 | 497.881 | 40.091 | BCST DCSTc | 2 | -107.238 |
| 104.130.155.160/ | 12 | -227.116 | 497.891 | 40.1 | BCST DCSTc | 2 | -166.141 |
| 88.95.137.155/ | 11 | -232.066 | 497.895 | 40.105 | BCST DCSTc | 2 | -131.723 |
| 88.130.137.147.155/ | 14 | -225.417 | 497.919 | 40.129 | BCST DCSTc | 2 | -137.1 |
| 104.130.137.147/108 | 16 | -222.539 | 498.052 | 40.262 | BVAR DCSTc | 3 | -60.749 |
| 104.130.137.147/ | 12 | -227.731 | 498.098 | 40.308 | BCST DCSTc | 2 | -166.562 |
| 104.111.130.137.147.155/ | 17 | -219.99 | 498.103 | 40.313 | BVAR DCSTc | 3 | -112.544 |
| 104.137.155/ | 9 | -232.959 | 498.114 | 40.324 | BCST DCSTc | 2 | -191.78 |
| 88.130.137.147.155/108 | 18 | -220.043 | 498.174 | 40.384 | BVAR DCSTc | 3 | -44.968 |
| 88.104.130.147/ | 11 | -229.588 | 498.203 | 40.412 | BCST DCSTc | 2 | -163.415 |
| 104.130.147.155.160/108 | 19 | -217.471 | 498.217 | 40.427 | BVAR DCSTc | 3 | -60.749 |
| 88.104.111.147.155/ | 12 | -229.182 | 498.289 | 40.499 | BCST DCSTc | 2 | -130.796 |
| 104.137.147.155/95 | 13 | -226.835 | 498.342 | 40.552 | BCST DCSTc | 2 | -132.594 |
| 104.147.155/95 | 10 | -231.873 | 498.368 | 40.578 | BVARc | 2 | -155.926 |
| 111.130.137.147.155/108 | 17 | -222.348 | 498.429 | 40.639 | BVAR DCSTc | 3 | -72.804 |
| 88.95.155/ | 9 | -236.277 | 498.499 | 40.708 | BCST DVARc | 3 | -154.229 |
| 88.104.130.155.160/ | 14 | -224.203 | 498.507 | 40.717 | BCST DCSTc | 2 | -139.93 |
| 88.104.130.137/ | 12 | -227.956 | 498.547 | 40.757 | BCST DCSTc | 2 | -162.783 |
| 88.95.130.155/ | 11 | -231.422 | 498.596 | 40.806 | BCST DCSTc | 2 | -135.31 |
| 88.104/95 | 9 | -234.898 | 498.602 | 40.812 | BCST DVARc | 3 | -168.314 |
| 104.130.137.155.160/ | 16 | -221.346 | 498.705 | 40.915 | BVAR DCSTc | 3 | -142.077 |
| 104.147.155.160/108 | 15 | -225.186 | 498.752 | 40.962 | BVAR DCSTc | 3 | -60.749 |
| 88.104.137.147/95 | 13 | -227.837 | 498.76 | 40.97 | BCST DCSTc | 2 | -123.665 |
| 88.95.147.155/ | 10 | -234.358 | 498.868 | 41.077 | BCST DCSTc | 2 | -133.015 |
| 88.104.155.160/95 | 13 | -227.381 | 498.87 | 41.079 | BCST DCSTc | 2 | -123.404 |
| 104.130.147.155/108 | 14 | -225.174 | 498.91 | 41.119 | BVAR DCSTc | 3 | -60.749 |

| | | | | | | | |
|---|---|---|---|---|---|---|---|
| 88.104.155.160/ | 11 | -230.425 | 498.909 | 41.119 | BCST DCSTc | 2 | -160.216 |
| 88.104.111.130.147.155/ | 15 | -223.466 | 498.947 | 41.156 | BCST DCSTc | 2 | -111.017 |
| 104.137.147.155.160/ | 15 | -224.307 | 499.028 | 41.238 | BVAR DCSTc | 3 | -139.808 |
| 130.137.147.155/108 | 16 | -224.21 | 499.043 | 41.253 | BVAR DCSTc | 3 | -72.804 |
| 88.104.137/95 | 11 | -231.295 | 499.229 | 41.439 | BCST DCSTc | 2 | -146.417 |
| 88.104.111.130.137.147.155/ | 19 | -217.149 | 499.231 | 41.441 | BVAR DCSTc | 3 | -86.405 |
| 88.104.130.137.147.155/95 | 19 | -216.788 | 499.265 | 41.474 | BVAR DCSTc | 3 | -85.185 |
| 104.111.147.155.160/108 | 18 | -221.084 | 499.309 | 41.518 | BVAR DCSTc | 3 | -60.749 |
| 88.104.111.130.137.155/ | 16 | -221.833 | 499.31 | 41.52 | BCST DCSTc | 2 | -110.383 |
| 88.104.130.137.147/ | 14 | -225.122 | 499.323 | 41.533 | BCST DCSTc | 2 | -140.655 |
| 88.104.130.147/95 | 13 | -227.126 | 499.325 | 41.535 | BCST DCSTc | 2 | -127.185 |
| 104.130.147.155/95 | 13 | -226.366 | 499.392 | 41.602 | BCST DCSTc | 2 | -136.355 |
| 104.130.137.147.155/95 | 16 | -221.488 | 499.71 | 41.92 | BCST DCSTc | 2 | -113.184 |
| 104.130.155/ | 9 | -232.818 | 499.826 | 42.035 | BCST DCSTc | 2 | -195.869 |
| 104.130.147.155.160/ | 15 | -223.725 | 499.837 | 42.047 | BVAR DCSTc | 3 | -143.457 |
| 88.104.111.137.147.155/95 | 18 | -220.427 | 499.865 | 42.075 | BVAR DCSTc | 3 | -69.978 |
| 104.130.137.155/95 | 14 | -224.803 | 499.878 | 42.088 | BCST DCSTc | 2 | -135.792 |
| 88.104.111.130.147.155/95 | 18 | -219.566 | 499.995 | 42.205 | BVAR DCSTc | 3 | -73.348 |
| 104.130.137.155/108 | 15 | -223.917 | 500.012 | 42.222 | BVAR DCSTc | 3 | -60.749 |
| 88.104.130.137/95 | 14 | -225.708 | 500.102 | 42.312 | BCST DCSTc | 2 | -126.767 |
| 88.104.130/95 | 11 | -230.75 | 500.131 | 42.341 | BCST DCSTc | 2 | -150.103 |
| 88.104.147/95 | 10 | -233.731 | 500.492 | 42.702 | BCST DCSTc | 2 | -147.852 |
| 88.104.147.155.160/ | 13 | -228.011 | 500.524 | 42.733 | BCST DCSTc | 2 | -138.508 |
| 104.137.147.155/ | 11 | -231.013 | 500.653 | 42.862 | BCST DCSTc | 2 | -170.54 |
| 104.147.155.160/ | 11 | -231.334 | 500.728 | 42.938 | BCST DCSTc | 2 | -165.129 |
| 88.104.111.130.137.147.155/108 | 21 | -213.62 | 500.84 | 43.05 | BVAR DCSTc | 3 | -31.012 |
| 88.104.111.130.137.155/95 | 19 | -218.145 | 500.902 | 43.111 | BVAR DCSTc | 3 | -72.927 |
| 104.147.155/ | 8 | -236.163 | 500.917 | 43.126 | BCST DCSTc | 2 | -193.984 |
| 88.104.137.155/ | 11 | -231.289 | 501.205 | 43.415 | BCST DCSTc | 2 | -166.813 |
| 88.104.155/ | 8 | -236.371 | 501.333 | 43.542 | BCST DCSTc | 2 | -190.188 |
| 88.104.130.137.147.155/108 | 20 | -215.604 | 501.697 | 43.907 | BVAR DCSTc | 3 | -31.012 |
| 88.104.111.137.155/95 | 15 | -226.021 | 501.704 | 43.914 | BCST DCSTc | 2 | -94.867 |
| 88.104.111.130.155/95 | 15 | -225.228 | 502.085 | 44.295 | BCST DCSTc | 2 | -98.305 |
| 104.130.147.155/ | 11 | -230.742 | 502.102 | 44.311 | BCST DCSTc | 2 | -174.499 |
| 104.130.137.155/ | 12 | -229.026 | 502.279 | 44.489 | BCST DCSTc | 2 | -173.784 |
| 88.104.111.155/95 | 12 | -231.448 | 502.429 | 44.638 | BCST DCSTc | 2 | -118.587 |
| 88.104.130.155/ | 11 | -230.988 | 502.594 | 44.804 | BCST DCSTc | 2 | -170.742 |
| 88.104.111.147.155/95 | 14 | -228.347 | 502.722 | 44.932 | BCST DCSTc | 2 | -96.192 |
| 88.104.137.147.155/ | 13 | -228.853 | 502.773 | 44.983 | BCST DCSTc | 2 | -145.082 |
| 88.104.137.147.155/95 | 15 | -226.016 | 503.166 | 45.376 | BCST DCSTc | 2 | -108.477 |
| 88.104.147.155/ | 10 | -234.209 | 503.436 | 45.646 | BCST DCSTc | 2 | -168.732 |
| 88.104.130.147.155/95 | 15 | -225.175 | 503.466 | 45.676 | BCST DCSTc | 2 | -111.867 |
| 88.104.130.147.155/ | 13 | -228.389 | 503.833 | 46.043 | BCST DCSTc | 2 | -148.848 |

| | | | | | | | |
|---|---|---|---|---|---|---|---|
| 104.111.130.137.147.155/108 | 19 | -219.811 | 503.868 | 46.078 | BVAR DCSTc | 3 | -60.749 |
| 104.130.137.147.155/ | 14 | -226.62 | 503.906 | 46.116 | BCST DCSTc | 2 | -152.083 |
| 88.104.130.137.155/ | 14 | -226.813 | 504.292 | 46.502 | BCST DCSTc | 2 | -148.272 |
| 88.104.130.137.155/95 | 16 | -223.809 | 504.351 | 46.561 | BCST DCSTc | 2 | -111.501 |
| 88.104.130.137.147.155/ | 16 | -223.664 | 504.453 | 46.663 | BCST DCSTc | 2 | -125.83 |
| 104.130.137.147.155/108 | 18 | -221.673 | 504.482 | 46.692 | BVAR DCSTc | 3 | -60.749 |
| 88.104.137.155/95 | 13 | -229.968 | 504.609 | 46.819 | BCST DCSTc | 2 | -131.723 |
| 88.104.155/95 | 11 | -234.18 | 505.213 | 47.422 | BCST DVARc | 3 | -154.229 |
| 88.104.130.155/95 | 13 | -229.325 | 505.31 | 47.52 | BCST DCSTc | 2 | -135.31 |
| 88.104.147.155/95 | 12 | -232.26 | 505.582 | 47.791 | BCST DCSTc | 2 | -133.015 |

ıt the crown and with a maximum rate value
iple backbones, only the deepest is displayed
re highlighted in bold. NP = Number of
of speciation rate, μ = extinction rate, β =

**ɔackbone when multiple backbones)**

| AICc | λ | α | μ | β |
|---|---|---|---|---|
| **42.316** | **0.138** | **0.069** | **0.736** | **–** |
| **50.24** | **1.532** | **–** | **2** | **-0.014** |
| **42.316** | **0.138** | **0.069** | **0.736** | **–** |
| 42.316 | 0.138 | 0.069 | 0.736 | – |
| 42.316 | 0.138 | 0.069 | 0.736 | – |
| 88.232 | 1.367 | 0.01 | 1.685 | – |
| 42.316 | 0.138 | 0.069 | 0.736 | – |
| 42.316 | 0.138 | 0.069 | 0.736 | – |
| 42.316 | 0.138 | 0.069 | 0.736 | – |
| 42.316 | 0.138 | 0.069 | 0.736 | – |
| 42.316 | 0.138 | 0.069 | 0.736 | – |
| 42.316 | 0.138 | 0.069 | 0.736 | – |
| 165.584 | 0.138 | – | 0.108 | – |
| 42.316 | 0.138 | 0.069 | 0.736 | – |
| 42.316 | 0.138 | 0.069 | 0.736 | – |
| 42.316 | 0.138 | 0.069 | 0.736 | – |
| 42.316 | 0.138 | 0.069 | 0.736 | – |
| 50.24 | 1.532 | – | 2 | -0.014 |
| 42.316 | 0.138 | 0.069 | 0.736 | – |
| 42.316 | 0.138 | 0.069 | 0.736 | – |
| 165.584 | 0.138 | – | 0.108 | – |
| 42.316 | 0.138 | 0.069 | 0.736 | – |
| 42.316 | 0.138 | 0.069 | 0.736 | – |
| 71.496 | 0.23 | – | 0.23 | – |
| 42.316 | 0.138 | 0.069 | 0.736 | – |
| 42.316 | 0.138 | 0.069 | 0.736 | – |
| 42.316 | 0.138 | 0.069 | 0.736 | – |
| 42.316 | 0.138 | 0.069 | 0.736 | – |
| 115.419 | 0.228 | 0.027 | 0.384 | – |
| 129.428 | 1.421 | 0.009 | 1.714 | – |
| 42.316 | 0.138 | 0.069 | 0.736 | – |
| 88.232 | 1.367 | 0.01 | 1.685 | – |
| 42.316 | 0.138 | 0.069 | 0.736 | – |
| 42.316 | 0.138 | 0.069 | 0.736 | – |
| 69.558 | 1.26 | 0.012 | 1.629 | – |
| 42.316 | 0.138 | 0.069 | 0.736 | – |
| 71.496 | 0.23 | – | 0.23 | – |
| 42.316 | 0.138 | 0.069 | 0.736 | – |

| | | | | |
|---|---|---|---|---|
| 65.649 | 1.602 | – | 2 | -0.011 |
| 42.316 | 0.138 | 0.069 | 0.736 | – |
| 72.207 | 0.737 | 0.026 | 1.292 | – |
| 42.316 | 0.138 | 0.069 | 0.736 | – |
| 42.316 | 0.138 | 0.069 | 0.736 | – |
| 42.316 | 0.138 | 0.069 | 0.736 | – |
| 115.419 | 0.228 | 0.027 | 0.384 | – |
| 42.316 | 0.138 | 0.069 | 0.736 | – |
| 165.584 | 0.138 | – | 0.108 | – |
| 42.316 | 0.138 | 0.069 | 0.736 | – |
| 42.316 | 0.138 | 0.069 | 0.736 | – |
| 165.584 | 0.138 | – | 0.108 | – |
| 42.316 | 0.138 | 0.069 | 0.736 | – |
| 42.316 | 0.138 | 0.069 | 0.736 | – |
| 165.584 | 0.138 | – | 0.108 | – |
| 42.316 | 0.138 | 0.069 | 0.736 | – |
| 108.379 | 1.316 | 0.011 | 1.654 | – |
| 165.584 | 0.138 | – | 0.108 | – |
| 165.584 | 0.138 | – | 0.108 | – |
| 42.316 | 0.138 | 0.069 | 0.736 | – |
| 102.207 | 1.136 | 0.013 | 1.51 | – |
| 165.584 | 0.138 | – | 0.108 | – |
| 42.316 | 0.138 | 0.069 | 0.736 | – |
| 42.316 | 0.138 | 0.069 | 0.736 | – |
| 42.316 | 0.138 | 0.069 | 0.736 | – |
| 98.337 | 0.347 | 0.03 | 0.651 | – |
| 143.132 | 0.067 | – | – | – |
| 165.584 | 0.138 | – | 0.108 | – |
| 42.316 | 0.138 | 0.069 | 0.736 | – |
| 42.316 | 0.138 | 0.069 | 0.736 | – |
| 42.316 | 0.138 | 0.069 | 0.736 | – |
| 42.316 | 0.138 | 0.069 | 0.736 | – |
| 81.096 | 1.514 | 0.007 | 1.759 | – |
| 42.316 | 0.138 | 0.069 | 0.736 | – |
| 165.584 | 0.138 | – | 0.108 | – |
| 42.316 | 0.138 | 0.069 | 0.736 | – |
| 71.496 | 0.23 | – | 0.23 | – |
| 108.079 | 1.343 | 0.01 | 1.668 | – |
| 165.584 | 0.138 | – | 0.108 | – |
| 152.94 | 0.195 | 0.024 | 0.296 | – |
| 42.316 | 0.138 | 0.069 | 0.736 | – |
| 42.316 | 0.138 | 0.069 | 0.736 | – |
| 42.316 | 0.138 | 0.069 | 0.736 | – |

| | | | | |
|---|---|---|---|---|
| 42.316 | 0.138 | 0.069 | 0.736 | – |
| 42.316 | 0.138 | 0.069 | 0.736 | – |
| 71.496 | 0.23 | – | 0.23 | – |
| 42.316 | 0.138 | 0.069 | 0.736 | – |
| 98.337 | 0.347 | 0.03 | 0.651 | – |
| 143.132 | 0.067 | – | – | – |
| 71.496 | 0.23 | – | 0.23 | – |
| 115.419 | 0.228 | 0.027 | 0.384 | – |
| 42.316 | 0.138 | 0.069 | 0.736 | – |
| 71.496 | 0.23 | – | 0.23 | – |
| 91.868 | 0.235 | 0.031 | 0.437 | – |
| 42.316 | 0.138 | 0.069 | 0.736 | – |
| 71.496 | 0.23 | – | 0.23 | – |
| 71.496 | 0.23 | – | 0.23 | – |
| 71.496 | 0.23 | – | 0.23 | – |
| 98.337 | 0.347 | 0.03 | 0.651 | – |
| 115.419 | 0.228 | 0.027 | 0.384 | – |
| 71.496 | 0.23 | – | 0.23 | – |
| 165.584 | 0.138 | – | 0.108 | – |
| 42.316 | 0.138 | 0.069 | 0.736 | – |
| 71.496 | 0.23 | – | 0.23 | – |
| 42.316 | 0.138 | 0.069 | 0.736 | – |
| 42.316 | 0.138 | 0.069 | 0.736 | – |
| 42.316 | 0.138 | 0.069 | 0.736 | – |
| 42.316 | 0.138 | 0.069 | 0.736 | – |
| 115.419 | 0.228 | 0.027 | 0.384 | – |
| 71.496 | 0.23 | – | 0.23 | – |
| 42.316 | 0.138 | 0.069 | 0.736 | – |
| 115.419 | 0.228 | 0.027 | 0.384 | – |
| 152.94 | 0.195 | 0.024 | 0.296 | – |
| 403.398 | 0.172 | -0.034 | – | – |
| 115.419 | 0.228 | 0.027 | 0.384 | – |
| 98.337 | 0.347 | 0.03 | 0.651 | – |
| 115.419 | 0.228 | 0.027 | 0.384 | – |
| 129.428 | 1.421 | 0.009 | 1.714 | – |
| 98.337 | 0.347 | 0.03 | 0.651 | – |
| 115.419 | 0.228 | 0.027 | 0.384 | – |
| 152.94 | 0.195 | 0.024 | 0.296 | – |
| 299.716 | 0.189 | – | 0.136 | – |
| 72.824 | 0.447 | 0.03 | 0.856 | – |
| 115.419 | 0.228 | 0.027 | 0.384 | – |
| 71.496 | 0.23 | – | 0.23 | – |
| 69.558 | 1.26 | 0.012 | 1.629 | – |

| | | | | |
|---|---|---|---|---|
| 42.316 | 0.138 | 0.069 | 0.736 | – |
| 346.613 | 0.173 | -0.038 | – | – |
| 115.419 | 0.228 | 0.027 | 0.384 | – |
| 42.316 | 0.138 | 0.069 | 0.736 | – |
| 91.868 | 0.235 | 0.031 | 0.437 | – |
| 42.316 | 0.138 | 0.069 | 0.736 | – |
| 42.316 | 0.138 | 0.069 | 0.736 | – |
| 204.681 | 0.265 | – | 0.234 | – |
| 165.584 | 0.138 | – | 0.108 | – |
| 119.606 | 0.757 | 0.01 | 0.933 | – |
| 42.316 | 0.138 | 0.069 | 0.736 | – |
| 42.316 | 0.138 | 0.069 | 0.736 | – |
| 152.94 | 0.195 | 0.024 | 0.296 | – |
| 71.496 | 0.23 | – | 0.23 | – |
| 42.316 | 0.138 | 0.069 | 0.736 | – |
| 358.317 | 0.154 | -0.03 | – | – |
| 42.316 | 0.138 | 0.069 | 0.736 | – |
| 65.649 | 1.602 | – | 2 | -0.011 |
| 251.683 | 0.264 | – | 0.208 | – |
| 42.316 | 0.138 | 0.069 | 0.736 | – |
| 72.207 | 0.737 | 0.026 | 1.292 | – |
| 309.292 | 0.198 | -0.035 | – | – |
| 72.824 | 0.447 | 0.03 | 0.856 | – |
| 152.94 | 0.195 | 0.024 | 0.296 | – |
| 42.316 | 0.138 | 0.069 | 0.736 | – |
| 165.584 | 0.138 | – | 0.108 | – |
| 165.584 | 0.138 | – | 0.108 | – |
| 98.337 | 0.347 | 0.03 | 0.651 | – |
| 143.132 | 0.067 | – | – | – |
| 42.316 | 0.138 | 0.069 | 0.736 | – |
| 160.617 | 0.343 | – | 0.342 | – |
| 165.584 | 0.138 | – | 0.108 | – |
| 42.316 | 0.138 | 0.069 | 0.736 | – |
| 98.337 | 0.347 | 0.03 | 0.651 | – |
| 143.132 | 0.067 | – | – | – |
| 42.316 | 0.138 | 0.069 | 0.736 | – |
| 308.131 | 0.212 | – | 0.16 | – |
| 212.3 | 0.306 | – | 0.274 | – |
| 42.316 | 0.138 | 0.069 | 0.736 | – |
| 42.316 | 0.138 | 0.069 | 0.736 | – |
| 115.419 | 0.228 | 0.027 | 0.384 | – |
| 167.484 | 0.409 | – | 0.407 | – |
| 72.824 | 0.447 | 0.03 | 0.856 | – |

| | | | | |
|---|---|---|---|---|
| 98.337 | 0.347 | 0.03 | 0.651 | – |
| 42.316 | 0.138 | 0.069 | 0.736 | – |
| 143.132 | 0.067 | – | – | – |
| 42.316 | 0.138 | 0.069 | 0.736 | – |
| 165.584 | 0.138 | – | 0.108 | – |
| 42.316 | 0.138 | 0.069 | 0.736 | – |
| 152.94 | 0.195 | 0.024 | 0.296 | – |
| 143.132 | 0.067 | – | – | – |
| 165.584 | 0.138 | – | 0.108 | – |
| 143.132 | 0.067 | – | – | – |
| 42.316 | 0.138 | 0.069 | 0.736 | – |
| 108.379 | 1.316 | 0.011 | 1.654 | – |
| 208.273 | 0.322 | – | 0.295 | – |
| 42.316 | 0.138 | 0.069 | 0.736 | – |
| 257.549 | 0.207 | – | 0.17 | – |
| 98.337 | 0.347 | 0.03 | 0.651 | – |
| 42.316 | 0.138 | 0.069 | 0.736 | – |
| 71.496 | 0.23 | – | 0.23 | – |
| 129.344 | 0.19 | 0.029 | 0.325 | – |
| 143.132 | 0.067 | – | – | – |
| 42.316 | 0.138 | 0.069 | 0.736 | – |
| 42.316 | 0.138 | 0.069 | 0.736 | – |
| 102.207 | 1.136 | 0.013 | 1.51 | – |
| 42.316 | 0.138 | 0.069 | 0.736 | – |
| 42.316 | 0.138 | 0.069 | 0.736 | – |
| 166.868 | 0.364 | – | 0.363 | – |
| 72.824 | 0.447 | 0.03 | 0.856 | – |
| 42.316 | 0.138 | 0.069 | 0.736 | – |
| 192.668 | 0.604 | – | 0.604 | – |
| 71.496 | 0.23 | – | 0.23 | – |
| 366.988 | 0.166 | -0.034 | – | – |
| 72.824 | 0.447 | 0.03 | 0.856 | – |
| 304.665 | 0.222 | – | 0.173 | – |
| 165.584 | 0.138 | – | 0.108 | – |
| 152.94 | 0.195 | 0.024 | 0.296 | – |
| 143.132 | 0.067 | – | – | – |
| 42.316 | 0.138 | 0.069 | 0.736 | – |
| 165.584 | 0.138 | – | 0.108 | – |
| 165.584 | 0.138 | – | 0.108 | – |
| 71.496 | 0.23 | – | 0.23 | – |
| 165.584 | 0.138 | – | 0.108 | – |
| 152.94 | 0.195 | 0.024 | 0.296 | – |
| 42.316 | 0.138 | 0.069 | 0.736 | – |

| | | | | |
|---|---|---|---|---|
| 101.544 | 1.675 | – | 2 | -0.011 |
| 143.132 | 0.067 | – | – | – |
| 98.337 | 0.347 | 0.03 | 0.651 | – |
| 42.316 | 0.138 | 0.069 | 0.736 | – |
| 42.316 | 0.138 | 0.069 | 0.736 | – |
| 81.096 | 1.514 | 0.007 | 1.759 | – |
| 250.528 | 0.228 | – | 0.19 | – |
| 42.316 | 0.138 | 0.069 | 0.736 | – |
| 145.585 | 1.302 | 0.009 | 1.589 | – |
| 42.316 | 0.138 | 0.069 | 0.736 | – |
| 71.496 | 0.23 | – | 0.23 | – |
| 382.389 | 0.24 | -0.045 | – | – |
| 98.337 | 0.347 | 0.03 | 0.651 | – |
| 91.868 | 0.235 | 0.031 | 0.437 | – |
| 265.248 | 0.177 | -0.032 | – | – |
| 98.337 | 0.347 | 0.03 | 0.651 | – |
| 143.132 | 0.067 | – | – | – |
| 152.94 | 0.195 | 0.024 | 0.296 | – |
| 325.012 | 0.259 | -0.054 | – | – |
| 115.419 | 0.228 | 0.027 | 0.384 | – |
| 152.215 | 1.4 | 0.008 | 1.663 | – |
| 42.316 | 0.138 | 0.069 | 0.736 | – |
| 108.079 | 1.343 | 0.01 | 1.668 | – |
| 71.496 | 0.23 | – | 0.23 | – |
| 71.496 | 0.23 | – | 0.23 | – |
| 71.496 | 0.23 | – | 0.23 | – |
| 98.337 | 0.347 | 0.03 | 0.651 | – |
| 42.316 | 0.138 | 0.069 | 0.736 | – |
| 71.496 | 0.23 | – | 0.23 | – |
| 165.86 | 0.485 | 0.016 | 0.646 | – |
| 91.868 | 0.235 | 0.031 | 0.437 | – |
| 297.568 | 0.237 | – | 0.184 | – |
| 115.419 | 0.228 | 0.027 | 0.384 | – |
| 265.577 | 0.239 | – | 0.203 | – |
| 233.91 | 0.429 | – | 0.41 | – |
| 42.316 | 0.138 | 0.069 | 0.736 | – |
| – | – | – | – | – |
| 264.082 | 0.213 | – | 0.176 | – |
| 279.014 | 0.344 | – | 0.299 | – |
| 71.496 | 0.23 | – | 0.23 | – |
| 98.337 | 0.347 | 0.03 | 0.651 | – |
| 71.496 | 0.23 | – | 0.23 | – |
| 71.496 | 0.23 | – | 0.23 | – |

| | | | | |
|---|---|---|---|---|
| 131.49 | 1.393 | 0.009 | 1.651 | – |
| 421.786 | 0.218 | -0.05 | – | – |
| 71.496 | 0.23 | – | 0.23 | – |
| 42.316 | 0.138 | 0.069 | 0.736 | – |
| 91.868 | 0.235 | 0.031 | 0.437 | – |
| 42.316 | 0.138 | 0.069 | 0.736 | – |
| 129.344 | 0.19 | 0.029 | 0.325 | – |
| 91.868 | 0.235 | 0.031 | 0.437 | – |
| 98.337 | 0.347 | 0.03 | 0.651 | – |
| 219.679 | 0.302 | 0.019 | 0.4 | – |
| 152.94 | 0.195 | 0.024 | 0.296 | – |
| 115.419 | 0.228 | 0.027 | 0.384 | – |
| 42.316 | 0.138 | 0.069 | 0.736 | – |
| 91.868 | 0.235 | 0.031 | 0.437 | – |
| 72.824 | 0.447 | 0.03 | 0.856 | – |
| 91.868 | 0.235 | 0.031 | 0.437 | – |
| 152.94 | 0.195 | 0.024 | 0.296 | – |
| 364.012 | 0.171 | -0.035 | – | – |
| 151.834 | 1.267 | 0.008 | 1.514 | – |
| 241.175 | 0.469 | – | 0.449 | – |
| 115.419 | 0.228 | 0.027 | 0.384 | – |
| 115.419 | 0.228 | 0.027 | 0.384 | – |
| 258.456 | 0.26 | – | 0.222 | – |
| 98.337 | 0.347 | 0.03 | 0.651 | – |
| 71.496 | 0.23 | – | 0.23 | – |
| 152.94 | 0.195 | 0.024 | 0.296 | – |
| 42.316 | 0.138 | 0.069 | 0.736 | – |
| 181.824 | 0.718 | – | 0.718 | – |
| 188.297 | 0.782 | – | 0.782 | – |
| 129.344 | 0.19 | 0.029 | 0.325 | – |
| 91.868 | 0.235 | 0.031 | 0.437 | – |
| 72.824 | 0.447 | 0.03 | 0.856 | – |
| 143.132 | 0.067 | – | – | – |
| 98.337 | 0.347 | 0.03 | 0.651 | – |
| 42.316 | 0.138 | 0.069 | 0.736 | – |
| 71.496 | 0.23 | – | 0.23 | – |
| 273.399 | 0.194 | -0.036 | – | – |
| 152.94 | 0.195 | 0.024 | 0.296 | – |
| 71.496 | 0.23 | – | 0.23 | – |
| 71.496 | 0.23 | – | 0.23 | – |
| 152.94 | 0.195 | 0.024 | 0.296 | – |
| 71.496 | 0.23 | – | 0.23 | – |
| 356.448 | 0.18 | -0.034 | – | – |

| | | | | |
|---|---|---|---|---|
| 91.868 | 0.235 | 0.031 | 0.437 | – |
| 42.316 | 0.138 | 0.069 | 0.736 | – |
| 115.419 | 0.228 | 0.027 | 0.384 | – |
| 286.624 | 0.373 | – | 0.327 | – |
| 207.421 | 0.269 | – | 0.251 | – |
| 123.438 | 1.272 | 0.011 | 1.604 | – |
| 317.951 | 0.178 | – | 0.117 | – |
| 72.824 | 0.447 | 0.03 | 0.856 | – |
| 42.316 | 0.138 | 0.069 | 0.736 | – |
| 71.496 | 0.23 | – | 0.23 | – |
| 240.239 | 0.444 | – | 0.425 | – |
| 165.584 | 0.138 | – | 0.108 | – |
| 91.868 | 0.235 | 0.031 | 0.437 | – |
| 165.584 | 0.138 | – | 0.108 | – |
| 98.337 | 0.347 | 0.03 | 0.651 | – |
| 71.496 | 0.23 | – | 0.23 | – |
| 282.165 | 0.386 | – | 0.342 | – |
| 376.552 | 0.258 | – | 0.202 | – |
| 187.596 | 0.843 | 0.008 | 0.988 | – |
| 98.337 | 0.347 | 0.03 | 0.651 | – |
| 254.678 | 0.273 | – | 0.239 | – |
| 72.824 | 0.447 | 0.03 | 0.856 | – |
| 129.344 | 0.19 | 0.029 | 0.325 | – |
| 165.584 | 0.138 | – | 0.108 | – |
| 152.94 | 0.195 | 0.024 | 0.296 | – |
| 214.762 | 0.316 | – | 0.298 | – |
| 115.419 | 0.228 | 0.027 | 0.384 | – |
| 98.337 | 0.347 | 0.03 | 0.651 | – |
| 157.534 | 0.534 | 0.013 | 0.693 | – |
| 161.307 | 1.156 | 0.006 | 1.312 | – |
| 115.419 | 0.228 | 0.027 | 0.384 | – |
| 115.419 | 0.228 | 0.027 | 0.384 | – |
| 98.337 | 0.347 | 0.03 | 0.651 | – |
| 159.194 | 0.655 | 0.011 | 0.824 | – |
| 165.584 | 0.138 | – | 0.108 | – |
| 311.029 | 0.19 | – | 0.124 | – |
| 129.344 | 0.19 | 0.029 | 0.325 | – |
| 164.252 | 0.555 | – | 0.555 | – |
| 434.754 | 0.198 | -0.043 | – | – |
| 42.316 | 0.138 | 0.069 | 0.736 | – |
| 339.04 | 0.23 | -0.045 | – | – |
| 115.419 | 0.228 | 0.027 | 0.384 | – |
| 270.081 | 0.2 | -0.038 | – | – |

| | | | | |
|---|---|---|---|---|
| 165.584 | 0.138 | – | 0.108 | – |
| 213.753 | 0.321 | 0.015 | 0.399 | – |
| 152.94 | 0.195 | 0.024 | 0.296 | – |
| 223.585 | 0.238 | – | 0.193 | – |
| 72.824 | 0.447 | 0.03 | 0.856 | – |
| 376.362 | 0.168 | -0.033 | – | – |
| 152.94 | 0.195 | 0.024 | 0.296 | – |
| 152.94 | 0.195 | 0.024 | 0.296 | – |
| 152.94 | 0.195 | 0.024 | 0.296 | – |
| 42.316 | 0.138 | 0.069 | 0.736 | – |
| 72.824 | 0.447 | 0.03 | 0.856 | – |
| 71.496 | 0.23 | – | 0.23 | – |
| 143.132 | 0.067 | – | – | – |
| 324.522 | 0.182 | – | 0.12 | – |
| 72.824 | 0.447 | 0.03 | 0.856 | – |
| 152.94 | 0.195 | 0.024 | 0.296 | – |
| 130.1 | 1.021 | 0.011 | 1.284 | – |
| 204.681 | 0.265 | – | 0.234 | – |
| 91.868 | 0.235 | 0.031 | 0.437 | – |
| 278.228 | 0.189 | – | 0.142 | – |
| 119.606 | 0.757 | 0.01 | 0.933 | – |
| 98.337 | 0.347 | 0.03 | 0.651 | – |
| 42.316 | 0.138 | 0.069 | 0.736 | – |
| 72.824 | 0.447 | 0.03 | 0.856 | – |
| 326.459 | 0.2 | – | 0.14 | – |
| 123.964 | 0.862 | 0.014 | 1.148 | – |
| 384.652 | 0.278 | – | 0.222 | – |
| 42.316 | 0.138 | 0.069 | 0.736 | – |
| 98.337 | 0.347 | 0.03 | 0.651 | – |
| 42.316 | 0.138 | 0.069 | 0.736 | – |
| 251.683 | 0.264 | – | 0.208 | – |
| 71.496 | 0.23 | – | 0.23 | – |
| 140.533 | 0.707 | 0.014 | 0.918 | – |
| 98.337 | 0.347 | 0.03 | 0.651 | – |
| 129.344 | 0.19 | 0.029 | 0.325 | – |
| 325.318 | 0.205 | – | 0.146 | – |
| 309.292 | 0.198 | -0.035 | – | – |
| 143.132 | 0.067 | – | – | – |
| 333.445 | 0.292 | – | 0.251 | – |
| 72.824 | 0.447 | 0.03 | 0.856 | – |
| 71.496 | 0.23 | – | 0.23 | – |
| 98.337 | 0.347 | 0.03 | 0.651 | – |
| 143.132 | 0.067 | – | – | – |

| | | | | |
|---|---|---|---|---|
| 96.742 | 1.306 | 0.011 | 1.652 | – |
| 98.337 | 0.347 | 0.03 | 0.651 | – |
| 152.94 | 0.195 | 0.024 | 0.296 | – |
| 143.132 | 0.067 | – | – | – |
| 42.316 | 0.138 | 0.069 | 0.736 | – |
| 42.316 | 0.138 | 0.069 | 0.736 | – |
| 346.821 | 0.243 | -0.048 | – | – |
| 165.584 | 0.138 | – | 0.108 | – |
| 165.584 | 0.138 | – | 0.108 | – |
| 160.617 | 0.343 | – | 0.342 | – |
| 383.223 | 0.167 | -0.033 | – | – |
| 98.337 | 0.347 | 0.03 | 0.651 | – |
| 72.824 | 0.447 | 0.03 | 0.856 | – |
| 231.368 | 0.273 | – | 0.228 | – |
| 325.918 | 0.307 | – | 0.262 | – |
| 71.496 | 0.23 | – | 0.23 | – |
| 331.091 | 0.168 | – | 0.098 | – |
| 98.337 | 0.347 | 0.03 | 0.651 | – |
| 212.3 | 0.306 | – | 0.274 | – |
| 442.981 | 0.209 | -0.046 | – | – |
| 380.643 | 0.287 | – | 0.232 | – |
| 319.436 | 0.212 | – | 0.146 | – |
| 152.94 | 0.195 | 0.024 | 0.296 | – |
| 230.025 | 0.246 | – | 0.2 | – |
| 72.824 | 0.447 | 0.03 | 0.856 | – |
| 42.316 | 0.138 | 0.069 | 0.736 | – |
| 129.344 | 0.19 | 0.029 | 0.325 | – |
| 167.484 | 0.409 | – | 0.407 | – |
| 139.542 | 0.642 | – | 0.642 | – |
| 178.094 | 0.322 | – | 0.309 | – |
| 152.94 | 0.195 | 0.024 | 0.296 | – |
| 143.132 | 0.067 | – | – | – |
| 115.419 | 0.228 | 0.027 | 0.384 | – |
| 72.824 | 0.447 | 0.03 | 0.856 | – |
| 42.316 | 0.138 | 0.069 | 0.736 | – |
| 42.316 | 0.138 | 0.069 | 0.736 | – |
| 152.94 | 0.195 | 0.024 | 0.296 | – |
| 129.344 | 0.19 | 0.029 | 0.325 | – |
| 373.045 | 0.297 | – | 0.236 | – |
| 206.569 | 0.416 | – | 0.4 | – |
| 165.584 | 0.138 | – | 0.108 | – |
| 115.419 | 0.228 | 0.027 | 0.384 | – |
| 342.789 | 0.249 | -0.049 | – | – |

| | | | | |
|---|---|---|---|---|
| 282.451 | 0.196 | -0.034 | – | – |
| 281.639 | 0.364 | – | 0.337 | – |
| 143.132 | 0.067 | – | – | – |
| 152.94 | 0.195 | 0.024 | 0.296 | – |
| 98.337 | 0.347 | 0.03 | 0.651 | – |
| 71.496 | 0.23 | – | 0.23 | – |
| 230.826 | 0.525 | – | 0.514 | – |
| 286.495 | 0.216 | – | 0.17 | – |
| 129.344 | 0.19 | 0.029 | 0.325 | – |
| 71.496 | 0.23 | – | 0.23 | – |
| 185.001 | 0.377 | – | 0.364 | – |
| 241.679 | 0.535 | 0.011 | 0.631 | – |
| 91.868 | 0.235 | 0.031 | 0.437 | – |
| 341.283 | 0.317 | – | 0.276 | – |
| 274.48 | 0.206 | – | 0.163 | – |
| 208.273 | 0.322 | – | 0.295 | – |
| 71.496 | 0.23 | – | 0.23 | – |
| 72.824 | 0.447 | 0.03 | 0.856 | – |
| 138.582 | 0.498 | – | 0.498 | – |
| 42.316 | 0.138 | 0.069 | 0.736 | – |
| 98.337 | 0.347 | 0.03 | 0.651 | – |
| 42.316 | 0.138 | 0.069 | 0.736 | – |
| 98.337 | 0.347 | 0.03 | 0.651 | – |
| 115.419 | 0.228 | 0.027 | 0.384 | – |
| 98.337 | 0.347 | 0.03 | 0.651 | – |
| 431.097 | 0.221 | -0.044 | – | – |
| 225.403 | 0.307 | – | 0.269 | – |
| 232.783 | 0.584 | – | 0.575 | – |
| 42.316 | 0.138 | 0.069 | 0.736 | – |
| 337.919 | 0.147 | -0.028 | – | – |
| 72.824 | 0.447 | 0.03 | 0.856 | – |
| 339.983 | 0.298 | – | 0.257 | – |
| 237.753 | 0.568 | – | 0.556 | – |
| 152.94 | 0.195 | 0.024 | 0.296 | – |
| 166.868 | 0.364 | – | 0.363 | – |
| 91.868 | 0.235 | 0.031 | 0.437 | – |
| 321.437 | 0.219 | – | 0.164 | – |
| 134.013 | 0.628 | 0.011 | 0.782 | – |
| 333.702 | 0.332 | – | 0.287 | – |
| 269.172 | 0.211 | – | 0.163 | – |
| 143.132 | 0.067 | – | – | – |
| 42.316 | 0.138 | 0.069 | 0.736 | – |
| 439.39 | 0.213 | -0.047 | – | – |

| | | | | |
|---|---|---|---|---|
| 71.496 | 0.23 | – | 0.23 | – |
| 192.668 | 0.604 | – | 0.604 | – |
| 152.94 | 0.195 | 0.024 | 0.296 | – |
| 71.496 | 0.23 | – | 0.23 | – |
| 98.337 | 0.347 | 0.03 | 0.651 | – |
| 152.94 | 0.195 | 0.024 | 0.296 | – |
| 143.132 | 0.067 | – | – | – |
| 143.132 | 0.067 | – | – | – |
| 98.337 | 0.347 | 0.03 | 0.651 | – |
| 42.316 | 0.138 | 0.069 | 0.736 | – |
| 72.824 | 0.447 | 0.03 | 0.856 | – |
| 316.008 | 0.221 | – | 0.158 | – |
| 143.132 | 0.067 | – | – | – |
| 98.337 | 0.347 | 0.03 | 0.651 | – |
| 71.496 | 0.23 | – | 0.23 | – |
| 235.931 | 0.21 | – | 0.179 | – |
| 72.824 | 0.447 | 0.03 | 0.856 | – |
| 165.584 | 0.138 | – | 0.108 | – |
| 289.101 | 0.397 | – | 0.369 | – |
| 42.316 | 0.138 | 0.069 | 0.736 | – |
| 152.94 | 0.195 | 0.024 | 0.296 | – |
| 115.419 | 0.228 | 0.027 | 0.384 | – |
| 186.697 | 0.308 | – | 0.292 | – |
| 152.94 | 0.195 | 0.024 | 0.296 | – |
| 329.423 | 0.343 | – | 0.299 | – |
| 145.585 | 1.302 | 0.009 | 1.589 | – |
| 196.11 | 0.342 | 0.02 | 0.47 | – |
| 129.344 | 0.19 | 0.029 | 0.325 | – |
| 382.389 | 0.24 | -0.045 | – | – |
| 282.673 | 0.198 | – | 0.153 | – |
| 115.419 | 0.228 | 0.027 | 0.384 | – |
| 295.815 | 0.352 | – | 0.326 | – |
| 91.868 | 0.235 | 0.031 | 0.437 | – |
| 42.316 | 0.138 | 0.069 | 0.736 | – |
| 71.496 | 0.23 | – | 0.23 | – |
| 129.344 | 0.19 | 0.029 | 0.325 | – |
| 265.248 | 0.177 | -0.032 | – | – |
| 133.862 | 0.849 | 0.012 | 1.091 | – |
| 98.337 | 0.347 | 0.03 | 0.651 | – |
| 325.012 | 0.259 | -0.054 | – | – |
| 71.496 | 0.23 | – | 0.23 | – |
| 152.215 | 1.4 | 0.008 | 1.663 | – |
| 98.337 | 0.347 | 0.03 | 0.651 | – |

| | | | | |
|---|---|---|---|---|
| 91.868 | 0.235 | 0.031 | 0.437 | – |
| 91.868 | 0.235 | 0.031 | 0.437 | – |
| 237.355 | 0.214 | – | 0.153 | – |
| 129.344 | 0.19 | 0.029 | 0.325 | – |
| 152.94 | 0.195 | 0.024 | 0.296 | – |
| 283.054 | 0.227 | – | 0.185 | – |
| 339.743 | 0.187 | – | 0.118 | – |
| 98.337 | 0.347 | 0.03 | 0.651 | – |
| 288.059 | 0.374 | – | 0.347 | – |
| 42.316 | 0.138 | 0.069 | 0.736 | – |
| 152.94 | 0.195 | 0.024 | 0.296 | – |
| 71.496 | 0.23 | – | 0.23 | – |
| 98.337 | 0.347 | 0.03 | 0.651 | – |
| 42.316 | 0.138 | 0.069 | 0.736 | – |
| 282.459 | 0.235 | – | 0.193 | – |
| 129.344 | 0.19 | 0.029 | 0.325 | – |
| 129.344 | 0.19 | 0.029 | 0.325 | – |
| 152.94 | 0.195 | 0.024 | 0.296 | – |
| 233.91 | 0.429 | – | 0.41 | – |
| 228.844 | 0.235 | – | 0.205 | – |
| 91.868 | 0.235 | 0.031 | 0.437 | – |
| 72.824 | 0.447 | 0.03 | 0.856 | – |
| 71.496 | 0.23 | – | 0.23 | – |
| 279.014 | 0.344 | – | 0.299 | – |
| 152.94 | 0.195 | 0.024 | 0.296 | – |
| 298.286 | 0.313 | – | 0.255 | – |
| 42.316 | 0.138 | 0.069 | 0.736 | – |
| 72.824 | 0.447 | 0.03 | 0.856 | – |
| 72.824 | 0.447 | 0.03 | 0.856 | – |
| 184.21 | 0.345 | 0.017 | 0.46 | – |
| 243.708 | 0.247 | – | 0.217 | – |
| 277.235 | 0.239 | – | 0.192 | – |
| 152.94 | 0.195 | 0.024 | 0.296 | – |
| 275.667 | 0.216 | – | 0.168 | – |
| 206.349 | 0.573 | – | 0.573 | – |
| 211.254 | 0.555 | – | 0.554 | – |
| 71.496 | 0.23 | – | 0.23 | – |
| 115.419 | 0.228 | 0.027 | 0.384 | – |
| 72.824 | 0.447 | 0.03 | 0.856 | – |
| 275.962 | 0.245 | – | 0.2 | – |
| 98.337 | 0.347 | 0.03 | 0.651 | – |
| 212.968 | 0.628 | – | 0.628 | – |
| 242.315 | 0.254 | 0.017 | 0.315 | – |

| | | | | |
|---|---|---|---|---|
| 115.419 | 0.228 | 0.027 | 0.384 | – |
| 72.824 | 0.447 | 0.03 | 0.856 | – |
| 182.526 | 0.441 | – | 0.437 | – |
| 152.94 | 0.195 | 0.024 | 0.296 | – |
| 152.94 | 0.195 | 0.024 | 0.296 | – |
| 279.888 | 0.456 | – | 0.421 | – |
| 355.668 | 0.242 | -0.045 | – | – |
| 71.496 | 0.23 | – | 0.23 | – |
| 129.344 | 0.19 | 0.029 | 0.325 | – |
| 151.834 | 1.267 | 0.008 | 1.514 | – |
| 241.175 | 0.469 | – | 0.449 | – |
| 251.723 | 0.403 | – | 0.374 | – |
| 72.824 | 0.447 | 0.03 | 0.856 | – |
| 346.65 | 0.16 | -0.032 | – | – |
| 71.496 | 0.23 | – | 0.23 | – |
| 71.496 | 0.23 | – | 0.23 | – |
| 181.824 | 0.718 | – | 0.718 | – |
| 188.297 | 0.782 | – | 0.782 | – |
| 152.94 | 0.195 | 0.024 | 0.296 | – |
| 98.337 | 0.347 | 0.03 | 0.651 | – |
| 91.868 | 0.235 | 0.031 | 0.437 | – |
| 42.316 | 0.138 | 0.069 | 0.736 | – |
| 71.496 | 0.23 | – | 0.23 | – |
| 289.597 | 0.18 | – | 0.126 | – |
| 165.584 | 0.138 | – | 0.108 | – |
| 91.868 | 0.235 | 0.031 | 0.437 | – |
| 191.339 | 0.415 | 0.015 | 0.527 | – |
| 91.868 | 0.235 | 0.031 | 0.437 | – |
| 72.824 | 0.447 | 0.03 | 0.856 | – |
| 152.94 | 0.195 | 0.024 | 0.296 | – |
| 245.355 | 0.243 | – | 0.183 | – |
| 273.399 | 0.194 | -0.036 | – | – |
| 395.152 | 0.243 | – | 0.179 | – |
| 236.517 | 0.273 | – | 0.244 | – |
| 286.624 | 0.373 | – | 0.327 | – |
| 71.496 | 0.23 | – | 0.23 | – |
| 123.438 | 1.272 | 0.011 | 1.604 | – |
| 452.108 | 0.209 | -0.044 | – | – |
| 387.284 | 0.209 | -0.044 | – | – |
| 143.132 | 0.067 | – | – | – |
| 336.559 | 0.195 | – | 0.129 | – |
| 237.404 | 0.283 | 0.018 | 0.361 | – |
| 305.983 | 0.339 | – | 0.281 | – |

| | | | | |
|---|---|---|---|---|
| 143.132 | 0.067 | – | – | – |
| 224.579 | 0.261 | – | 0.235 | – |
| 91.868 | 0.235 | 0.031 | 0.437 | – |
| 240.239 | 0.444 | – | 0.425 | – |
| 129.344 | 0.19 | 0.029 | 0.325 | – |
| 184.478 | 0.421 | 0.016 | 0.554 | – |
| 190.233 | 0.368 | 0.016 | 0.481 | – |
| 115.419 | 0.228 | 0.027 | 0.384 | – |
| 258.959 | 0.439 | – | 0.409 | – |
| 129.344 | 0.19 | 0.029 | 0.325 | – |
| 129.344 | 0.19 | 0.029 | 0.325 | – |
| 129.344 | 0.19 | 0.029 | 0.325 | – |
| 282.165 | 0.386 | – | 0.342 | – |
| 72.824 | 0.447 | 0.03 | 0.856 | – |
| 129.344 | 0.19 | 0.029 | 0.325 | – |
| 179.917 | 0.448 | 0.016 | 0.596 | – |
| 152.94 | 0.195 | 0.024 | 0.296 | – |
| 304.772 | 0.32 | – | 0.263 | – |
| 187.596 | 0.843 | 0.008 | 0.988 | – |
| 186.44 | 0.532 | 0.014 | 0.675 | – |
| 143.132 | 0.067 | – | – | – |
| 271.656 | 0.266 | – | 0.224 | – |
| 210.62 | 0.841 | 0.01 | 1.007 | – |
| 299.483 | 0.367 | – | 0.313 | – |
| 72.824 | 0.447 | 0.03 | 0.856 | – |
| 72.824 | 0.447 | 0.03 | 0.856 | – |
| 72.824 | 0.447 | 0.03 | 0.856 | – |
| 282.731 | 0.194 | – | 0.136 | – |
| 194.924 | 0.254 | – | 0.223 | – |
| 143.132 | 0.067 | – | – | – |
| 152.94 | 0.195 | 0.024 | 0.296 | – |
| 296.556 | 0.178 | – | 0.123 | – |
| 157.534 | 0.534 | 0.013 | 0.693 | – |
| 161.307 | 1.156 | 0.006 | 1.312 | – |
| 233.389 | 0.3 | 0.018 | 0.39 | – |
| 329.463 | 0.205 | – | 0.132 | – |
| 232.727 | 0.289 | – | 0.263 | – |
| 159.194 | 0.655 | 0.011 | 0.824 | – |
| 115.419 | 0.228 | 0.027 | 0.384 | – |
| 231.923 | 0.302 | – | 0.277 | – |
| 216.719 | 0.643 | 0.011 | 0.776 | – |
| 241.77 | 0.256 | – | 0.2 | – |
| 152.94 | 0.195 | 0.024 | 0.296 | – |

| | | | | |
|---|---|---|---|---|
| 260.433 | 0.393 | – | 0.362 | – |
| 72.824 | 0.447 | 0.03 | 0.856 | – |
| 193.505 | 0.595 | 0.016 | 0.777 | – |
| 164.252 | 0.555 | – | 0.555 | – |
| 98.337 | 0.347 | 0.03 | 0.651 | – |
| 72.824 | 0.447 | 0.03 | 0.856 | – |
| 143.132 | 0.067 | – | – | – |
| 343.514 | 0.193 | – | 0.126 | – |
| 459.122 | 0.207 | -0.044 | – | – |
| 72.824 | 0.447 | 0.03 | 0.856 | – |
| 355.064 | 0.267 | – | 0.218 | – |
| 129.344 | 0.19 | 0.029 | 0.325 | – |
| 296.104 | 0.184 | – | 0.129 | – |
| 339.04 | 0.23 | -0.045 | – | – |
| 270.081 | 0.2 | -0.038 | – | – |
| 233.006 | 0.25 | – | 0.222 | – |
| 98.337 | 0.347 | 0.03 | 0.651 | – |
| 223.585 | 0.238 | – | 0.193 | – |
| 297.954 | 0.204 | – | 0.151 | – |
| 403.322 | 0.261 | – | 0.197 | – |
| 401.744 | 0.247 | – | 0.183 | – |
| 200.827 | 0.656 | – | 0.656 | – |
| 207.3 | 0.713 | – | 0.713 | – |
| 395.378 | 0.22 | -0.046 | – | – |
| 98.337 | 0.347 | 0.03 | 0.651 | – |
| 401.998 | 0.265 | – | 0.203 | – |
| 72.824 | 0.447 | 0.03 | 0.856 | – |
| 130.1 | 1.021 | 0.011 | 1.284 | – |
| 72.824 | 0.447 | 0.03 | 0.856 | – |
| 129.344 | 0.19 | 0.029 | 0.325 | – |
| 240.73 | 0.349 | 0.016 | 0.437 | – |
| 202.378 | 0.296 | – | 0.266 | – |
| 71.496 | 0.23 | – | 0.23 | – |
| 72.824 | 0.447 | 0.03 | 0.856 | – |
| 202.712 | 0.784 | – | 0.784 | – |
| 98.337 | 0.347 | 0.03 | 0.651 | – |
| 71.496 | 0.23 | – | 0.23 | – |
| 182.854 | 1.138 | 0.009 | 1.343 | – |
| 129.344 | 0.19 | 0.029 | 0.325 | – |
| 72.824 | 0.447 | 0.03 | 0.856 | – |
| 71.496 | 0.23 | – | 0.23 | – |
| 91.868 | 0.235 | 0.031 | 0.437 | – |
| 255.856 | 0.483 | – | 0.459 | – |

| | | | | |
|---|---|---|---|---|
| 129.344 | 0.19 | 0.029 | 0.325 | – |
| 289.697 | 0.192 | – | 0.133 | – |
| 123.964 | 0.862 | 0.014 | 1.148 | – |
| 336.378 | 0.174 | -0.033 | – | – |
| 98.337 | 0.347 | 0.03 | 0.651 | – |
| 91.868 | 0.235 | 0.031 | 0.437 | – |
| 129.344 | 0.19 | 0.029 | 0.325 | – |
| 201.241 | 0.264 | – | 0.232 | – |
| 98.337 | 0.347 | 0.03 | 0.651 | – |
| 408.007 | 0.196 | -0.043 | – | – |
| 71.496 | 0.23 | – | 0.23 | – |
| 96.742 | 1.306 | 0.011 | 1.652 | – |
| 177.689 | 0.53 | – | 0.53 | – |
| 143.132 | 0.067 | – | – | – |
| 350.602 | 0.284 | – | 0.237 | – |
| 152.94 | 0.195 | 0.024 | 0.296 | – |
| 143.132 | 0.067 | – | – | – |
| 303.067 | 0.182 | – | 0.126 | – |
| 290.982 | 0.219 | – | 0.162 | – |
| 281.636 | 0.265 | – | 0.229 | – |
| 98.337 | 0.347 | 0.03 | 0.651 | – |
| 223.453 | 0.31 | – | 0.278 | – |
| 346.821 | 0.243 | -0.048 | – | – |
| 312.37 | 0.23 | -0.045 | – | – |
| 344.982 | 0.284 | – | 0.231 | – |
| 91.868 | 0.235 | 0.031 | 0.437 | – |
| 391.63 | 0.226 | -0.047 | – | – |
| 72.824 | 0.447 | 0.03 | 0.856 | – |
| 71.496 | 0.23 | – | 0.23 | – |
| 129.344 | 0.19 | 0.029 | 0.325 | – |
| 231.368 | 0.273 | – | 0.228 | – |
| 363.039 | 0.29 | – | 0.24 | – |
| 304.936 | 0.202 | – | 0.148 | – |
| 178.668 | 0.407 | – | 0.404 | – |
| 311.634 | 0.307 | – | 0.273 | – |
| 185.19 | 0.468 | – | 0.464 | – |
| 397.57 | 0.279 | – | 0.219 | – |
| 115.419 | 0.228 | 0.027 | 0.384 | – |
| 152.94 | 0.195 | 0.024 | 0.296 | – |
| 270.408 | 0.296 | – | 0.239 | – |
| 98.337 | 0.347 | 0.03 | 0.651 | – |
| 129.344 | 0.19 | 0.029 | 0.325 | – |
| 230.025 | 0.246 | – | 0.2 | – |

| | | | | |
|---|---|---|---|---|
| 129.344 | 0.19 | 0.029 | 0.325 | – |
| 72.824 | 0.447 | 0.03 | 0.856 | – |
| 71.496 | 0.23 | – | 0.23 | – |
| 139.542 | 0.642 | – | 0.642 | – |
| 178.094 | 0.322 | – | 0.309 | – |
| 143.132 | 0.067 | – | – | – |
| 152.94 | 0.195 | 0.024 | 0.296 | – |
| 304.061 | 0.326 | – | 0.289 | – |
| 157.262 | 0.352 | – | 0.352 | – |
| 129.344 | 0.19 | 0.029 | 0.325 | – |
| 206.569 | 0.416 | – | 0.4 | – |
| 230.678 | 0.349 | – | 0.317 | – |
| 129.344 | 0.19 | 0.029 | 0.325 | – |
| 259.242 | 0.397 | – | 0.379 | – |
| 91.868 | 0.235 | 0.031 | 0.437 | – |
| 342.789 | 0.249 | -0.049 | – | – |
| 152.94 | 0.195 | 0.024 | 0.296 | – |
| 209.815 | 0.648 | – | 0.648 | – |
| 282.451 | 0.196 | -0.034 | – | – |
| 42.316 | 0.138 | 0.069 | 0.736 | – |
| 152.94 | 0.195 | 0.024 | 0.296 | – |
| 230.826 | 0.525 | – | 0.514 | – |
| 356.09 | 0.163 | -0.031 | – | – |
| 91.868 | 0.235 | 0.031 | 0.437 | – |
| 98.337 | 0.347 | 0.03 | 0.651 | – |
| 358.996 | 0.299 | – | 0.252 | – |
| 185.001 | 0.377 | – | 0.364 | – |
| 240.614 | 0.217 | – | 0.179 | – |
| 98.337 | 0.347 | 0.03 | 0.651 | – |
| 414.962 | 0.195 | -0.043 | – | – |
| 98.337 | 0.347 | 0.03 | 0.651 | – |
| 129.344 | 0.19 | 0.029 | 0.325 | – |
| 98.337 | 0.347 | 0.03 | 0.651 | – |
| 167.106 | 0.923 | 0.013 | 1.15 | – |
| 309.715 | 0.167 | – | 0.103 | – |
| 138.582 | 0.498 | – | 0.498 | – |
| 160.727 | 0.717 | 0.013 | 0.906 | – |
| 129.344 | 0.19 | 0.029 | 0.325 | – |
| 272.51 | 0.443 | 0.013 | 0.53 | – |
| 358.407 | 0.307 | – | 0.26 | – |
| 359.045 | 0.277 | – | 0.229 | – |
| 297.971 | 0.217 | – | 0.158 | – |
| 225.403 | 0.307 | – | 0.269 | – |

| | | | | |
|---|---|---|---|---|
| 152.94 | 0.195 | 0.024 | 0.296 | – |
| 232.783 | 0.584 | – | 0.575 | – |
| 320.1 | 0.245 | -0.048 | – | – |
| 152.94 | 0.195 | 0.024 | 0.296 | – |
| 287.554 | 0.229 | – | 0.175 | – |
| 184.718 | 0.43 | – | 0.427 | – |
| 237.753 | 0.568 | – | 0.556 | – |
| 351.417 | 0.312 | – | 0.26 | – |
| 319.301 | 0.336 | – | 0.302 | – |
| 299.159 | 0.348 | – | 0.314 | – |
| 129.344 | 0.19 | 0.029 | 0.325 | – |
| 352.85 | 0.307 | – | 0.254 | – |
| 42.316 | 0.138 | 0.069 | 0.736 | – |
| 226.531 | 0.365 | – | 0.336 | – |
| 134.013 | 0.628 | 0.011 | 0.782 | – |
| 230.217 | 0.347 | – | 0.328 | – |
| 416.228 | 0.207 | -0.045 | – | – |
| 129.344 | 0.19 | 0.029 | 0.325 | – |
| 266.475 | 0.436 | – | 0.417 | – |
| 254.063 | 0.202 | – | 0.164 | – |
| 351.504 | 0.29 | – | 0.237 | – |
| 98.337 | 0.347 | 0.03 | 0.651 | – |
| 143.132 | 0.067 | – | – | – |
| 71.496 | 0.23 | – | 0.23 | – |
| 152.94 | 0.195 | 0.024 | 0.296 | – |
| 252.853 | 0.209 | – | 0.172 | – |
| 311.678 | 0.354 | – | 0.317 | – |
| 71.496 | 0.23 | – | 0.23 | – |
| 318.11 | 0.315 | – | 0.281 | – |
| 159.776 | 0.547 | 0.016 | 0.722 | – |
| 98.337 | 0.347 | 0.03 | 0.651 | – |
| 98.337 | 0.347 | 0.03 | 0.651 | – |
| 129.344 | 0.19 | 0.029 | 0.325 | – |
| 155.001 | 0.588 | 0.016 | 0.781 | – |
| 152.94 | 0.195 | 0.024 | 0.296 | – |
| 152.94 | 0.195 | 0.024 | 0.296 | – |
| 91.868 | 0.235 | 0.031 | 0.437 | – |
| 72.824 | 0.447 | 0.03 | 0.856 | – |
| 186.697 | 0.308 | – | 0.292 | – |
| 269.46 | 0.339 | – | 0.294 | – |
| 129.344 | 0.19 | 0.029 | 0.325 | – |
| 316.074 | 0.252 | -0.049 | – | – |
| 91.868 | 0.235 | 0.031 | 0.437 | – |

| | | | | |
|---|---|---|---|---|
| 129.344 | 0.19 | 0.029 | 0.325 | – |
| 299.855 | 0.224 | – | 0.175 | – |
| 133.862 | 0.849 | 0.012 | 1.091 | – |
| 265.565 | 0.411 | – | 0.393 | – |
| 346.626 | 0.33 | – | 0.281 | – |
| 307.338 | 0.367 | – | 0.332 | – |
| 247.63 | 0.215 | – | 0.175 | – |
| 72.824 | 0.447 | 0.03 | 0.856 | – |
| 237.355 | 0.214 | – | 0.153 | – |
| 343.034 | 0.304 | – | 0.159 | 0.017 |
| 248.432 | 0.251 | – | 0.213 | – |
| 129.344 | 0.19 | 0.029 | 0.325 | – |
| 306.597 | 0.378 | – | 0.344 | – |
| 98.337 | 0.347 | 0.03 | 0.651 | – |
| 72.824 | 0.447 | 0.03 | 0.856 | – |
| 71.496 | 0.23 | – | 0.23 | – |
| 251.616 | 0.463 | – | 0.441 | – |
| 260.138 | 0.445 | – | 0.429 | – |
| 247.017 | 0.224 | – | 0.185 | – |
| 152.94 | 0.195 | 0.024 | 0.296 | – |
| 129.344 | 0.19 | 0.029 | 0.325 | – |
| 404.391 | 0.221 | -0.044 | – | – |
| 152.94 | 0.195 | 0.024 | 0.296 | – |
| 251.087 | 0.543 | – | 0.524 | – |
| 294.557 | 0.227 | – | 0.172 | – |
| 313.27 | 0.337 | – | 0.306 | – |
| 255.518 | 0.46 | – | 0.447 | – |
| 72.824 | 0.447 | 0.03 | 0.856 | – |
| 423.214 | 0.205 | -0.045 | – | – |
| 412.678 | 0.212 | -0.047 | – | – |
| 152.94 | 0.195 | 0.024 | 0.296 | – |
| 261.073 | 0.2 | – | 0.161 | – |
| 298.286 | 0.313 | – | 0.255 | – |
| 307.771 | 0.34 | – | 0.304 | – |
| 98.337 | 0.347 | 0.03 | 0.651 | – |
| 297.073 | 0.372 | – | 0.324 | – |
| 177.26 | 0.892 | 0.008 | 1.062 | – |
| 72.824 | 0.447 | 0.03 | 0.856 | – |
| 260.601 | 0.242 | – | 0.206 | – |
| 258.642 | 0.501 | – | 0.478 | – |
| 214.028 | 0.312 | 0.019 | 0.418 | – |
| 152.94 | 0.195 | 0.024 | 0.296 | – |
| 91.868 | 0.235 | 0.031 | 0.437 | – |

| | | | | |
|---|---|---|---|---|
| 268.711 | 0.556 | 0.013 | 0.673 | – |
| 276.971 | 0.369 | – | 0.324 | – |
| 98.337 | 0.347 | 0.03 | 0.651 | – |
| 262.373 | 0.503 | – | 0.489 | – |
| 206.349 | 0.573 | – | 0.573 | – |
| 211.254 | 0.555 | – | 0.554 | – |
| 129.344 | 0.19 | 0.029 | 0.325 | – |
| 152.94 | 0.195 | 0.024 | 0.296 | – |
| 318.267 | 0.188 | – | 0.125 | – |
| 309.017 | 0.349 | – | 0.32 | – |
| 72.824 | 0.447 | 0.03 | 0.856 | – |
| 212.968 | 0.628 | – | 0.628 | – |
| 182.526 | 0.441 | – | 0.437 | – |
| 279.888 | 0.456 | – | 0.421 | – |
| 355.668 | 0.242 | -0.045 | – | – |
| 230.701 | 0.443 | – | 0.426 | – |
| 72.824 | 0.447 | 0.03 | 0.856 | – |
| 251.723 | 0.403 | – | 0.374 | – |
| 366.791 | 0.252 | – | 0.195 | – |
| 143.132 | 0.067 | – | – | – |
| 147.369 | 0.538 | 0.014 | 0.726 | – |
| 275.857 | 0.348 | – | 0.304 | – |
| 71.496 | 0.23 | – | 0.23 | – |
| 129.344 | 0.19 | 0.029 | 0.325 | – |
| 98.337 | 0.347 | 0.03 | 0.651 | – |
| 255.477 | 0.248 | – | 0.209 | – |
| 153.96 | 0.64 | 0.011 | 0.809 | – |
| 152.94 | 0.195 | 0.024 | 0.296 | – |
| 91.868 | 0.235 | 0.031 | 0.437 | – |
| 209.92 | 0.333 | 0.019 | 0.452 | – |
| 98.337 | 0.347 | 0.03 | 0.651 | – |
| 257.819 | 0.479 | – | 0.457 | – |
| 245.355 | 0.243 | – | 0.183 | – |
| 254.042 | 0.221 | – | 0.181 | – |
| 304.436 | 0.4 | – | 0.351 | – |
| 129.344 | 0.19 | 0.029 | 0.325 | – |
| 72.824 | 0.447 | 0.03 | 0.856 | – |
| 152.94 | 0.195 | 0.024 | 0.296 | – |
| 411.476 | 0.218 | -0.044 | – | – |
| 202.158 | 0.317 | 0.016 | 0.407 | – |
| 359.311 | 0.263 | – | 0.2 | – |
| 316.323 | 0.38 | – | 0.351 | – |
| 305.983 | 0.339 | – | 0.281 | – |

| | | | | |
|---|---|---|---|---|
| 419.688 | 0.21 | -0.047 | – | – |
| 203.69 | 0.305 | 0.016 | 0.391 | – |
| 356.543 | 0.333 | – | 0.29 | – |
| 299.936 | 0.412 | – | 0.365 | – |
| 216.826 | 0.398 | 0.017 | 0.516 | – |
| 152.94 | 0.195 | 0.024 | 0.296 | – |
| 258.959 | 0.439 | – | 0.409 | – |
| 72.824 | 0.447 | 0.03 | 0.856 | – |
| 268.073 | 0.179 | – | 0.131 | – |
| 304.772 | 0.32 | – | 0.263 | – |
| 72.824 | 0.447 | 0.03 | 0.856 | – |
| 72.824 | 0.447 | 0.03 | 0.856 | – |
| 72.824 | 0.447 | 0.03 | 0.856 | – |
| 153.266 | 0.585 | 0.012 | 0.756 | – |
| 304.012 | 0.413 | – | 0.383 | – |
| 299.483 | 0.367 | – | 0.313 | – |
| 209.355 | 0.375 | 0.014 | 0.462 | – |
| 373.81 | 0.25 | – | 0.193 | – |
| 194.924 | 0.254 | – | 0.223 | – |
| 315.127 | 0.197 | – | 0.137 | – |
| 152.94 | 0.195 | 0.024 | 0.296 | – |
| 249.757 | 0.279 | – | 0.246 | – |
| 241.77 | 0.256 | – | 0.2 | – |
| 374.836 | 0.272 | – | 0.216 | – |
| 260.433 | 0.393 | – | 0.362 | – |
| 72.824 | 0.447 | 0.03 | 0.856 | – |
| 373.333 | 0.256 | – | 0.2 | – |
| 129.344 | 0.19 | 0.029 | 0.325 | – |
| 72.824 | 0.447 | 0.03 | 0.856 | – |
| 72.824 | 0.447 | 0.03 | 0.856 | – |
| 366.336 | 0.261 | – | 0.198 | – |
| 194.196 | 0.343 | – | 0.326 | – |
| 71.496 | 0.23 | – | 0.23 | – |
| 367.295 | 0.283 | – | 0.22 | – |
| 200.827 | 0.656 | – | 0.656 | – |
| 207.3 | 0.713 | – | 0.713 | – |
| 261.241 | 0.196 | – | 0.146 | – |
| 210.882 | 0.305 | 0.016 | 0.384 | – |
| 129.344 | 0.19 | 0.029 | 0.325 | – |
| 91.868 | 0.235 | 0.031 | 0.437 | – |
| 201.038 | 0.393 | – | 0.375 | – |
| 202.378 | 0.296 | – | 0.266 | – |
| 202.712 | 0.784 | – | 0.784 | – |

| | | | | |
|---|---|---|---|---|
| 308.113 | 0.21 | – | 0.145 | – |
| 72.824 | 0.447 | 0.03 | 0.856 | – |
| 316.283 | 0.301 | – | 0.256 | – |
| 129.344 | 0.19 | 0.029 | 0.325 | – |
| 274.505 | 0.183 | – | 0.135 | – |
| 255.856 | 0.483 | – | 0.459 | – |
| 241.508 | 0.322 | – | 0.278 | – |
| 276.267 | 0.206 | – | 0.16 | – |
| 129.344 | 0.19 | 0.029 | 0.325 | – |
| 129.344 | 0.19 | 0.029 | 0.325 | – |
| 201.241 | 0.264 | – | 0.232 | – |
| 196.813 | 0.412 | – | 0.398 | – |
| 72.824 | 0.447 | 0.03 | 0.856 | – |
| 381.867 | 0.271 | – | 0.214 | – |
| 363.312 | 0.292 | – | 0.231 | – |
| 177.689 | 0.53 | – | 0.53 | – |
| 380.356 | 0.255 | – | 0.198 | – |
| 432.364 | 0.206 | -0.044 | – | – |
| 72.824 | 0.447 | 0.03 | 0.856 | – |
| 223.453 | 0.31 | – | 0.278 | – |
| 129.344 | 0.19 | 0.029 | 0.325 | – |
| 312.37 | 0.23 | -0.045 | – | – |
| 129.344 | 0.19 | 0.029 | 0.325 | – |
| 206.514 | 0.445 | 0.014 | 0.556 | – |
| 243.429 | 0.663 | 0.013 | 0.81 | – |
| 178.668 | 0.407 | – | 0.404 | – |
| 221.288 | 0.529 | – | 0.52 | – |
| 374.331 | 0.281 | – | 0.218 | – |
| 185.19 | 0.468 | – | 0.464 | – |
| 259.635 | 0.314 | 0.015 | 0.376 | – |
| 270.408 | 0.296 | – | 0.239 | – |
| 328.871 | 0.297 | – | 0.258 | – |
| 129.344 | 0.19 | 0.029 | 0.325 | – |
| 330.302 | 0.292 | – | 0.251 | – |
| 228.049 | 0.575 | – | 0.565 | – |
| 72.824 | 0.447 | 0.03 | 0.856 | – |
| 230.934 | 0.596 | 0.011 | 0.715 | – |
| 269.322 | 0.224 | – | 0.175 | – |
| 323.988 | 0.327 | – | 0.282 | – |
| 235.9 | 0.576 | 0.011 | 0.685 | – |
| 157.262 | 0.352 | – | 0.352 | – |
| 72.824 | 0.447 | 0.03 | 0.856 | – |
| 323.34 | 0.299 | – | 0.254 | – |

| | | | | |
|---|---|---|---|---|
| 129.344 | 0.19 | 0.029 | 0.325 | – |
| 129.344 | 0.19 | 0.029 | 0.325 | – |
| 230.678 | 0.349 | – | 0.317 | – |
| 209.815 | 0.648 | – | 0.648 | – |
| 237.792 | 0.652 | 0.01 | 0.765 | – |
| 322.738 | 0.308 | – | 0.264 | – |
| 376.001 | 0.291 | – | 0.238 | – |
| 98.337 | 0.347 | 0.03 | 0.651 | – |
| 370.356 | 0.29 | – | 0.229 | – |
| 320.1 | 0.245 | -0.048 | – | – |
| 129.344 | 0.19 | 0.029 | 0.325 | – |
| 129.344 | 0.19 | 0.029 | 0.325 | – |
| 184.718 | 0.43 | – | 0.427 | – |
| 227.334 | 0.551 | – | 0.543 | – |
| 226.531 | 0.365 | – | 0.336 | – |
| 276.911 | 0.376 | – | 0.351 | – |
| 129.344 | 0.19 | 0.029 | 0.325 | – |
| 218.921 | 0.222 | – | 0.192 | – |
| 336.513 | 0.324 | – | 0.284 | – |
| 267.719 | 0.409 | – | 0.374 | – |
| 278.486 | 0.369 | – | 0.343 | – |
| 129.344 | 0.19 | 0.029 | 0.325 | – |
| 337.359 | 0.29 | – | 0.248 | – |
| 232.012 | 0.238 | 0.019 | 0.302 | – |
| 387.757 | 0.234 | – | 0.168 | – |
| 98.337 | 0.347 | 0.03 | 0.651 | – |
| 331.056 | 0.325 | – | 0.28 | – |
| 129.344 | 0.19 | 0.029 | 0.325 | – |
| 265.917 | 0.236 | – | 0.191 | – |
| 269.46 | 0.339 | – | 0.294 | – |
| 316.074 | 0.252 | -0.049 | – | – |
| 152.94 | 0.195 | 0.024 | 0.296 | – |
| 314.911 | 0.328 | – | 0.196 | 0.013 |
| 284.134 | 0.411 | – | 0.386 | – |
| 329.801 | 0.306 | – | 0.261 | – |
| 274.881 | 0.442 | – | 0.406 | – |
| 343.034 | 0.304 | – | 0.159 | 0.017 |
| 290.74 | 0.411 | 0.012 | 0.479 | – |
| 129.344 | 0.19 | 0.029 | 0.325 | – |
| 251.616 | 0.463 | – | 0.441 | – |
| 270.291 | 0.456 | – | 0.423 | – |
| 251.087 | 0.543 | – | 0.524 | – |
| 129.344 | 0.19 | 0.029 | 0.325 | – |

| | | | | |
|---|---|---|---|---|
| 324.662 | 0.351 | – | 0.31 | – |
| 226.447 | 0.262 | – | 0.232 | – |
| 286.202 | 0.426 | 0.012 | 0.502 | – |
| 152.94 | 0.195 | 0.024 | 0.296 | – |
| 297.073 | 0.372 | – | 0.324 | – |
| 180.144 | 0.349 | 0.018 | 0.469 | – |
| 177.26 | 0.892 | 0.008 | 1.062 | – |
| 129.344 | 0.19 | 0.029 | 0.325 | – |
| 225.211 | 0.23 | – | 0.199 | – |
| 285.588 | 0.367 | – | 0.34 | – |
| 258.642 | 0.501 | – | 0.478 | – |
| 276.971 | 0.369 | – | 0.324 | – |
| 230.701 | 0.443 | – | 0.426 | – |
| 395.929 | 0.252 | – | 0.187 | – |
| 293.472 | 0.465 | 0.011 | 0.538 | – |
| 147.369 | 0.538 | 0.014 | 0.726 | – |
| 275.857 | 0.348 | – | 0.304 | – |
| 153.96 | 0.64 | 0.011 | 0.809 | – |
| 129.344 | 0.19 | 0.029 | 0.325 | – |
| 257.819 | 0.479 | – | 0.457 | – |
| 304.436 | 0.4 | – | 0.351 | – |
| 299.936 | 0.412 | – | 0.365 | – |
| 281.289 | 0.454 | – | 0.432 | – |
| 345.307 | 0.261 | – | 0.211 | – |
| 334.49 | 0.354 | – | 0.318 | – |
| 72.824 | 0.447 | 0.03 | 0.856 | – |
| 153.266 | 0.585 | 0.012 | 0.756 | – |
| 392.159 | 0.26 | – | 0.197 | – |
| 337.852 | 0.274 | – | 0.219 | – |
| 384.567 | 0.269 | – | 0.198 | – |
| 72.824 | 0.447 | 0.03 | 0.856 | – |
| 194.196 | 0.343 | – | 0.326 | – |
| 201.038 | 0.393 | – | 0.375 | – |
| 353.217 | 0.284 | – | 0.234 | – |
| 351.795 | 0.266 | – | 0.216 | – |
| 241.508 | 0.322 | – | 0.278 | – |
| 345.702 | 0.297 | – | 0.242 | – |
| 196.813 | 0.412 | – | 0.398 | – |
| 294.432 | 0.32 | – | 0.283 | – |
| 221.288 | 0.529 | – | 0.52 | – |
| 341.682 | 0.307 | – | 0.255 | – |
| 228.049 | 0.575 | – | 0.565 | – |
| 301.952 | 0.35 | – | 0.313 | – |

| | | | | |
|---|---|---|---|---|
| 129.344 | 0.19 | 0.029 | 0.325 | – |
| 308.433 | 0.308 | – | 0.274 | – |
| 300.811 | 0.328 | – | 0.292 | – |
| 227.334 | 0.551 | – | 0.543 | – |
| 255.985 | 0.407 | – | 0.389 | – |
| 129.344 | 0.19 | 0.029 | 0.325 | – |
| 267.719 | 0.409 | – | 0.374 | – |
| 314.911 | 0.328 | – | 0.196 | 0.013 |
| 274.881 | 0.442 | – | 0.406 | – |
| 270.291 | 0.456 | – | 0.423 | – |

**Table S4:** A) Global comparison of combinations of diversification shifts for Cycadales (shifts are tested at the crown) with B) the rates of the diversification model for their backbones. The best combination of shift and the phylogeny analysed with no shift are highlighted in bold. NP = Number of parameters, logL = log(Likelihood), λ = speciation rate (at present if variable), α = dependency parameter of speciation rate, μ = extinction rate, β = dependency parameter of extinction rate

| A) Total | | | | | B) Backbone | | | | | | | |
|---|---|---|---|---|---|---|---|---|---|---|---|---|
| **Combination** | **NP** | **logL** | **AICc** | **ΔAICc** | **Model** | **NP** | **logL** | **AICc** | **λ** | **α** | **μ** | **β** |
| **233.309.333.372.399/** | **15** | **-615.069** | **1264.914** | **0** | **BCST DVAR** | **3** | **-80.582** | **169.564** | **0.257** | **–** | **0.294** | **-0.001** |
| 233.333.372.399/ | 13 | -624.182 | 1276.874 | 11.96 | BVAR DCST | 3 | -158.543 | 323.791 | 0.219 | 0.001 | 0.243 | – |
| 233.309.333.399/ | 13 | -625.542 | 1279.605 | 14.691 | BVAR DCST | 3 | -148.438 | 303.542 | 0.416 | 0.001 | 0.445 | – |
| 233.309.372.399/ | 12 | -631.37 | 1288.92 | 24.005 | BVAR DCST | 3 | -199.835 | 406.16 | 0.342 | 0 | 0.363 | – |
| 233.333.399/ | 11 | -634.871 | 1293.424 | 28.51 | BVAR DCST | 3 | -226.614 | 459.628 | 0.312 | 0 | 0.331 | – |
| 309.333.372.399/296 | 16 | -628.944 | 1296.708 | 31.793 | BCST DVAR | 4 | -73.326 | 159.096 | 0.209 | – | 0.291 | -0.005 |
| 233.372.399/ | 9 | -640.201 | 1299.684 | 34.769 | BCST DCST | 2 | -277.514 | 559.191 | 0.276 | – | 0.276 | – |
| 309.333.372.399/ | 12 | -637.246 | 1300.831 | 35.917 | BCST DCST | 2 | -267.364 | 538.887 | 0.33 | – | 0.33 | – |
| 233.309.399/ | 9 | -643.131 | 1305.587 | 40.673 | BCST DCST | 2 | -268.978 | 542.115 | 0.369 | – | 0.369 | – |
| 333.372.399/ | 10 | -644.543 | 1310.815 | 45.901 | BCST DCST | 2 | -343.508 | 691.138 | 0.294 | – | 0.294 | – |
| 333.372.399/296 | 14 | -640.993 | 1313.002 | 48.088 | BVAR DCST | 4 | -154.223 | 317.658 | 0.205 | 0.002 | 0.243 | – |
| 233.309.333.372/ | 12 | -644.074 | 1314.448 | 49.534 | BVAR DCST | 3 | -250.287 | 506.898 | 0.338 | 0 | 0.36 | – |
| 233.399/ | 7 | -650.62 | 1315.956 | 51.042 | BCST DCST | 2 | -345.315 | 694.751 | 0.32 | – | 0.32 | – |
| 309.333.399/ | 10 | -647.122 | 1316.02 | 51.105 | BCST DCST | 2 | -334.622 | 673.362 | 0.364 | – | 0.364 | – |
| 309.333.399/296 | 14 | -642.88 | 1316.757 | 51.842 | BCST DVAR | 4 | -144.645 | 298.433 | 0.4 | – | 0.458 | -0.002 |
| 372.399/ | 7 | -651.937 | 1318.883 | 53.969 | BVAR | 2 | -453.855 | 911.796 | 0.204 | -0.038 | – | – |
| 333.399/ | 8 | -652.363 | 1321.907 | 56.992 | BVAR | 2 | -408.71 | 821.517 | 0.211 | -0.04 | – | – |
| 309.372.399/ | 9 | -651.355 | 1322.309 | 57.395 | BVAR | 2 | -384.425 | 772.955 | 0.211 | -0.04 | – | – |
| 399/ | 5 | -656.845 | 1324.163 | 59.249 | BVAR | 2 | -516.145 | 1036.363 | 0.223 | -0.04 | – | – |
| 233.333.372/ | 9 | -653.286 | 1326.097 | 61.183 | BCST DCST | 2 | -328.346 | 660.814 | 0.285 | – | 0.285 | – |
| 309.399/ | 7 | -656.143 | 1327.343 | 62.428 | BVAR | 2 | -446.595 | 897.276 | 0.232 | -0.042 | – | – |
| 233.309.333/ | 9 | -654.213 | 1327.998 | 63.084 | BCST DCST | 2 | -317.808 | 639.735 | 0.36 | – | 0.36 | – |
| 309.372.399/296 | 13 | -649.949 | 1328.421 | 63.506 | BVAR DCST | 4 | -197.283 | 403.4 | 0.323 | 0.001 | 0.353 | – |
| 333.399/296 | 12 | -653.371 | 1332.702 | 67.788 | BVAR DCST | 4 | -223.984 | 456.645 | 0.297 | 0.001 | 0.323 | – |
| 233.309.372/ | 8 | -660.162 | 1337.719 | 72.805 | BCST DCST | 2 | -369.327 | 742.759 | 0.343 | – | 0.343 | – |
| 233.333/ | 7 | -662.208 | 1339.395 | 74.481 | BCST DCST | 2 | -394.651 | 793.399 | 0.321 | – | 0.321 | – |
| 333.372/ | 7 | -662.282 | 1339.846 | 74.931 | BVAR | 2 | -501.947 | 1007.968 | 0.219 | -0.04 | – | – |
| 372.399/296 | 11 | -658.844 | 1341.362 | 76.448 | BVAR DCST | 4 | -275.026 | 558.608 | 0.264 | 0 | 0.267 | – |

| | | | | | | | | | | | | |
|---|---|---|---|---|---|---|---|---|---|---|---|---|
| 309.333.372/ | 9 | -660.96 | 1341.785 | 76.871 | BVAR | 2 | -431.777 | 867.641 | 0.23 | -0.042 | – | – |
| 372/ | 4 | -666.665 | 1341.912 | 76.997 | BVAR | 2 | -609.282 | 1222.624 | 0.228 | -0.04 | – | – |
| 333/ | 5 | -665.582 | 1341.913 | 76.999 | BVAR | 2 | -562.63 | 1129.323 | 0.236 | -0.042 | – | – |
| 309.333/ | 7 | -663.969 | 1343.267 | 78.353 | BVAR | 2 | -492.169 | 988.41 | 0.249 | -0.043 | – | – |
| **No shift** | **2** | **-669.909** | **1343.871** | **78.956** | **–** | **–** | **–** | **–** | **–** | **–** | **–** | **–** |
| 309.372/ | 6 | -665.437 | 1344.036 | 79.121 | BVAR | 2 | -539.207 | 1082.481 | 0.238 | -0.041 | – | – |
| 309/ | 4 | -668.436 | 1345.502 | 80.588 | BVAR | 2 | -599.588 | 1203.235 | 0.251 | -0.042 | – | – |
| 309.399/296 | 11 | -661.906 | 1347.52 | 82.606 | BVAR DCST | 4 | -266.623 | 541.786 | 0.361 | 0.001 | 0.376 | – |
| 233.372/ | 6 | -667.73 | 1348.266 | 83.352 | BCST DCST | 2 | -445.743 | 895.573 | 0.311 | – | 0.31 | – |
| 233.309/ | 6 | -669.396 | 1351.645 | 86.731 | BCST DCST | 2 | -435.943 | 875.972 | 0.366 | – | 0.366 | – |
| 309.333.372/296 | 13 | -662.742 | 1354.007 | 89.093 | BCST DVAR | 4 | -247.824 | 504.196 | 0.327 | – | 0.357 | -0.001 |
| 399/296 | 8 | -669.455 | 1355.749 | 90.835 | BVAR | 3 | -343.02 | 692.282 | 0.198 | -0.04 | – | – |
| 233/ | 4 | -677.083 | 1362.436 | 97.522 | BCST DCST | 2 | -512.479 | 1029.03 | 0.333 | – | 0.331 | – |
| 333.372/296 | 11 | -671.859 | 1367.535 | 102.621 | BVAR DCST | 4 | -325.789 | 659.991 | 0.143 | -0.006 | 0.054 | – |
| 309.333/296 | 11 | -672.973 | 1369.803 | 104.889 | BVAR DCST | 4 | -315.438 | 639.279 | 0.356 | 0.001 | 0.37 | – |
| 372/296 | 7 | -680.469 | 1375.832 | 110.918 | BVAR | 3 | -437.351 | 880.878 | 0.211 | -0.041 | – | – |
| 333/296 | 8 | -679.552 | 1376.179 | 111.265 | BVAR | 3 | -390.864 | 787.922 | 0.218 | -0.043 | – | – |
| 309.372/296 | 9 | -679.548 | 1378.598 | 113.684 | BVAR | 3 | -367.582 | 741.377 | 0.221 | -0.043 | – | – |
| 296/ | 5 | -684.369 | 1379.082 | 114.167 | BVAR | 2 | -185.735 | 375.667 | 0.259 | -0.035 | – | – |
| 309/296 | 7 | -683.181 | 1381.303 | 116.389 | BVAR | 3 | -428.598 | 863.369 | 0.243 | -0.045 | – | – |

**Table S5:** A) Global comparison of combinations of diversification shifts for Cetacea (shifts are tested at the ster of the diversification model for their backbones. When the combination has multiple backbones, only the deepest details in Appendix S4). The best combinations of shifts and the phylogeny analysed with no shift are highlig Number of parameters, logL = log(Likelihood), λ = speciation rate (at present if variable), α = dependency para rate, μ = extinction rate, β = dependency parameter of extinction rate

| **A) Total** | | | | | **B) Backbone** | | | | |
|---|---|---|---|---|---|---|---|---|---|
| **Combination** | **NP** | **logL** | **AICc** | **ΔAICc** | **Model** | **NP** | **logL** | **AICc** | **λ** |
| **95.109.135.140/** | **12** | **-252.099** | **540.041** | **0** | **BVAR DCST** | **3** | **-58.299** | **124.598** | **0.228** |
| **140/** | **4** | **-265.784** | **540.187** | **0.147** | **BVAR** | **2** | **-180.045** | **364.334** | **0.06** |
| **95.109.140/** | **10** | **-256.898** | **540.971** | **0.931** | **BVAR DCST** | **3** | **-77.879** | **163.092** | **0.162** |
| 109.140/ | 7 | -263.658 | 543.246 | 3.206 | BVAR DCST | 3 | -110.4 | 227.689 | 0.088 |
| 109.135.140/ | 9 | -260.7 | 545.584 | 5.543 | BVAR DCST | 3 | -92.66 | 192.463 | 0.088 |
| 89.109.140/ | 9 | -261.82 | 545.682 | 5.641 | BVAR DCST | 3 | -57.419 | 122.839 | 0.147 |
| 135.140/ | 6 | -264.535 | 545.725 | 5.684 | BVAR | 2 | -164.015 | 332.308 | 0.053 |
| 89.109.135.140/ | 11 | -257.023 | 546.087 | 6.046 | BVAR DCST | 3 | -37.84 | 85.68 | 0.218 |
| 95.140/ | 6 | -264.542 | 546.357 | 6.316 | BCST | 1 | -153.042 | 308.182 | 0.077 |
| 89.140/ | 6 | -267.61 | 548.947 | 8.907 | BVAR | 2 | -130.728 | 265.809 | 0.057 |
| 140/103 | 5 | -269.477 | 549.99 | 9.95 | BCST | 1 | -53.761 | 109.83 | 0.075 |
| 95.140/103 | 9 | -261.288 | 550.104 | 10.063 | BCST DCST | 2 | -19.811 | 47.621 | 0.924 |
| 95.135.140/ | 10 | -261.11 | 552.122 | 12.082 | BVAR DCST | 3 | -134.828 | 276.384 | 0.083 |
| 109.140/103 | 9 | -265.482 | 553.95 | 13.91 | BCST | 1 | -53.761 | 109.83 | 0.075 |
| 95.109.140/103 | 13 | -257.293 | 554.064 | 14.023 | BCST DCST | 2 | -19.811 | 47.621 | 0.924 |
| 95.109.140/89 | 12 | -259.267 | 554.376 | 14.336 | BVAR DCST | 3 | -57.419 | 122.839 | 0.147 |
| 89.135.140/ | 8 | -266.295 | 554.394 | 14.353 | BVAR | 2 | -114.631 | 233.691 | 0.048 |
| 95.109.135.140/89 | 14 | -254.47 | 554.781 | 14.741 | BVAR DCST | 3 | -37.84 | 85.68 | 0.218 |
| **No shift** | **1** | **-276.789** | **555.626** | **15.585** | – | – | – | – | – |
| 135.140/103 | 8 | -267.24 | 556.052 | 16.011 | BCST | 1 | -53.761 | 109.83 | 0.075 |
| 95.135.140/103 | 12 | -259.051 | 556.165 | 16.125 | BCST DCST | 2 | -19.811 | 47.621 | 0.924 |
| 109.135.140/103 | 11 | -261.016 | 557.381 | 17.341 | BCST | 1 | -53.761 | 109.83 | 0.075 |
| 95.109.135.140/103 | 15 | -252.827 | 557.495 | 17.454 | BCST DCST | 2 | -19.811 | 47.621 | 0.924 |
| 95.140/89 | 9 | -265.057 | 557.642 | 17.601 | BVAR | 2 | -130.728 | 265.809 | 0.057 |
| 109/ | 3 | -276.668 | 560.066 | 20.025 | BCST | 1 | -209.15 | 420.362 | 0.111 |
| 95/ | 4 | -274.059 | 560.971 | 20.931 | BCST | 1 | -248.298 | 498.649 | 0.11 |
| 89/ | 3 | -277.307 | 561.672 | 21.631 | BCST | 1 | -226.164 | 454.386 | 0.119 |
| 95.109/ | 7 | -271.141 | 561.971 | 21.931 | BCST DCST | 2 | -177.861 | 359.945 | 0.186 |
| 103/ | 3 | -277.792 | 562.066 | 22.025 | BCST | 1 | -53.761 | 109.83 | 0.075 |
| 95.103/ | 7 | -269.603 | 562.18 | 22.139 | BCST DCST | 2 | -19.811 | 47.621 | 0.924 |
| 89.109/ | 6 | -274.371 | 562.659 | 22.618 | BCST DCST | 2 | -155.709 | 315.668 | 0.206 |
| 95.135.140/89 | 11 | -263.742 | 563.088 | 23.047 | BVAR | 2 | -114.631 | 233.691 | 0.048 |
| 135/ | 3 | -276.569 | 563.188 | 23.148 | BCST | 1 | -261.787 | 525.625 | 0.104 |
| 109/103 | 5 | -277.117 | 565.459 | 25.418 | BCST | 1 | -53.761 | 109.83 | 0.075 |
| 95.109/103 | 9 | -268.928 | 565.572 | 25.532 | BCST DCST | 2 | -19.811 | 47.621 | 0.924 |
| 95.109.135/ | 10 | -268 | 565.976 | 25.936 | BVAR DCST | 3 | -159.938 | 326.386 | 0.24 |
| 109.135/ | 6 | -275.14 | 567.158 | 27.117 | BCST DCST | 2 | -192.84 | 389.89 | 0.153 |

| | | | | | | | | | |
|---|---|---|---|---|---|---|---|---|---|
| 89.109.135/ | 9 | -271.76 | 567.773 | 27.732 | BVAR DCST | 3 | -138.317 | 283.219 | 0.248 |
| 95.135/ | 6 | -273.882 | 568.621 | 28.581 | BCST | 1 | -233.339 | 468.736 | 0.108 |
| 89.135/ | 5 | -277.218 | 569.498 | 29.458 | BCST | 1 | -211.293 | 424.649 | 0.118 |
| 135/103 | 5 | -277.691 | 569.879 | 29.839 | BCST | 1 | -53.761 | 109.83 | 0.075 |
| 95.135/103 | 9 | -269.501 | 569.993 | 29.952 | BCST DCST | 2 | -19.811 | 47.621 | 0.924 |
| 95/89 | 6 | -274.754 | 570.366 | 30.326 | BCST | 1 | -226.164 | 454.386 | 0.119 |
| 95.109/89 | 9 | -271.818 | 571.353 | 31.312 | BCST DCST | 2 | -155.709 | 315.668 | 0.206 |
| 109.135/103 | 7 | -277.107 | 573.474 | 33.434 | BCST | 1 | -53.761 | 109.83 | 0.075 |
| 95.109.135/103 | 11 | -268.918 | 573.588 | 33.547 | BCST DCST | 2 | -19.811 | 47.621 | 0.924 |
| 95.109.135/89 | 12 | -269.207 | 576.467 | 36.426 | BVAR DCST | 3 | -138.317 | 283.219 | 0.248 |
| 95.135/89 | 8 | -274.665 | 578.192 | 38.152 | BCST | 1 | -211.293 | 424.649 | 0.118 |

n) with B) the rates
: is displayed (more
hted in bold. NP =
meter of speciation

| α | μ | β |
|---|---|---|
| **0.043** | **0.474** | **–** |
| **0.024** | **–** | **–** |
| **0.044** | **0.333** | **–** |
| 0.045 | 0.146 | – |
| 0.051 | 0.183 | – |
| 0.046 | 0.309 | – |
| 0.03 | – | – |
| 0.045 | 0.476 | – |
| – | – | – |
| 0.033 | – | – |
| – | – | – |
| – | 0.924 | – |
| 0.043 | 0.125 | – |
| – | – | – |
| – | 0.924 | – |
| 0.046 | 0.309 | – |
| 0.041 | – | – |
| 0.045 | 0.476 | – |
| – | – | – |
| – | – | – |
| – | 0.924 | – |
| – | – | – |
| – | 0.924 | – |
| 0.033 | – | – |
| – | – | – |
| – | – | – |
| – | – | – |
| – | 0.124 | – |
| – | – | – |
| – | 0.924 | – |
| – | 0.134 | – |
| 0.041 | – | – |
| – | – | – |
| – | – | – |
| – | 0.924 | – |
| 0.022 | 0.302 | – |
| – | 0.081 | – |

| | | |
|---|---|---|
| 0.018 | 0.277 | – |
| – | – | – |
| – | – | – |
| – | – | – |
| – | 0.924 | – |
| – | – | – |
| – | 0.134 | – |
| – | – | – |
| – | 0.924 | – |
| 0.018 | 0.277 | – |
| – | – | – |
