## Appendix S3 for "Estimating clade-specific diversification rates and palaeodiversity dynamics from reconstructed phylogenies"

### Appendix S3: Simulation framework

Estimating clade-specific diversification rates and paleodiversity dynamics from reconstructed phylogenies

Mazet Nathan, Morlon Hélène, Fabre Pierre-Henri, Condamine Fabien

2023-06-30

#### Contents

This document reproduces the simulation analyses of Mazet *et al.* (in prep). The first part generates the parameter values with constrains explained in the main manuscript. Then, the second part sets up the `Snakemake` pipeline in bash. Finally, the third part analyses the results from the `Snakemake` pipeline.

#### Loading packages

In this part, we install different packages we need. For RPANDA, you can either install the version from the GitHub repository of RPANDA with its specific branch (named clade.shift.model) or used the archive file available in the folder (RPANDA_2.2_clade.shift.model_300623.tar.gz).

#### I - Generating the parameter combinations

In this first part, we selected the parameter values for the four diversification models we tested in the simulations following the explanations provided in the main manuscript. The simulated trees were created with the function `tess.sim.age()` from the package TESS. In TESS functions, time is defined forward in time while RPANDA defines time backward in time. In other words, for model with rate varying exponentially through time, in `TESS`, $\lambda$ gives the rate value at the root while in `RPANDA`, it gives the rate value at present. In both case $\alpha$ gives the dependency in the same direction than the definition of time. Forward in time (TESS), a positive $\alpha$ corresponds to an increase of the speciation rate through time. Backward in time (RPANDA), a positive $\alpha$ corresponds to a **decrease** of the speciation rate through time. To avoid the confusion, we keep the `TESS` format for now and defined `lambda0` and `mu0` as the rate values at present. For the sake of reproductivness, we set up a seed.

```
set.seed(1612)
param <- data.frame(lambda = rep(NA, 4),
                    lambda0 = rep(NA, 4),
                    alpha = rep(NA, 4),
                    mu = rep(NA, 4),
                    mu0 = rep(NA, 4),
                    beta = rep(NA, 4),
                    tmrca = rep(NA, 4))
```

##### 1. Models

The parameter values for the four models were selected with the following code to follow the conditions as explained in the main text (section *2.3 / Simulations*)

```

j = 1

tmrca = 100
t <- 0:tmrca

tree.expanding <- list()
seq_lambda <- seq(0.05, 0.25, 0.01)

while(length(tree.expanding) <= 80){
  cat("next \n")
  lambda = seq_lambda[1]
```

```
  seq_lambda <- seq_lambda[-1]
  mu = 0

  tree.expanding <- tess.sim.age(n = 100, age = tmrca, lambda = lambda, mu = mu)

  if(length(tree.expanding) != 0 ){
    tree.expanding <- tree.expanding[Ntip.multiPhylo(tree.expanding) > 40]
  }

  if(length(tree.expanding) != 0 ){
    tree.expanding <- tree.expanding[sapply(tree.expanding,
                                            function(x)
                                              any(branching.times(x) > 25
                                                  & branching.times(x) < 33))]
  }

  if(length(tree.expanding) != 0 ){
    enough_tips <- c()
    for(i in 1:length(tree.expanding)){
      node <- branching.times(tree.expanding[[i]])[
        branching.times(tree.expanding[[i]]) > 25 &
          branching.times(tree.expanding[[i]]) < 33]
      node_desc <- sapply(names(node),
                          function(x) length(Descendants(tree.expanding[[i]],
                                                         as.numeric(x))[[1]]))
      node <- node[node_desc > 5]
      if(length(node) >= 1){
        node <- as.numeric(names(node)[node == min(node)])
        enough_tips <- c(enough_tips, T)
      } else {
        enough_tips <- c(enough_tips, F)
      }

    }
    tree.expanding <- tree.expanding[enough_tips]
  }
}

length(tree.expanding)
#hist(Ntip.multiPhylo(tree.expanding))
summary(Ntip.multiPhylo(tree.expanding))

param[j,] <- c(lambda, lambda, NA, NA, NA, NA, tmrca)
j = j + 1
```

###### Model 1: Expanding diversity (BCST)

```
tree.saturing <- list()
seq_lambda <- seq(0.05, 0.25, 0.01)
```

```
while(length(tree.saturing) <= 80){
  cat("next \n")
  lambda = seq_lambda[1]
  seq_lambda <- seq_lambda[-1]
  alpha = log(0.5)/sample(seq(tmrca*2/3, tmrca*3/4, 1), 1)

  speciation <- function(t) lambda*exp(alpha*t)

  tree.saturing <- tess.sim.age(n = 100, age = tmrca, lambda = speciation, mu = 0)

  if(length(tree.saturing) != 0 ){
    tree.saturing <- tree.saturing[Ntip.multiPhylo(tree.saturing) > 40]
  }

  if(length(tree.saturing) != 0 ){
    tree.saturing <- tree.saturing[sapply(tree.saturing,
                                          function(x) any(branching.times(x) > 25 &
                                                           branching.times(x) < 33))]
  }

  if(length(tree.saturing) != 0 ){
    enough_tips <- c()
    for(i in 1:length(tree.saturing)){
      node <- branching.times(tree.saturing[[i]])[
        branching.times(tree.saturing[[i]]) > 25 &
          branching.times(tree.saturing[[i]]) < 33]
      node_desc <- sapply(names(node),
                          function(x) length(Descendants(tree.saturing[[i]],
                                                          as.numeric(x))[[1]]))
      node <- node[node_desc > 5]
      if(length(node) >= 1){
        node <- as.numeric(names(node)[node == min(node)])
        enough_tips <- c(enough_tips, T)
      } else {
        enough_tips <- c(enough_tips, F)
      }
    }
    tree.saturing <- tree.saturing[enough_tips]
  }

}

lambda0 <- lambda*exp(alpha*t)[length(t)]
param[j,] <- c(lambda, lambda0, -alpha, NA, NA, NA, tmrca)
j = j + 1
```

###### Model 2: Saturing diversity (BVAR)

```
tree.ww.s <- list()
seq_lambda <- seq(0.2, 0.4, 0.01)
```

```
seq_mu <- seq_lambda/2

while(length(tree.ww.s) <= 80){
  cat("next \n")
  lambda <- seq_lambda[1]
  mu = seq_mu[1]

  seq_lambda <- seq_lambda[-1]
  seq_mu <- seq_mu[-1]

  alpha = log(mu/lambda)/sample(seq(tmrca*2/3, tmrca*3/4, 1),1)

  speciation <- function(t) lambda*exp(alpha*t)

  # a minimum of 40 species
  tree.ww.s <- tess.sim.age(n = 100, age = tmrca, lambda = speciation, mu = mu)

  if(length(tree.ww.s) != 0 ){
    tree.ww.s <- tree.ww.s[Ntip.multiPhylo(tree.ww.s) > 40]
  }

  if(length(tree.ww.s) != 0 ){
    tree.ww.s <- tree.ww.s[sapply(tree.ww.s,
                               function(x) any(branching.times(x) > 25 &
                                                 branching.times(x) < 33))]
  }

  if(length(tree.ww.s) != 0){
    enough_tips <- c()
    for(i in 1:length(tree.ww.s)){
      node <- branching.times(tree.ww.s[[i]])[branching.times(tree.ww.s[[i]]) > 25 &
                                                branching.times(tree.ww.s[[i]]) < 33]
      node_desc <- sapply(names(node),
                          function(x) length(Descendants(tree.ww.s[[i]],
                                                         as.numeric(x))[[1]]))
      node <- node[node_desc > 5]
      if(length(node) >= 1){
        node <- as.numeric(names(node)[node == min(node)])
        enough_tips <- c(enough_tips, T)
      } else {
        enough_tips <- c(enough_tips, F)
      }
    }
    tree.ww.s <- tree.ww.s[enough_tips]
  }
}

lambda0 <- lambda*exp(alpha*t)[length(t)]

param[j,] <- c(lambda, lambda0, -alpha, mu, mu, NA, tmrca)
j = j + 1
```

**Model 3: Waxing-waning with speciation rate decreasing (BVAR DCST)**

```
seq_lambda <- seq(0.2, 0.4, 0.01)
seq_mu <- seq_lambda/2
#seq_mu <- rep(0.01, length(seq_lambda))

tree.ww.e <- list()
while(length(tree.ww.e) < 80){
  cat("next \n")
  lambda <- seq_lambda[1]
  mu <- seq_mu[1]

  seq_lambda <- seq_lambda[-1]
  seq_mu <- seq_mu[-1]

  beta = log(lambda/mu)/sample(seq(tmrca*2/3, tmrca*3/4, 1),1)

  extinction <- function(t) mu*exp(beta*t)

  lambda - extinction(t)

  tree.ww.e <- tess.sim.age(n = 100, age = tmrca, lambda = lambda, mu = extinction)

  tree.ww.e <- tree.ww.e[Ntip.multiPhylo(tree.ww.e) > 40]

  if(length(tree.ww.e) > 0){
    tree.ww.e <- tree.ww.e[sapply(tree.ww.e, function(x) any(branching.times(x) > 25 &
                                                             branching.times(x) < 33))]
  }

  if(length(tree.ww.e) != 0 ){
    enough_tips <- c()
    for(i in 1:length(tree.ww.e)){
      node <- branching.times(tree.ww.e[[i]])[branching.times(tree.ww.e[[i]]) > 25 &
                                               branching.times(tree.ww.e[[i]]) < 33]
      node_desc <- sapply(names(node), function(x) length(Descendants(tree.ww.e[[i]],
                                                                       as.numeric(x))[[1]]))
      node <- node[node_desc > 5]
      if(length(node) >= 1){
        node <- as.numeric(names(node)[node == min(node)])
        enough_tips <- c(enough_tips, T)
      } else {
        enough_tips <- c(enough_tips, F)
      }
    }
    tree.ww.e <- tree.ww.e[enough_tips]
  }
}

mu0 <- mu*exp(beta*t)[length(t)]

param[j,] <- c(lambda, lambda, NA, mu, mu0, -beta, tmrca)

param$models <- c("BCST", "BVAR", "BVAR_DCST", "BCST_DVAR")
```

**Model 4: Waxing-waning with extinction rate increasing (BCST DVAR)**

##### 2. Output

We can represent the evolution of rates through time for the four models.

```
par(mfrow = c(2,2), cex = 0.5)
# Scenario 1
plot(0:tmrca, rep(param$lambda[1],length(tmrca:0)), xaxt = "n", type = "l",
     main = "Model 1: Expanding diversity (BCST)", col = "blue",
     ylab = "Rates (Event / Lineage / Myr)", xlab = "Time (Myrs)", ylim = c(0,0.3),
     lwd = 1, las = 1)
axis(side = 1, las = 1, line = 0, xpd = TRUE, labels = -rev(seq(t[1], t[length(t)], 5)),
     at = seq(t[1], t[length(t)], 5))
legend("topleft", legend = c("Speciation rate", "Extinction rate"),
       col = c("blue", "red"), lty = 1, lwd = 1, bty = "n")
# Scenario 2
speciation <- function(t) param$lambda[2]*exp(-param$alpha[2]*t)
plot(tmrca:0, speciation(tmrca:0), xaxt = "n", type = "l",
     main = "Model 2: Saturing diversity (BVAR)", col = "blue",
     ylab = "Rates (Event / Lineage / Myr)", xlab = "Time (Myrs)", ylim = c(0,0.3),
     lwd = 1, las = 1)
axis(side = 1, las = 1, line = 0, xpd = TRUE, labels = -rev(seq(t[1], t[length(t)], 5)),
     at = seq(t[1], t[length(t)], 5))

# Scenario 3
speciation <- function(t) param$lambda[3]*exp(-param$alpha[3]*t) # forward in time

plot(tmrca:0, speciation(tmrca:0), xaxt = "n", type = "l",
     main = "Model 3: Waxing-waning dynamic (BVAR DCST)", col = "blue",
     ylab = "Rates (Event / Lineage / Myr)", xlab = "Time (Myrs)", ylim = c(0,0.3),
     lwd = 1, las = 1)
axis(side = 1, las = 1, line = 0, xpd = TRUE, labels = -rev(seq(t[1], t[length(t)], 5)),
     at = seq(t[1], t[length(t)], 5))
lines(tmrca:0, rep(param$mu[3], length(tmrca:0)), col = "red", lwd = 1)

# Scenario 4
extinction <- function(t) param$mu[4]*exp(-param$beta[4]*t)

plot(tmrca:0, extinction(tmrca:0), xaxt = "n", type = "l",
     main = "Model 4: Waxing-waning dynamic (BCST DVAR)", col = "red",
     ylab = "Rates (Event / Lineage / Myr)", xlab = "Time (Myrs)", ylim = c(0,0.3),
     lwd = 1, las = 1)
axis(side = 1, las = 1, line = 0, xpd = TRUE, labels = -rev(seq(t[1], t[length(t)], 5)),
     at = seq(t[1], t[length(t)], 5))
lines(tmrca:0, rep(param$lambda[4], length(tmrca:0)), col = "blue", lwd = 1)
```

Now, we can export the table with rate values of the four models for the next step made with `Snakemake`. Because we only used the `TESS` package for simulations and that analyses are made with the `RPANDA` package, the rates will be convert backward in time such as $\lambda$ and $\mu$ gives respectively the values of speciation and extinction rates at present.

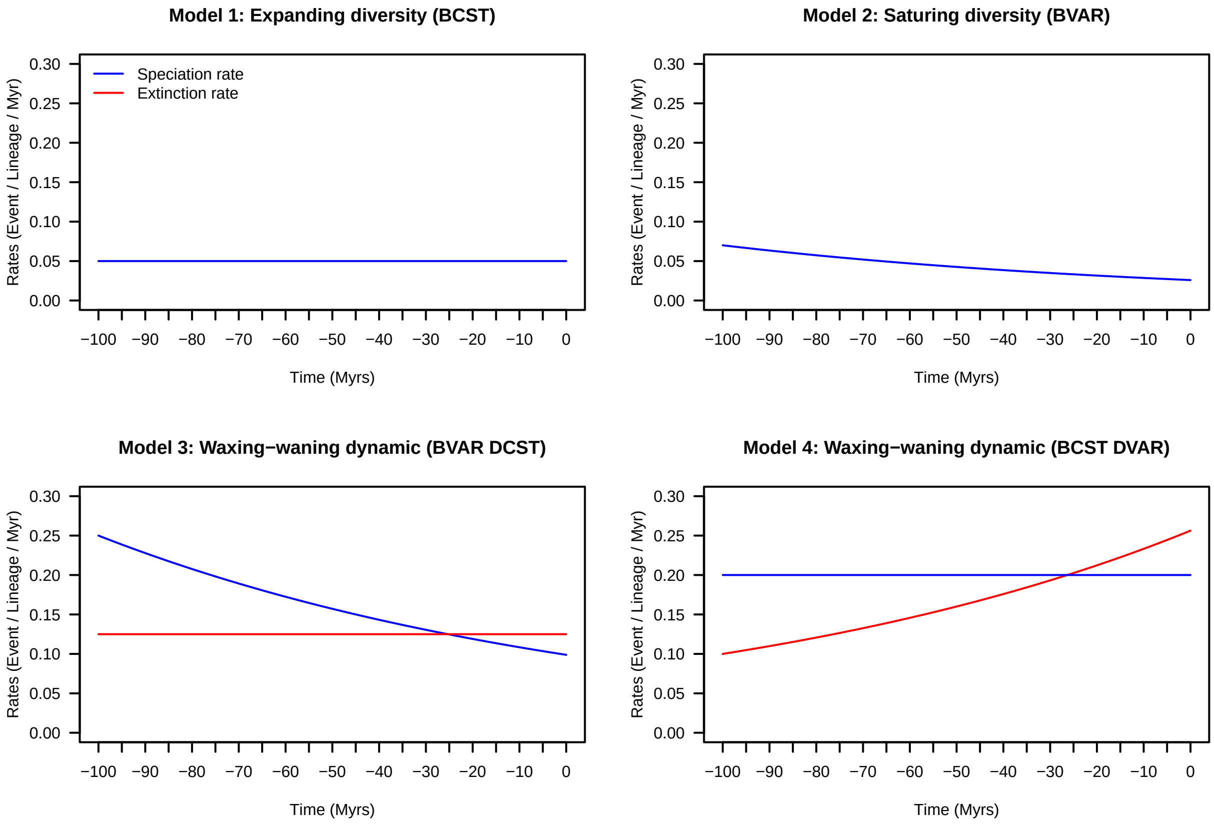

Figure 1: Evolution of diversification rates through time for the four simulated models.

```r
# saving the full config file
write.csv(param, "config_comb_full.csv", row.names = F)

# keeping RPANDA format of rate values
param$tmrca <- 100
param <- param[,c("models", "lambda0", "alpha", "mu0", "beta", "tmrca")]
names(param) <- c("models", "lambda", "alpha", "mu", "beta", "tmrca")

write.csv(param, "config_comb_bigmem.csv", row.names = F)
```

#### II - Launching the `Snakemake` pipeline

The pipeline is composed of two Snakefiles that call R scripts stored in the folder scripts. The Snakefile titled `Snakefile_simul` simulates the raw phylogenies from model values contained in the table config_comb_bigmem.csv thanks to the script tree_sim.R. This Snakefile also creates phylogenies with the shift of diversification from raw phylogenies using the script create_comb.R. The second Snakefile, `Snakefile_comb`, applies the automation as described in the Fig. 1 of the main manuscript for both raw phylogenies and phylogenies with a shift. It returns outputs in rds files for each analysis. We split the Snakefiles into two files in order to check simulated trees before analysing them.

Simulations were launched on a bigmem system shared with other users. Consequently, we created a conda environment with all the R packages needed for the simulations. This environment called `simshift_env` can be set up by executing the script `simshifts_setup.sh` as follow:

```
# in a terminal
sh simshifts_setup.sh
```

This script uses an environment file (environment.yaml) to create the conda environment. It also runs the R script `pkgs.R` located in the folder `scripts` to install all necessary packages.

Once the conda envrionment is setup, we can activate it and launch the `Snakefile_simul` to simulate trees.

```
# in a terminal
conda activate simshift_env
snakemake -s Snakefile_simul -j 30
```

Here, the argument `-j 30` corresponds to the number of cores we want to allocate.

After checking that simulated trees corresponds to what we wanted, we can start analysing trees by launching the `Snakefile_comb`:

```
# in a terminal
conda activate simshift_env
snakemake -s Snakefile_comb -j 30
```

#### III - Analyses

When the simulations are over, we can start analysing them.

##### 1. Loading trees and results

In the following lines, we loaded trees and results.

```
# Models
models <- c("BCST", "BVAR", "BVAR_DCST", "BCST_DVAR")

# storage
all_tree_r <- list()
all_tree_s <- list()

all_res_r <- list()
all_res_s <- list()

# Trees
for(res.i in 1:length(models)){
  cat("\n", res.i, "/", length(models))
  names_tree <- list.files("./data/trees", pattern = paste0(models[res.i],"@"),
                          full.names = T)
  names_tree_r <- names_tree[grepl("raw", names_tree)]
  names_tree_r <- names_tree_r[gsub("_raw.tre", "", sapply(strsplit(names_tree_r,
                                                                    split = "_n"),
                                                           "[[", 2)) %in%
                                as.character(1:1000)]
  names_tree_s <- names_tree[grepl("shift", names_tree)]
  names_tree_s <- names_tree_s[gsub("_shift.tre", "", sapply(strsplit(names_tree_s,
                                                                      split = "_n"),
                                                             "[[", 2)) %in%
                                as.character(1:1000)]

  tree_r <- rep(list(NULL), 1000)
  tree_s <- rep(list(NULL), 1000)
  names(tree_r) <- gsub(".tre", "", sapply(strsplit(names_tree_r, "_n"), "[[", 2))
  names(tree_s) <- gsub(".tre", "", sapply(strsplit(names_tree_s, "_n"), "[[", 2))

  # Res
  names_res <- list.files("./data/res", pattern = paste0(models[res.i],"@"),
                         full.names = T)
  names_res_r <- names_res[grepl("raw", names_res)]
  names_res_s <- names_res[grepl("shift", names_res)]

  res_r <- rep(list(NULL), 1000)
  res_s <- rep(list(NULL), 1000)
  names(res_r) <- gsub(".rds", "", sapply(strsplit(names_res_r, "_n"), "[[", 2))
  names(res_s) <- gsub(".rds", "", sapply(strsplit(names_res_s, "_n"), "[[", 2))

  # load trees and results
  for(i in 1:1000){
    cat("\n\t",i, "/ 1000")
    tree_r[i] <- list(ladderize(read.tree(names_tree_r[i]), F))
    tree_s[i] <- list(ladderize(read.tree(names_tree_s[i]), F))

    res_r[i] <- list(readRDS(names_res_r[i]))
    res_s[i] <- list(readRDS(names_res_s[i]))
```

```
  }

  all_tree_s[res.i] <- list(tree_s)
  all_tree_r[res.i] <- list(tree_r)

  all_res_r[res.i] <- list(res_r)
  all_res_s[res.i] <- list(res_s)
}

names(all_tree_r) <- names(all_tree_s) <- names(all_res_s) <- names(all_res_r) <- models

all_res_r_tot <- lapply(1:length(models),
                        function(res.i) lapply(all_res_r[[res.i]],
                                               function(x) x$total))
all_res_s_tot <- lapply(1:length(models),
                        function(res.i) lapply(all_res_s[[res.i]],
                                               function(x) x$total))

names(all_res_r_tot) <- names(all_res_s_tot) <- models
```

##### 2. Shift detection

In this part, we looked at the shift detection and the rate of false detection of shift.

First, we defined false detections of shift as simulations for which the combination with a shift gets a better fit ($\Delta$AICc $\geq 2$) than the whole tree analysed without shift (with the same diversification model for the whole tree). As explained in the main text, we found that the rates of false detections of shift with simulated models fixed were higher than 5 %, the expected value from the model development from (Morlon et al. 2011). This shows that local optimisation of a same model of diversification (the simulated one) is not penalised enough to avoid selecting a shift whereas it has not been simulated. To solve this issue, we penalised the likelihood of phylogeny with shift by adding a parameter for the location of the shift as in previous methods such as MEDUSA (Alfaro et al. 2009). The following code recreates the results of the old version of the method (without this additional parameter) to compare the rate of false detection of shift.

```
# function to detect a shift
detect_shift <- function(all_res, threshold = 2){

  summary <- c()
  for(i in 1:length(all_res)){
    if(which(all_res[[i]]$Combination == "whole_tree") == 2){
      if(all_res[[i]]$delta_AICc[2] > threshold){
        res <- "True"
      } else {
        res <- "True_ns"
      }
    } else {
      if(all_res[[i]]$delta_AICc[2] > threshold){
        res <- "False"
      } else {
        res <- "False_ns"
      }
    }
    summary[i] <- res
```

```
  }
  return(summary)
}

#### b) type of error with 1 parameter for the shift ####
th = 2
# raw
all_res_r_old <- all_res_r
names(all_res_r_old) <- models
all_res_r_tot_sim <- all_res_r_tot

all_res_r_tot_sim_old <- all_res_r_tot
all_res_r_tot_old <- all_res_r_tot

for(res.i in 1:length(all_res_r)){
  cat(res.i, "/", length(all_res_r), "\n")
  for(i in 1:length(all_res_r[[res.i]])){
    cat("\t", i, "/", length(all_res_r[[res.i]]), "\n")

    total_values <- all_res_r[[res.i]][[i]]$total[
      all_res_r[[res.i]][[i]]$total$Combination != "whole_tree",
      c("Parameters", "logL", "AICc")]

    # simulated model
    whole_values <- all_res_r[[res.i]][[i]]$whole_tree[
      all_res_r[[res.i]][[i]]$whole_tree$Models == models[res.i],
      c("Parameters", "logL", "AICc")]
    backbone_values <- all_res_r[[res.i]][[i]]$backbones[[1]][[1]][
      all_res_r[[res.i]][[i]]$backbones[[1]][[1]]$Models == models[res.i],
      c("Parameters", "logL", "AICc")]
    sub_values <- all_res_r[[res.i]][[i]]$subclades[[1]][
      all_res_r[[res.i]][[i]]$subclades[[1]]$Models == models[res.i],
      c("Parameters", "logL", "AICc")]

    all_res_r_tot_sim[[res.i]][[i]][
      all_res_r_tot_sim[[res.i]][[i]]$Combination != "whole_tree",
      c("Parameters", "logL", "AICc")] <- sub_values + backbone_values
    all_res_r_tot_sim[[res.i]][[i]][
      all_res_r_tot_sim[[res.i]][[i]]$Combination == "whole_tree",
      c("Parameters", "logL", "AICc")] <- whole_values
    all_res_r_tot_sim[[res.i]][[i]]$delta_AICc <-
      all_res_r_tot_sim[[res.i]][[i]]$AICc - min(all_res_r_tot_sim[[res.i]][[i]]$AICc)
    all_res_r_tot_sim[[res.i]][[i]] <- all_res_r_tot_sim[[res.i]][[i]][
      order(all_res_r_tot_sim[[res.i]][[i]]$delta_AICc),]

    # simulated model old version (-1 parameter)
    phy <- all_tree_r[[res.i]][[i]]
    phy_s <- all_tree_s[[res.i]][[i]]

    getMRCA(phy, phy$tip.label[!phy$tip.label %in% phy_s$tip.label])

    nodes <- branching.times(phy)[branching.times(phy) > 25 & branching.times(phy) < 33]
    node_desc <- sapply(Descendants(phy, as.numeric(names(nodes))), length)
```

```
nodes <- nodes[node_desc > 5]
node <- as.numeric(names(nodes[nodes == min(nodes)]))

backbone.phy <- drop.tip(phy, phy$tip.label[Descendants(phy, as.numeric(node))[[1]]])
subclade.phy <- subtree(phy, phy$tip.label[Descendants(phy, as.numeric(node))[[1]]])

NP_sim_subclade <- all_res_r[[res.i]][[i]]$subclades[[1]]$Parameters[
  all_res_r[[res.i]][[i]]$subclades[[1]]$Models == models[res.i]]
NP_sim_backbone <- all_res_r[[res.i]][[i]]$backbones[[1]][[1]]$Parameters[
  all_res_r[[res.i]][[i]]$backbones[[1]][[1]]$Models == models[res.i]]

LH_sim_local_subclade <- all_res_r[[res.i]][[i]]$subclades[[1]]$logL[
  all_res_r[[res.i]][[i]]$subclades[[1]]$Models == models[res.i]]
LH_sim_local_backbone <- all_res_r[[res.i]][[i]]$backbones[[1]][[1]]$logL[
  all_res_r[[res.i]][[i]]$backbones[[1]][[1]]$Models == models[res.i]]
LH_sim_local <- LH_sim_local_backbone+LH_sim_local_subclade

AIC_sim_local_subclade_old <- 2 * -LH_sim_local_subclade +
  2 * (NP_sim_subclade-1) +
  (2 * (NP_sim_subclade-1) * ((NP_sim_subclade-1) + 1))/
  (Ntip(subclade.phy) - (NP_sim_subclade-1) - 1)
AIC_sim_local_backbone <- 2 * -LH_sim_local_backbone +
  2 * (NP_sim_backbone) +
  (2 * (NP_sim_backbone) *
     ((NP_sim_backbone) + 1))/(Ntip(backbone.phy) - (NP_sim_backbone) - 1)

AICc_sim_local_old <- AIC_sim_local_backbone + AIC_sim_local_subclade_old

all_res_r_tot_sim_old[[res.i]][[i]][
  all_res_r_tot_sim_old[[res.i]][[i]]$Combination == "whole_tree",
  c("Parameters", "logL", "AICc")] <- whole_values
all_res_r_tot_sim_old[[res.i]][[i]][
  all_res_r_tot_sim_old[[res.i]][[i]]$Combination != "whole_tree",
  c("Parameters", "logL", "AICc")] <- c(NP_sim_backbone+NP_sim_subclade-1,
                                        LH_sim_local, AICc_sim_local_old)
all_res_r_tot_sim_old[[res.i]][[i]]$delta_AICc <-
  all_res_r_tot_sim_old[[res.i]][[i]]$AICc -
  min(all_res_r_tot_sim_old[[res.i]][[i]]$AICc)
all_res_r_tot_sim_old[[res.i]][[i]] <-
  all_res_r_tot_sim_old[[res.i]][[i]][
    order(all_res_r_tot_sim_old[[res.i]][[i]]$delta_AICc),]

# Best model old method (-1 parameter)
NP_best_subclade <- all_res_r[[res.i]][[i]]$subclades[[1]]$Parameters[
  all_res_r[[res.i]][[i]]$subclades[[1]]$AICc ==
    min(all_res_r[[res.i]][[i]]$subclades[[1]]$AICc)]
NP_best_backbone <-
  all_res_r[[res.i]][[i]]$backbones[[1]][[1]]$Parameters[
    all_res_r[[res.i]][[i]]$backbones[[1]][[1]]$AICc ==
      min(all_res_r[[res.i]][[i]]$backbones[[1]][[1]]$AICc)]

LH_best_local_subclade <- all_res_r[[res.i]][[i]]$subclades[[1]]$logL[
  all_res_r[[res.i]][[i]]$subclades[[1]]$AICc ==
```

```
    min(all_res_r[[res.i]][[i]]$subclades[[1]]$AICc)]
LH_best_local_backbone <- all_res_r[[res.i]][[i]]$backbones[[1]][[1]]$logL[
    all_res_r[[res.i]][[i]]$backbones[[1]][[1]]$AICc ==
      min(all_res_r[[res.i]][[i]]$backbones[[1]][[1]]$AICc)]
LH_best_local <- LH_best_local_backbone+LH_best_local_subclade

AIC_best_local_subclade_old <- 2 * -LH_best_local_subclade +
  2 * (NP_best_subclade-1) +
  (2 * (NP_best_subclade-1) *
     ((NP_best_subclade-1) + 1))/(Ntip(subclade.phy) -
                                    (NP_best_subclade-1) - 1)
AIC_best_local_backbone <- 2 * -LH_best_local_backbone + 2 *
  (NP_best_backbone) +
  (2 * (NP_best_backbone) *
     ((NP_best_backbone) + 1))/(Ntip(backbone.phy) - (NP_best_backbone) - 1)

AICc_best_local_old <- AIC_best_local_backbone + AIC_best_local_subclade_old

all_res_r_tot_old[[res.i]][[i]][
  all_res_r_tot_old[[res.i]][[i]]$Combination != "whole_tree",
  c("Parameters", "logL", "AICc")] <-
  c(NP_best_backbone+NP_best_subclade-1, LH_best_local, AICc_best_local_old)
all_res_r_tot_old[[res.i]][[i]]$delta_AICc <-
  all_res_r_tot_old[[res.i]][[i]]$AICc - min(all_res_r_tot_old[[res.i]][[i]]$AICc)
all_res_r_tot_old[[res.i]][[i]] <-
  all_res_r_tot_old[[res.i]][[i]][order(all_res_r_tot_old[[res.i]][[i]]$delta_AICc),]

# old method (-1 parameter) for all models

NP_best_subclade_old <- all_res_r[[res.i]][[i]]$subclades[[1]]$Parameters
NP_best_backbone_old <- all_res_r[[res.i]][[i]]$backbones[[1]][[1]]$Parameters

LH_best_local_subclade_old <- all_res_r[[res.i]][[i]]$subclades[[1]]$logL
LH_best_local_backbone_old <- all_res_r[[res.i]][[i]]$backbones[[1]][[1]]$logL
LH_best_local_old <- LH_best_local_backbone_old+LH_best_local_subclade_old

all_res_r_old[[res.i]][[i]]$total[
  all_res_r_old[[res.i]][[i]]$total$Combination !=
    "whole_tree", c("Parameters", "logL", "AICc")] <-
  c(NP_best_backbone+NP_best_subclade-1, LH_best_local, AICc_best_local_old)
all_res_r_old[[res.i]][[i]]$total$delta_AICc <-
  all_res_r_old[[res.i]][[i]]$total$AICc -
  min(all_res_r_old[[res.i]][[i]]$total$AICc)
all_res_r_old[[res.i]][[i]]$total <-
  all_res_r_old[[res.i]][[i]]$total[
    order(all_res_r_old[[res.i]][[i]]$total$delta_AICc),]

AIC_best_local_subclade_old <- 2 * -LH_best_local_subclade_old +
  2 * (NP_best_subclade_old-1) +
  (2 * (NP_best_subclade_old-1) * ((NP_best_subclade_old-1) + 1))/
  (Ntip(subclade.phy) - (NP_best_subclade_old-1) - 1)
AIC_best_local_backbone_old <- 2 * -LH_best_local_backbone_old +
  2 * (NP_best_backbone_old) +
```

```
      (2 * (NP_best_backbone_old) * ((NP_best_backbone) + 1))/
      (Ntip(backbone.phy) - (NP_best_backbone_old) - 1)
    AICc_best_local_old <- AIC_best_local_backbone_old +
      AIC_best_local_subclade_old

    all_res_r_old[[res.i]][[i]]$subclades[[1]][,c("Parameters", "logL", "AICc")] <-
      cbind(NP_best_subclade_old-1,
            LH_best_local_subclade_old,
            AIC_best_local_subclade_old)

    all_res_r_old[[res.i]][[i]]$subclades[[1]]$delta_AICc <-
      all_res_r_old[[res.i]][[i]]$subclades[[1]]$AICc -
      min(all_res_r_old[[res.i]][[i]]$subclades[[1]]$AICc)

    all_res_r_old[[res.i]][[i]]$subclades[[1]][
      order(all_res_r_old[[res.i]][[i]]$subclades[[1]]$delta_AICc),]
  }
}
```

We then create a data.frame to summarize the results and reproduce the Figure 3 of the main text (here Figure 2):

```
# data.frame for the output
summary_best_sim_df <- data.frame(FP_best_old = sapply(all_res_r_tot_old,
                                                       function(x)
                                                         sum(detect_shift(x, th) ==
                                                              "True")/length(x)),
                                  FP_best = sapply(all_res_r_tot,
                                                   function(x)
                                                     sum(detect_shift(x, th) ==
                                                          "True")/length(x)),
                                  FP_sim_old = sapply(all_res_r_tot_sim_old,
                                                      function(x)
                                                        sum(detect_shift(x, th) ==
                                                            "True")/length(x)),
                                  FP_sim = sapply(all_res_r_tot_sim,
                                                  function(x)
                                                    sum(detect_shift(x, th) ==
                                                         "True")/length(x)))

par(mfrow = c(1,1), mar = c(5,5,4,4))
a <- barplot(t(summary_best_sim_df), beside = T,
             names.arg = c("BCST", "BVAR", "BVAR DCST", "BCST DVAR"),
             ylim = c(0,1), ylab = "Frequency", xlab = "Models",
             main = "",
             col = c("darkorange2", "darkorange4", "deepskyblue1", "deepskyblue4"),
             las = 1,
             space = c(0.1, 1), cex.names = 1, cex.axis = 1, cex.lab = 1)
mtext(side = 3, text = expression(Delta), line = 1, cex = 1, at = median(a[1,]))
mtext(side = 3, text = paste0("                 AICc < ", th), line = 1, cex = 1,
      at = median(a[1,]))
text(a,unlist(c(summary_best_sim_df[1,], summary_best_sim_df[2,],
                summary_best_sim_df[3,], summary_best_sim_df[4,]))+0.03,
```

```
     labels=c(summary_best_sim_df[1,], summary_best_sim_df[2,],
              summary_best_sim_df[3,], summary_best_sim_df[4,]), cex=0.5)

legend("topright", legend = c("best model (old version)",
                       "best model (+1 parameter)",
                       "simulated model (old version)",
                       "simulated model (+1 parameter)"),
       fill = c("darkorange2", "darkorange4",
                "deepskyblue1", "deepskyblue4"),
       bty = "n", cex = 0.7)
mtext(text = "False positive rates with different methods",
      side = 3, outer = T,
      cex = 1, line = 1)
```

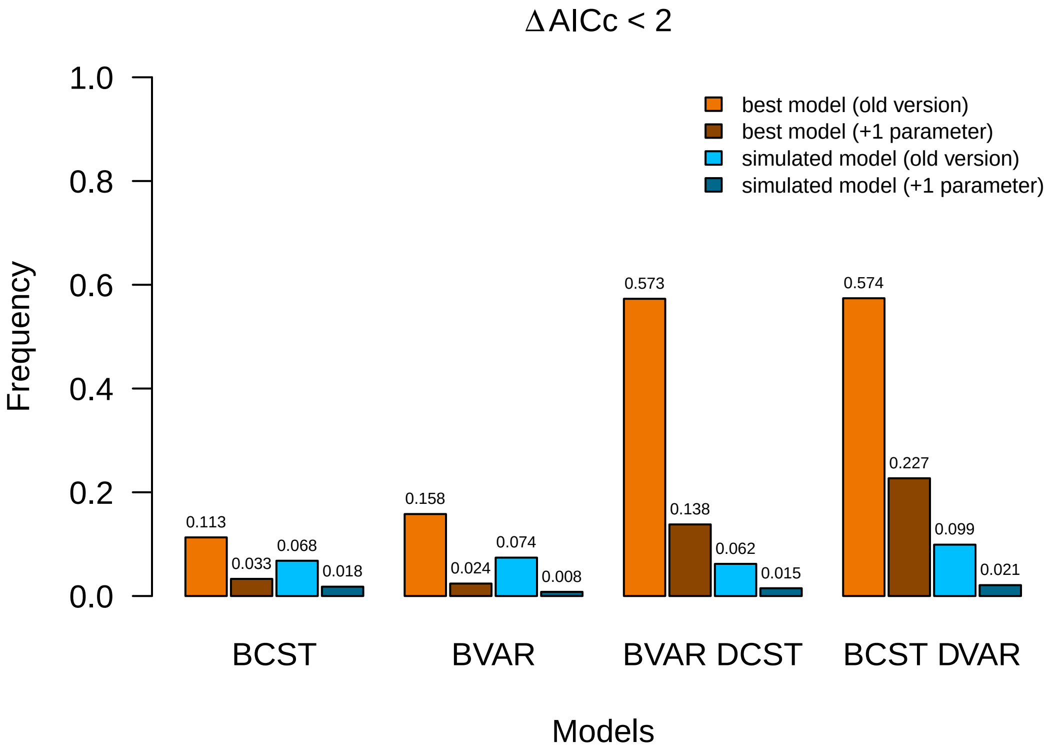

Figure 2: False positive rates with different methods

We also had a look at the differences in rate values between the true model and the best models for the backbones. Here are the code to do so:

```
all_param_s_bck_true <- lapply(1:length(models),
                                function(res.i)
                                  lapply(all_res_s[[res.i]],
                                         function(x) x$backbone[[1]][[1]][
                                           x$backbone[[1]][[1]]$Models == models[res.i],]))
all_param_s_bck_true_df <- lapply(all_param_s_bck_true, function(x) do.call(rbind, x))
```

```
true_lambdas <- sapply(all_param_s_bck_true_df, function(x) x$Lambda)
true_alpha <- sapply(all_param_s_bck_true_df, function(x) x$Alpha)
true_mu <- sapply(all_param_s_bck_true_df, function(x) x$Mu)
true_beta <- sapply(all_param_s_bck_true_df, function(x) x$Beta)

true_param_df <- data.frame(rates = c(true_lambdas, true_alpha, true_mu, true_beta))
true_param_df$param <- rep(c("Lambda", "Alpha", "Mu", "Beta"), each = 4000)
true_param_df$models <- rep(rep(c("BCST", "BVAR", "BVAR_DCST", "BCST_DVAR"),
                                each = 1000), 4)
true_param_df$param <- factor(true_param_df$param)
true_param_df$models <- factor(true_param_df$models)

all_param_s_bck_best <- lapply(1:length(models),
                               function(res.i)
                                 lapply(all_res_s[[res.i]],
                                        function(x) x$backbone[[1]][[1]][1,]))

all_param_s_bck_best_df <- lapply(all_param_s_bck_best, function(x) do.call(rbind, x))

best_lambdas <- sapply(all_param_s_bck_best_df, function(x) x$Lambda)
best_alpha <- sapply(all_param_s_bck_best_df, function(x) x$Alpha)
best_mu <- sapply(all_param_s_bck_best_df, function(x) x$Mu)
best_beta <- sapply(all_param_s_bck_best_df, function(x) x$Beta)

best_param_df <- data.frame(rates = c(best_lambdas, best_alpha, best_mu, best_beta))
best_param_df$param <- rep(c("Lambda", "Alpha", "Mu", "Beta"), each = 4000)
best_param_df$models <- rep(rep(c("BCST", "BVAR", "BVAR_DCST", "BCST_DVAR"),
                                each = 1000), 4)
best_param_df$param <- factor(best_param_df$param)
best_param_df$models <- factor(best_param_df$models)
```

We then illustrate these differences with the following boxplots by parameter and by simulated model.

```
# BCST
true_param_df_BCST <- true_param_df[true_param_df$models == "BCST",]
true_param_df_BCST$selection <- "True"

best_param_df_BCST <- best_param_df[best_param_df$models == "BCST",]
best_param_df_BCST$selection <- "Best"

param_df_BCST <- rbind(true_param_df_BCST, best_param_df_BCST)
param_df_BCST$param <- factor(param_df_BCST$param,
                              levels = c("Lambda", "Alpha", "Mu", "Beta"))

# BVAR
true_param_df_BVAR <- true_param_df[true_param_df$models == "BVAR",]
true_param_df_BVAR$selection <- "True"

best_param_df_BVAR <- best_param_df[best_param_df$models == "BVAR",]
best_param_df_BVAR$selection <- "Best"

param_df_BVAR <- rbind(true_param_df_BVAR, best_param_df_BVAR)
param_df_BVAR$param <- factor(param_df_BVAR$param,
```

```
                              levels = c("Lambda", "Alpha", "Mu", "Beta"))

# BVAR_DCST
true_param_df_BVAR_DCST <- true_param_df[true_param_df$models == "BVAR_DCST",]
true_param_df_BVAR_DCST$selection <- "True"

best_param_df_BVAR_DCST <- best_param_df[best_param_df$models == "BVAR_DCST",]
best_param_df_BVAR_DCST$selection <- "Best"

param_df_BVAR_DCST <- rbind(true_param_df_BVAR_DCST, best_param_df_BVAR_DCST)
param_df_BVAR_DCST$param <- factor(param_df_BVAR_DCST$param,
                                   levels = c("Lambda", "Alpha", "Mu", "Beta"))

# BCST_DVAR
true_param_df_BCST_DVAR <- true_param_df[true_param_df$models == "BCST_DVAR",]
true_param_df_BCST_DVAR$selection <- "True"

best_param_df_BCST_DVAR <- best_param_df[best_param_df$models == "BCST_DVAR",]
best_param_df_BCST_DVAR$selection <- "Best"

param_df_BCST_DVAR <- rbind(true_param_df_BCST_DVAR, best_param_df_BCST_DVAR)
param_df_BCST_DVAR$param <- factor(param_df_BCST_DVAR$param,
                                   levels = c("Lambda", "Alpha", "Mu", "Beta"))

par(mfrow = c(2,2), cex = 0.5)
boxplot(rates ~ selection + param, data = param_df_BCST, main = "BCST",
        at = c(1:2, 4:5, 7:8, 10:11), col = c("darkorange4", "deepskyblue4"),
        names = rep("", 8), xlab = "Parameters", ylab = "Rate values",
        xaxt = "n", ylim = c(-0.4, 0.2), cex.axis = 1, las = 1)
axis(1, at = c(1.5, 4.5, 7.5, 10.5), labels = levels(param_df_BCST$param))
points(c(1.5, 4.5, 7.5, 10.5), param[1, c("lambda", "alpha", "mu", "beta")],
       col = "red", pch = 19)
legend("bottomleft", legend = c("Best", "True", "True values"),
       fill = c("darkorange4", "deepskyblue4", NA), pch = c(NA, NA, 19),
       border = c("black", "black", NA), col = c(NA, NA, "red"),
       bty = "n", cex = 1)

boxplot(rates ~ selection + param, data = param_df_BVAR, main = "BVAR",
        names = rep("", 8), xlab = "Parameters", ylab = "Rate values",
        at = c(1:2, 4:5, 7:8, 10:11), col = c("darkorange4", "deepskyblue4"),
        xaxt = "n", ylim = c(-2, 1.5), cex.axis = 1, las = 1)
axis(1, at = c(1.5, 4.5, 7.5, 10.5), labels = levels(param_df_BCST$param))
points(c(1.5, 4.5, 7.5, 10.5), param[2, c("lambda", "alpha", "mu", "beta")],
       col = "red", pch = 19)

boxplot(rates ~ selection + param, data = param_df_BVAR_DCST, main = "BVAR_DCST",
        at = c(1:2, 4:5, 7:8, 10:11), col = c("darkorange4", "deepskyblue4"),
        names = rep("", 8), xlab = "Parameters", ylab = "Rate values",
        xaxt = "n", ylim = c(-0.1, 0.4), cex.axis = 1, las = 1)
axis(1, at = c(1.5, 4.5, 7.5, 10.5), labels = levels(param_df_BCST$param))
points(c(1.5, 4.5, 7.5, 10.5), param[3, c("lambda", "alpha", "mu", "beta")],
       col = "red", pch = 19)
```

```
boxplot(rates ~ selection + param, data = param_df_BCST_DVAR, main = "BCST_DVAR",
        names = rep("", 8), xlab = "Parameters", ylab = "Rate values",
        at = c(1:2, 4:5, 7:8, 10:11), col = c("darkorange4", "deepskyblue4"),
        xaxt = "n", ylim = c(-0.1, 0.5), cex.axis = 1, las = 1)
axis(1, at = c(1.5, 4.5, 7.5, 10.5), labels = levels(param_df_BCST$param))
points(c(1.5, 4.5, 7.5, 10.5), param[4, c("lambda", "alpha", "mu", "beta")],
       col = "red", pch = 19)
```

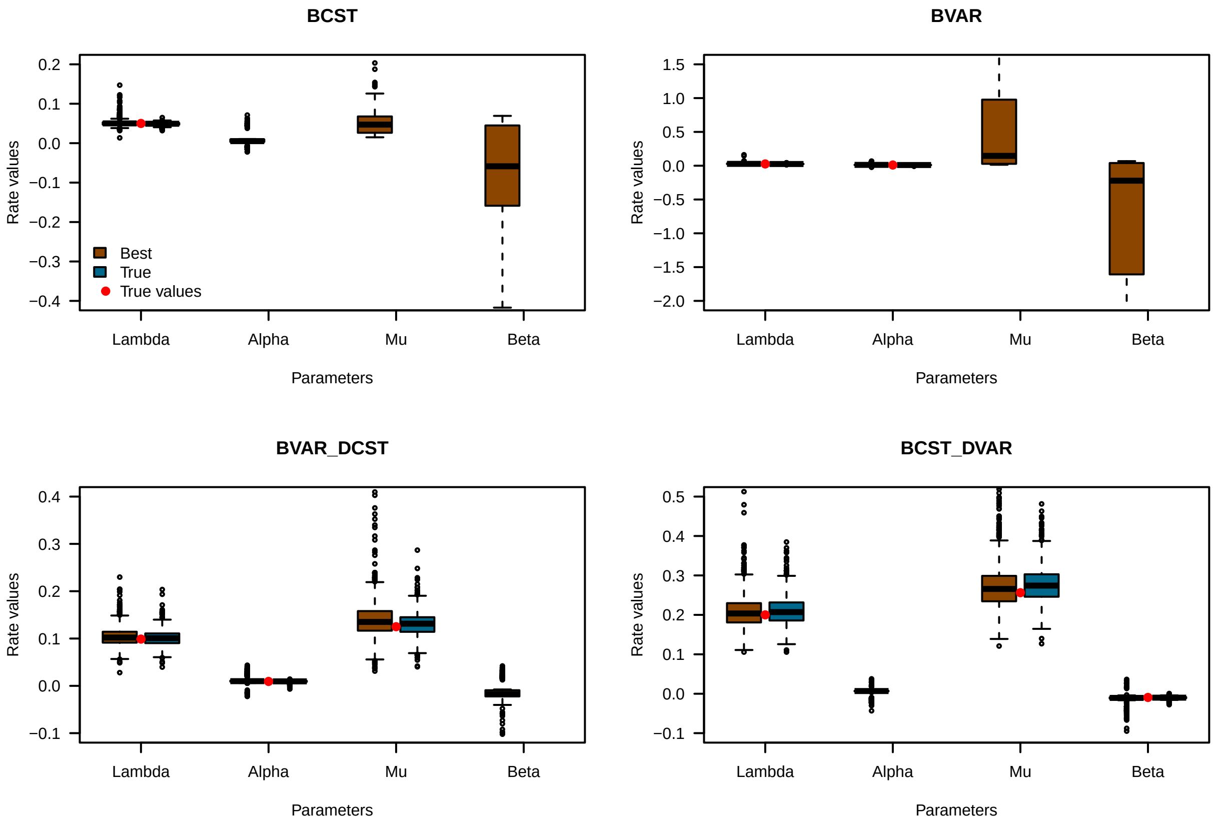

Figure 3: Rate values of best and true models for the backbones

We can see that the ranges values are quite large for the best models when the parameter is not in the true model (when there is a brownish boxplot without a blue boxplot in comparison because the parameter is not included in the true model).

#### 3. Diversity dynamics

After measuring the rate of false detection of shift per models, we wanted to look at the difference between estimated and simulated diversity dynamics. To do so, we calculated the global D error as in Billaud et al. (2020):

$$D = \frac{\Sigma_{t=0}^{T_{max}} |(N_{obs}(t) - N_{th}(t))|/N_{obs}(t)}{T_{max} + 1}$$

where $N_{obs}(t)$ and $N_{th}(t)$ are respectively the estimated and simulated number of species at time $t$ and $T_{max}$ is the total time of the simulation (100 Myrs in our case).

The following code calculates the observed and expected diversity dynamic with the deterministic approach. First on the simulation without shifts:

```
# function for expected paleodiversity dynamics
source("scripts/sim_diversity.R")

all_div_r <- lapply(1:length(models),
                    function(i) lapply(all_tree_r[[i]],
                                       function(x) sim_diversity(phylo =  x,
                                                                 model = models[i],
                                                                 param = param[i,])))
names(all_div_r) <- models

# RAW ####

all_paleodiv_adequacy_r <- rep(list(NULL),4)
all_estim_div_r <- rep(list(NULL),4)
all_simul_div_r <- rep(list(NULL),4)

names <- c("BCST", "BVAR", "BVAR_DCST", "BCST_DVAR")

quartile_names <- c("first quartile", "median", "third quartile")
th = 2

for(res.i in 1:length(all_res_r)){
  cat("Model",res.i,":", names[res.i], "\n")
  # Loop on simulations ####
  paleodiv_adequacy_r <- data.frame(
    Ntip = rep(NA, length(all_tree_r[[res.i]])),
    Ntip_shift = rep(NA, length(all_tree_r[[res.i]])),
    D = rep(NA, length(all_tree_r[[res.i]])),
    model.sub = rep(NA, length(all_tree_r[[res.i]])),
    model.bck = rep(NA, length(all_tree_r[[res.i]])),
    rates.bck = rep(NA, length(all_tree_r[[res.i]])),
    rates.sub = rep(NA, length(all_tree_r[[res.i]])))

  estim_div <- rep(list(NULL), 1000)
  simul_div <- rep(list(NULL), 1000)

  for(tr in 1:length(all_tree_r[[res.i]])){
    cat(paste0("\t",tr,"/",length(all_tree_r[[res.i]])), "\n")
    tree <- all_tree_r[[res.i]][[tr]]
    paleodiv_adequacy_r$Ntip[tr] <- Ntip(tree)

    taxo <- data.frame(Species = tree$tip.label,
                       Genus = ifelse(grepl("t", tree$tip.label), "t", "s"))

    if(all(!grepl("s", tree$tip.label))){

      node <- branching.times(tree)[branching.times(tree) > 25 &
                                      branching.times(tree) < 33]
      node_desc <- sapply(Descendants(tree, as.numeric(names(node))), length)
```

```r
    node <- node[node_desc > 5]
    node <- names(node[node == min(node)])
    node <- as.numeric(node[1])

    sp_shift <- tree$tip.label[unlist(Descendants(tree, node))]
    taxo$Genus[taxo$Species %in% sp_shift] <- "S"

    paleodiv_adequacy_r$Ntip_shift[tr] <- length(sp_shift)

  } else {
    paleodiv_adequacy_r$Ntip_shift[tr] <-
      length(tree$tip.label[
        grepl("s", tree$tip.label)])
  }

  # significant only
  if(all_res_r[[res.i]][[tr]]$total$Combination[1] == "whole_tree" |
     all_res_r[[res.i]][[tr]]$total$delta_AICc[2] < th){

    model_whole <- all_res_r[[res.i]][[tr]]$whole_tree$Models[
      all_res_r[[res.i]][[tr]]$whole_tree$AICc ==
        min(all_res_r[[res.i]][[tr]]$whole_tree$AICc)]
    paleodiv_adequacy_r$model.bck[tr] <- model_whole
    paleodiv_adequacy_r$rates.bck[tr] <- paste(all_res_r[[res.i]][[tr]]$whole_tree[
      all_res_r[[res.i]][[tr]]$whole_tree$AICc ==
        min(all_res_r[[res.i]][[tr]]$whole_tree$AICc),
      c("Lambda", "Alpha", 'Mu', "Beta")], collapse = "/")
    combi <- which(all_res_r[[res.i]][[tr]]$total$Combination == "whole_tree")
  } else {

    paleodiv_adequacy_r$model.bck[tr] <-
      all_res_r[[res.i]][[tr]]$backbones[[1]][[1]]$Models[1]
    paleodiv_adequacy_r$model.sub[tr] <-
      all_res_r[[res.i]][[tr]]$subclades[[1]]$Models[1]

    rates.bck <- paste(all_res_r[[res.i]][[tr]]$backbones[[1]][[1]][
      1, c("Lambda", "Alpha", 'Mu', "Beta")], collapse = "/")
    paleodiv_adequacy_r$rates.bck[tr] <- rates.bck

    rates.sub <- paste(all_res_r[[res.i]][[tr]]$subclades[[1]][
      1, c("Lambda", "Alpha", 'Mu', "Beta")], collapse = "/")
    paleodiv_adequacy_r$rates.sub[tr] <- rates.sub

    combi <- which(all_res_r[[res.i]][[tr]]$total$Combination != "whole_tree")
  }

  f.df <- get.sampling.fractions(phylo = tree, data = taxo, clade.size = 5)

  # Estimated diversity
  diversity <- paleodiv(phylo = tree, data = taxo, sampling.fractions = f.df,
                        shift.res = all_res_r[[res.i]][[tr]], combi = combi)

  estim_div[[tr]] <- diversity
```

```
    # Simulated diversity
    sim_div <- sim_diversity(phylo = tree,
                             model = models[res.i],
                             param = param[res.i,])
    simul_div[[tr]] <- sim_div

    # Global error D
    paleodiv_adequacy_r$D[tr] <- sum(abs(diversity - sim_div)/
                                     diversity)/length(diversity)

  }
  # three quartiles by model ####
  paleodiv_adequacy_r$D <- round(paleodiv_adequacy_r$D, 3)

  all_paleodiv_adequacy_r[[res.i]] <- paleodiv_adequacy_r

  names(simul_div) <- names(estim_div) <- names(all_res_r[[res.i]])

  all_simul_div_r[res.i] <- list(simul_div)
  all_estim_div_r[res.i] <- list(estim_div)
}
names(all_estim_div_r) <- models
names(all_simul_div_r) <- models

save(all_paleodiv_adequacy_r, all_estim_div_r, all_simul_div_r,
     file = "./simul_res_raw.RData")
```

Then on simulations with a shift:

```
all_div_s <- lapply(1:length(models),
                    function(i) lapply(all_tree_s[[i]],
                                       function(x) sim_diversity(phylo =  x,
                                                                 model = models[i],
                                                                 param = param[i,])))
names(all_div_s) <- models

# SHIFT ####
all_paleodiv_adequacy_s <- rep(list(NULL),4)
all_simul_div_s <- rep(list(NULL),4)
all_estim_div_s <- rep(list(NULL),4)

for(res.i in 1:length(all_res_s)){
  cat("Model",res.i,":", models[res.i], "\n")
  # Loop on simulations ####
  paleodiv_adequacy_s <- data.frame(
    Ntip = rep(NA, length(all_tree_s[[res.i]])),
    Ntip_shift = rep(NA, length(all_tree_s[[res.i]])),
    D = rep(NA, length(all_tree_s[[res.i]])),
    model.sub = rep(NA, length(all_tree_s[[res.i]])),
    model.bck = rep(NA, length(all_tree_s[[res.i]])),
    rates.bck = rep(NA, length(all_tree_s[[res.i]])),
    rates.sub = rep(NA, length(all_tree_s[[res.i]])))
```

```
estim_div <- rep(list(NULL), 1000)
simul_div <- rep(list(NULL), 1000)

for(tr in 1:length(all_tree_s[[res.i]])){
  cat(paste0("\t",tr,"/",length(all_tree_s[[res.i]])), "\n")
  tree <- all_tree_s[[res.i]][[tr]]
  paleodiv_adequacy_s$Ntip[tr] <- Ntip(tree)

  taxo <- data.frame(Species = tree$tip.label,
                     Genus = ifelse(grepl("t", tree$tip.label), "t", "s"))

  if(all(!grepl("s", tree$tip.label))){

    node <- branching.times(tree)[branching.times(tree) > 25 &
                                  branching.times(tree) < 33]
    node <- as.numeric(names(node[node == min(node)]))
    sp_shift <- tree$tip.label[unlist(Descendants(tree, node))]
    taxo$Genus[taxo$Species %in% sp_shift] <- "S"

    paleodiv_adequacy_s$Ntip_shift[tr] <- length(sp_shift)

  } else {
    paleodiv_adequacy_s$Ntip_shift[tr] <- length(tree$tip.label[grepl("s",
                                                                  tree$tip.label)])
  }

  # significant only
  if(all_res_s[[res.i]][[tr]]$total$Combination[1] == "whole_tree" |
     all_res_s[[res.i]][[tr]]$total$delta_AICc[2] < th){

    model_whole <- all_res_s[[res.i]][[tr]]$whole_tree$Models[
      all_res_s[[res.i]][[tr]]$whole_tree$AICc ==
        min(all_res_s[[res.i]][[tr]]$whole_tree$AICc)]
    paleodiv_adequacy_s$model.bck[tr] <- model_whole
    paleodiv_adequacy_s$rates.bck[tr] <- paste(all_res_s[[res.i]][[tr]]$whole_tree[
      all_res_s[[res.i]][[tr]]$whole_tree$AICc ==
        min(all_res_s[[res.i]][[tr]]$whole_tree$AICc),
      c("Lambda", "Alpha", 'Mu', "Beta")], collapse = "/")
    combi <- which(all_res_r[[res.i]][[tr]]$total$Combination == "whole_tree")
  } else {

    paleodiv_adequacy_s$model.bck[tr] <-
      all_res_s[[res.i]][[tr]]$backbones[[1]][[1]]$Models[1]
    paleodiv_adequacy_s$model.sub[tr] <-
      all_res_s[[res.i]][[tr]]$subclades[[1]]$Models[1]

    rates.bck <- paste(all_res_s[[res.i]][[tr]]$backbones[[1]][[1]][
      1,
      c("Lambda", "Alpha", 'Mu', "Beta")], collapse = "/")
    paleodiv_adequacy_s$rates.bck[tr] <- rates.bck

    rates.sub <- paste(all_res_s[[res.i]][[tr]]$subclades[[1]][
      1,
```

```
        c("Lambda", "Alpha", 'Mu', "Beta")],
        collapse = "/")
      paleodiv_adequacy_s$rates.sub[tr] <- rates.sub
      combi <- which(all_res_r[[res.i]][[tr]]$total$Combination != "whole_tree")
    }

    f.df <- get.sampling.fractions(phy = tree, data = taxo, clade.size = 5)

    # Estimated diversity
    diversity <- paleodiv(phy = tree, data = taxo, sampling.fractions = f.df,
                          shift.res = all_res_s[[res.i]][[tr]], combi = combi)

    estim_div[[tr]] <- diversity
    # Simulated diversity
    sim_div <- sim_diversity(phy = tree, model = names[res.i], param = param)
    simul_div[[tr]] <- sim_div
    # Global error D
    paleodiv_adequacy_s$D[tr] <- sum(abs(diversity - sim_div)/diversity)/length(diversity)

  }
  # three quartile by model ####
  paleodiv_adequacy_s$D <- round(paleodiv_adequacy_s$D, 3)

  all_paleodiv_adequacy_s[[res.i]] <- paleodiv_adequacy_s
  names(simul_div) <- names(estim_div) <- names(all_res_s[[res.i]])

  all_simul_div_s[res.i] <- list(simul_div)
  all_estim_div_s[res.i] <- list(estim_div)

}
names(all_simul_div_s) <- models
names(all_estim_div_s) <- models

save(all_paleodiv_adequacy_s, all_estim_div_s, all_simul_div_s,
     file = "./simul_res_shift.RData")

load("./data/simul_res_raw.RData")
```

We can now merge all simulations to represent the observed and expected paleodiversity dynamics for the three quartiles of $D$ values for the four models as the Figure 4.

Here are the codes to merge the simulations with and without shift:

```
all_paleodiv_adequacy_r.df <-  do.call(rbind, all_paleodiv_adequacy_r)
all_paleodiv_adequacy_r.df <- as.data.frame(all_paleodiv_adequacy_r.df)
all_paleodiv_adequacy_r.df$general.model <- rep(c("Model1", "Model2", "Model3", "Model4"),
                                                each = 1000)
all_paleodiv_adequacy_r.df$simulated_shift <- rep(FALSE, nrow(all_paleodiv_adequacy_r.df))
all_paleodiv_adequacy_r.df$FP <- ifelse(!is.na(all_paleodiv_adequacy_r.df$model.sub),
                                        T, F)
all_paleodiv_adequacy_r.df$FN <- F

all_paleodiv_adequacy_s.df <-  do.call(rbind, all_paleodiv_adequacy_s)
all_paleodiv_adequacy_s.df <- as.data.frame(all_paleodiv_adequacy_s.df)
```

```
all_paleodiv_adequacy_s.df$general.model <- rep(c("Model1", "Model2", "Model3", "Model4"),
                                                 each = 1000)
all_paleodiv_adequacy_s.df$simulated_shift <- rep(TRUE, nrow(all_paleodiv_adequacy_s.df))
all_paleodiv_adequacy_s.df$FN <- ifelse(is.na(all_paleodiv_adequacy_s.df$model.sub),
                                        T, F)
all_paleodiv_adequacy_s.df$FP <- F

all_adequacy <- as.data.frame(rbind(all_paleodiv_adequacy_r.df,
                                    all_paleodiv_adequacy_s.df))
all_adequacy$general.model <- factor(all_adequacy$general.model)

names(all_simul_div_r) <- models

all_simul_paleodiv <- c(do.call(c, all_simul_div_r), do.call(c, all_simul_div_s))
names(all_simul_paleodiv) <- gsub("[.]", "_n", names(all_simul_paleodiv))

all_estim_paleodiv <- c(do.call(c, all_estim_div_r), do.call(c, all_estim_div_s))
names(all_estim_paleodiv) <- gsub("[.]", "_n", names(all_estim_paleodiv))

all_trees <- c(do.call(c, all_tree_r), do.call(c, all_tree_s))
names(all_trees) <- paste0(rep(rep(paste0(names, "_n"), each = 1000),2),
                           names(all_trees))
```

And here are the lines to reproduce the Figure 4 of the main manuscript (here Figure 3):

```
par(mfrow = c(4,3), oma = c(2,6,3,2), mar = c(3,3,3,1), cex = 0.7, cex.main = 1)
no_decline <- function(x){
  dif <- c()
  for(i in 2:length(x)-1){
    dif <- c(dif, x[i+1] - x[i] > 0)
  }
  return(all(dif))
}

no_decline_check <- c()
for(mod in 1:nlevels(all_adequacy$general.model)){

  df <- all_adequacy[all_adequacy$general.model ==
                     levels(all_adequacy$general.model)[mod],]
  D_quant <- quantile(df$D, probs = c(0.25, 0.5, 0.75))

  trees <- all_trees[grepl(paste0(models[mod],"_n"), names(all_trees))]
  diversities <- all_estim_paleodiv[grepl(paste0(models[mod],"_n"),
                                         names(all_estim_paleodiv))]
  sim_diversities <- all_simul_paleodiv[grepl(paste0(models[mod],"_n"),
                                             names(all_simul_paleodiv))]

  # decline
  no_decline_check[mod] <- sum(sapply(diversities, no_decline))/2000

  which_quant <- c()
  for(i in 1:length(D_quant)){
    which_quant <- c(which_quant,
```

```
                    which(df$D == min(df$D[round(df$D,2) %in%
                                             round(D_quant[i], 2)]))[1])
  }

  for(tr in which_quant){
    # Estimated diversity
    diversity <- diversities[[tr]]
    # Simulated diversity
    sim_div <- sim_diversities[[tr]]

    t <- seq(0, 100,1)
    plot(t, diversity, type = "l", lwd = 3, xaxt = "n",
         ylim = c(0, max(diversity, sim_div)+1),
         col = "darkorchid2",
         main = paste0(quartile_names[which(which_quant == tr)], "\n D = ",
                       df$D[tr], 3), las = 1)
    axis(side = 1, las = 1, line = 0, xpd = TRUE,
         labels = -rev(seq(t[1], t[length(t)], 5)), at = seq(t[1], t[length(t)], 5),
         cex.axis = 0.8, padj = -0.5)
    mtext(text = "Time (Myrs)", side = 1, line = 2, cex = 0.5)
    mtext(text = "Number of species (log)", side = 2, line = 2.4, cex = 0.5)
    lines(t, sim_div, col = "chartreuse3", lwd = 3, lty = 1)
    legend("topleft", legend = c("Inferred", "Simulated"), lty = 1,
           col = c("darkorchid2", "chartreuse3"), bty = "n", lwd = 3, cex = 0.8)
  }
}
mtext(text = gsub("_", " ", rev(models)), at = c(0.12, 0.35, 0.62, 0.85),
      side = 2, line = 2, outer = T, cex = 0.8)
mtext(text = "Paleodiversity dynamics", side = 3, line = 0.5,
      outer = T, cex = 1)
```

At first glance, the diversity curves might seem to be overestimated. We looked more into details (1) at the potential bias between the simulated and the expected curves at (2) the peak and at the largest difference of the curves.

```
overestimation <- lapply(1:length(all_estim_div_s), function(i){
  sapply(1:length(all_estim_div_s$BCST), function(j){
    sum(all_estim_div_s[[i]][[j]] > all_simul_div_s[[i]][[j]])/101
  })
})
names(overestimation) <- names(all_estim_div_s)
sapply(overestimation, summary)
```

```
##              BCST      BVAR BVAR_DCST BCST_DVAR
## Min.    0.0000000 0.0000000 0.0000000 0.0000000
## 1st Qu. 0.2871287 0.4257426 0.2277228 0.3465347
## Median  0.7623762 0.8564356 0.6039604 0.7029703
## Mean    0.6296436 0.7030198 0.5811188 0.6290891
## 3rd Qu. 0.9900990 0.9900990 0.9900990 0.9727723
## Max.    0.9900990 0.9900990 0.9900990 0.9900990
```

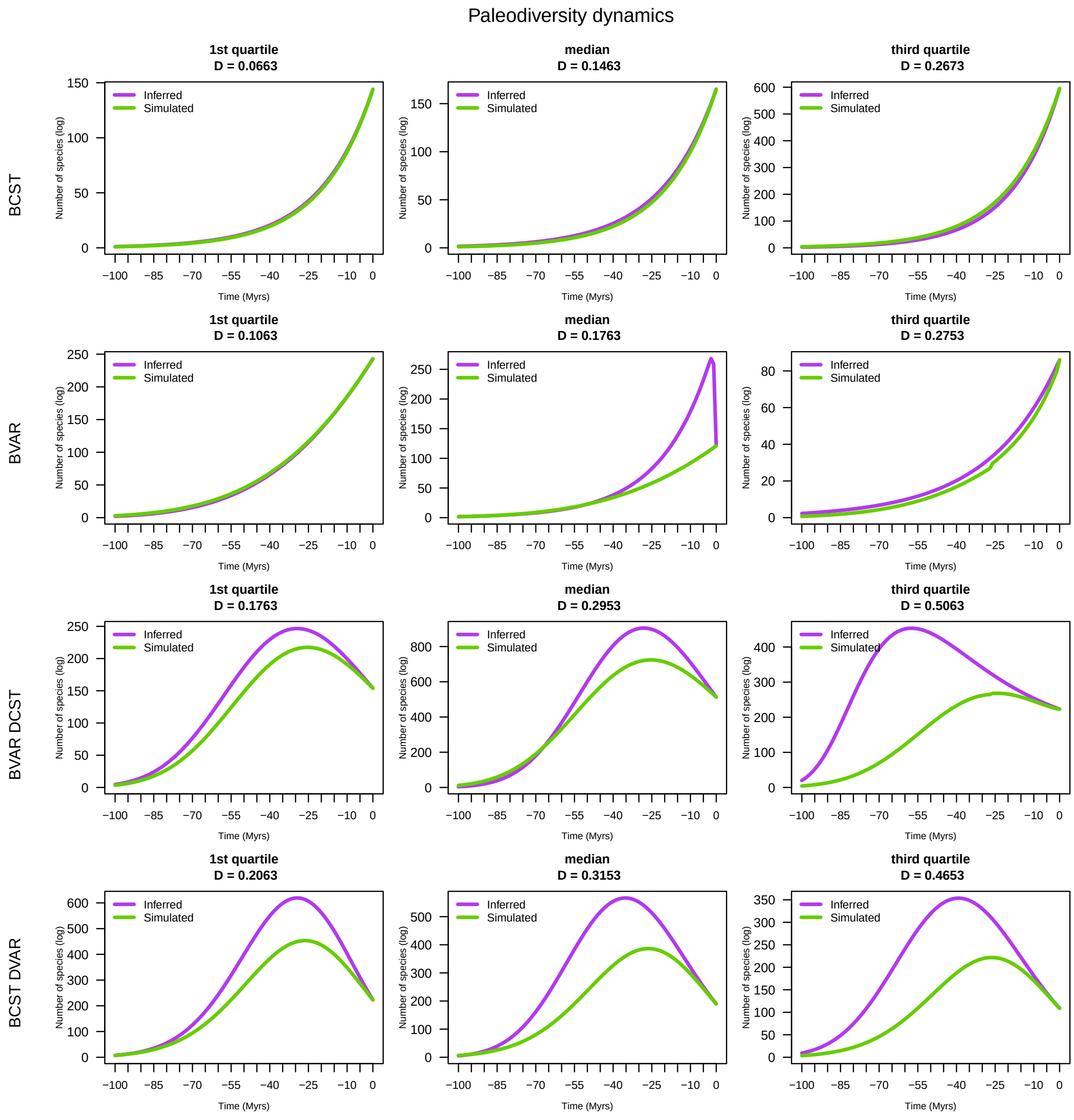

Figure 4: Comparison of inferred versus simulated paleodiversity dynamics for the three quartiles of global error D by model

```r
# the peak
# simulation without shift
peak_overestimation <- lapply(1:length(all_estim_div_r), function(i){
  sapply(1:length(all_estim_div_r$BCST), function(j){
    max(all_estim_div_r[[i]][[j]]) > max(all_simul_div_r[[i]][[j]])
  })
})
names(peak_overestimation) <- names(all_estim_div_r)
peak_sup <- sapply(peak_overestimation, sum)

peak_good_estimation <- lapply(1:length(all_estim_div_r), function(i){
  sapply(1:length(all_estim_div_r$BCST), function(j){
    max(all_estim_div_r[[i]][[j]]) == max(all_simul_div_r[[i]][[j]])
  })
})
names(peak_good_estimation) <- names(all_estim_div_r)
peak_equal <- sapply(peak_good_estimation, sum)

peak_underestimation <- lapply(1:length(all_estim_div_r), function(i){
  sapply(1:length(all_estim_div_r$BCST), function(j){
    max(all_estim_div_r[[i]][[j]]) < max(all_simul_div_r[[i]][[j]])
  })
})
names(peak_underestimation) <- names(all_estim_div_r)
peak_inf <- sapply(peak_underestimation, sum)
peak_r <- rbind(peak_inf, peak_equal, peak_sup)
apply(peak_r, 2, sum)
```

```
##      BCST      BVAR BVAR_DCST BCST_DVAR
##      1000      1000      1000      1000
```

```r
# simulation with a shift
peak_overestimation <- lapply(1:length(all_estim_div_s), function(i){
  sapply(1:length(all_estim_div_s$BCST), function(j){
    max(all_estim_div_s[[i]][[j]]) > max(all_simul_div_s[[i]][[j]])
  })
})
names(peak_overestimation) <- names(all_estim_div_s)
peak_sup <- sapply(peak_overestimation, sum)

peak_good_estimation <- lapply(1:length(all_estim_div_s), function(i){
  sapply(1:length(all_estim_div_s$BCST), function(j){
    max(all_estim_div_s[[i]][[j]]) == max(all_simul_div_s[[i]][[j]])
  })
})
names(peak_good_estimation) <- names(all_estim_div_s)
peak_equal <- sapply(peak_good_estimation, sum)

peak_underestimation <- lapply(1:length(all_estim_div_s), function(i){
  sapply(1:length(all_estim_div_s$BCST), function(j){
    max(all_estim_div_s[[i]][[j]]) < max(all_simul_div_s[[i]][[j]])
  })
})
```

```
names(peak_underestimation) <- names(all_estim_div_s)
peak_inf <- sapply(peak_underestimation, sum)
peak_s <- rbind(peak_inf, peak_equal, peak_sup)
apply(peak_s, 2, sum)
```

```
##      BCST      BVAR BVAR_DCST BCST_DVAR
##      1000      1000      1000      1000
```

```
peak <- peak_r + peak_s
peak <- peak/2000

# largest difference
# raw
biais_r <- lapply(1:length(all_estim_div_r), function(i){
  sapply(1:length(all_estim_div_r$BCST), function(j){
    res <- all_estim_div_r[[i]][[j]] - all_simul_div_r[[i]][[j]]
    res <- ifelse(max(abs(res)) %in% res, max(abs(res)), -max(abs(res)))
    return(res)
  })
})
names(biais_r) <- names(all_estim_div_r)

biais_r_inf <- sapply(biais_r, function(x) sum(x < 0))
biais_r_eq <- sapply(biais_r, function(x) sum(x == 0))
biais_r_sup <- sapply(biais_r, function(x) sum(x > 0))

biais_r <- rbind(biais_r_inf, biais_r_eq, biais_r_sup)

# shift
biais_s <- lapply(1:length(all_estim_div_s), function(i){
  sapply(1:length(all_estim_div_s$BCST), function(j){
    res <- all_estim_div_s[[i]][[j]] - all_simul_div_s[[i]][[j]]
    res <- ifelse(max(abs(res)) %in% res, max(abs(res)), -max(abs(res)))
    return(res)
  })
})
names(biais_s) <- names(all_estim_div_s)

biais_s_inf <- sapply(biais_s, function(x) sum(x < 0))
biais_s_eq <- sapply(biais_s, function(x) sum(x == 0))
biais_s_sup <- sapply(biais_s, function(x) sum(x > 0))

biais_s <- rbind(biais_s_inf, biais_s_eq, biais_s_sup)

biais <- biais_r + biais_s
biais <- biais/2000
```

Here the code for representing the results by models

```
par(mfrow = c(1,2), mar = c(4,7,4,2), oma = c(0,0,0,4), xpd = F)
barplot(as.matrix(peak),
        col = c("dodgerblue", "grey", "darkred"), las = 1,
```

```
        main = "Bias at the peak", ylab = "Proportions",
        cex.names = 0.6)
par(mar = c(4,2,4,7))
barplot(as.matrix(biais),
        col = c("dodgerblue", "grey", "darkred"), las = 1,
        main = "Bias at the highest difference", ylab = "Proportions",
        cex.names = 0.6)
par(xpd = T)
legend(4.8, 0.6, legend = c("overestimed", "same value","underestimated"),
       fill = c("darkred", "grey", "dodgerblue"), bty = "n", cex=0.8, xpd = T)
```

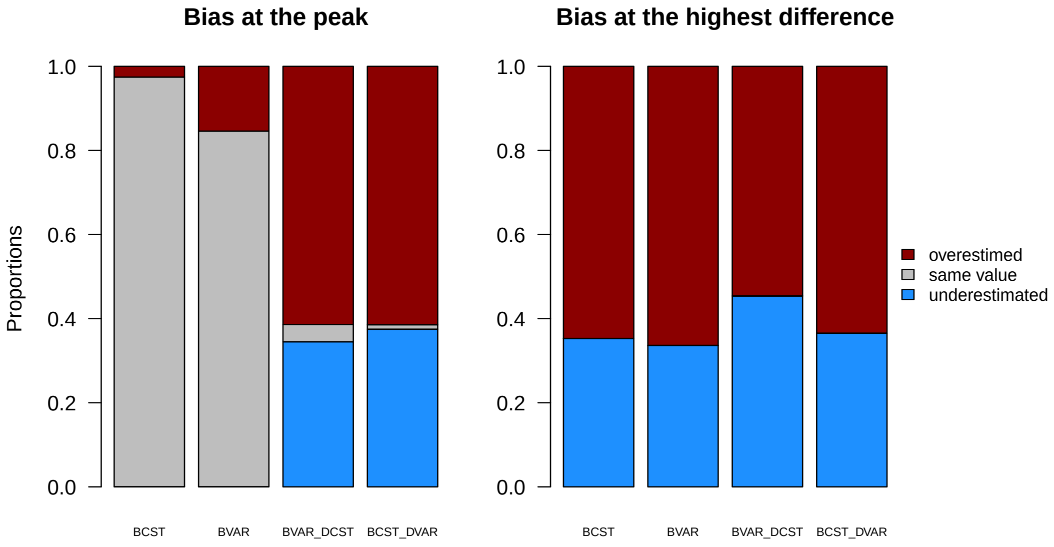

Figure 5: Comparison of inferred versus simulated paleodiversity dynamics for the three quartiles of global error D by model

We can see that the paleodiversity is well estimated at the peak for simpler models and not always overestimated, with almost 40% of underestimation for both most complex models at the peak and on average for bias at the highest difference.

#### 4. Extinction in subclades

The method we improved and implemented into the RPANDA package has been developed to better estimate extinction rate in deep parts of phylogenies. In the simulations, we decided to simulate the shift of diversification with a pure-birth model (BCST). One might be interested in testing whether extinction rate is properly detected and estimated in subclades.

In order to do so, we led a small set of simulations (500 trees) following the same pipeline with the Model 3 (BVAR DCST) but with a shift simulated by grafting a subclade simulated by a birth-death model (BCST_DCST) instead of a pure-birth model (BCST). All the other criteria of simulations were the same. We chose to keep the same net diversification rate with a speciation rate equal to 0.2 and an extinction rate equal to 0.1.

The simulations can be done by launching the Snakefile `Snakefile_comp_BCST_DCST` as the other Snakefiles.

```
# in a terminal
conda activate simshift_env
snakemake -s Snakefile_comp_BCST_DCST -j 30
```

After downloading both the trees and the results we can have a look at the detection of the shift and see how many times the shift is detected.

```
names_trees_BCST_DCST <- list.files("data/trees", pattern = "BCST_DCST.t", full.names = T)
names_res_BCST_DCST <- list.files("data/res", pattern = "BCST_DCST.r", full.names = T)

trees_BCST_DCST <- lapply(names_trees_BCST_DCST, read.tree)
res_BCST_DCST <- lapply(names_res_BCST_DCST, readRDS)

res_BCST_DCST_noBCST <- lapply(res_BCST_DCST, remove.model, "BCST")
res_BCST_DCST_noBCST <- lapply(res_BCST_DCST_noBCST, remove.model, "BVAR")

best_model <- list()
shift <- c()
lambda_BCST_DCST <- c()
mu_BCST_DCST <- c()
lambda_BCST <- c()
for(nres in 1:length(res_BCST_DCST)){
  best_model[[nres]] <- res_BCST_DCST[[nres]]$subclades[[1]][
    res_BCST_DCST[[nres]]$subclades[[1]]$delta_AICc < 2,]
  shift <- c(shift, ifelse(res_BCST_DCST[[nres]]$total$delta_AICc[2] > 2,
                           res_BCST_DCST[[nres]]$total$Combination[1],
                           "no_shift"))
  lambda_BCST_DCST <- c(lambda_BCST_DCST, res_BCST_DCST[[nres]]$subclades[[1]][
    res_BCST_DCST[[nres]]$subclades[[1]]$Models == "BCST_DCST", c("Lambda")])
  mu_BCST_DCST <- c(mu_BCST_DCST, res_BCST_DCST[[nres]]$subclades[[1]][
    res_BCST_DCST[[nres]]$subclades[[1]]$Models == "BCST_DCST", c("Mu")])
  lambda_BCST <- c(lambda_BCST, res_BCST_DCST[[nres]]$subclades[[1]][
    res_BCST_DCST[[nres]]$subclades[[1]]$Models == "BCST", c("Lambda")])
}
```

```
sum(sapply(res_BCST_DCST, function(x)
  x$total$Combination[1] != "whole_tree" & x$total$delta_AICc[2] > 2))/500
```

As we can see, the simulated shift is well recovered.

```
model_recovery <- table(sapply(best_model, function(x) x$Models[1]))/500
par(mfrow = c(1,2))
a <- barplot(model_recovery, cex.names = 0.45,
             ylim = c(0,1), las = 2, xlab = "Models", ylab = "Recovery rate",
             main = "A) Best models for simulations\nof BCST_DCST model")
text(a[,1], table(sapply(best_model, function(x) x$Models[1]))/500+0.03,
     labels = as.character(table(sapply(best_model, function(x) x$Models[1]))/500))

boxplot(lambda_BCST_DCST, mu_BCST_DCST,
        lambda_BCST,
        main = "B) Estimated rate values",
        ylab = "Rate values",
```

```
         names = c("lambda", "mu", "lambda (=r)"),
         col = c("darkorange", "darkorange", "gold"), las = 1)
points(c(1:3), c(0.2, 0.1, 0.1), col = "red", pch = 19,
       cex = 1)
legend("topright", legend = c("BCST_DCST", "BCST", "Simulated values"),
       fill= c("darkorange", "gold", NA), pch = c(NA, NA, 19),
       border = c("black", "black", NA), col = c(NA,NA,"red"), bty = "n")
```

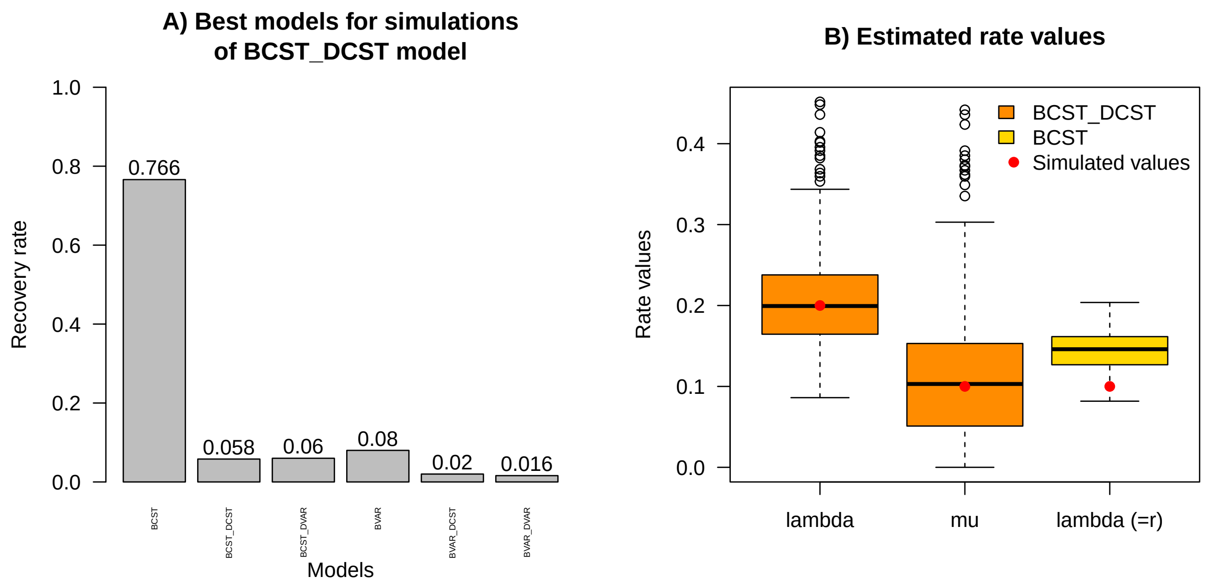

Figure 6: Preliminary results on simulations with a subclade simulated under a constant birth-death model

Unfortunately, the BCST_DCST model is not recovered for the subclades in most of the case (only 5.8%).

Instead, the BCST model is recovered in 76.6% of the cases (Figure 6A). If we look at the rate estimates, the BCST_DCST model estimated rates more accurately than the BCST model (Figure 6B, orange). However, speciation rate estimated with the BCST model (Figure 6B, yellow) is underestimated. This is a preliminary result that requires further studies to better understand this pattern. We hypothesize that it can be caused by a higher pull of the present effect because the net diversification rate (r = $\lambda$ - $\mu$) is overestimated with the BCST model (r = $\lambda$).
