## Appendix S4 for "Estimating clade-specific diversification rates and palaeodiversity dynamics from reconstructed phylogenies"

### Appendix S4: Empirical cases

Estimating clade-specific diversification rates and paleodiversity dynamics from reconstructed phylogenies

Mazet Nathan, Morlon Hélène, Fabre Pierre-Henri, Condamine Fabien

2023-06-30

#### Contents

This document reproduces the empirical analyses of Mazet *et al.* (in prep) to illustrate how to use the automation of the method developed by Morlon *et al.* (2011) and improved by Billaud *et al.* (2020). The first step consists in cleaning the taxonomic databases and the species names in the phylogeny. Then, the second step shows how the analyses have been made on each of the four groups. The third and final step details how to make the same plots as in the manuscript.

#### Loading packages

In this part, we install different packages we need. For RPANDA, you can either install the version from the GitHub repository of RPANDA with its specific branch (named clade.shift.model) or used the archive file available in the folder (RPANDA_2.2_clade.shift.model_300623.tar.gz).

```
# set directory
setwd("./empirical_cases/")

# installing RPANDA from github with devtools
install.packages("devtools")
devtools::install_github("https://github.com/hmorlon/PANDA.git",
                         ref = "clade.shift.model")

# installing RPANDA from the archive file
install.packages(pkgs = "../RPANDA_2.2_clade.shift.model_300623.tar.gz",
                 repos = NULL)

# Packages ####
pkgs <- c("ape", "phytools", "phangorn","RPANDA", "Hmisc", "treebalance",
          "parallel", "ParallelLogger", "RColorBrewer", "devtools")
# Packages not installed yet
new.pkgs <- pkgs[!(pkgs %in% installed.packages()[,"Package"])]
# Installing and loading each package
if(length(new.pkgs) != 0){
  for(i in 1:length(new.pkgs)){
    install.packages(new.pkgs[i], dependencies = T)
  }
}
sapply(pkgs, require, character.only = T)
```

#### I - Cleaning data

The data for the Cetacea and the Vangidae were already cleaned so we used data from the related papers (Steeman *et al.* 2009 for Cetacea and Jønsson *et al.* 2012 for Vangidae).

##### 1. Parnassiinae

The raw data comes from Condamine et al. (2018). We cleaned the taxonomic database with more recent names according to Nakae (2021).

```
tree_parnassiinae <- read.tree("data/raw/Parnassiinae_MolMorpho_BD.tre")
taxo_parnassiinae <- read.csv("data/raw/taxo_parnassiinae.csv")
taxo_parnassiinae$spe_suffixe <- taxo_parnassiinae$Species
taxo_parnassiinae$Species <- paste(taxo_parnassiinae$Genus,
```

```r
                                    taxo_parnassiinae$spe_suffixe, sep = "_")
taxo_parnassiinae$Species[taxo_parnassiinae$Species %in%
                            taxo_parnassiinae$Name.to.use.in.trees]

matching_names <- data.frame(tree_names = tree_parnassiinae$tip.label,
                             taxo_names = rep(NA, length(tree_parnassiinae$tip.label)))
matching_names$tree_names <- sort(matching_names$tree_names)

tip_label_with_subgenus <-
  matching_names$tree_names[sapply(strsplit(matching_names$tree_names,
                                             split = "_"),
                                    length) >= 3]
tip_label_without_subgenus <- sapply(strsplit(tip_label_with_subgenus, "_"),
                                      function(x){
  if(length(x[-2][!x[-2] %in% as.character(1:5)]) < 2){
    name <- paste(x, collapse = "_")
  } else {
    name <- paste(x[-2], collapse = "_")
  }
  return(name)
})

matching_names$taxo_names[matching_names$tree_names %in%
                            tip_label_with_subgenus] <- tip_label_without_subgenus

tip_label_without_subgenus[
  tip_label_without_subgenus %in% taxo_parnassiinae$Species]

# Nazari & Sperling 2007
matching_names$taxo_names[
  matching_names$tree_names == "Archon_apollinus_1"] <- "Archon_apollinus"

matching_names$taxo_names[
  matching_names$tree_names == "Archon_apollinus_2"] <- "Archon_bellargus"

matching_names$taxo_names[
  matching_names$tree_names == "Hypermnestra_helios_1"] <- "Hypermnestra_helios"

matching_names$taxo_names[
  matching_names$tree_names == "Hypermnestra_helios_2"] <- "Hypermnestra_persica"

# Weiss J.C. 1991-2005
matching_names$taxo_names[
  matching_names$tree_names == "Parnassius_staudingeri_1"] <- "Parnassius_staudingeri"

matching_names$taxo_names[
  matching_names$tree_names == "Parnassius_staudingeri_2"] <- "Parnassius_infernalis"

matching_names$taxo_names[
  matching_names$tree_names == "Parnassius_staudingeri_3"] <- "Parnassius_hunza"

# Dapporto 2010
matching_names$taxo_names[
```

```r
  matching_names$tree_names == "Zerynthia_polyxena_1"] <- "Zerynthia_polyxena"

matching_names$taxo_names[
  matching_names$tree_names == "Zerynthia_polyxena_2"] <- "Zerynthia_cassandra"

# Leraut 2016
matching_names$taxo_names[
  matching_names$tree_names == "Zerynthia_rumina_1"] <- "Zerynthia_rumina"

matching_names$taxo_names[
  matching_names$tree_names == "Zerynthia_rumina_2"] <- "Zerynthia_africana"

species_in_taxo <- taxo_parnassiinae$Species[
  taxo_parnassiinae$Species %in%
    matching_names$tree_names[is.na(matching_names$taxo_names)]]
species_in_taxo <- sort(species_in_taxo)

matching_names$taxo_names[
  which(matching_names$tree_names %in% species_in_taxo)] <- species_in_taxo

matching_names$taxo_names[
  matching_names$tree_names == "Luehdorfia_taibai"] <- "Luehdorfia_longicaudata"

matching_names$taxo_names[
  matching_names$tree_names == "Allancastria_cerisyi"] <- "Allancastria_cerisyi"

taxo_parnassiinae$Species[
  taxo_parnassiinae$Species == "Allancastria_cerisy"] <- "Allancastria_cerisyi"

tree_parnassiinae$tip.label[
  tree_parnassiinae$tip.label == "Allancastria_cerisy"] <- "Allancastria_cerisyi"

taxo_parnassiinae$spe_suffixe[
  taxo_parnassiinae$spe_suffixe == "cerisy"] <- "cerisyi"

matching_names$taxo_names[
  !matching_names$taxo_names %in% taxo_parnassiinae$Species]

taxo_parnassiinae$Species[
  !taxo_parnassiinae$Species %in% matching_names$taxo_names]

# from the list given by Fabien
sp_in_tree_no_name <- matching_names$taxo_names[
  !matching_names$taxo_names %in% taxo_parnassiinae$Species]

matching_names$taxo_names[
  matching_names$taxo_names == "Parnassius_mnemosyne_1"] <- "Parnassius_mnemosyne"

matching_names$taxo_names[
  matching_names$taxo_names == "Parnassius_mnemosyne_2"] <- "Parnassius_turatii"

# new name not published yet because uncertain
matching_names$taxo_names[
```

```r
  matching_names$taxo_names == "Parnassius_mnemosyne_3"] <- "Parnassius_mnemosyne_3"

# nothing changes yet
matching_names$taxo_names[
  matching_names$taxo_names == "Parnassius_augustus"] <- "Parnassius_augustus"

matching_names$taxo_names[
  matching_names$taxo_names == "Parnassius_apollo_1"] <- "Parnassius_apollo"

matching_names$taxo_names[
  matching_names$taxo_names == "Parnassius_apollo_2"] <- "Parnassius_apollo_2"

matching_names$taxo_names[
  matching_names$taxo_names == "Parnassius_apollo_3"] <- "Parnassius_apollo_3"

matching_names$taxo_names[
  matching_names$taxo_names == "Parnassius_apollo_4"] <- "Parnassius_apollo_4"

matching_names$taxo_names[
  matching_names$taxo_names == "Parnassius_apollo_5"] <- "Parnassius_apollo_5"
# species suffix changed
matching_names$taxo_names[
  matching_names$taxo_names == "Parnassius_stubbendorfi"] <- "Parnassius_stubbendorfii"

matching_names$taxo_names[
  matching_names$taxo_names == "Parnassius_arcticus"] <- "Parnassius_arctica"

# five new species to add
new_sp_taxo_parnassiinae <- data.frame(Species = c("Parnassius_felderi",
                                                   "Parnassius_stenosemus",
                                                   "Parnassius_rueckbeili",
                                                   "Parnassius_turatii",
                                                   "Parnassius_augustus"),
                                       Subgenus = rep("", 5),
                                       Genus = rep("Parnassius", 5),
                                       Tribe = rep("Parnassius", 5))

taxo_parnassiinae2 <- taxo_parnassiinae[, c("Species",
                                            "Subgenus",
                                            "Genus",
                                            "Tribe")]

taxo_parnassiinae2 <- rbind(taxo_parnassiinae2,new_sp_taxo_parnassiinae)

taxo_parnassiinae2 <- taxo_parnassiinae2[!taxo_parnassiinae2$Species %in%
                                           c("Parnassius_felderi",
                                             "Parnassius_stenosemus",
                                             "Parnassius_rueckbeili"),]

c("Parnassius_labeyriei", "Parnassius_corybas", "Parnassius_dongalaicus") %in%
  taxo_parnassiinae2$Species

for(sp in 1:nrow(matching_names)){
```

```r
  tree_parnassiinae$tip.label[
    tree_parnassiinae$tip.label == matching_names$tree_names[sp]] <-
    matching_names$taxo_names[sp]
}

new_sp <- tree_parnassiinae$tip.label[
  !tree_parnassiinae$tip.label %in% taxo_parnassiinae2$Species]

new_sp_taxo_parnassiinae <- data.frame(Species = new_sp,
                                       Subgenus = c(rep("Parnassius", 4),
                                                    rep("Koramius",3),
                                                    "Driopa"),
                                       Genus = rep("Parnassius", 8),
                                       Tribe = rep("Parnassiini", 8))

taxo_parnassiinae2 <- rbind(taxo_parnassiinae2, new_sp_taxo_parnassiinae)

taxo_parnassiinae2$Species[!taxo_parnassiinae2$Species %in% tree_parnassiinae$tip.label]
rownames(taxo_parnassiinae2) <- NULL

tree_parnassiinae$tip.label %in% taxo_parnassiinae2$Species

taxo_parnassiinae2$Tribe[taxo_parnassiinae2$Tribe == "Parnassius"] <- "Parnassiini"

taxo_parnassiinae2$Subgenus[
  taxo_parnassiinae2$Species == "Parnassius_turatii"] <- "Driopa"

taxo_parnassiinae2$Subgenus[
  grepl("Parnassius_apollo", taxo_parnassiinae2$Species)] <- "Parnassius"

taxo_parnassiinae2$Subgenus[
  taxo_parnassiinae2$Species == "Parnassius_augustus"] <- "Kailasius"

taxo_parnassiinae2$Subgenus[taxo_parnassiinae2$Subgenus == ""] <-
  taxo_parnassiinae2$Genus[taxo_parnassiinae2$Subgenus == ""]

setdiff(taxo_parnassiinae2$Species, tree_parnassiinae$tip.label)

taxo_parnassiinae <- taxo_parnassiinae2

write.csv(taxo_parnassiinae, "data/cleaned/taxo_parnassiinae_updated.csv",
          row.names = F)
write.tree(tree_parnassiinae, "data/cleaned/tree_parnassiinae_updated.tre")
```

##### 2. Cycadales

The raw data comes from Condamine et al. (2015). We also used posterior trees to evaluate the uncertainty of the dating process. We analyzed the data with the taxonomy from the IUCN Red List database to rename, drop or add species depending on their taxonomic situation.

We also illustrated how the method can be applied to a sample of posteriors trees to take into account for the uncertainty of the dating process in the paleodiversity dynamic estimate.

```r
tree_cycads <- read.tree("data/raw/Cycadales_6FC_BD_100M_MCC.tre")

tree_cycads_posteriors <- read.nexus("data/raw/Cycadales_6FC_BD_100M.trees")
# this file was too heavy to be loaded on github
# but the object is in the environment empirical_cases.RData

tree_cycads_posteriors <- tree_cycads_posteriors[-c(1:2500)]

set.seed(1)
sub_sample <- sample(1:length(tree_cycads_posteriors), 1000)

tree_cycads_posteriors <- tree_cycads_posteriors[sub_sample]

# The most Consensus Tree is stored as the last element of
# tree_cycads_posteriors
tree_cycads_posteriors <- c(tree_cycads_posteriors, tree_cycads)

taxo_cycads <- read.csv("data/raw/iucn_data_taxo_cycads/taxonomy.csv")

taxo_cycads <- taxo_cycads[,names(taxo_cycads) %in% c("scientificName",
                                                      "genusName",
                                                      "familyName")]
taxo_cycads <- taxo_cycads[,c(1,3,2)]
names(taxo_cycads) <- c("Species", "Genus", "Family")

taxo_cycads$Species <- gsub(" ", "_", taxo_cycads$Species)

synonyms_cycads <- read.csv("data/raw/iucn_data_taxo_cycads/synonyms.csv")
synonyms_cycads <- synonyms_cycads[, names(synonyms_cycads) %in%
                                     c("genusName",
                                       "speciesName",
                                       "scientificName")]

synonyms_cycads$scientificName <- gsub(" ", "_", synonyms_cycads$scientificName)
synonyms_cycads$Synonym <- paste(synonyms_cycads$genusName,
                                 synonyms_cycads$speciesName, sep = "_")
names(synonyms_cycads)[1] <- "Species"
synonyms_cycads <- synonyms_cycads[, c("Species", "Synonym")]

synonyms_cycads <- unique(synonyms_cycads)

missing_sp_cycads <- tree_cycads$tip.label[!tree_cycads$tip.label %in%
                                             taxo_cycads$Species]

# Cycas
missing_sp_cycads[grepl("Cycas", missing_sp_cycads)]
synonyms_cycads[grepl("Cycas", synonyms_cycads$Species), c("Species", "Synonym")]

species_to_add <- data.frame(Species = NA, Genus= NA, Family = NA)
species_to_add[1,] <- c("Cycas_miquelii", "Cycas", "CYCADACEAE")

synonyms_cycads[grepl("pectinata", synonyms_cycads$Species),]
# Cycas_pectinata seems to be a synonym of Cycas_jenkinsiana but two studies
```

```r
# showed very different phylogenetic results with the same name
# so we keep Cycas_pectinata_1 and Cycas_pectinata_2 as different species.

taxo_cycads$Species[
  taxo_cycads$Species == "Cycas_pectinata"] <- "Cycas_pectinata_1"
species_to_add[2,] <- c("Cycas_pectinata_2", "Cycas", "CYCADACEAE")

synonyms_cycads[grepl("litoralis", synonyms_cycads$Synonym),
                c("Species", "Synonym")]
# Cycas_litoralis seems to be a synonym of Cycas_edentata but
# very different in the phylogeny: to add

species_to_add[3,] <- c("Cycas_litoralis", "Cycas", "CYCADACEAE")

# Cycas_chamberlainii is a synonym of its sister species Cycas_riuminiana: to drop
species_to_drop <- c("Cycas_chamberlainii")
species_to_add[4,] <- c("Cycas_ensata", "Cycas", "CYCADACEAE")

# Macrozamia
missing_sp_cycads[grepl("Macrozamia", missing_sp_cycads)]
taxo_cycads$Species[taxo_cycads$Genus == "Macrozamia"]
taxo_cycads$Species[
  taxo_cycads$Species == "Macrozamia_pauli-guilielmi"] <-
  "Macrozamia_pauliguilielmi"
# typo error for this species

# Ceratozamia
missing_sp_cycads[grepl("Ceratozamia", missing_sp_cycads)]
sort(taxo_cycads$Species[taxo_cycads$Genus == "Ceratozamia"])
# Ceratozamia_chimalapensis and Ceratozamia_decumbens are in the phylogeny

# new species Medina-Villarreal et al. 2019
species_to_add[5,] <- c("Ceratozamia_totonacorum", "Ceratozamia", "ZAMIACEAE")
species_to_add[6,] <- c("Ceratozamia_chimalapensis", "Ceratozamia", "ZAMIACEAE")
species_to_add[7,] <- c("Ceratozamia_decumbens", "Ceratozamia", "ZAMIACEAE")
species_to_add[8,] <- c("Ceratozamia_hondurensis", "Ceratozamia", "ZAMIACEAE")
species_to_add[9,] <- c("Ceratozamia_santillanii", "Ceratozamia", "ZAMIACEAE")
species_to_add[10,] <- c("Ceratozamia_tenuis", "Ceratozamia", "ZAMIACEAE")

# Encephalartos
missing_sp_cycads[grepl("Encephalartos", missing_sp_cycads)]
sort(taxo_cycads$Species[taxo_cycads$Genus == "Encephalartos"])

taxo_cycads$Species[taxo_cycads$Species == "Encephalartos_eugene-maraisii"] <-
  "Encephalartos_eugenemaraisii"

taxo_cycads$Species[taxo_cycads$Species == "Encephalartos_friderici-guilielmi"] <-
  "Encephalartos_fridericiguilielmi"

# Encephalartos allochrous can be elevated to species level
# because it is very divergent from Encephalartos_barteri
species_to_add[11,] <- c("Encephalartos_allochrous", "Encephalartos", "ZAMIACEAE")
```

```r
# Encephalartos_barteri to remove
species_to_drop <- c(species_to_drop, "Encephalartos_barteri_barteri")

# Zamia
sort(taxo_cycads$Species[taxo_cycads$Genus == "Zamia"])

missing_sp_cycads[grepl("Zamia", missing_sp_cycads)]
synonyms_cycads[synonyms_cycads$Synonym == "Zamia_lindenii",]
# too divergent (8 Myrs and not sister species) to be synonyms
species_to_add[12,] <- c("Zamia_lindenii", "Zamia", "ZAMIACEAE")

missing_sp_cycads[grepl("Zamia", missing_sp_cycads)]
synonyms_cycads[synonyms_cycads$Synonym == "Zamia_splendens",]
# no synonym for Zamia_splendens: to add

# other species to add
species_to_add[13,] <- c("Zamia_brasiliensis", "Zamia", "ZAMIACEAE")
species_to_add[14,] <- c("Zamia_paucifoliolata", "Zamia", "ZAMIACEAE")
species_to_add[15,] <- c("Zamia_pyrophylla", "Zamia", "ZAMIACEAE")
species_to_add[16,] <- c("Zamia_imbricata", "Zamia", "ZAMIACEAE")
species_to_add[17,] <- c("Zamia_sinuensis", "Zamia", "ZAMIACEAE")
species_to_add[18,] <- c("Zamia_stenophyllidia", "Zamia", "ZAMIACEAE")
species_to_add[19,] <- c("Zamia_lindosensis", "Zamia", "ZAMIACEAE")
species_to_add[20,] <- c("Zamia_splendens", "Zamia", "ZAMIACEAE")

species_to_drop <- c(species_to_drop, "Zamia_picta")
species_to_drop <- c(species_to_drop, "Zamia_lawsoniana")
species_to_drop <- c(species_to_drop, "Zamia_furfuraceaBL")
species_to_drop <- c(species_to_drop, "Zamia_kickxii")

taxo_cycads <- rbind(taxo_cycads, species_to_add)

# update family name
taxo_cycads$Family[taxo_cycads$Family == "STANGERIACEAE"] <- "ZAMIACEAE"

taxo_cycads$Family <- capitalize(tolower(taxo_cycads$Family))

taxo_cycads$Genus[taxo_cycads$Genus == "Dioon"] <-
  taxo_cycads$Species[taxo_cycads$Genus == "Dioon"]

for(i_trees in 1:length(tree_cycads_posteriors)){
  cat("\n", i_trees, "/", length(tree_cycads_posteriors))
  # Cycas

  # Cycas_ensata is a new species because too divergent from
  # Cycas_media_media that becomes Cycas_media
  tree_cycads_posteriors[[i_trees]]$tip.label[
    tree_cycads_posteriors[[i_trees]]$tip.label == "Cycas_media_media"] <- "Cycas_media"

  tree_cycads_posteriors[[i_trees]]$tip.label[
    tree_cycads_posteriors[[i_trees]]$tip.label == "Cycas_media_ensata"] <- "Cycas_ensata"

  # Encephalartos
```

```
  tree_cycads_posteriors[[i_trees]]$tip.label[
    tree_cycads_posteriors[[i_trees]]$tip.label == "Encephalartos_barteri_allochrous"] <-
    "Encephalartos_allochrous"

  # Zamia
  tree_cycads_posteriors[[i_trees]]$tip.label[
    tree_cycads_posteriors[[i_trees]]$tip.label == "Zamia_furfuraceaAL"] <-
    "Zamia_furfuracea"

  tree_cycads_posteriors[[i_trees]]$tip.label[
    tree_cycads_posteriors[[i_trees]]$tip.label == "Zamia_amblyphyllidia"] <-
    "Zamia_erosa"

  # final
  tree_cycads_posteriors[[i_trees]] <- drop.tip(tree_cycads_posteriors[[i_trees]],
                                                c(species_to_drop, "Ginkgo_biloba"))
  print(all(tree_cycads_posteriors[[i_trees]]$tip.label %in% taxo_cycads$Species))
}
write.csv(taxo_cycads, "data/cleaned/taxo_cycads_cleaned.csv", row.names = F)
write.tree(tree_cycads_posteriors, "data/cleaned/cycads_posteriors_and_MCC.tre")
```

#### II - Analyses

The main procedure consists in four steps: (1) calculating the sampling fractions for the selected subclades, (2) calculating all the combinations of selected subclades that can be tested together, (3) applying diversification models to the parts of all combination and selecting best combinations from their global AICc comparisons, (4) calculating the paleodiversity dynamic with the deterministic or the probabilistic approach (see Figure 1 from the main text). To avoid being repetitive, these four steps will only be detailed for the Cetacea example.

##### 1. Cetacea

For the case study of the Cetacea, we only tested shifts for the four main families of the group (Balaenopteridae, Delphinidae, Phocoenidae and Ziphiidae) and two the parvorders (Mysticeti and Odontoceti) as in the original paper of the method (Morlon *et al.* 2011). To do so, we removed the column "Genus" from the taxonomic database and calculated the sampling fractions for these groups.

```
data("Cetacea")
data("taxo_cetacea")
taxo_cetacea <- taxo_cetacea[,!names(taxo_cetacea) %in% c("Genus")]
par(xpd = T)
f_cetacea <- get.sampling.fractions(Cetacea, taxo_cetacea, clade.size = 5,
                                    plot = T, cex = 0.3, text.cex = 0.5,
                                    pch.cex = 1,
                                    edge.width = 1)
```

Now, we can visualize the shifts that can be tested and display their sampling fractions ($f$) thanks to this line:

```
f_cetacea[!is.na(f_cetacea$to_test),]
```

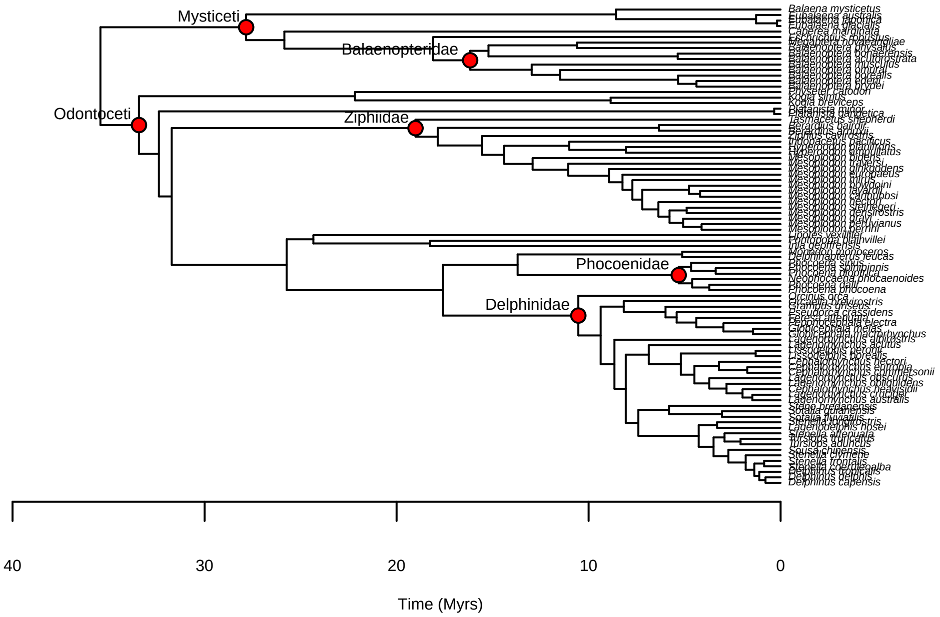

Figure 1: Phylogeny of Cetacea with testable nodes

Then, we can calculate the combinations of subclades where shifts can be tested thanks to `get.comb.shift()`. The output is a vector of node IDs. Node IDs where shifts are tested are at the left of the "/" while node IDs of shifts located in the backbone are at the right.

```
comb_cetacea <- get.comb.shift(Cetacea, taxo_cetacea, f_cetacea,
                               clade.size = 5, Ncores = 4)
```

Here are two examples: the second element of `comb_cetacea` is the combination with a shift at node 95 (Balaenopteridae) and with a shift in the backbone at node 89 (Mysticeti). The thirty senventh element is the combination with nodes 89, 109 and 140 (Mysticeti, Ziphiidae, Delphinidae) and no possible shift in the backbone (`NULL` in the vector).

```
comb_cetacea[2]
```

```
## [1] "95/89"
```

```
comb_cetacea[37]
```

```
## [1] "89.109.140/"
```

Now, we can use the function `shift.estimates()` to apply diversification models to all parts of all combinations and compare the global AICc of all combinations. The first analysis is done without multiple backbones and by including the probabilities of survival of the stems of subclades in the backbone analysis (`backbone.option = "crown.shift"`, by default). We removed the model `"BVAR_DVAR"` as it can be problematic to estimate rates (see Burin et al. 2019).

```
shifts_cetacea <- shift.estimates(phylo = Cetacea, data = taxo_cetacea,
                                  sampling.fractions = f_cetacea,
                                  models = c("BCST", "BCST_DCST", "BVAR",
                                             "BVAR_DCST", "BCST_DVAR"),
                                  comb.shift = comb_cetacea,
                                  Ncores = 12, np.sub = 3,
                                  multi.backbone = "all")
```

The output of `shift.estimates()` is a list with four elements (see help files for more details). What interests us here is the comparison of global AICc. We can print the head of this comparison thanks to the following line:

```
head(shifts_cetacea$total)
```

Here, we can see that the best combination is better than the second best combination ($\Delta$AICc = 8.5). We may also want to look at rates estimated for all parts of the best combination. We can do this with the following lines by selecting the backbone and subclade elements that correspond to the best combination.

```
# For the backbone
shifts_cetacea$backbones[shifts_cetacea$total$Combination[1]]
```

```
## $`95.109.135.140/`
## $`95.109.135.140/`$`95.109.135.140_bck`
##      Models Parameters      logL     AICc     Lambda      Alpha          Mu
## 5 BCST_DVAR          3 -57.11860 122.2372 0.33241221         NA 1.24529302
```

```
## 4 BVAR_DCST          3 -59.59423 127.1885 0.26664678 0.04150521 0.54468135
## 1      BCST          1 -70.63969 143.5651 0.05371094         NA         NA
## 3      BVAR          2 -69.38357 143.6902 0.02846093 0.04023431         NA
## 2 BCST_DCST          2 -70.25991 145.4429 0.08649512         NA 0.05550311
##         Beta delta_AICc
## 5 -0.1327028   0.000000
## 4         NA   4.951265
## 1         NA  21.327907
## 3         NA  21.453016
## 2         NA  23.205706
```

```
subclades <- strsplit(shifts_cetacea$total$Combination[1], "[.]")[[1]]

# For subclades with node 95 (Balaenopteridae) as an example
shifts_cetacea$subclades[subclades]$`95`
```

We did the same analysis but with the inclusion of the probabilities of survival of the stems of subclades in the likelihood of subclades (`backbone.option = "stem.shift"`) as in Morlon et al. (2011). This analysis is illustrated in Appendix S1.

```
shifts_cetacea_stem.shift <- shift.estimates(phylo = Cetacea,
                                             data = taxo_cetacea,
                                             sampling.fractions = f_cetacea,
                                             models = c("BCST", "BCST_DCST", "BVAR",
                                                        "BVAR_DCST", "BCST_DVAR"),
                                             comb.shift = comb_cetacea,
                                             Ncores = 12, np.sub = 3,
                                             backbone.option = "stem.shift",
                                             multi.backbone = "all")
```

Now that the best combinations are known, we can calculate the paleodiversity with the probabilistic approach from `shift.estimates()` output thanks to the function `apply_prob_dtt()` as below. This method allows to calculate uncertainty around paleodiversity dynamic estimates. To do so, `apply_prob_dtt()` calls `prob_dtt()` from RPANDA and returns a list of `prob_dtt()` outputs (which are matrices). As explained in Billaud et al. (2020), all the sum of probabilities per million year must be equal to 1. However, it can be difficult to reach one for groups showing a decline in the paleodiversity dynamic (all except Vangidae) because the range of paleodiversity over which we need to calculate the probabilities can be very large. To circumvent this issue, `apply_prob_dtt()` set the range of the paleodiversity to the maximum of the deterministic estimate from the function `paleodiv()` and successively multiplies this maximum by 2, 3, 5, 7 and 10 until the sums of probabilities for each million year reach a minimum of 95%. Because of this way to proceed, `apply_prob_dtt()` might take time.

```
# Probabilistic approach
prob_dtt_cetacea <- apply_prob_dtt(phylo = Cetacea,
                                   data = taxo_cetacea,
                                   sampling.fractions = f_cetacea,
                                   shift.res = shifts_cetacea)
```

```
##
## DTT calculation for subclade(s):
##   1 / 4
##   2 / 4
##   3 / 4
```

```
##   4 / 4
##
## DTT calculation for backbone(s):
##   1 / 1
## with a maximum value of m = 1140 (max. deterministic value x2)
##  -> minimum value of the sum of probabilities/Myr= 0.9900762
```

We can check the minimum of the sum of the probability with these following lines.

```
# checking probabilities
lapply(prob_dtt_cetacea, function(x) sapply(x, function(x) min(colSums(x))))
```

```
## $backbones
##        88
## 0.9900762
##
## $subclades
##  95 109 135 140
##   1   1   1   1
```

Here, this minimum is equal to 99% of the probabilities.

In few cases, this value of 95% is not reached for few million years (see help files and next example for more details).

We did the same for the analysis with `backbone.option = "stem.shift"`:

```
# Probabilistic approach
prob_dtt_cetacea_stem <- apply_prob_dtt(phylo = Cetacea,
                                        data = taxo_cetacea,
                                        sampling.fractions = f_cetacea,
                                        backbone.option = "stem.shift",
                                        shift.res = shifts_cetacea_stem.shift)
```

Unfortunately, we can see that the sum of the probabilities did not reach 95% for all Myrs.

```
# checking probabilities
lapply(prob_dtt_cetacea_stem, function(x) sapply(x, function(x) min(colSums(x))))
```

```
## $backbones
##        88
## 0.9033125
##
## $subclades
##  95 109 135 140
##   1   1   1   1
```

We can identify for which time point these sums are below 95%.

```
which(colSums(prob_dtt_cetacea_stem$backbones[[1]]) ==
        min(colSums(prob_dtt_cetacea_stem$backbones[[1]])))
```

```
## 34.424789
##         1
```

In this case, this is only the inference at the the crown age that is below 95%, which is reassuring for the rest of the estimations.

##### 2. Vangidae

For the case study of the Vangidae, we analyzed the two morphological shifts found in Jønsson et al. (2012). We also tested the monophyletic group corresponding to the malagasy species. We did not detail the steps as they are the same than for the Cetacea example. We did not used `backbone.option = "stem.shift"` because we expected the shift to be at the crown of the subclades.

```
# loading taxonomy
taxo_vangidae <- read.csv("data/raw/Vangidae_taxonomy.csv")
tree_vangidae <- read.tree("data/raw/Vanga_CONS.tre")

taxo_vangidae$genus <- factor(taxo_vangidae$genus)
colnames(taxo_vangidae) <- Hmisc::capitalize(colnames(taxo_vangidae))

significant_shift <- c("Xenopirostris_polleni", "Xenopirostris_xenopirostris",
                       "Xenopirostris_damii", "Oriolia_bernieri",
                       "Falculea_palliata", "Artamella_viridis")

marginal_shift <- c("Xenopirostris_polleni", "Xenopirostris_xenopirostris",
                    "Xenopirostris_damii", "Oriolia_bernieri",
                    "Falculea_palliata", "Artamella_viridis",
                    "Schetba_rufa", "Euryceros_prevostii")

taxo_vangidae$marginal_shift[
  taxo_vangidae$Species %in% marginal_shift] <- "marginal_shift"
taxo_vangidae$marginal_shift[is.na(taxo_vangidae$marginal_shift)] <- ""
taxo_vangidae$significant_shift[
  taxo_vangidae$Species %in% significant_shift] <- "significant_shift"
taxo_vangidae$significant_shift[is.na(taxo_vangidae$significant_shift)] <- ""

# getting sampling fractions
f_vangidae <- get.sampling.fractions(phylo = tree_vangidae,
                                     data = taxo_vangidae)

# getting combinations
comb_vangidae <- get.comb.shift(phylo = tree_vangidae,
                                data = taxo_vangidae,
                                sampling.fractions = f_vangidae,
                                Ncores = 5, clade.size = 5)

# simple backbone
t1 <- Sys.time()
shift.vangidae <- shift.estimates(phylo = tree_vangidae,
                                  data = taxo_vangidae,
                                  sampling.fractions = f_vangidae,
                                  models = c("BCST", "BCST_DCST", "BVAR",
                                             "BVAR_DCST", "BCST_DVAR"),
```

```
                                     comb.shift = comb_vangidae, np.sub = 2,
                                     Ncores = 5, multi.backbone = "all")
t2 <- Sys.time()
time_vangidae <- t2 - t1

# Probabilistic approach
prob_dtt_vangidae <- apply_prob_dtt(phylo = tree_vangidae,
                                       data = taxo_vangidae,
                                       sampling.fractions = f_vangidae,
                                       shift.res = shift.vangidae)
# checking probabilities
sapply(prob_dtt_vangidae, function(x) min(colSums(x)))
```

##### 3. Parnassiinae

For the case study of the Parnassiinae, we tested taxonomic groups corresponding to host-plant shifts or mountain colonizations (Condamine et al. 2018).

```
# loading taxonomy and phylogeny
tree_parnassiinae <- read.tree("data/cleaned/tree_parnassiinae_updated.tre")
taxo_parnassiinae <- read.csv("data/cleaned/taxo_parnassiinae_updated.csv")

# getting sampling fractions
f_parnassiinae <- get.sampling.fractions(tree_parnassiinae, taxo_parnassiinae)
f_parnassiinae[!is.na(f_parnassiinae$to_test),]
```

Because we need the sampling fractions, testing shifts based on ecological cannot be totally disconnected from taxonomy. In our case of Parnassiinae, taxonomy and ecology overlap for some clades. For few examples, the tribes *Luehdorfiini* and *Zerynthiini* feed on Aristochiaceae while the genus *Panassius* matches with the colonisation of Hymalian and Tibetan Plateau (Condamine et al. 2018). If you work with an well-sampled phylogeny, some genus or family might be fully sampled. In this case, it is easy to tests shifts inside this taxa based on ecological or morphological features.

The first analysis made on this phylogeny has highlight that some combinations with a better global fit but with very unrealistic rates.

```
# getting combinations
comb_parnassiinae <- get.comb.shift(phylo = tree_parnassiinae,
                                       data = taxo_parnassiinae,
                                       sampling.fractions = f_parnassiinae,
                                       Ncores = 12)

# simple backbone
t1 <- Sys.time()
shifts_parnassiinae <- shift.estimates(phylo = tree_parnassiinae,
                                          data = taxo_parnassiinae,
                                          sampling.fractions = f_parnassiinae,
                                          comb.shift = comb_parnassiinae,
                                          models = c("BCST", "BCST_DCST", "BVAR",
                                                "BVAR_DCST", "BCST_DVAR"),
                                          Ncores = 15, np.sub = 3)
t2 <- Sys.time()
time_parnassiinae <- t2 - t1
```

We can see if we look at the backbone model comparison of the best combination is extremely higher than other combinations. We see that the model BVAR_DVAR behaves oddly compared to all other models.

```r
# The head of the global comparison
head(shifts_parnassiinae$total)

# the backbone model comparisons of the best combination
shifts_parnassiinae$backbones$`95.111.130.137.147.155.160`
```

We can see that both speciation and extinction rate values are extremely high and unrealistic.

Seeing this, we can first remove the model `"BCST_DCST"` from the model comparison thanks to the function `remove.models()` to see if global AICc of combinations recalculated without `"BCST_DCST"` still show results with unrealistic rate values with large difference of AICc with other models.

```r
# removing BCST_DCST
shifts_parnassiinae_noBCST_DCST <- remove.model(shifts_parnassiinae, "BCST_DCST")

head(shifts_parnassiinae_noBCST_DCST$total)
```

```r
shifts_parnassiinae_noBCST_DCST$backbones$`88.95.111.130.137.147.160`
```

```
## $‘88.95.111.130.137.147.160_bck‘
##      Models Parameters      logL     AICc     Lambda         Alpha       Mu
## 4 BVAR_DCST          3 -10.20117 32.40235 205.089963  0.0001028021 205.5301
## 5 BCST_DVAR          3 -10.28406 32.56812 188.523528            NA 188.9267
## 1      BCST          1 -46.64697 95.96061   0.121250            NA       NA
## 3      BVAR          2 -45.88804 98.17608   0.181652 -0.0292331003       NA
##            Beta delta_AICc
## 4            NA  0.0000000
## 5 -9.725786e-05  0.1657715
## 1            NA 63.5582624
## 3            NA 65.7737296
```

We can see that this is still the case with the model `"BVAR_DCST"` which estimates extremely important values for $\lambda$ and $\mu$.

To circumvent this issue, we applied constrains on diversification rate by limiting rate values to 2. This constrain allowed us to find a more realistic and reliable results that is discussed in the main manuscript. We led the analysis with all combinations (the ones with multiple backbones included) and we did not used `backbone.option = "stem.shift"` because we expected the shift to be at the crown of the subclades.

```r
# adding constrains
t1 <- Sys.time()
shifts_parnassiinae_rmax2 <- shift.estimates(phylo = tree_parnassiinae,
                                             data = taxo_parnassiinae,
                                             sampling.fractions = f_parnassiinae,
                                             comb.shift = comb_parnassiinae,
                                             models = c("BCST", "BCST_DCST", "BVAR",
                                                        "BVAR_DCST", "BCST_DVAR"),
                                             Ncores = 15, np.sub = 3, rate.max = 2,
                                             multi.backbone = "all")
t2 <- Sys.time()
time_parnassiinae_rmax2 <- t2 - t1
```

We can see that there are three best combinations with a $\Delta$AICc $< 2$.

```
head(shifts_parnassiinae_rmax2$total)
```

Now we can quickly estimates the palaeodiversity dynamics corresponding to these three combinations of shifts to see if there are more realistic than without the constrain.

```
paleodiv(phylo = tree_parnassiinae,
         data = taxo_parnassiinae,
         sampling.fractions = f_parnassiinae,
         shift.res = shifts_parnassiinae_rmax2,
         combi = 1)
```

```
##  [1]    37.51136    78.07039   238.62022   644.24462  1549.25435  3344.16260
##  [7]  6526.55507 11594.22480 18867.97734 28288.60886 39294.06326 50826.30205
## [13] 61513.04403 69967.39588 75106.49333 76382.78201 73862.70730 68143.26565
## [19] 60167.41460 50993.73595 41598.86022 32746.78959 24935.61315 18408.18921
## [25] 13202.59905  9217.95796  6277.30385  4177.35970  2721.96426  1740.57255
## [31]  1095.41937   681.34761   423.26571   264.67951   170.91709   117.90433
## [37]    90.52207    79.52885    79.59380    88.00000
```

```
paleodiv(phylo = tree_parnassiinae,
         data = taxo_parnassiinae,
         sampling.fractions = f_parnassiinae,
         shift.res = shifts_parnassiinae_rmax2,
         combi = 2)
```

```
##  [1]  19.53880  24.03555  33.51588  45.97229  62.01440  82.25080 107.23540
##  [8] 137.39923 174.35008 215.31009 261.28252 311.35652 364.23579 418.17943
## [15] 471.06232 521.63348 565.12439 600.31559 625.02969 637.66772 637.31098
## [22] 623.83043 597.91817 561.03407 515.27253 463.16638 407.45353 350.83704
## [29] 295.76852 246.29166 201.55760 162.05132 131.32336 109.40068  91.98879
## [36]  82.32858  77.78089  77.41679  80.80220  88.00000
```

```
paleodiv(phylo = tree_parnassiinae,
         data = taxo_parnassiinae,
         sampling.fractions = f_parnassiinae,
         shift.res = shifts_parnassiinae_rmax2,
         combi = 3)
```

```
##  [1]    37.51136    78.07039   238.62022   644.24462  1549.25435  3344.16260
##  [7]  6526.55507 11594.22480 18867.97734 28288.60886 39294.06326 50826.30205
## [13] 61513.04403 69967.39588 75106.49333 76382.78201 73862.29184 68142.81728
## [19] 60166.90882 50993.16875 41598.22855 32746.09195 24934.85042 18407.36577
## [25] 13201.72424  9217.04807  6276.38470  4176.47006  2721.16029  1739.93356
## [31]  1095.05533   681.40878   422.39624   264.34787   170.92892   118.29787
## [37]    91.27522    80.50184    80.43583    88.00000
```

As we can see, the combination 1 and 3 involving a shift at the node 108 provide an highly unrealistic maximum for their paleodiversity dynamics. In consequence, we only applied the probabilistic approach for calculating the paleodiversity dynamic with the second combination which is the more reliable scenario in terms of the number of species estimated.

```r
# Probabilistic approach
prob_dtt_parnassiinae_rmax2 <- apply_prob_dtt(phylo = tree_parnassiinae,
                                              data = taxo_parnassiinae,
                                              sampling.fractions = f_parnassiinae,
                                              shift.res = shifts_parnassiinae_rmax2,
                                              combi = 2)
# checking probabilities
lapply(prob_dtt_parnassiinae_rmax2, function(x) sapply(x, function(x) min(colSums(x))))

# calculating the overall maximum
mat <- prob_dtt_parnassiinae_rmax2$backbones[[1]]
mean.val <- t(as.numeric(rownames(mat))) %*% mat
max_mean_div <- max(mean.val)
```

###### 4. Cycadales

The phylogeny of Cycadales harbors an extremely specific shape due the ancient origin of the group dating from the Permian and the recent radiations that have created the extant paleodiversity. We decided to test shift for the main radiating genera of the group excepting the genus *Dioon* which is the poorest with at least five species to avoid creating a very poor backbone with such long branches that can be difficult to analyze. To do so, we simply replaced the genus name in the genus column of the taxonomy by the species names. Because, the analyses on the posteriors are time-consuming and have been made on a more powerful machine, we here only applied the approach to the most consensus tree and downloaded the result for posterior tree analyses. We did not apply the model BVAR_DVAR and we only analyzed the phylogeny with `backbone.option = "crown.shift"` which is the default option.

```r
taxo_cycads <- read.csv("data/cleaned/taxo_cycads_cleaned.csv")
tree_cycads_posteriors <- read.tree("data/cleaned/cycads_posteriors_and_MCC.tre")

# Most Consensus Tree has been stored as the last of element of tree_cycads_posteriors
# (see Cleaning data part)
tree_cycads <- tree_cycads_posteriors[[length(tree_cycads_posteriors)]]

f_cycads <- get.sampling.fractions(phylo = tree_cycads, data = taxo_cycads)

# testing only genus
taxo_cycads <- taxo_cycads[,names(taxo_cycads) %in% c("Species", "Genus")]

comb_cycads <- get.comb.shift(phylo = tree_cycads,
                              data = taxo_cycads,
                              sampling.fractions = f_cycads,
                              Ncores = 5)

shift_cycads <- shift.estimates(phylo = tree_cycads,
                                data = taxo_cycads,
                                sampling.fractions = f_cycads,
                                comb.shift = comb_cycads,
                                models = c("BCST", "BCST_DCST", "BVAR",
                                           "BVAR_DCST", "BCST_DVAR"),
                                Ncores = 12, multi.backbone = "all")

# same analysis made on posteriors (already done)
load("res/cycads_bigmem_1.RData")
```

```
load("res/cycads_bigmem_2.RData")
# merge all posteriors
all_shift_cycads[1:500] <- all_shift_cycads1

all_shift_cycads <- lapply(all_shift_cycads, remove.model, "BVAR_DVAR")
# merge all posteriors (and removing the analysis on the MCC)
shift_cycads_posteriors <- all_shift_cycads[-1001]

# Probabilistic approach
prob_dtt_cycads <- apply_prob_dtt(phylo = tree_cycads, data = taxo_cycads,
                                  sampling.fractions = f_cycads,
                                  shift.res = shift_cycads)
# checking the probabilities
lapply(prob_dtt_cycads, function(x) sapply(x, function(x) min(colSums(x))))
```

##### 5. Point about the model BVAR_DVAR

In all previous analyses, we did not include the model `BVAR_DVAR` in which both speciation and extinction vary. This model has already been criticised by Burin et al. (2018) and empirical tests have comfort that this model can be problematic. We quickly develop this point with an example.

Here is the command line to run the analysis on Cetacea with all models.

```
shifts_cetacea_allmodels <- shift.estimates(phylo = Cetacea,
                                             data = taxo_cetacea,
                                             sampling.fractions = f_cetacea,
                                             comb.shift = comb_cetacea,
                                             Ncores = 12,
                                             multi.backbone = "all")
```

If we look at the comparison of combinations, we can see that there is only one combination involving multiple backbones (denoted by the /103, which means that the shift at the node 103 creates two backbones).

If we look at the model comparison for the backbones, we can see that BVAR_DVAR behaves strangely.

```
shifts_cetacea_allmodels$backbones$`109.135.140/103`$`103_bck`
```

Indeed, the difference of logL and AICc with other models are really large but more importantly, the rate values are also extremely high. If we compute the paleodiversity dynamic from this combination, it will reach unrealistic values.

```
paleodiv(phylo = Cetacea,
         data = taxo_cetacea,
         sampling.fractions = f_cetacea,
         shift.res = shifts_cetacea_allmodels,
         combi = 1)
```

```
##  [1]  9.346447e+15  2.678421e+80 4.923070e+173 3.533150e+211 8.353907e+216
##  [6] 2.820721e+204 1.872791e+183 8.337547e+158 3.947571e+134 1.039911e+112
## [11]  9.611509e+91  5.527864e+74  1.703136e+60  1.658451e+48  2.551684e+38
## [16]  2.980508e+30  1.309617e+24  1.146401e+19  1.148138e+15  8.209148e+11
## [21]  2.827403e+09  3.394845e+07  1.092891e+06  7.659114e+04  1.012192e+04
```

```
## [26]  2.450102e+03  1.147338e+03  9.127101e+02  9.290872e+02  1.009729e+03
## [31]  1.079000e+03  1.072283e+03  9.398884e+02  6.754358e+02  3.681822e+02
## [36]  1.581777e+02  8.900000e+01
```

More investigations on the behavior of this model would be interesting to see whether the observed behavior of some backbone trees correspond to overfitting. For now, we advocate not to use the model BVAR_DVAR by default or to use it with constraints, although it does not ensure to avoid overfitting.

#### III - Testing model adequacy

The tendency to false positive showed thanks from simulations (see Appendix S3) might let us skeptical empirical results. One way to confirm whether models are reliable is to test for model adequacy. The idea is to simulate new data from the parameter of empirical results (thanks to the rate values of backbone(s) and subclade(s) and the shift ages) to see how the empirical data are close to the new simulated data.

##### 1. Cetacea

We used the function `simul.comb.shift()` to simulate posterior trees under the best combinations of the Cetacea analysis.

Here the code to do so:

```
# to edit with updates (option for the number of simulations)
all_posteriors_cetacea <- simul.comb.shift(phylo = Cetacea,
                                           sampling.fractions = f_cetacea,
                                           shift.res = shifts_cetacea)
cetacea_posterior_trees <- all_posteriors_cetacea[1:500]
```

We can look at the number of tips in each part of the simulated trees to see whether simulated data are similar to the empirical tree of Cetacea.

```
ntip_by_groups <- lapply(cetacea_posterior_trees,
                         function(x) table(sapply(strsplit(x$tip.label,
                                                           split = ""), "[[", 1)))
ntip_by_groups_df <- as.data.frame(do.call(rbind, ntip_by_groups))
ntip_by_groups_cetacea <- c(f_cetacea$sp_in[!is.na(f_cetacea$to_test)][c(2,4:6)], 87-71)

par(mfrow = c(1,2), cex = 0.6)
boxplot(ntip_by_groups_df, las = 1, main = "Species richness by group",
        ylab = "Number of species", xlab = "Groups")
points(c(1:5), ntip_by_groups_cetacea, pch = 19, col = "red")
legend("topright", bty = "n", legend = "Empirical values", pch = 19, col = "red", cex = 1)

avgLeafDepI_Cetacea <- avgLeafDepI(Cetacea)
avgLeafDepI_Cetacea_posteriors <- sapply(cetacea_posterior_trees, avgLeafDepI)
hist(avgLeafDepI_Cetacea_posteriors,
     main = "Distribution of average leaf depth index",
     xlab = "Average leaf depth", las = 1)
abline(v = avgLeafDepI_Cetacea, col = "red")
```

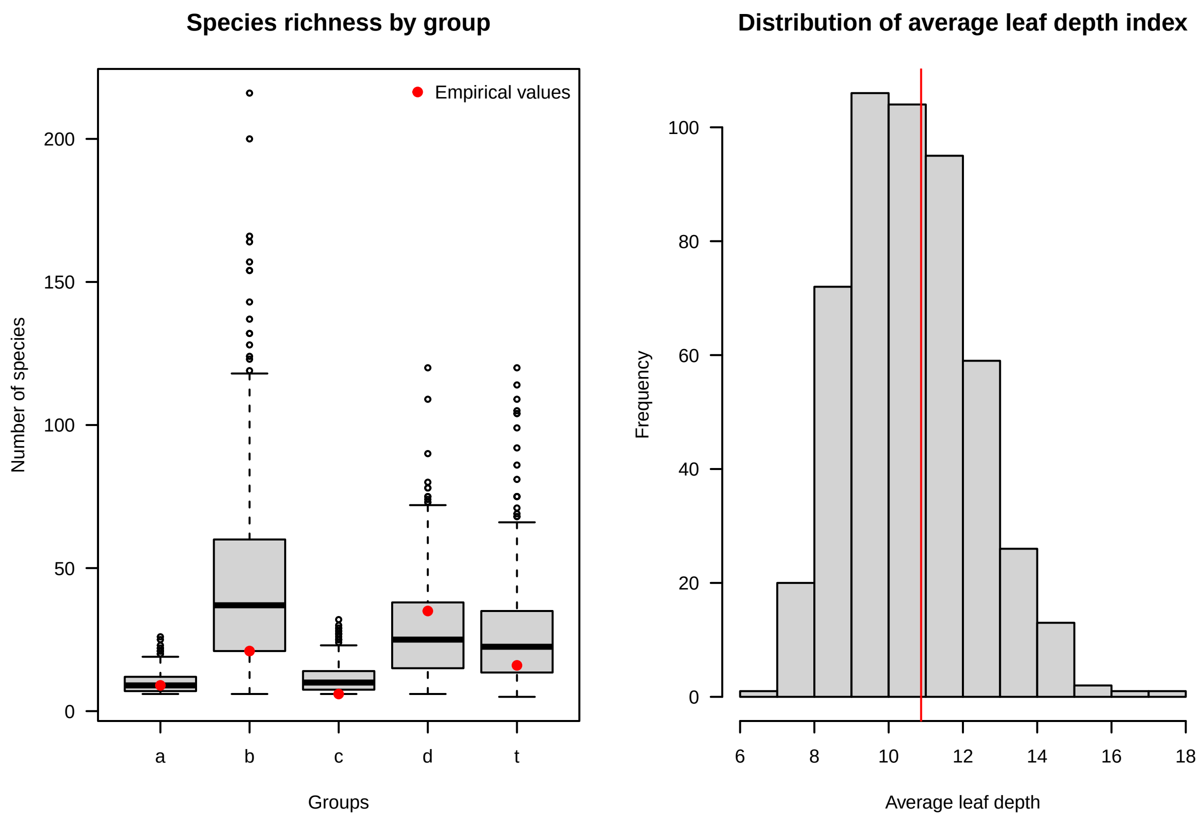

We can see that the empirical values fall into the interquartile of simulated data. We also tested the average leaf depth index which measure the imbalance fo the tree, the higher the index, the lower the balance of the tree. The empirical value falls into the distribution of simulated values.

To test not only the tips we can measure the proportion of values of the empirical Lineages-Through-Time (LTT) that are contained in a 95% confidence interval obtained from the LTT of simulated trees.

```
ltt_cetacea <- ltt(Cetacea, plot = F)
ltt_cetacea_df <- data.frame(times = round(ltt_cetacea$times, 4),
                             ltt = ltt_cetacea$ltt)

cetacea_posterior_trees_mp <- as.multiPhylo(cetacea_posterior_trees[[1]])
for(i in 2:length(cetacea_posterior_trees)){
  cetacea_posterior_trees_mp[[i]] <- cetacea_posterior_trees[[i]]
}
ltt95_CI <- ltt95(cetacea_posterior_trees_mp, log = T,  las = 1)
lines(ltt_cetacea_df$times, ltt_cetacea_df$ltt, type = "s", col = "red")
mtext(text = "Cetacea", side = 3, line = 1)

ltt95_CI_df <- as.data.frame(ltt95_CI[,c("time", "low(lineages)", "high(lineages)")])
ltt95_CI_df$time <- round(ltt95_CI_df$time, 4)

points_in_cetacea <- c()
for(i in 1:nrow(ltt_cetacea_df)){
  int_max <- sort(ltt95_CI_df$time[
    ltt95_CI_df$time >= ltt_cetacea_df$times[i]][1])
  int_min <- sort(ltt95_CI_df$time[ltt95_CI_df$time <= ltt_cetacea_df$times[i]],
```

```
        decreasing = T)[1]
  ltt_min <- ltt95_CI_df$`low(lineages)`[ltt95_CI_df$time == int_min]
  ltt_max <- ltt95_CI_df$`high(lineages)`[ltt95_CI_df$time == int_max]

  points_in_cetacea[i] <- ifelse(ltt_min <= ltt_cetacea_df$ltt[i] &
                                   ltt_cetacea_df$ltt[i] <= ltt_max, T, F)
}
legend("topleft",
       legend = c("95% of the distribution of simulated trees around the median",
                  "Median of simulated trees",
                  "Empirical data"),
       lty = c(3,1,1), lwd = c(1,2,1), col = c("black","black","red"),
       bty = "n", cex = 0.8)
```

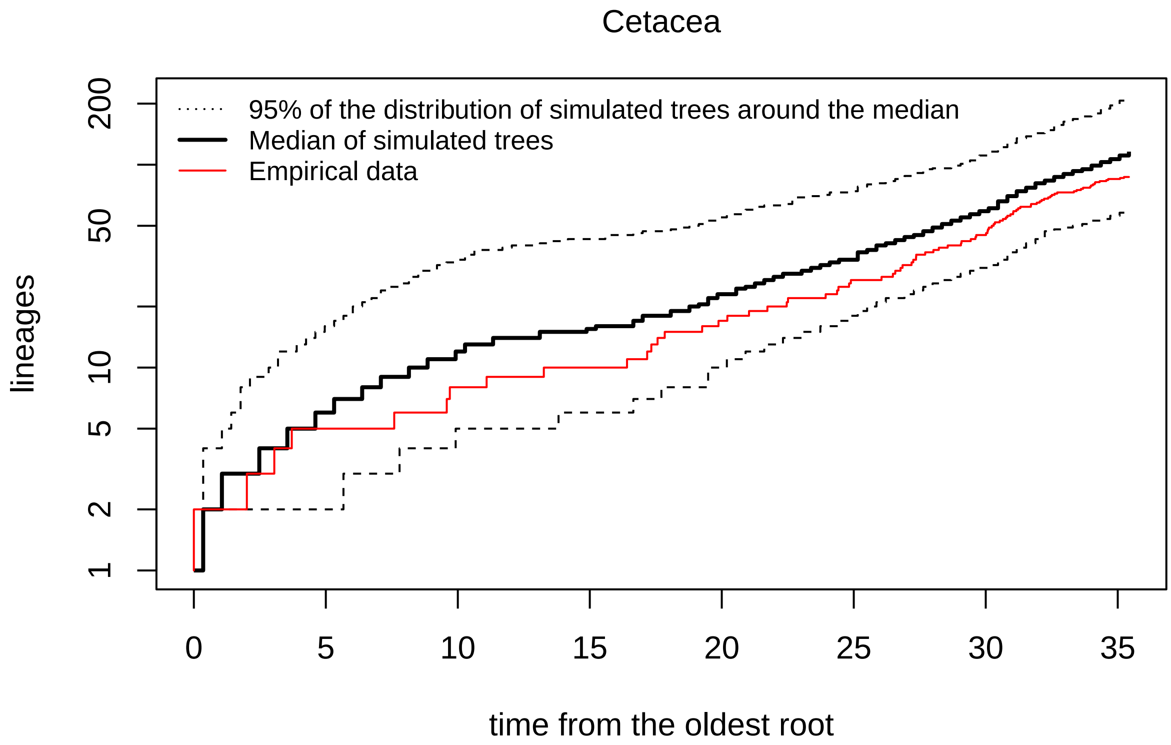

The plot above shows the median LTT plot of the posterior trees, the confidence interval at 95% in doted lines and the empirical LTT in red. It seems that the empirical LTT is below the lower bound of the confidence interval for a larger part of the history than for the previous case of Cetacea.

```
sum(points_in_cetacea)/length(points_in_cetacea)
```

```
## [1] 0.9886364
```

For Cetacea, 98.8% of the values of the empirical LTT fall in the a 95% confidence interval.

**2. Parnassiinae**

As for Cetacea, we used the function `simul.comb.shift()` to simulate trees under the best combination of the Parnassiinae analysis we kept (the second best combination).

Here the code to do so:

```
# to edit with updates (option for the number of simulations)
all_posteriors_parnassinae <- simul.comb.shift(n = 150000,
                                               phylo = tree_parnassiinae,
                                               sampling.fractions = f_parnassiinae,
                                               shift.res = shifts_parnassiinae_rmax2,
                                               combi = 2)

parnassinae_posterior_trees <- all_posteriors_parnassinae[1:500]
```

We can look at the number of tips in each part of the simulated trees to see whether simulated data are similar to the empirical tree of Parnassiinae.

```
ntip_by_groups <- lapply(parnassinae_posterior_trees,
                         function(x) table(sapply(strsplit(x$tip.label,
                                                           split = ""), "[[", 1)))
ntip_by_groups_df <- as.data.frame(do.call(rbind, ntip_by_groups))
ntip_by_groups_parnassiinae <-
  c(f_parnassiinae$sp_in[!is.na(f_parnassiinae$to_test)][c(1,2,6:9,11)],
                                 85-77)
par(mfrow = c(1,2), cex = 0.6)
boxplot(ntip_by_groups_df, las = 1, main = "Species richness by group",
        ylab = "Number of species", xlab = "Groups")
points(c(1:8), ntip_by_groups_parnassiinae, pch = 19, col = "red")
legend("topright", bty = "n", legend = "Empirical values", pch = 19,
       col = "red", cex = 1)

avgLeafDepI_Parnassiinae <- avgLeafDepI(tree_parnassiinae)
avgLeafDepI_Parnassiinae_posteriors <- sapply(all_posteriors_parnassinae, avgLeafDepI)
hist(avgLeafDepI_Parnassiinae_posteriors, bty = "o",
     main = "Distribution of average leaf depth index",
     xlab = "Average leaf depth", las = 1)
abline(v = avgLeafDepI_Parnassiinae, col = "red")
legend("topright", bty = "n", legend = "Empirical index value", lty = 1,
       col = "red", cex = 1)
```

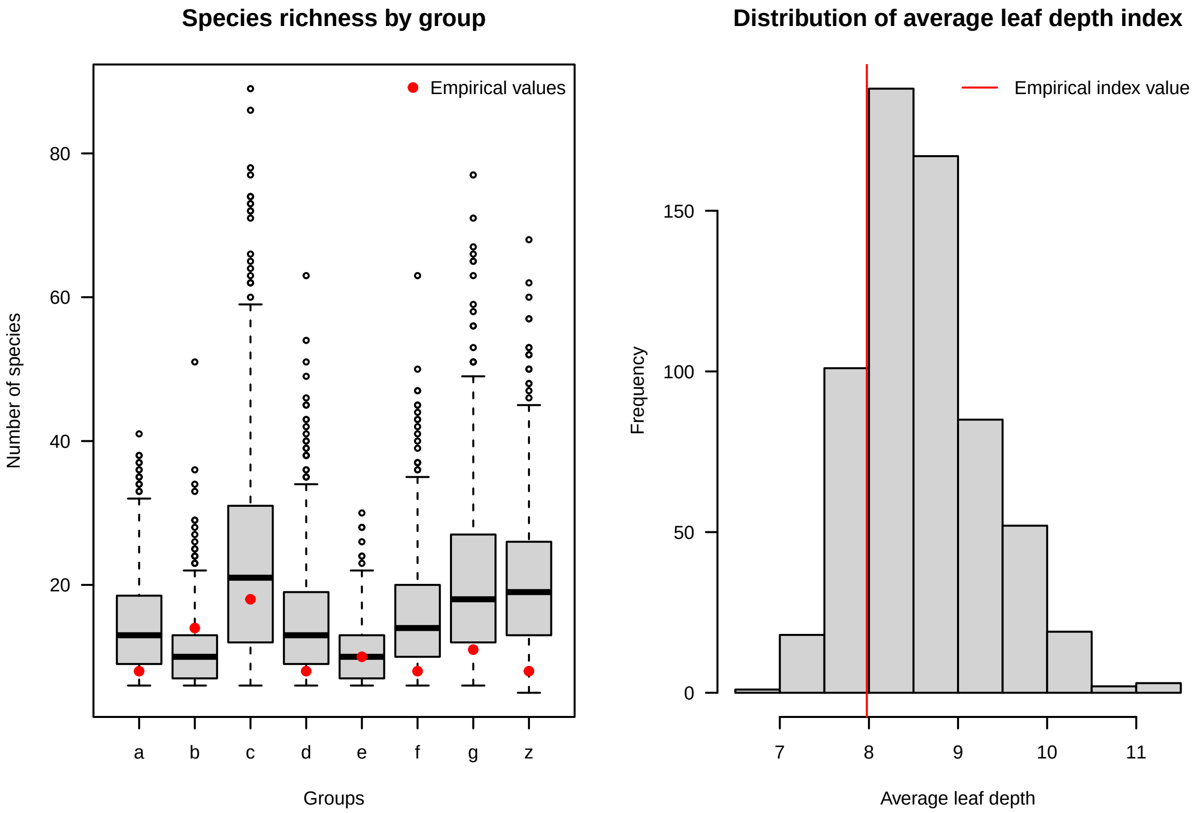

We can see that the empirical values fall less often into the interquartile of simulated data (groups d, f, g and z the backbone). However, the empirical value of the average leaf depth index falls into the distribution of simulated values.

We then measure the proportion of the values of the empirical LTT that are contained in a 95% confidence interval obtained from the LTT of simulated trees.

```
# Parnassiinae
ltt_parnassiinae <- ltt(tree_parnassiinae, plot = F)
ltt_parnassiinae_df <- data.frame(times = ltt_parnassiinae$times,
                                  ltt = ltt_parnassiinae$ltt)

parnassiinae_posterior_trees_mp <- as.multiPhylo(parnassinae_posterior_trees[[1]])
for(i in 2:length(parnassinae_posterior_trees)){
  parnassiinae_posterior_trees_mp[[i]] <- parnassinae_posterior_trees[[i]]
}

ltt95_CI_parnassiinae <- ltt95(parnassiinae_posterior_trees_mp,
                               log = T)
lines(ltt_parnassiinae_df$times, ltt_parnassiinae_df$ltt, type = "s", col = "red")
mtext(text = "Lineage-Through-Time plot for Parnassiinae", side = 3, line = 0.5)
legend("topleft",
       legend = c("95% of the distribution of simulated trees around the median",
                  "Median of simulated trees",
                  "Empirical data"),
       lty = c(3,1,1), lwd = c(1,2,1), col = c("black","black","red"),
       bty = "n", cex = 0.8)
```

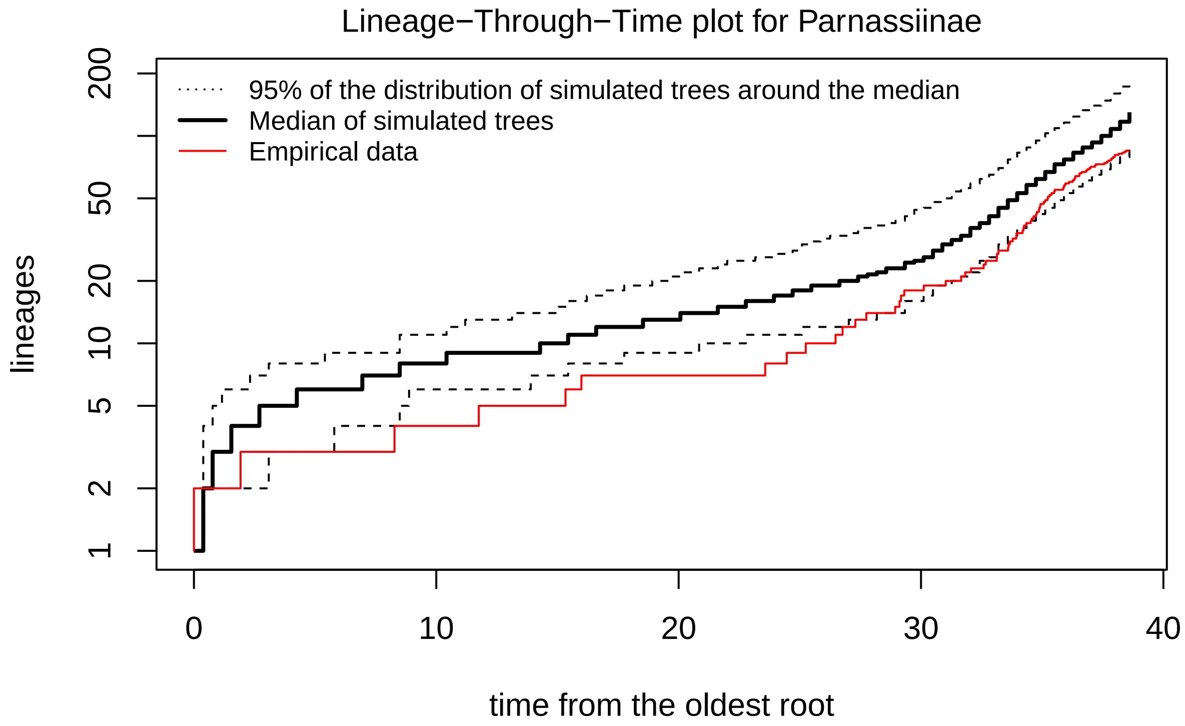

Figure 2: Lineages-Through-Time plot of Parnassiinae posterior predictives analyses. y-axis is log-scaled.

```r
ltt95_CI_parnassiinae_df <- as.data.frame(ltt95_CI_parnassiinae[,c("time",
                                                                   "low(lineages)",
                                                                   "high(lineages)")])

points_in_parnassiinae <- c()
for(i in 1:nrow(ltt_parnassiinae_df)){
  int_max <- sort(ltt95_CI_parnassiinae_df$time[
    ltt95_CI_parnassiinae_df$time > ltt_parnassiinae_df$times[i]][1])
  int_min <- sort(ltt95_CI_parnassiinae_df$time[
    ltt95_CI_parnassiinae_df$time <= ltt_parnassiinae_df$times[i]],
                  decreasing = T)[1]
  ltt_min <- ltt95_CI_parnassiinae_df$`low(lineages)`[
    ltt95_CI_parnassiinae_df$time == int_min]
  ltt_max <- ltt95_CI_parnassiinae_df$`high(lineages)`[
    ltt95_CI_parnassiinae_df$time == int_max]

  points_in_parnassiinae[i] <- ifelse(ltt_min <= ltt_parnassiinae_df$ltt[i] &
                                        ltt_parnassiinae_df$ltt[i] <= ltt_max,
                                      T, F)
}
```

The plot above shows the median LTT plot of the posterior trees, the confidence interval at 95% in doted lines and the empirical LTT in red. It seems that the empirical LTT is below the lower bound of the confidence interval for a larger part of the history than for the previous case of Cetacea.

```r
sum(points_in_parnassiinae)/length(points_in_parnassiinae)
```

```
## [1] 0.8488372
```

Indeed, 82.6% of the values of the empirical LTT fall in the a 95% confidence interval. This result echoes to the comparison of the number of species by groups, few empirical groups have low diversity compared to their distribution of rate values. We can also link this to the fact that, in the shift analysis, three combinations of shifts were equivalent for this phylogeny. It might highlight that there is more uncertainty on the location of shifts for this case than for Cetacea.

##### 3. Cycadales

Finally, we used the same procedure and analyses for the best combination of the Parnassiinae analysis we kept (the second best combination).

Here the code to do so:

```r
# to edit with updates (option for the number of simulations)
all_posteriors_cycads <- simul.comb.shift(phylo = tree_cycads,
                                          sampling.fractions = f_cycads,
                                          shift.res = shift_cycads,
                                          combi = 1)

cycads_posterior_trees <- all_posteriors_cycads[1:500]
```

We can look at the number of tips in each part of the simulated trees to see whether simulated data are similar to the empirical tree of Cycadales.

```
ntip_by_groups <- lapply(cycads_posterior_trees,
                         function(x) table(sapply(strsplit(x$tip.label,
                                                  split = ""), "[[", 1)))
ntip_by_groups_df <- as.data.frame(do.call(rbind, ntip_by_groups))
ntip_by_groups_cycads <- c(f_cycads$sp_in[!is.na(f_cycads$to_test)][c(1,3:6)],
                           231-217)

par(mfrow = c(1,2), cex = 0.6)
boxplot(ntip_by_groups_df, las = 1, main = "Species richness by group",
        ylab = "Number of species", xlab = "Groups")
points(c(1:6), ntip_by_groups_cycads, pch = 19, col = "red")
legend("topright", bty = "n", legend = "Empirical values",
       pch = 19, col = "red", cex = 1)

avgLeafDepI_cycads <- avgLeafDepI(tree_cycads)
avgLeafDepI_cycads_posteriors <- sapply(cycads_posterior_trees, avgLeafDepI)
hist(avgLeafDepI_cycads_posteriors,
     main = "Distribution of average leaf depth index",
     xlab = "Average leaf depth", las = 1)
abline(v = avgLeafDepI_cycads, col = "red")
legend("topright", bty = "n", legend = "Empirical index value", lty = 1,
       col = "red", cex = 1)
```

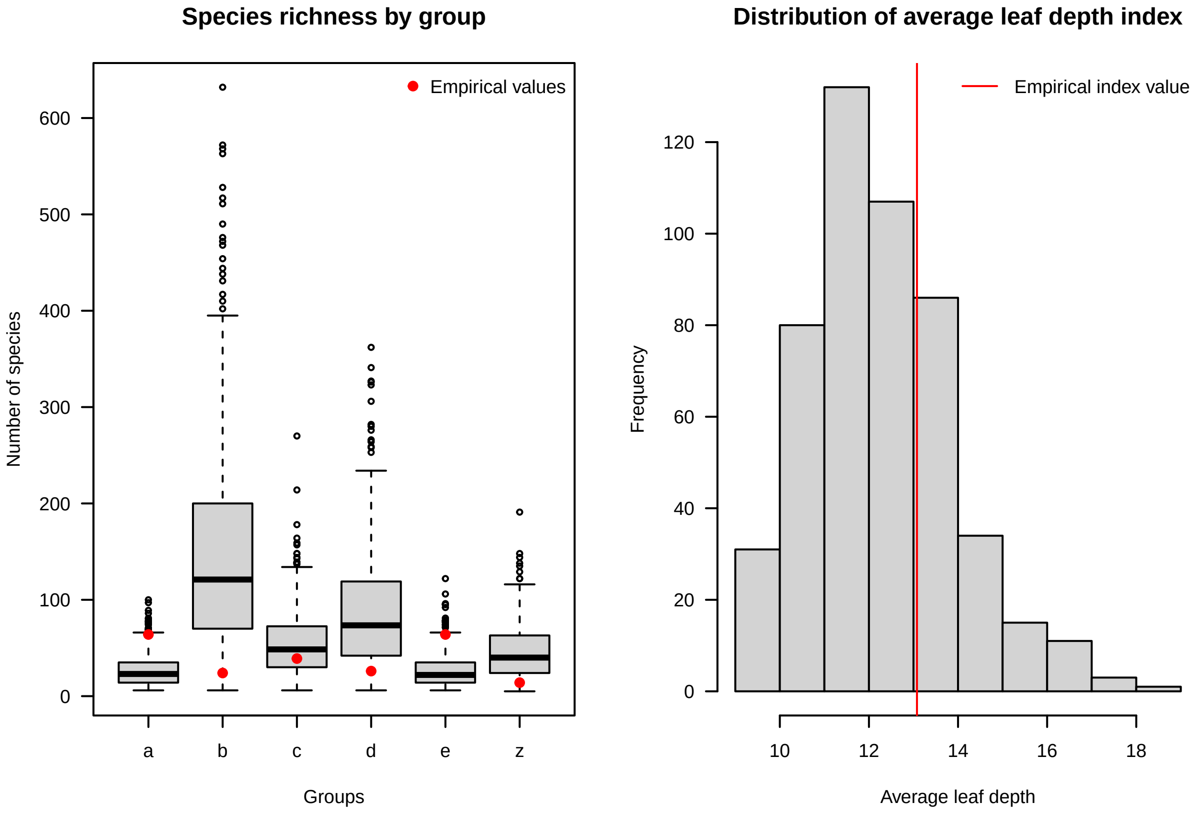

We can see that the empirical values fall into the interquartile of simulated data only for the group c, the other groups having more extreme empirical values of the number of tips. As previously, the empirical value of the average leaf depth index falls into the distribution of simulated values.

We then measure the proportion of the values of the empirical LTT that are contained in a 95% confidence interval obtained from the LTT of simulated trees.

```
ltt_cycads <- ltt(tree_cycads, plot = F)
ltt_cycads_df <- data.frame(times = round(ltt_cycads$times, 4),
                            ltt = ltt_cycads$ltt)

cycads_posterior_trees_mp <- as.multiPhylo(cycads_posterior_trees[[1]])
for(i in 2:length(cycads_posterior_trees)){
  cycads_posterior_trees_mp[[i]] <- cycads_posterior_trees[[i]]
}

ltt95_CI_cycads <- ltt95(cycads_posterior_trees_mp,
                         log = T)
lines(ltt_cycads_df$times, ltt_cycads_df$ltt, type = "s", col = "red")
mtext(text = "Lineage-Through-Time plot for Cycadales", side = 3, line = 0.5)

ltt95_CI_cycads_df <- as.data.frame(ltt95_CI_cycads[,c("time",
                                                       "low(lineages)",
                                                       "high(lineages)")])
ltt95_CI_cycads_df$time <- round(ltt95_CI_cycads_df$time, 4)

points_in_cycads <- c()
for(i in 1:nrow(ltt_cycads_df)){
  int_max <- sort(ltt95_CI_cycads_df$time[
    ltt95_CI_cycads_df$time >= ltt_cycads_df$times[i]][1])
  int_min <- sort(ltt95_CI_cycads_df$time[
    ltt95_CI_cycads_df$time <= ltt_cycads_df$times[i]],
                  decreasing = T)[1]
  ltt_min <- ltt95_CI_cycads_df$`low(lineages)`[
    ltt95_CI_cycads_df$time == int_min]
  ltt_max <- ltt95_CI_cycads_df$`high(lineages)`[
    ltt95_CI_cycads_df$time == int_max]

  points_in_cycads[i] <- ifelse(ltt_min <= ltt_cycads_df$ltt[i] &
                                  ltt_cycads_df$ltt[i] <= ltt_max, T, F)
}
legend("topleft",
       legend = c("95% of the distribution of simulated trees around the median",
                  "Median of simulated trees",
                  "Empirical data"),
       lty = c(3,1,1), lwd = c(1,2,1), col = c("black","black","red"),
       bty = "n", cex = 0.8)
```

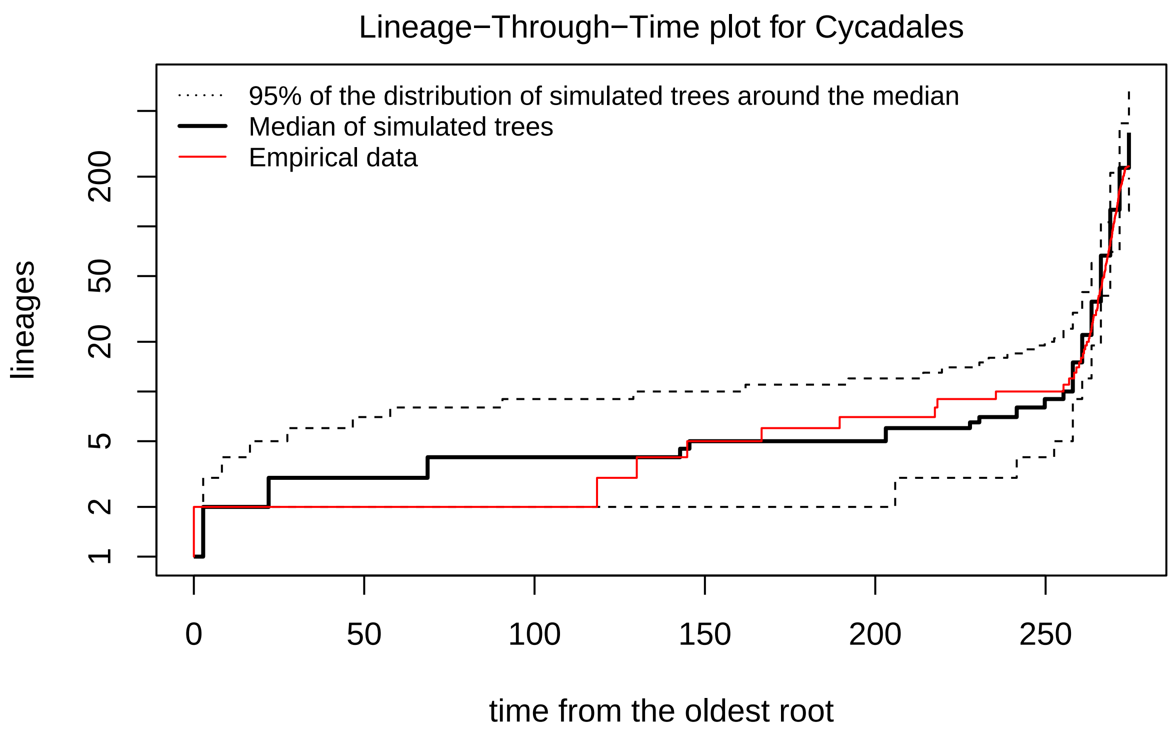

The plot above shows the median LTT plot of the posterior trees, the confidence interval at 95% in doted lines and the empirical LTT in red. The empirical LTT seems to be inside the confidence interval for almost all the history of the clade.

```
sum(points_in_cycads)/length(points_in_cycads)
```

```
## [1] 0.9956897
```

Indeed, 99.6% of the values of the empirical LTT fall in the a 95% confidence interval.

#### IV - Making plots

This part explains how to represent the shifts of diversification on the phylogeny, the diversification rates of the backbone through time and the corresponding paleodiversity dynamics.

##### 1. Cetacea

The following code shows how to reproduce the Figure 5 and Figure S1 for the analyses of Cetacea.

With `backbone.option = "crown.shift"` (Figure 5):

```
# diversification rates through time
rates <- div.rates(phylo = Cetacea,
                   shift.res = shifts_cetacea,
                   combi = 1, part = "all")
```

```
group_colors <- c(brewer.pal(8, "Dark2")[1:length(prob_dtt_cetacea$subclades)],
                  c("black"))

# plot
par(mar = c(6,2,4,0), xpd = T)
layout(matrix(c(1,1,2,2,
                1,1,2,2,
                1,1,3,3,
                1,1,3,3), 4, 4, byrow = T))
# phylogeny
phylo_p <- plot.phylo.comb(phylo = Cetacea, data = taxo_cetacea,
                           sampling.fractions = f_cetacea,
                           shift.res =  shifts_cetacea,
                           combi = 1, tip.color = "white",
                           label.offset = 0.3,
                           main = "",
                           edge.width = 1, cex = 0.4,
                           tested_nodes = T, text.cex = 0.8, pch.cex = 1)

add.gts(thickness = -6, quaternary = T, cex = 1, is.phylo = T,
        names = c("Q.", "P.", "Miocene", "Oligocene", "E."), xpd.x = F)

mtext("Time (Myrs)", side = 1, line = 3.5, cex = 0.7, at = 17)

par(new = T, usr = par("usr"), mar = c(6,2,4,0))

phylo_p <- plot.phylo.comb(phylo = Cetacea, data = taxo_cetacea,
                           sampling.fractions = f_cetacea,
                           shift.res =  shifts_cetacea,
                           combi = 1, label.offset = 0.3,
                           main = "Phylogeny of Cetacea\n(option crown.shift)",
                           edge.width = 1, cex = 0.4,
                           tested_nodes = T, text.cex = 0.8, pch.cex = 1)

legend(-1.5, 91, legend = c("Tested nodes"), xpd = T,
       pch = 21, col = "black", pt.bg = c("red"),
       bty = "n", cex = 0.8)

# rates
root.age <- max(branching.times(Cetacea))
time <- -c(root.age,floor(root.age):0)

par(mar = c(6,4,4,4), xpd = T)
plot(time, rates[[length(rates)]][1,], type = "n", lwd = 1,
     ylim = c(0,1.2),
     col = "blue", bty = "n",
     cex.lab = 1, cex.axis = 1, cex.main = 1,
     las = 1, axes = F, xlab = "", ylab = "",
     main = "",
     col.main = group_colors[length(rates)])

add.gts(thickness = -0.3, quaternary = T, cex = 1,
        names = c("Q.", "P.", "Miocene", "Oligocene", "E."),
```

```
        xpd.x = F)
mtext("Time (Myrs)", side = 1, line = 3.5, cex = 0.7)

par(new = T)
plot(time, rates[[length(rates)]][1,], type = "l", lwd = 1,
     ylim = c(0,1.5),
     col = "blue", bty = "n",
     cex.lab = 1, cex.axis = 1, cex.main = 1.3,
     las = 1, xaxt = "n",
     xlab = "", ylab = "Rates (Event / Lineage / Myr)",
     main = "Diversification rates of the backbone")
lines(time,
      rates[[length(rates)]][2,], type = "l", lwd = 1,
      col = "red")
legend(-35, 1.5, legend = c("Speciation rate", "Extinction rate"),
       col = c("blue", "red"), lty = 1, bty = "n", lwd= 1, cex = 1)

# Paleodiversity dynamic with probabilistic approach
par(xpd = T)
plot_prob_dtt(mat = prob_dtt_cetacea$backbones[[1]], lwd = 1, grain = 0.05,
              xlab = "", ylab = "", col.mean = "white", bty = "n",
              main = "Diversity through time of Cetacea\n(probabilistic approach)",
              plot.prob = F,
              ylim = c(0, 1000), las = 1, plot.bound = F, col.bound = "white",
              lty.bound = 2, axes = F, add.present = T)

add.gts(thickness = -250, quaternary = T, cex = 1,
        names = c("Q.", "P.", "Miocene", "Oligocene", "E."),
        xpd.x = F)

par(new = T)
plot_prob_dtt(mat = prob_dtt_cetacea$backbones[[1]], lwd = 1, grain = 0.05,
              xlab = "", ylab = "", col.mean = "black", bty = "n",
              ylim = c(0, 1000), las = 1, plot.bound = T, plot.prob = F,
              col.bound = "black", lty.bound = 2, xaxt = "n", add.present = T)

par(new = T)
plot_prob_dtt(mat = prob_dtt_cetacea$backbones[[1]], lwd = 1, grain = 0.05,
              col.mean = scales::alpha("white", 0), bty = "n",
              ylim = c(0, 40), las = 1, plot.bound = F, plot.mean = T,
              plot.prob = F, axes = F, xlab = "", ylab = "", add.present = T)

axis(side = 4, las = 1, xpd = TRUE, cex.axis = 1, line = 0)

mtext("Number of species (subclades)", side = 4, line = 2, cex = 0.7)
for(i in 1:length(prob_dtt_cetacea$subclades)){
  plot_prob_dtt(mat = prob_dtt_cetacea$subclades[i][[1]], lwd = 1,
                plot.prob = F, grain = 0.1, add = T,
                col.mean = group_colors[i], add.present = T)
}

mtext("Number of Species", side = 2, line = 3, cex = 0.7)
mtext("Time (Myrs)", side = 1, line = 3.5, cex = 0.7)
```

```r
legend("topleft", lty = 2, legend = "Confidence interval at 95%", bty = "n", cex = 0.8)
```

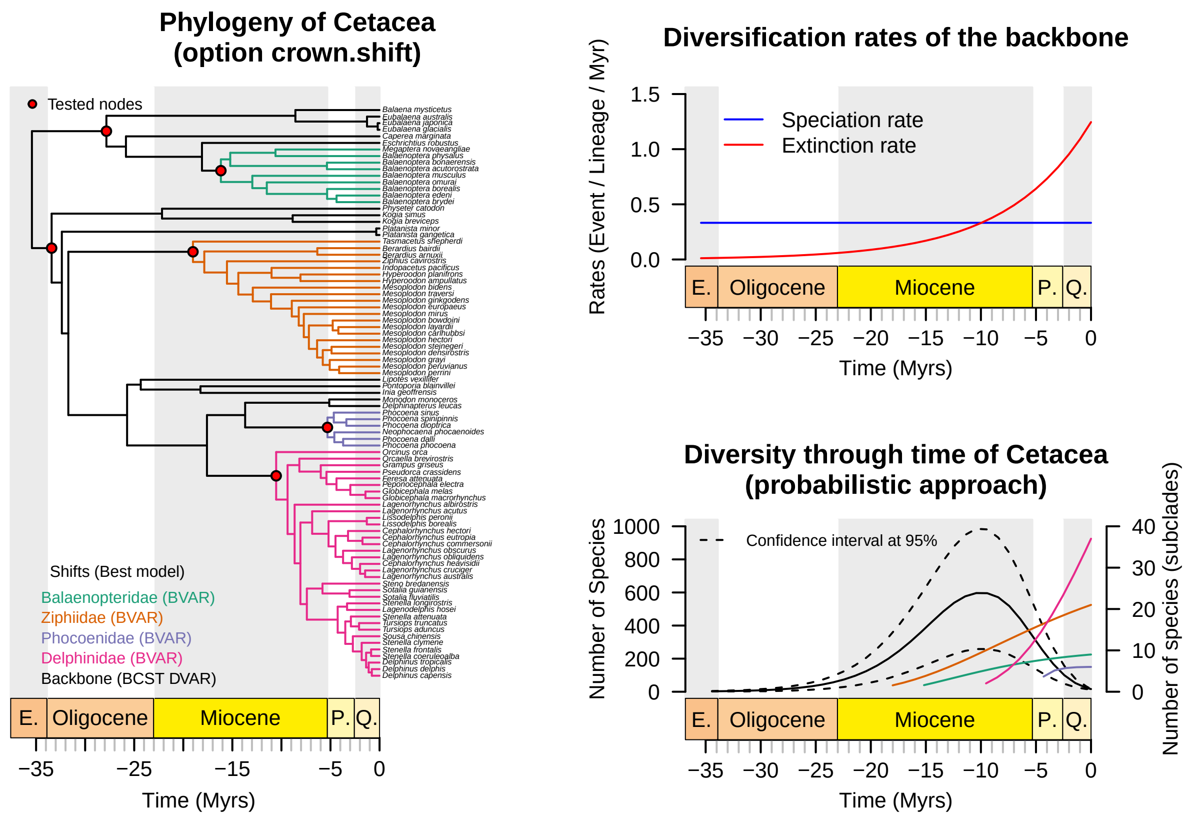

Figure 3: Paleodiversity dynamics of Cetacea (with stems included into the backbone analysis)

With `backbone.option = "stem.shift"` (Figure S1):

```
diversity <- paleodiv(phylo = Cetacea,
                      data = taxo_cetacea, sampling.fractions = f_cetacea,
                      shift.res = shifts_cetacea_stem.shift,
                      combi = 1, split.div = T)

rates <- div.rates(phylo = Cetacea,
                   shift.res = shifts_cetacea_stem.shift,
                   combi = 1, part = "all")

group_colors <- c(brewer.pal(8,"Dark2")[1:length(prob_dtt_cetacea_stem$subclades)],
                  c("black"))

# plot
par(mar = c(6,2,4,0), xpd = T)
layout(matrix(c(1,1,2,2,
                1,1,2,2,
                1,1,3,3,
                1,1,3,3), 4, 4, byrow = T))
# phylogeny
phylo_p <- plot.phylo.comb(phylo = Cetacea,
                           data = taxo_cetacea,
                           sampling.fractions = f_cetacea,
                           shift.res = shifts_cetacea_stem.shift,
                           tip.color = "white",
                           label.offset = 0.3,
                           main = "",
                           combi = 1, edge.width = 1, cex = 0.4, tested_nodes = F,
                           text.cex = 0.8, pch.cex = 1,
                           backbone.option = "stem.shift")

add.gts(thickness = -5.5, quaternary = T, cex = 1, is.phylo = T,
        names = c("Q.", "Pl.", "Miocene", "Oligocene", "E."), xpd.x = F)

mtext("Time (Myrs)", side = 1, line = 3.5, cex = 0.7, at = 17)

par(new = T, mar = c(6,2,4,0))
phylo_p <- plot.phylo.comb(phylo = Cetacea,
                           data = taxo_cetacea,
                           sampling.fractions = f_cetacea,
                           shift.res = shifts_cetacea_stem.shift,
                           backbone.option = "stem.shift",
                           main = "Phylogeny of Cetacea\n(option stem.shift)",
                           label.offset = 0.3,
                           combi = 1, edge.width = 1, cex = 0.4,
                           tested_nodes = F, text.cex = 0.8, pch.cex = 1)

# rates
root.age <- max(branching.times(Cetacea))
time <- -c(root.age,floor(root.age):0)

par(mar = c(6,4,4,4), xpd = T)
plot(time, rates[[length(rates)]][1,],
```

```
     type = "n", lwd = 1, ylim = c(0,1.2),
     col = "blue", bty = "n",
     cex.lab = 1, cex.axis = 1, cex.main = 1.3,
     las = 1, axes = F, xlab = "", ylab = "",
     main = "Diversification rates of the backbone",
     col.main = group_colors[length(rates)])

add.gts(thickness = -0.30, quaternary = T, cex = 1,
        names = c("Q.", "Pl.", "Miocene", "Oligocene", "E."), xpd.x = F)
mtext("Time (Myrs)", side = 1, line = 3.5, cex = 0.7)

par(new = T)
plot(time, rates[[length(rates)]][1,],
     type = "l", lwd = 1, ylim = c(0,1.2),
     col = "blue", bty = "n",
     cex.lab = 1, cex.axis = 1, cex.main = 1,
     las = 1, xaxt = "n",
     xlab = "", ylab = "Rates (Event / Lineage / Myr)",
     main = "")
lines(time, rates[[length(rates)]][2,],
      type = "l", lwd = 1, col = "red")
legend(-25, 1.2, legend = c("Speciation rate", "Extinction rate"),
       col = c("blue", "red"), lty = 1, bty = "n", lwd= 1, cex = 1)

# Paleodiversity dynamic with probabilistic approach
par(xpd = T)
plot_prob_dtt(mat = prob_dtt_cetacea_stem$backbones[[1]],
              lwd = 1, grain = 0.05, col.mean = "black", bty = "n",
              ylim = c(0, 700), las = 1, plot.bound = F, col.bound = "black",
              lty.bound = 2, axes = F,
              plot.prob = F,
              main = "Diversity-through-time plot\n(probabilistic approach)",
              xlab = "", ylab = "")

add.gts(thickness = -150, quaternary = T, cex = 1,
        names = c("Q.", "P.", "Miocene", "Oligocene", "E."),
        xpd.x = F)

par(new = T)
plot_prob_dtt(mat = prob_dtt_cetacea_stem$backbones[[1]],
              lwd = 1, grain = 0.05, col.mean = "black",
              bty = "n", xlab = "", ylab = "",
              ylim = c(0, 700), las = 1, plot.bound = T,
              plot.prob = F, col.bound = "black", lty.bound = 2, xaxt = "n")

par(new = T)
plot_prob_dtt(mat = prob_dtt_cetacea_stem$backbones[[1]],
              lwd = 1, grain = 0.05, col.mean = scales::alpha("white", 0), bty = "n",
              ylim = c(0, 40), las = 1, plot.bound = F, plot.mean = T,
              plot.prob = F, axes = F, xlab = "", ylab = "")

axis(side = 4, las = 1, xpd = TRUE, cex.axis = 1, line = 0)
```

```r
mtext("Number of species (subclades)", side = 4, line = 2, cex = 0.7)
for(i in 1:length(prob_dtt_cetacea_stem$subclades)){
  plot_prob_dtt(mat = prob_dtt_cetacea_stem$subclades[i][[1]],
                lwd = 1, plot.prob = F, grain = 0.1, add = T,
                col.mean = group_colors[i])
}

mtext("Number of Species", side = 2, line = 3, cex = 0.7)
mtext("Time (Myrs)", side = 1, line = 3.5, cex = 0.7)
legend("topleft", lty = 2, legend = "Confidence interval at 95%", bty = "n")
```

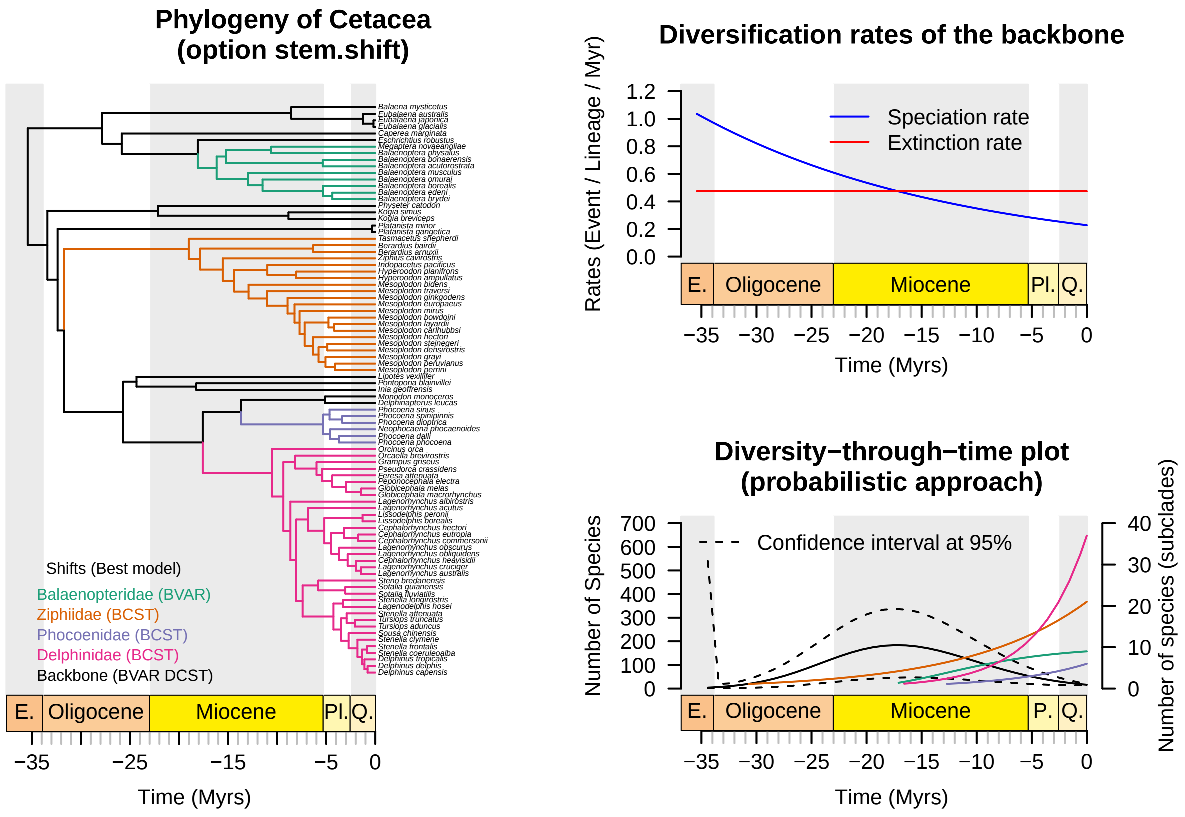

Figure 4: Paleodiversity dynamics of Cetacea (with stems included into subclade analyses)

#### 2. Vangidae

The following code shows how to reproduce the Figure 6 for the Vangidae analysis from the main manuscript.

```r
# diversification rates through time
rates <- div.rates(phylo = tree_vangidae, shift.res = shift.vangidae,
                   combi = 1, part = "all")

# plot
par(mar = c(6,2,4,0), xpd = T)
```

```r
layout(matrix(c(1,1,2,2,
                1,1,2,2,
                1,1,3,3,
                1,1,3,3), 4, 4, byrow = T))

phylo_p <- plot.phylo.comb(phylo = tree_vangidae,
                           data = taxo_vangidae,
                           sampling.fractions = f_vangidae,
                           shift.res = shift.vangidae,
                           label.offset = 0.3,
                           tip.color = "white",
                           main = "Phylogeny of Vangidae",
                           combi = 1, edge.width = 1, cex = 0.5, tested_nodes = T)

# phylogeny
plot_dim <- par("usr")
par(new = T, usr = plot_dim,  xpd = T)

add.gts(thickness = -2, quaternary = T, cex = 1, is.phylo = T,
        names = c("Q.", "P.", "Miocene"), xpd.x = F)
mtext(text = "Time (Myrs)", side = 1, line = 3.5, cex = 0.7, at = 9)

par(new = T, usr = plot_dim, mar = c(6,2,4,0))
phylo_p <- plot.phylo.comb(phylo = tree_vangidae,
                           data = taxo_vangidae,
                           sampling.fractions = f_vangidae,
                           shift.res = shift.vangidae, leg = F,
                           label.offset = 0.3,
                           main = "Phylogeny of Vangidae",
                           combi = 1, edge.width = 1, cex = 0.5,
                           tested_nodes = T, pch.cex = 1.5)
legend(0,34, legend = "Tested nodes", pch = 21, col = "black",
       pt.bg = "red", bty = "n", cex = 1)

# diversification rates through time
root.age <- max(branching.times(tree_vangidae))
time <- -c(root.age,floor(root.age):0)
par(mar = c(6,4,4,4), xpd = T)
plot(time, rates[[length(rates)]][1,], xlim = c(-20,0),
     type = "l", lwd = 2, ylim = c(0,0.4),
     col = "blue", bty = "n", cex.lab = 1.5, cex.axis = 1.4,
     cex.main = 1.3, las = 1, axes = F,
     xlab = "", ylab = "",
     main = "Diversification rates of the whole phylogeny")
lines(1:length(rates[[length(rates)]][1,]),
      rates[[length(rates)]][2,], type = "l", lwd = 2,
      col = "red")

add.gts(thickness = -0.1, quaternary = T, cex = 1,
        names = c("Q.", "P.", "Miocene"), xpd.x = F)
mtext("Time (Myrs)", side = 1, line = 3.7, cex = 0.7)
```

```r
par(new = T)
plot(time, rates[[length(rates)]][1,],
     type = "l", lwd = 1, ylim = c(0,0.4),
     col = "blue", bty = "n", cex.lab = 1, cex.axis = 1,
     cex.main = 1, las = 1, xaxt = "n",
     xlab = "", ylab = "",
     main = "")
mtext("Rates (Event / Lineage / Myr)", side = 2, line = 2.5, cex = 0.7)
lines(1:length(rates[[length(rates)]][1,]),
      rates[[length(rates)]][2,], type = "l", lwd = 1,
      col = "red")
legend(-12, 0.44, legend = c("Speciation rate"),
       col = c("blue"), lty = 1, bty = "n", lwd= 1, cex = 1)

# Paleodiversity dynamic with probabilistic approach
plot_prob_dtt(mat = prob_dtt_vangidae[[1]], lwd = 1, grain = 0.05, axes = F,
              plot.prob = F,
              xlab = "", ylab = "", col.mean = "black", bty = "n", ylim = c(0,35),
              las = 1, plot.bound = T, col.bound = "black", lty.bound = 2,
              main = "Diversity through time of Vangidae\n(probabilistic approach)")

par(new = T)
present = as.numeric(colnames(prob_dtt_vangidae[[1]])[
  length(colnames(prob_dtt_vangidae[[1]]))])
root.age = as.numeric(colnames(prob_dtt_vangidae[[1]])[1])

add.gts(thickness = -8, quaternary = T, cex = 1,
              names = c("Q.", "P.", "Miocene"), xpd.x = F)

par(new = T)
plot_prob_dtt(mat = prob_dtt_vangidae[[1]], lwd = 1, grain = 0.05,
              col.mean = "black", bty = "n", xlab = "", ylab = "",
              las = 1, plot.bound = T, plot.prob = F, ylim = c(0,35),
              col.bound = "black", lty.bound = 2, xaxt = "n")

mtext("", side = 3, line = 1, cex = 0.7)
mtext("Number of Species", side = 2, line = 2.5, cex = 0.7)
mtext("Time (Myrs)", side = 1, line = 3.5, cex = 0.7)
```

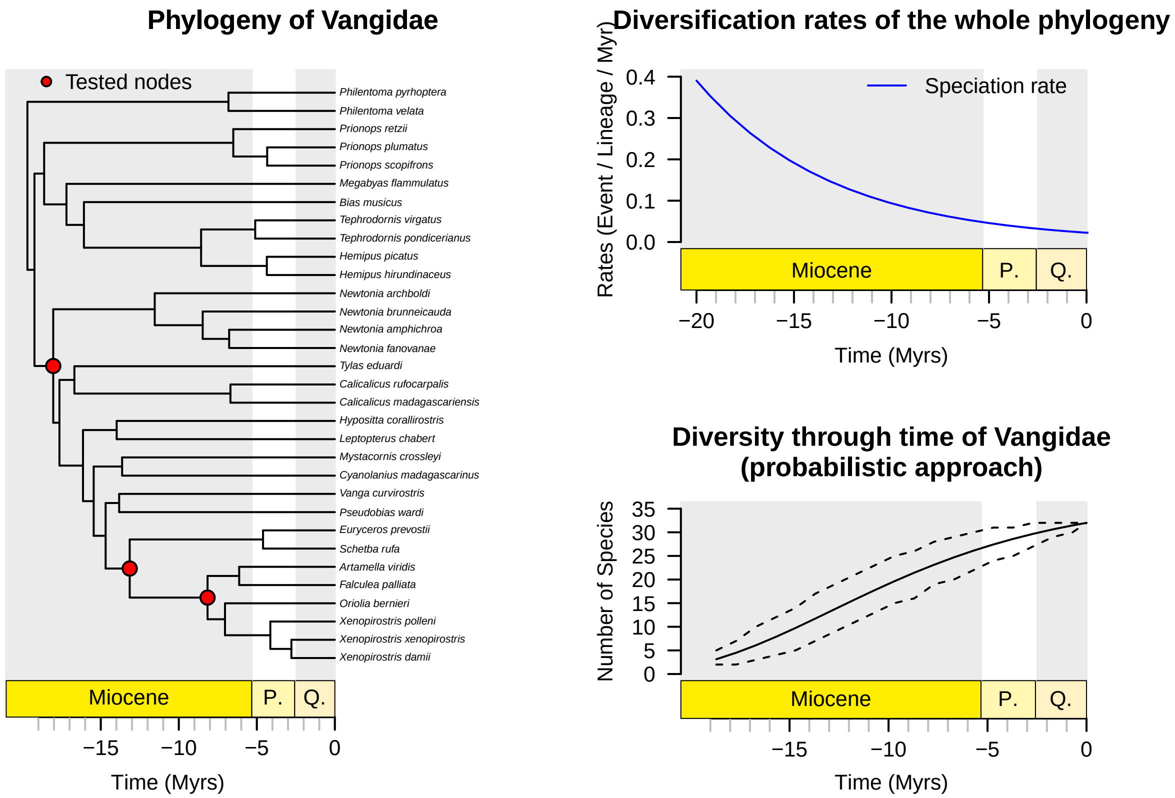

Figure 5: Paleodiversity dynamic of Vangidae

##### 3. Parnassiinae

The following code shows how to reproduce the Figure 7 for the Parnassiinae analyses from the main manuscript.

```
# diversification rates
rates <- div.rates(phylo = tree_parnassiinae,
                   shift.res = shifts_parnassiinae_rmax2,
                   combi = 2, part = "all")

group_colors <- c(brewer.pal(8,"Dark2")[1:length(prob_dtt_parnassiinae_rmax2$subclades)],
                  c("black"))

# plot
par(mar = c(6,2,4,0), xpd = T)
layout(matrix(c(1,1,2,2,
                1,1,2,2,
                1,1,3,3,
                1,1,3,3), 4, 4, byrow = T))

# phylogeny
phylo_p <- plot.phylo.comb(phylo = tree_parnassiinae,
                           data = taxo_parnassiinae,
                           sampling.fractions = f_parnassiinae,
                           shift.res = shifts_parnassiinae_rmax2,
                           tip.color = "white", main = "Phylogeny of Parnassiinae",
                           combi = 2, edge.width = 1, cex = 0.4,
                           label.offset = 0.3,
                           tested_nodes = T, text.cex = 0.8, pch.cex = 1)

add.gts(thickness = -5.3, quaternary = T, cex = 0.9, is.phylo = T,
        names = c("Q", "P", "Miocene", "Oligocene", "Eoc."), xpd.x = F)

mtext("Time (Myrs)", side = 1, line = 3.5, cex = 0.7, at = 17)

par(new = T, usr = par("usr"), mar = c(6,2,4,0))
phylo_p <- plot.phylo.comb(phylo = tree_parnassiinae,
                           data = taxo_parnassiinae,
                           sampling.fractions = f_parnassiinae,
                           shift.res = shifts_parnassiinae_rmax2,
                           combi = 2, edge.width = 1, cex = 0.4,
                           label.offset = 0.3,
                           tested_nodes = T, text.cex = 0.8, pch.cex = 1,
                           main = "Phylogeny of Parnassiinae")

legend(-1.7, 89, legend = c("Tested nodes"), xpd = T, pch = 21,
       col = "black", pt.bg = c("red"), bty = "n", cex = 1)

root.age <- max(branching.times(tree_parnassiinae))
time <- -c(root.age,floor(root.age):0)
# diversification rates through time
par(mar = c(6,4,4,4), xpd = T)
plot(time, rates[[length(rates)]][1,],
     type = "l", lwd = 2, ylim = c(1.5,2),
```

```r
     col = "blue", bty = "n", cex.lab = 1.5, cex.axis = 1.4,
     cex.main = 1.3, las = 1, axes = F,
     xlab = "", ylab = "",
     main = "Diversification rates of the backbone",
     col.main = group_colors[length(rates)])

add.gts(thickness = -0.12, quaternary = T, cex = 0.9,
        names = c("Q", "P", "Miocene", "Oligocene", "Eoc."), xpd.x = F)

mtext("Time (Myrs)", side = 1, line = 3.5, cex = 0.7)
par(new = T)
plot(time, rates[[length(rates)]][1,], type = "l", lwd = 1,
     ylim = c(1,2),
     col = "blue", bty = "n", cex.lab = 1, cex.axis = 1,
     cex.main = 1, las = 1, xaxt = "n",
     xlab = "", ylab = "",
     main = "")
lines(time, rates[[length(rates)]][2,],
      type = "l", lwd = 1, col = "red")
legend(-25, 10.1, legend = c("Speciation rate", "Extinction rate"),
       col = c("blue", "red"), lty = 1, bty = "n", lwd= 1, cex = 1)
mtext("Rates (Event / Lineage / Ma)", side = 2, line = 3, cex = 0.7)

# Paleodiversity dynamic with probabilistic approach
plot_prob_dtt(mat = prob_dtt_parnassiinae_rmax2$backbones[[1]], lwd = 1, grain = 0.05,
              col.mean = "black", bty = "n",
              ylim = c(0, 2000), las = 1,
              plot.bound = T, col.bound = "black", lty.bound = 2,
              plot.mean = T, plot.prob = F, axes = F, xlab = "", ylab = "",
              main = "Diversity through time of Parnassiinae\n(probabilistic approach)")

add.gts(thickness = -430, quaternary = T, cex = 0.9,
        names = c("Q", "P", "Miocene", "Oligocene", "Eoc."),
        xpd.x = F)

par(new = T)
plot_prob_dtt(mat = prob_dtt_parnassiinae_rmax2$backbones[[1]],
              lwd = 1, grain = 0.05, col.mean = "black", bty = "n",
              ylim = c(0, 2000), plot.prob = F, xlab = "", ylab = "",
              las = 1, plot.bound = T, col.bound = "black",
              lty.bound = 2, xaxt = "n")

par(new = T)
plot_prob_dtt(mat = prob_dtt_parnassiinae_rmax2$backbones[[1]],
              lwd = 1, grain = 0.05, col.mean = scales::alpha("white", 0),
              bty = "n",  ylim = c(0, 20), las = 1, plot.bound = F, plot.mean = T,
              plot.prob = F, axes = F, xlab = "", ylab = "")

axis(side = 4, las = 1, xpd = TRUE, cex.axis = 1, line = 0)
mtext("Number of species (subclades)", side = 4, line = 2, cex = 0.7)
for(i in 1:length(prob_dtt_parnassiinae_rmax2$subclades)){
  plot_prob_dtt(mat = prob_dtt_parnassiinae_rmax2$subclades[i][[1]],
                lwd = 1, plot.prob = F, grain = 0.1, add = T,
```

```
                    col.mean = group_colors[i])
}

mtext("Number of Species", side = 2, line = 3, cex = 0.7)
mtext("Time (Myrs)", side = 1, line = 3.5, cex = 0.7)
legend("topleft", lty = 2, legend = "Confidence interval at 95%", bty = "n")
```

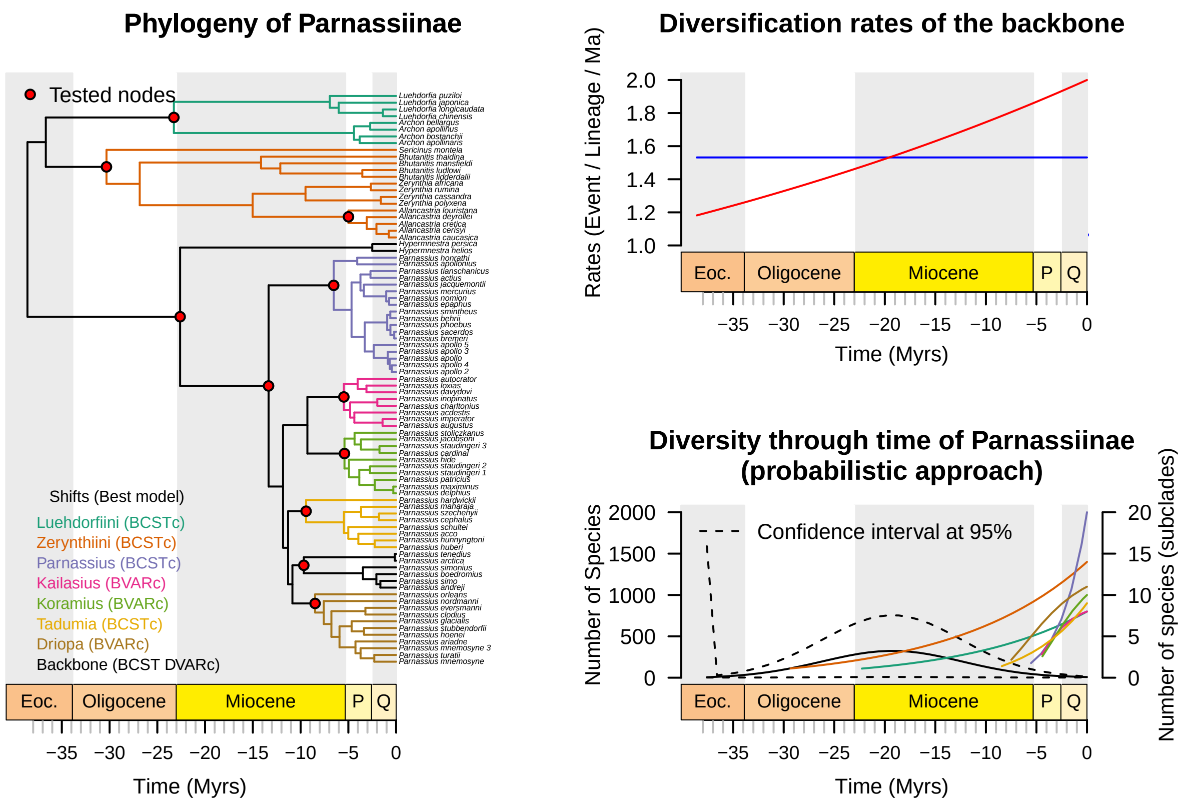

Figure 6: Paleodiversity dynamics of Parnassiinae

#### 4. Cycadales

The following code shows how to reproduce the Figure 8 for analyses of Cycadales from the main manuscript.

First, we calculate the confidence interval and the paleodiversity with both the deterministic and the probabilistic approach.

Now that estimates have been calculated, we can represent the Figure 8 summarising the results for Cycadales.

```
# > plot ####
par(mar = c(4,4,3,2), xpd = T)
layout(matrix(c(1,1,2,2,
                1,1,2,2,
                1,1,3,3,
```

```
                 1,1,3,3,
                 1,1,4,4,
                 1,1,4,4), 6, 4, byrow = T))

# phylogeny
plot.phylo.comb(phylo = tree_cycads,
                data = taxo_cycads,
                sampling.fractions = f_cycads,
                shift.res = shift_cycads, show.tip.label = F, cex.main = 1,
                combi = 1, edge.width = 0.7,
                tip.color = "white", main = "",
                cex = 1, tested_nodes = T, text.cex = 1, pch.cex = 1)

par(new = T)
add.gts(thickness = -12, quaternary = T, cex = 0.4, is.phylo = T,
              names = c("", "", "M", "O", "E", "P", "UC",
                        "LC", "UJ","MJ", "LJ", "T", "P"),
              time.interval = 10, padj = -4, xpd.x = F)
mtext(text = "Time (Myrs)", side = 1, line = 2.5, cex = 0.4)

par(new = T, usr = par("usr"), mar = c(6,4,3,2), xpd = T)

plot.phylo.comb(phylo = tree_cycads,
                data = taxo_cycads,
                sampling.fractions = f_cycads,
                shift.res = shift_cycads, show.tip.label = F, cex.main = 1,
                combi = 1, edge.width = 0.7,
                main = "Phylogeny of Cycadales",
                cex = 1, tested_nodes = T, text.cex = 1, pch.cex = 1)

legend(-1.5, 240, legend = c("Tested nodes"), xpd = T, pch = 21,
       col = "black", pt.bg = c("red"), bty = "n", cex = 0.8)

# diversification rates through time
root.age <- max(branching.times(tree_cycads))
time <- -c(root.age,floor(root.age):0)

par(mar = c(4,4,3,4), xpd = T)
plot(time, rates_cycads[[length(rates_cycads)]][1,],
     type = "l", lwd = 1, #ylim = c(0,2),
     col = "blue", bty = "n", cex.lab = 0.6, cex.axis = 0.6,
     cex.main = 1, las = 1, axes = F,
     xlab = "", ylab = "",
     main = "Diversification rates of the backbone")

add.gts(thickness = -0.07, quaternary = T, is.phylo = F, cex = 0.5,
        names = c("", "", "M", "Ol", "E", "Pa", "UC",
                  "LC", "UJ","MJ", "LJ", "T", "P"),
        time.interval = 10, padj = -3, xpd.x = F)

mtext(text = "Time (Myrs)", side = 1, line = 2.5, cex = 0.5)

par(new = T)
```

```r
plot(time, rates_cycads[[length(rates_cycads)]][1,],
     type = "l", lwd = 1, #ylim = c(0,2),
     col = "blue", bty = "n", cex.lab = 0.6, cex.axis = 0.6,
     cex.main = 1, las = 1, xaxt = "n",
     xlab = "", ylab = "", main = "")
lines(time, rates_cycads[[length(rates_cycads)]][2,],
      type = "l", lwd = 1, col = "red")
legend(-285, 0.38, legend = c("Speciation rate", "Extinction rate"),
       col = c("blue", "red"), lty = 1, bty = "n", lwd= 1, cex = 0.8)
mtext(text = "Rates (Event / Lineage / Myr)", side = 2, line = 2.5, cex = 0.5)

# Paleodiversity dynamic with deterministic approach
max_root.age <- max(sapply(tree_cycads_posteriors,
                           function(x) max(branching.times(x))))
time <- -c(max_root.age,floor(max_root.age):0)

par(mar = c(4,4,3,4), xpd = T)
plot(time, MCC_diversity,
     bty = "n", axes = F, las = 1,
     main = "C) Diversity through time of Cycadales \n(deterministic approach)",
     cex.main = 1, cex.axis = 0.5, cex.lab = 0.5,
     xlab = "", ylab = "", lwd = 1, type = "n", lty = 1, ylim = c(0,500))

add.gts(thickness = -180, quaternary = T, is.phylo = F, cex = 0.5,
        names = c("", "", "M", "O", "E", "Pa", "UC",
                  "LC", "UJ","MJ", "LJ", "T", "P", "C"),
        time.interval = 10, padj = -2.5, xpd.x = F)

mtext(text = "Time (Myrs)", side = 1, line = 2.5, cex = 0.5, at = 180)

par(new = T, mar = c(4,4,3,4), xpd = T)
plot(time, MCC_diversity, bty = "n", xaxt = "n",
     main = "", las = 1, cex.main = 1, cex.axis = 0.5, cex.lab = 0.5,
     xlab = "", ylab = "", lwd = 1, type = "l", lty = 1, ylim = c(0,500))

mtext(text = "Number of species", side = 2, line = 2.5, cex = 0.6)

polygon(x = -c(length(MCC_diversity):1, rev(length(MCC_diversity):1)),
        y = c(range_div[1,], rev(range_div[2,])),
        col = alpha("red", 0.3), border = alpha("red", 0.3))
legend("topleft", fill =  alpha("red", 0.3), bty = "n",
       legend = "Range of posterior\ndiversities",
       cex = 0.8)
mtext("Time (Myrs)", side = 1, line = 2.5, cex = 0.5)

# Paleodiversity dynamic with probabilistic approach
par(mar = c(4,4,2,4), xpd = T)
plot_prob_dtt(mat = prob_dtt_cycads$backbones[[1]],
              lwd = 1, grain = 0.05, col.mean = "black", bty = "n",
              ylim = c(0, 600), las = 1,
              plot.bound = T, col.bound = "black", lty.bound = 2,
              plot.mean = T, plot.prob = F, axes = F, cex.main = 1,
              xlab = "", ylab = "",
```

```r
             main = "D) Diversity through time of Cycadales \n(probabilistic approach)")

add.gts(thickness = -180, quaternary = T, cex = 0.5, is.phylo = F,
        names = c("", "", "M", "Ol", "E", "Pa", "UC",
                  "LC", "UJ","MJ", "LJ", "T", "P"),
        xpd.x = F, time.interval = 10, padj = -2.5)

par(new = T)
plot_prob_dtt(mat = prob_dtt_cycads$backbones[[1]],
              lwd = 1, grain = 0.05, col.mean = "black", bty = "n",
              ylim = c(0, 600), plot.prob = F,
              xlab = "", ylab = "",
              las = 1, plot.bound = T, col.bound = "black",
              lty.bound = 2, xaxt = "n", cex.axis = 0.6)

par(new = T)
plot_prob_dtt(mat = prob_dtt_cycads$backbones[[1]], lwd = 1, grain = 0.05,
              col.mean = scales::alpha("white", 0), bty = "n",
              ylim = c(0, 100), las = 1, plot.bound = F, plot.mean = T,
              plot.prob = F, axes = F, xlab = "", ylab = "")

axis(side = 4, las = 1, xpd = TRUE, cex.axis = 0.6, line = 0, cex = 0.5)
mtext("Number of species (subclades)", side = 4, line = 2.5, cex = 0.4)
for(i in 1:length(prob_dtt_cycads$subclades)){
  plot_prob_dtt(mat = prob_dtt_cycads$subclades[i][[1]],
                lwd = 1, plot.prob = F, grain = 0.1, add = T,
                col.mean = group_colors[i],
                xlab = "", ylab = "")
}
mtext("Number of species", side = 2, line = 2.5, cex = 0.5)
mtext("Time (Myrs)", side = 1, line = 2.5, cex = 0.5)
legend("top", lty = 2, legend = "Confidence interval at 95%", bty = "n", cex = 0.8)
```

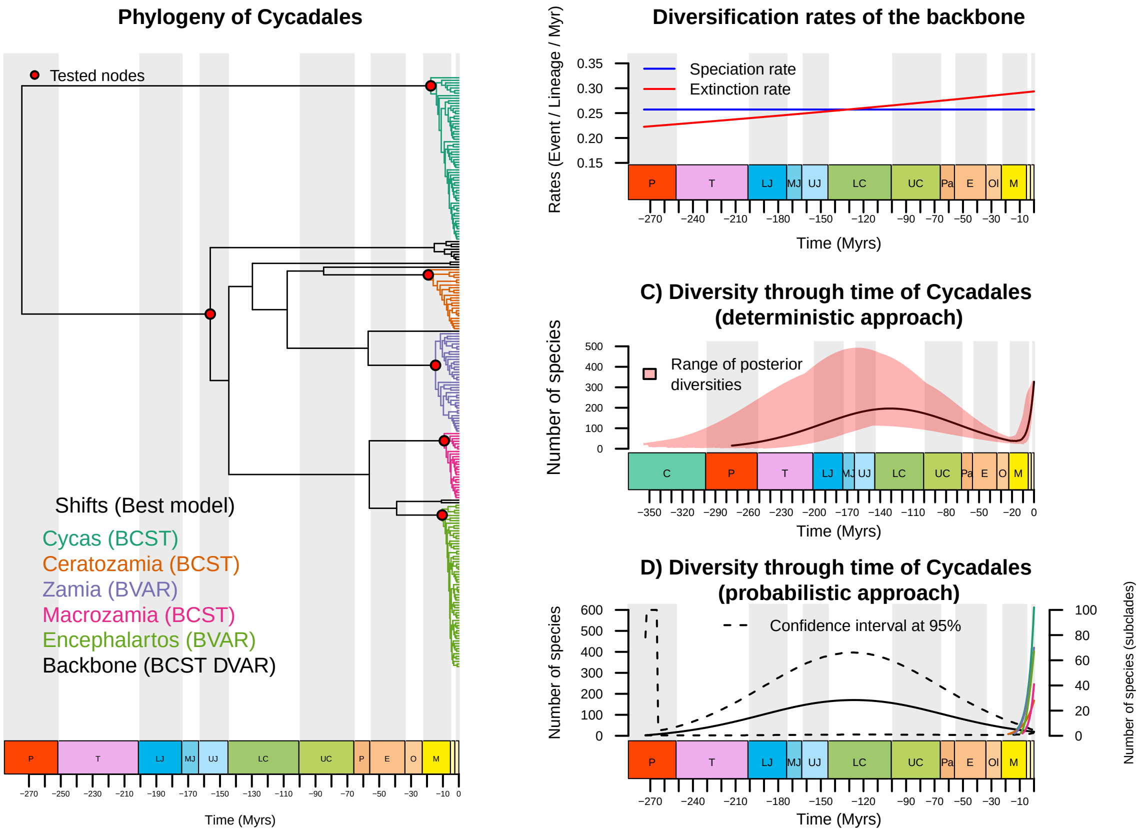

Figure 7: Paleodiversity dynamics of Cycadales

Jonsson, K.A., Fabre, P.-H., Fritz, S.A., Etienne, R.S., Ricklefs, R.E., Jorgensen, T.B., Fjeldsa, J., Rahbek, C., Ericson, P.G.P., Woog, F., Pasquet, E., Irestedt, M., (2012). Ecological and evolutionary determinants for the adaptive radiation of the Madagascan vangas. *Proceedings of the National Academy of Sciences* 109, 6620–6625. https://doi.org/10.1073/pnas.1115835109

Leraut (2016) - Papillons de jour d'Europe et des contrées voisines. 1116 p., N.A.P. Éditions

Medina-Villarreal, A., González-Astorga, J., Espinosa de los Monteros, A., (2019). Evolution of Ceratozamia cycads: A proximate-ultimate approach. Molecular Phylogenetics and Evolution 139, 106530. https://doi.org/10.1016/j.ympev.2019.106530

Morlon, H., Parsons, T.L., Plotkin, J.B., (2011). Reconciling molecular phylogenies with the fossil record. *Proceedings of the National Academy of Sciences of The United States of America* 108, 16327–16332. https://doi.org/10.1073/pnas.1102543108

Nakae, M. (2021) Papilionidae of the World. Roppon-Ashi Entomological Books.

Steeman, M.E., Hebsgaard, M.B., Fordyce, R.E., Ho, S.Y.W., Rabosky, D.L., Nielsen, R., Rahbek, C., Glenner, H., Sørensen, M.V., Willerslev, E., (2009). Radiation of Extant Cetaceans Driven by Restructuring of the Oceans. *Systematic Biology* 58, 573–585. https://doi.org/10.1093/sysbio/syp060

Weiss J.C. 1991–2005. The Parnassiinae of the World. Part I 1991, II 1992, III 1999, IV 2005. Venette: Sciences Nat. p. 400.
