## Appendix S5 for "Estimating clade-specific diversification rates and palaeodiversity dynamics from reconstructed phylogenies"

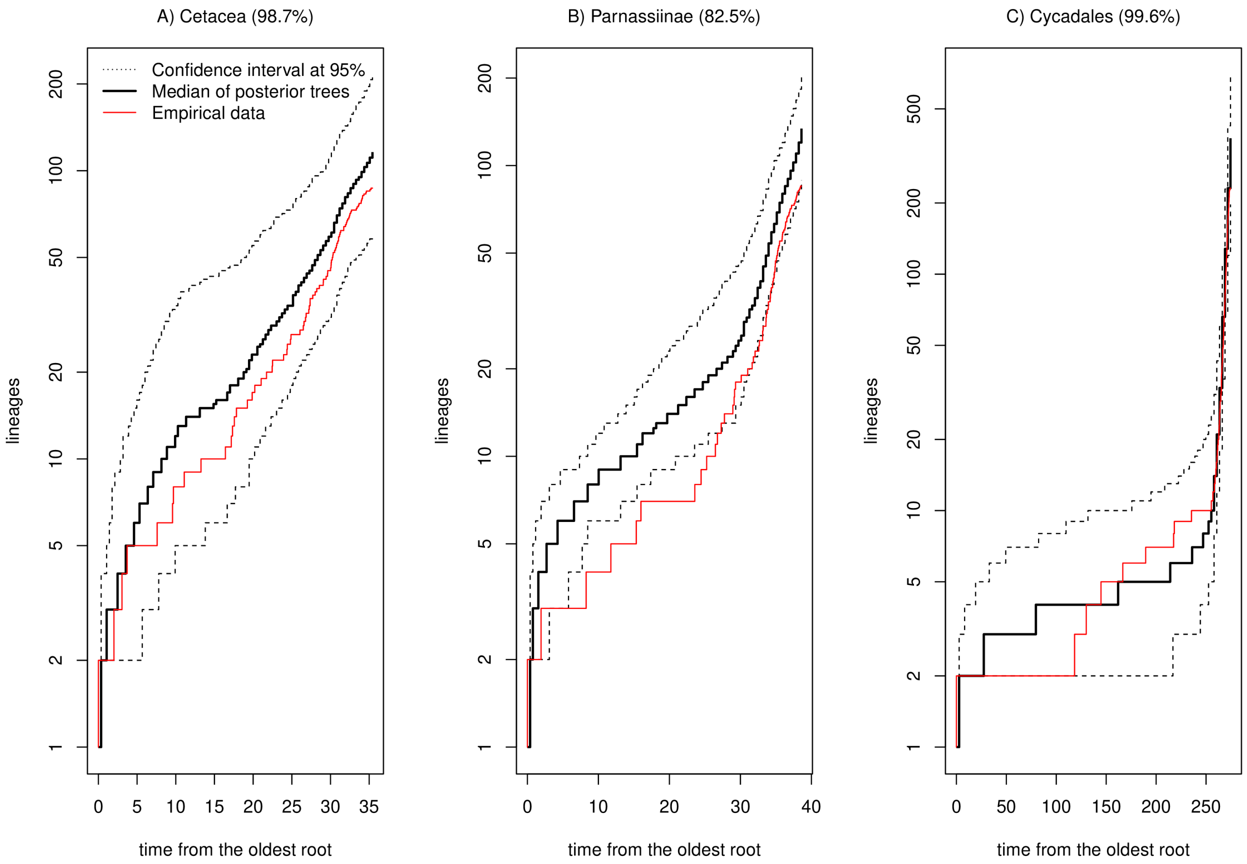


**Figure S5: Lineages-Through-Time plot of Cetacea posterior predictives analyses for A) Cetacea, B) Parnassiinae and C) Cycades.** Dotted lines represent the confidence interval at 95%, solid black line correspond to the median of the poserior trees and solid red line is the empirical LTT. y-axis is log-scaled. We then measure the proportion of values of the empirical Lineages-Through-Time (LTT) that are contained in a 95% confidence interval obtained from the LTT of simulated trees. These values are in parentheses in each title.

We used the function **simul.comb.shift()** to simulate trees under the best combinations of the Cetacea analysis (See appendix S4, III – Testing model adequacy for more details).
